# Nanoliter-scale selection of optimized bioengineered peptide antibiotics that rescue mice with bacterial lung infection

**DOI:** 10.1101/2024.05.30.596569

**Authors:** Nils Böhringer, Zerlina G. Wuisan, Michael Marner, I Dewa Made Kresna, Ute Mettal, Steven Schmitt, Silke Reiter, Yang Liu, Karina Brinkrolf, Oliver Rupp, Oliver Schwengers, Julia Findeisen, Susanne Herold, Ulrich Matt, Till F. Schäberle

**Author notes:** These authors contributed equally (Zerlina G. Wuisan, Michael Marner). **Data and materials availability:** The data that support the findings of this study are available in the supplementary material of this article. **Ethical approval statement:**Animal experiments were conducted according to the legal regulations of the German Animal Welfare Act (Tierschutzgesetz) and approved by the regional authorities of the State of Hesse (Regierungspräsidium Giessen).

## Abstract

Increasing numbers of multi-drug resistant pathogens call for new chemical scaffolds, addressing novel targets, that can serve as lead structures for the development of life-saving drugs. For antibiotics, natural product-inspired molecules represent a most promising resource. Natural products evolved to high chemical complexity and occupy a chemical space different than synthetic libraries. However, clinical translation of promising natural products is often impeded by their relative inaccessibility to medicinal chemistry optimization, *e.g.* iterative synthesis of large series of derivatives. Here, this limitation is addressed with a randomized library of bicyclic heptapeptides based on the natural product darobactin that hits the clinically not addressed target BamA. Variants of the ribosomally synthesized and post-translationally modified peptides were generated using heterologous mutasynthesis. A parallelized screening assay is adapted in nanoliter-scale beads to test the darobactin derivatives against our sensor strain. Loss of fluorescence sorting prioritized 563 events out of the analyzed ∼500k beads. Re-testing confirmed 48 hit events, of which 40 proved to produce distinct darobactin-type molecules. Most promising structures were isolated and the growth inhibitory effects against Gram-negative pathogens validated. One of our current frontrunner compounds (*i.e.*, darobactin B) was reinforced by the randomized screen. While microbiological investigations of the new derivatives is ongoing, darobactin B was profiled in later tier assays and compared to another promising, rationally-designed analog (*i.e.*, darobactin B9, “D22”). Early ADMET profiling and efficacy tests in a mouse pneumonia model were performed. Darobactin B reduced bacterial load of *Pseudomonas aeruginosa* and *Klebsiella pneumoniae* by intraperitoneal, as well as intratracheal administration. Our study showcases the potential of mutasynthetic libraries for high-throughput screening and identification of functional peptides for drug lead discovery.

## INTRODUCTION

The emergence and spread of multi-drug resistant (MDR) pathogens poses a threat to the global health systems. (*1–3*) Resistance transmission renders the ‘magic bullet’ less effective at combating bacterial infections. On a global level 1.27 million deaths were directly attributed to antimicrobial resistance (AMR) and 4.95 million deaths linked to AMR in 2019. (*4*) Among bacterial pathogens, Gram-negatives such as *Pseudomonas aeruginosa*, *Acinetobacter baumannii* and *Klebsiella pneumoniae* are of particular concern. Research and development of new antibiotics against these critical pathogens is named as priority one by the world health organization (WHO). (*5*) Even though the awareness for this silent pandemic is growing, economic incentives are still lacking resulting in a nowadays desiccated antibiotic development pipeline. (*6–9*)

Most antibiotics in clinical use are either microbial natural products (NPs) or semisynthetic derivatives of them. (*10*) Darobactin (DAR) A exemplifies the latest discovery of NPs specifically active against Gram-negative pathogens. (*11*) DAR is a ribosomally and posttranslational modified peptide (RiPP) encoded by a biosynthetic gene cluster (BGC) consisting of the five genes *darA-darE*. These translate into the precursor peptide DarA, an ABC transporter system (DarBCD) and the radical SAM (RaS) enzyme DarE. (*12*) The latter is catalyzing the formation of two fused intramolecular rings in the DAR precursor. The precursor peptide is subsequently trimmed down by proteases, resulting in the rigid β-strand-like bicyclic heptapeptide. (*11, 13*) This enables the interaction of the peptide backbone with the β-barrel assembly machinery (BAM) complex, an essential component of the Gram-negative outer membrane. The transmembrane protein BamA is a so far unaddressed pharmaceutical target and particularly attractive as it is directly accessible from the extracellular space. (*14, 15*)

To engineer functional heterologous expression systems for new DAR-like BamA inhibitors only the precursor peptide and the modifying RaS enzyme are essential. (*13, 16–18*) The small core BGC (*darAE)* make the bicyclic heptapeptidic BamA inhibitors a prime target for mutasynthetic engineering. Certainly, not all theoretically possible 1.28 billion precursor heptamers will result in an active bicyclic molecule. However, despite biosynthetic constraints, thousands of distinct DAR-type compounds could potentially be generated and evaluated by bioactivity screening. Diversity-driven approaches need robust high-throughput technology to be viable for discovery and optimization campaigns. Miniaturized screening systems such as the nanoFleming technology (*19*) can be applied to test the inhibitory activity against a target pathogen in whole cell assays. Fluorescence-labelled test strains, challenged with respective antibiotic producers in such a bead-based system, enable quick and sensitive detection of active molecules.

Here, we demonstrate that cloning a randomized peptide library in combination with a miniaturized high throughput test system can be efficiently used to select the most active variants for further evaluation. First, we tested the substrate flexibility of the modifying RaS enzyme. Informed by these results, we designed a library of 16k variants covering all heptapeptide sequences suitable for macrocyclization by DarE and introduced it into our expression host. In total, about 500k beads were screened for loss of fluorescence and 563 (∼0.1%) recovered from which 48 (∼0.096 ‰) were selected based on their inhibitory activity. After the high-throughput enrichment step, 8.5% of the beads passed the conformational assay representing a high hit-rate compared to <<0.1% in regular screening campaigns. This approach identified new bicyclic heptapeptides active against relevant Gram-negative pathogens *in vitro* and justified *in vivo* lung infection models for the current frontrunner compounds.

## RESULTS

### Generation and screening of a randomized darobactin-type BamA inhibitor library

Using our established platform for heterologous expression of bicyclic heptapeptides (*13*), we first tested whether the ring size of the macrocycles could be expanded (*i.e.*, W**A**NWSKSF, WN**A**WSKSF, WNW**A**SKSF, WNWS**A**KSF), contracted (*i.e.*, WWSKSF, WNWKSF), or if the heptapeptide could be elongated (*i.e.*, **X**WNWSKSF**X**, WNWSKS**WA**) or shortened (*i.e.*, removal of C-terminal amino acids not involved in ring closures). The biosynthetic machinery seemed intolerant to the macrocycle variations, which is in agreement with recent findings using an *in vitro* system. (*20*) Additionally, biosynthesis is also intolerant to N- or C-terminal extension, *i.e*. shorter peptides were not detected while longer peptides were trimmed down to heptamers. Furthermore, a comprehensive one-by-one aminoacid (AA) exchange of the DAR core peptide was performed to test if (i) the macrocycles were formed and (ii) resulting compounds exhibit antibacterial activity (Table S5). We observed that position W^1^, W^3^ and K^5^ are conserved in all bioactive bicyclic heptapeptides. C-terminal (position F^7^) incorporation of hydrophobic and/or aromatic AAs, preferably W or F was proven essential for bioactivity (Fig. S4), even though DarE would accept nearly all proteinogenic AAs at this position. These findings guided the design of the randomized library to access the plenitude of possible bicyclic heptapeptides. Two distinct libraries, *i.e.*, W^1^-X^2^-W^3^-X^4^-K^5^-X^6^-F^7^ and W^1^-X^2^-W^3^-X^4^-K^5^-X^6^-W^7^, were generated by site saturation mutagenesis using NNK-codon degenerated primers. The pooled Master Library contained ∼44.6 bn molecules/µL. Quality control of the library by amplicon sequencing revealed a minimal coverage of 83.96 % (6,675 distinct W^1^-X^2^-W^3^-X^4^-K^5^-X^6^-F^7^ heptapeptides and 6,758 distinct W^1^-X^2^-W^3^-X^4^-K^5^-X^6^-W^7^ heptapeptides; in total the coverage for W^1^-X^2^-W^3^-X^4^-K^5^-X^6^-F/W^7^ is 97.95 %). Out of the 120,683 sequencing reads, 29,677 included W^1^-X^2^-W^3^-X^4^-K^5^-X^6^-F^7^ heptapeptides and 24,886 included W^1^-X^2^-W^3^-X^4^-K^5^-X^6^-W^7^ heptapeptides, which results in a total fidelity of 45.21% at a sampling multiplier of 3.7-fold the library size. (*21*) For heterologous expression of the library, a DAR-resistant *E. coli* host strain was generated. Therefore, point mutations that were reported previously to result in resistant strains, (*11*) were introduced into *bamA* by λ-Red recombineering. Then, the resulting expression host strain *E. coli* Bap1^DarR^ was transformed using this master library.

To overcome the inherent limitation of variable production titers of different derivatives, an ultra-sensitive, fluorescence-labelled reporter *E. coli* MG1655 sensor strain was created and used on our HTS approach (Figure 1). In a BamA mutant strain, (*22, 23*) the gene *bamB* was deleted by λ-Red recombination. This resulted in a >130-fold increase in susceptibility against DAR A with an MIC of <0.03 µg mL^−1^ for *E. coli* MG1655 *ΔbamB::aac*(*3*)IV in comparison to 4-8 µg mL^−1^ for *E. coli* MG1655 wild type (Table S1). The activity of the putative BamA inhibitor molecules was screened in nanoscale bioreactors (*i.e*., alginate-based beads 150 µm in diameter) in which producer and sensor strain were co-encapsulated in a ratio of 1:100 cells per bead (Figure 1). The growth rates of the producer and sensor strains were fine-tuned, to prevent overgrowth and nutrient depletion. Because the producer strains carrying the pRSF-Duet constructs had a longer lag-phase than the strains without the vector (Figure S2), the growth of the sensor strain was restricted by introduction of a riboflavin auxotrophy. Therefore, the *ribC* (encoding the riboflavin synthase) was replaced by the riboflavin transporter *ribM* from *Streptomyces davawensis* by homologous recombination. This causes growth arrest of the sensor cells once riboflavin is depleted, giving the non-auxotroph library producer strains a growth advantage.

**Figure 1.**
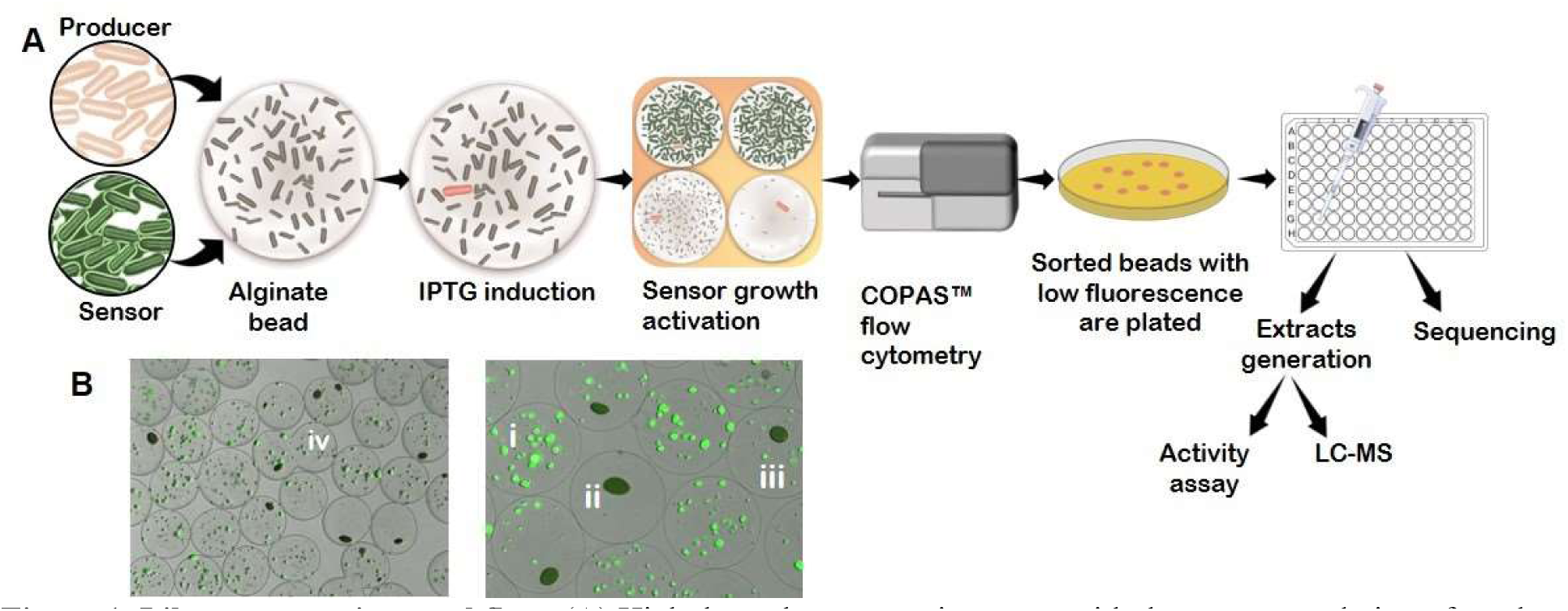
Library screening workflow. (**A**) High throughput screening starts with the co-encapsulation of producer (red) and sensor (green) strains in alginate bead at a 1:100 ratio. After five hours incubation in riboflavin-depleted medium, production of the encoded BamA inhibitor variants is induced by IPTG and incubation is continued for three hours. Then, sensor growth is activated by supplementation of riboflavin. Growth is allowed for 30 minutes, before the growth medium is replaced with paraffin oil and incubated overnight. The beads are then analyzed by COPAS flow-cytometry for colony size and fluorescence level. Low-fluorescent beads are selected and spotted on riboflavin-depleted agar plates (LBKan for selection). Colonies that grow on agar plates are transferred into 96-well plates for extract generation and sequencing. The resulting extracts are used for activity confirmation screening against MG1655 *bamB::aac*(*3*)IV and analysis by UPLC-MS. (**B**) Microscopic pictures of beads. The producer strains that were allowed to have a head start show bigger colonies (dark green) and can be distinguished from the sensor strain (smaller colonies, green fluorescence). The different beads are exemplified: (i) intense fluorescence, with or without producer strain, (ii) only producer strain, no sensor strain (low fluorescence), (iii) producer strain, reduced amount of sensor strain (low fluorescence), (iv) special cases, like merged beads. Representative beads are labeled, white letters.

Following co-encapsulation, the producer was allowed to grow in riboflavin depleted LB medium for 5 h at 30 °C, before DAR production was induced by IPTG addition. After 8 h, sensor strain growth was activated by addition of riboflavin and after 30 min incubation, beads were submerged in paraffin oil to prevent inter-bead-diffusion. Following overnight incubation, beads were collected and remaining paraffin oil was removed using tenside-containing buffer. Next, the beads (∼500k) were analyzed using a COPAS flow-cytometry system for adsorption and fluorescence. One dominant bead population showed high fluorescence, and a smaller subpopulation (∼1.5 %) showed low fluorescence, indicating inhibition of the sensor strain. Low-fluorescent beads (n=7668) were individually spotted to LB_Kan_ agar plates. That way, 563 colonies were recovered, and plasmid sequencing allowed quick dereplication of 380 distinct monoclonal encapsulated producer colonies (112 with ≥2 different clones, 71 did not give any sequencing result). At the same time, these clones were cultivated for arrayed mL-scale production of the respective DAR derivative. Whole culture broths were quenched with MeOH and concentrated 1:20 in H_2_O. Crude extracts were analyzed by UPLC-MS to link *darA* sequencing with analytical results. Additionally, micro-broth dilution assays were performed to evaluate the degree of anti *E. coli* activity. Out of the 563 selected clones, 48 extracts inhibited the sensitive test strain *E. coli bamB::aac*(*3*)IV; thereof, 40 contained a single DAR-type BamA inhibitor sequence (Figure 2D). Within these 48 HITs, some sequences were represented more than once, resulting in identification of 30 distinct HIT molecules.

**Figure 2:**
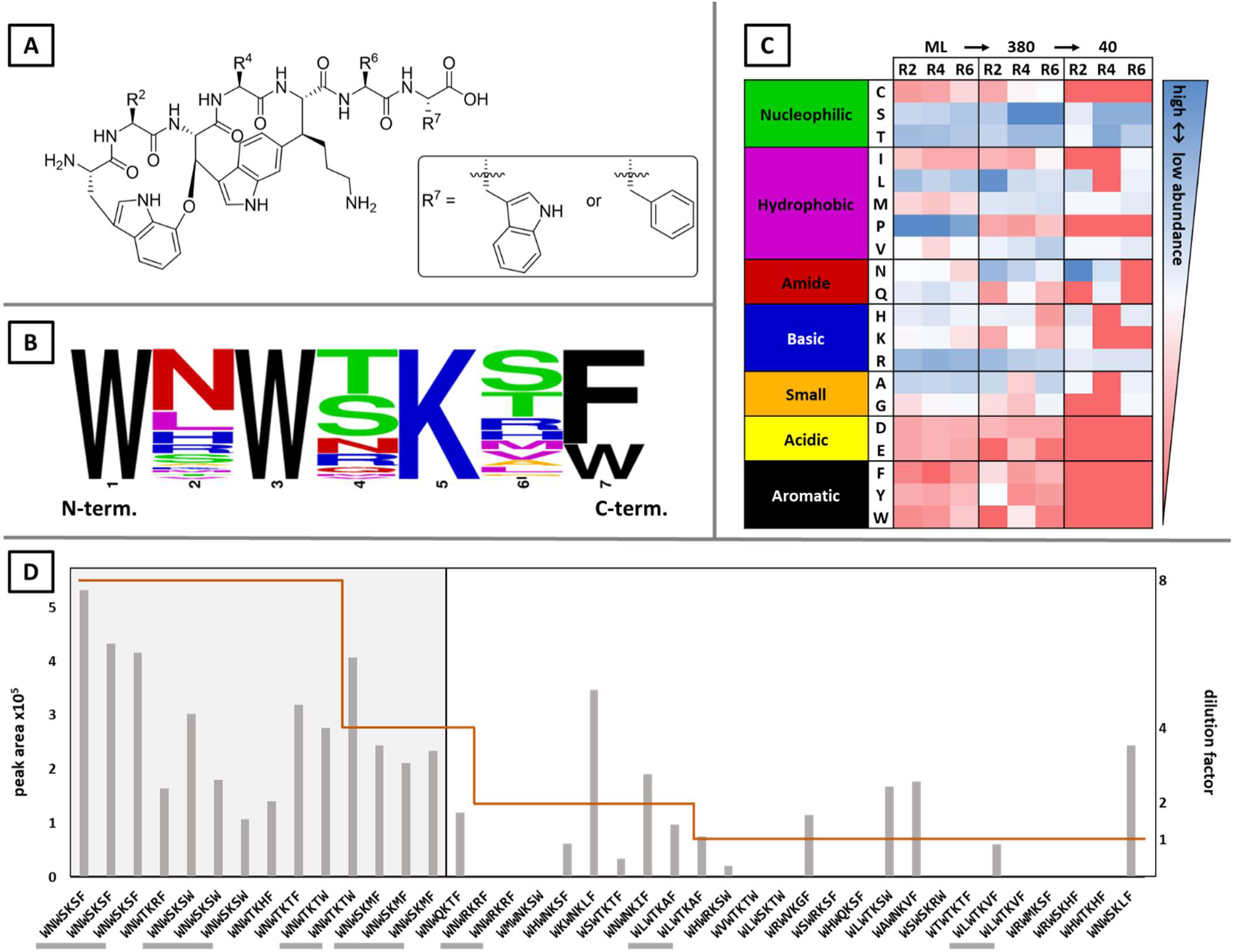
**A)** Chemical structure of the scaffold, randomized amino acid positions are indicated by R2, R4, R6, and R**7** as either F or W. **B**) Sequence logo of amino acid abundance in the enriched beads that were selected based on activity screening (weblogo.berkeley.edu). **C**) Heat map of amino acid distribution, the residues are categorized based on their chemical properties; shown are the randomized residues at position 2, 4 and 6 (R2, R4, R6) of the master library (ML), the 380 enriched monoclonal beads and the 40 verified hits strain (30 compounds). **D**) Ion abundance ([peak area], grey columns) and activity score ([dilution factor], brown line) of verified hits. Sequences that were detected more than once are highlighted (vertical bars below the amino acid sequence). Top scoring hits that were selected for production and MIC determination are marked by the grey box.

The comparison of the relative abundance of specific AAs at a particular position in the master library and the verified HITs revealed selective enrichment of a small number of AAs per randomized position (Figure 2B-C). Position 2 displayed 11-fold enrichment of N in the HITs, while A, H, K, L, M, R, S, T and V were observed in lower abundances. Position 4 showed less variability in the incorporated AAs and highest enrichment was observed for S and T, with a 4.8- and 4.4-fold increase in relative abundance, respectively. Further AAs observed at this position within the HIT-group were N, R, M, Q and V (relative abundance 2.8, 1.1, 0.7, 0.8 and 0.7-fold). In line with that, S and T were observed to be the most favorable residues at this position: they were present in two third (67.5%) of HITs and all HIT-derived crude extracts that inhibited the test strain down to the 8-fold dilution step carried an S^4^ or T^4^ (Figure 2D). For position 6, a similar variability as for position 2 could be observed; however, instead of N, we could see an enrichment of S and T with a 4.3- and 2.7-fold increase in relative abundance. The residues M, V, H and R were also enriched (2, 1.7, 1.5- and 1.2-fold), while the abundance of I, A, L and G decreased (0.8, 0.7, 0.7 and 0.6-fold).

In the top ten HITs that inhibited *E. coli* up to an 8-fold dilution, the natural variant DAR A (WNWSKSF) and WNWSKSW (DAR 9) (*17*) were both recovered three times. For these two, the only difference is in position 7 that was predetermined to be either F or W by the library design. Furthermore, WNWTKRF (DAR B), WNWTKHF, WNWTKTF, and WNWTKTW (D38, (*18*)) are top ten variants. The observed relative abundance of F^7^:W^7^ in the master library was 52.5:47.5, which is in good accordance with the envisaged equal distribution of the different residues. Within the HITs, abundance of W^7^ was reduced to 29%; however, within the top 10 hits 40% of the clones carried W^7^.

To evaluate the results of the HIT confirmation assays against *E. coli*, the compounds causing inhibition down to 8-fold dilution were short-listed for fermentative production and purification. In addition to the heptapeptides showing inhibition at the highest dilution tested (WNWTKHF, WNWTKTF and WNWTKTW (D38), WNWSKMF (inhibition at 4-fold dilution step) was selected, since it was recovered 3 times in the screening assay. MICs were determined against a panel of clinically relevant Gram-negative pathogens. Overall, the purified DAR variants showed activity against the test panel. Variations in the MIC against specific strains were observed (Table 1). The compounds WNWTKTF and WNWTKTW (D38) that are structurally very similar to DAR A and DAR D9, also showed similar antimicrobial behavior. MICs against *E. coli* strains were 4-16 µg mL^−1^ and *P. aeruginosa* PA01 was inhibited at 1-2 µg mL^−1^. However, the growth inhibitory activity was less pronounced against other *P. aeruginosa* isolates, such as the cystic fibrosis isolate EXT111762. (*24*) The activity of these F^7^ or W^7^ variants against these *P. aeruginosa* isolates was similar, *i.e.* identical or only one dilution step difference (Table 1, Table S3). In contrast, WNWTKHF exhibited improved activity against *E. coli* and *K. pneumoniae* test strains as well as *P. aeruginosa* PA01 (MICs of 0.5 - 2 µg mL^−1^). While these results are comparable to DAR B, the MICs against the other *P. aeruginosa* and *A. baumannii* isolates indicate a weaker performance (again compared to DAR B). For WNWSKMF, spontaneous oxidation to the corresponding methionine-sulfoxide was observed, reducing compound stability. Based on these MIC data, DAR B was selected as the current frontrunner to undergo further investigation. Its already established accessibility by fermentation enabled the following detailed investigation against *P. aeruginosa* and *K. pneumoniae*. Furthermore, DAR B9 (=D22) was selected to be tested in parallel. Even though it was not detected in the HTS performed here, it shows high structural similarity and was reported from rationale-based DAR engineering approaches as a potent DAR derivative. (*16, 17, 24*)

**Table 1.**
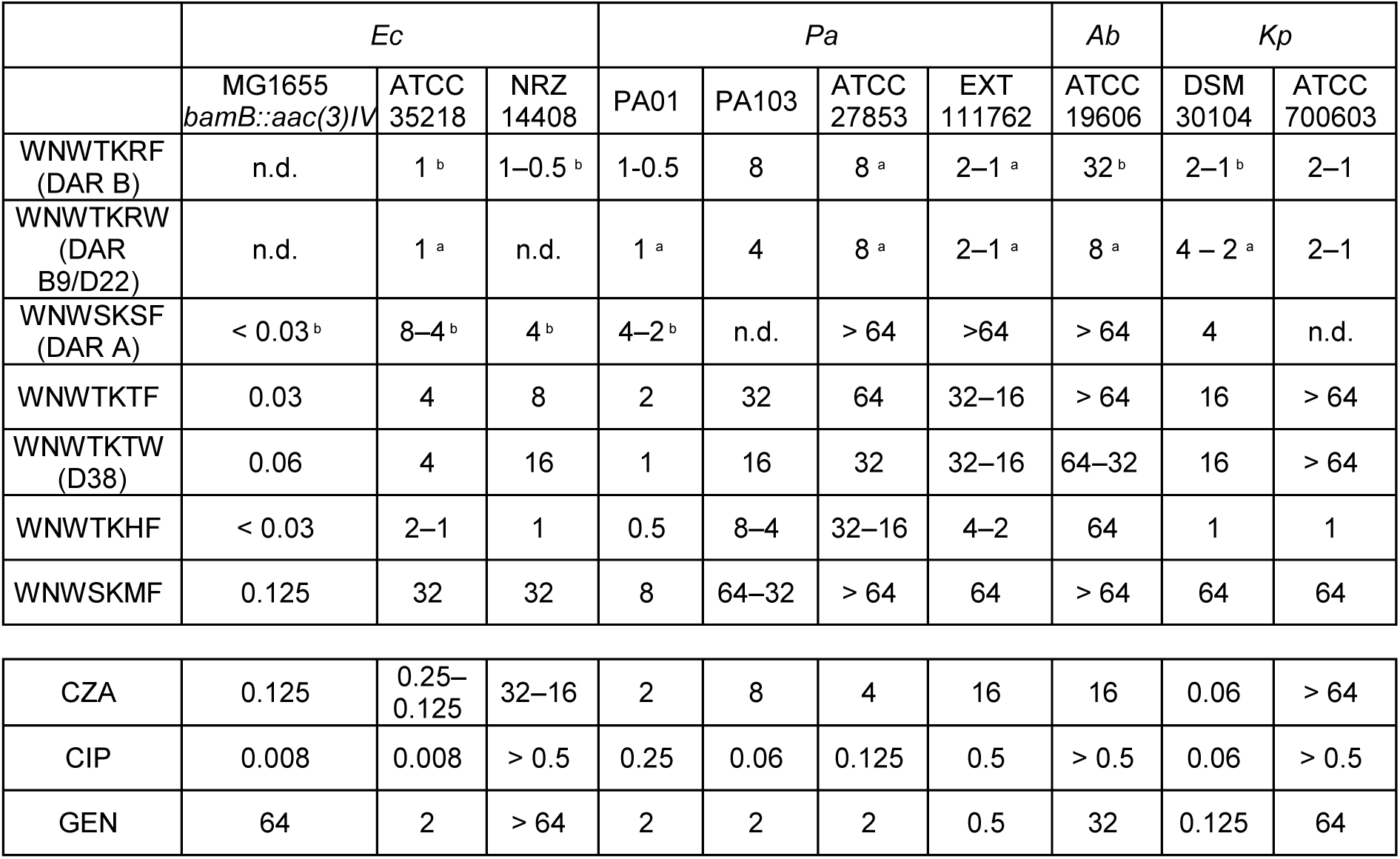
Minimum inhibitory concentration of selected darobactin variants. and reference compounds dDarobactin A (DAR A), darobactin B (DAR B), darobactin B9 (DAR B9, D22) and standard antibiotics ceftazidime (CZA), ciprofloxacin (CIP) and gentamicin (GEN) were tested against *Escherichia coli* (Ec), *Pseudomonas aeruginosa* (Pa), *Acinetobacter baumannii* (Ab) and *Klebsiella pneumoniae* (Kp) isolates. Each MIC determination was carried out in triplicate. Values are given in µg mL^−1^.

### Time-kill-curve of darobactin B

Time-kill-curves were recorded against quality control strain *P. aeruginosa* ATCC27853 and the 4MRGN cystic fibrosis isolate EXT111762. Before MIC determination and time-kill-curve generation, we characterized the isolate EXT111762 by antimicrobial susceptibility testing and genome sequencing. Strikingly, AMRfinder and CART annotated 59 genes related to antibiotic resistance including 3-component signal transduction systems (cprS/cprR, parS/parR, basS/basR), (*25–27*) RND-type multidrug efflux systems (*28, 29*) and drug specific genes like *fosA*, (*30*) oxa-904 carbapenemase, pdc-3 cephalosporinase, (*31*) catB7 chloramphenicol acetyltransferase, (*32*) gyrA (quinolone resistant gyrase) (*33*) and lipid A-modifying arnA. (*34*) (*genome sequencing data GENBANK ID will be released on acceptance*). For both strains the previously determined MICs (Table 1) were used to investigate the killing dynamics between DAR B and *Pseudomonas* (the number of colony-forming units (CFUs) was determined at seven time points in triplicates).

Most importantly, the results (Figure 3) indicate a 3-log reduction of CFUs per mL after 8h for both investigated *P. aeruginosa* strains if DAR B is administered at 4x or 8x of the respective MIC (light and dark green lines in Figure 3). After an initial decline, the data series of ATCC27583 treated with 0.5x MIC ascends identical to the growth control (no DAR supplementation, black line in Figure 3). The effect of 1x MIC treatment on *Pseudomonas* survival was more pronounced. For both strains, the determined CFU concentration followed a steep decline similar to the higher DAR treatments. However, different from the 4x and 8x cultures, the 1x MIC supplemented cultures recovered and reached the upper limit of CFU detection by the end of the experiment. As the growth control of the clinical isolate EXT111762 needed 24h to reach the upper limit of detection (LOD), we decided to monitor the experiment for 48h. Remarkably, even during this elongated time series only single CFUs (ATCC27853) or none at all (EXT111762) were observed in the *Pseudomonas* cultures treated with 4x and 8x the respective MIC. Hence, the observed minimum bactericidal concentrations (MBC) for DAR B were ∼32 µg mL^−1^ (ATCC27853) and 8 µg mL^−1^ for the clinical cystic fibrosis isolate EXT111762.

**Figure 3.**
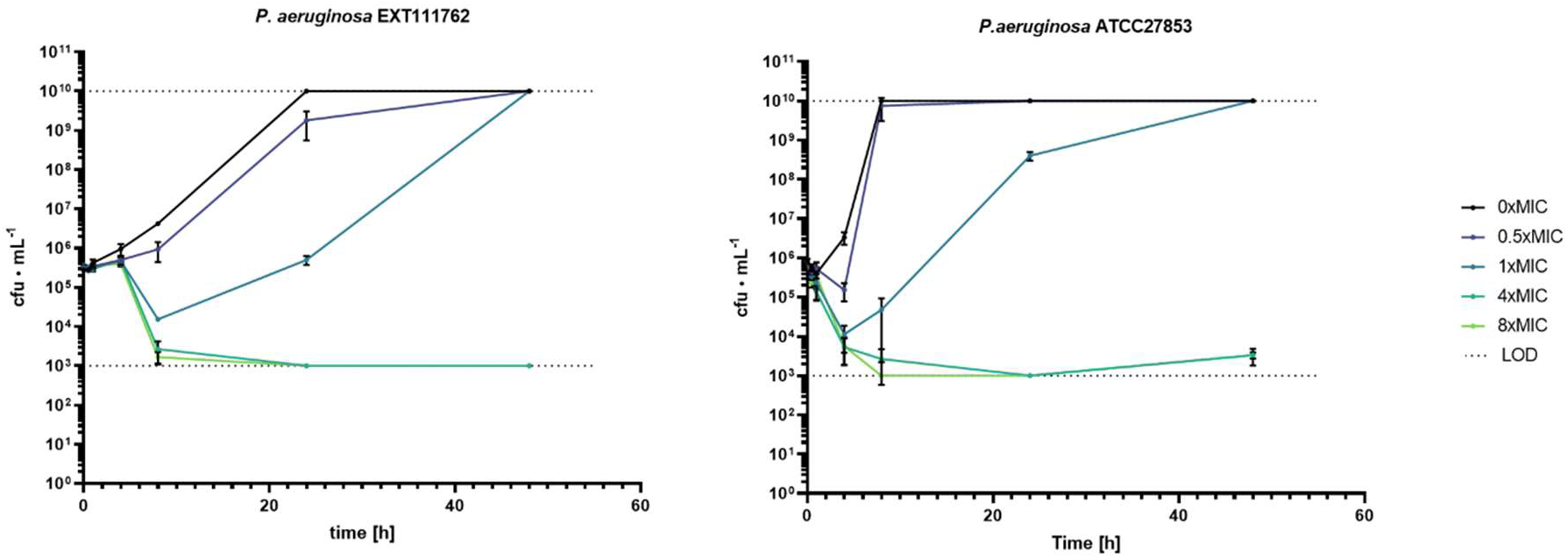
Time-kill curves of darobactin B against *Pseudomonas aeruginosa*. EXT111762 (left) and ATCC27853 (right). The compound was supplemented at 4 different concentrations (0.5x, 1x, 4x and 8x MIC) to the cultures (each treatment n=3) and the number colony forming units (CFUs) was determined at seven time points (0h, 0.5h, 1h, 4h, 8h, 24h and 48h). LOD: Limit of detection.

### Early ADMET of darobactin B and B9 (D22)

Further characterization towards physicochemical and ADMET (absorption, distribution, metabolism, excretion, and toxicity) properties were carried out for the current frontrunner compounds. The hydrophilicity of DAR B and DAR B9 (D22) was confirmed by the determination of its distribution coefficient at pH 7.4 (*logD_7.4_* = −0.39 and 0.20, respectively), indicating higher hydrophilicity compared to the reference compound rifampicin (*logD_7.4_* = −1.12). DAR B and B9 (D22) showed very low permeability in Caco-2 cells, from apical to basolateral and basolateral to apical. To predict the possibility of metabolism-mediated drug-drug interaction, CYP inhibition was studied. CYP2D6 and CYP3A were tested as members of the cytochrome P450 superfamily that catalyze the majority of drug metabolism in the liver. (*35*) We observed low CYP inhibition (−1.7-17.2%) in response to DAR B and DAR B9 (10 µM) exposure (Table 2). (*35*) We observed low CYP inhibition (−1.7-17.2%) in response to DAR B and DAR B9 (D22) (10 µM) exposure (Table 2). For the highest validated test concentration of 100 µM DAR B and DAR B9 (D22), HepG2 cell viability was only reduced by 12.2% and 12.9%, respectively. (*35*) We observed low CYP inhibition (−1.7-17.2%) in response to DAR B and DAR B9 (10 µM) exposure (Table 2). For the highest validated test concentration of 100 µM DAR B and DAR B9, HepG2 cell viability was only reduced by 12.2% and 12.9%, respectively. Therefore, no IC_50/90_ values could be calculated. The inhibition of potassium ion channel encoded by hERG (the human Ether-à-go-go-Related Gene) was determined to predict the risk of cardiac arrhythmia. Electrophysiological assays showed low inhibition of the ion channel by DAR B and DAR B9 (D22) (30 µM), representing a non-significant effect (Table 2).

**Table 2.** Early-ADMET data for darobactin B and B9 (D22)

| Assay | Test concentration | Darobactin B | Darobactin B9 (D22) | Reference |  |
| --- | --- | --- | --- | --- | --- |
| **Distribution coefficient (log D)** | 100 μM | -0.39 | 0.20 | Rifampicin | 1.12 |
| **In vitro absorption (Caco-2)** |  |  |  |  |  |
| A-B permeability ($10^{-6}$ cm s<sup>-1</sup>) | 10 μM | <0.095 | <0.286 | Propranolol | 38.0 |
| B-A permeability ($10^{-6}$ cm s<sup>-1</sup>) | 10 μM | <0.016 | <0.042 | Propranolol | 17.2 |
| **CYP inhibition (substrate)** |  |  |  |  | **IC<sub>50</sub> (μM)** |
| CYP2D6 (dextromethorphan) | 10 μM | 17.2% | 15% | Quinidine | 0.33 |
| CYP3A (midazolam) | 10 μM | 16.7% | 9.2% | Ketoconazole | 0.26 |
| CYP3A (testosterone) | 10 μM | 13.4% | -1.7% | Ketoconazole | 0.19 |
| **Cell growth inhibition (HepG2)** | 100 μM | 12.2% | 12.8% | Staurosporine | 0.45 |
| **Ion channel inhibition (hERG)** | 30 μM | 7.34% | 8.71% | Verapamil | 0.235 |

### *In vivo* efficacy of darobactin B in a Gram-negative pneumonia mouse model

In a next step, DAR B and B9 were tested in mouse lung infection models. We used pneumonia models with *K. pneumoniae* and *P. aeruginosa*, which represent two of the most important bacterial lung pathogens due to their pathogenicity and increasing antibiotic resistance. (*5*) After infection, mice were treated systemically with DAR B, B9 (D22) or carrier, and the bacterial outgrowth was determined in whole lung homogenates 24 hours after infection. DAR B clearly reduced the bacterial load in the lungs in both, *P. aeruginosa* and *K. pneumoniae* pneumonia experiments while DAR B9 (D22) only reduced the *P. aeruginosa*, and not *K. pneumoniae* bacterial load in our study (Fig. 4). In addition, *in vivo* efficacy against the MDR *P. aeruginosa* EXT111762 was observed (Fig. 4C). Next, we tested if the compounds would also be suitable for inhalational administration. Such topical administration is primarily performed in cystic fibrosis patients or as an add-on in patients with pneumonia (*36*) and was recently reported as a means to prevent ventilator associated pneumonia. (*37*) Again, we induced pneumonia with *K. pneumoniae* or *P. aeruginosa*, but this time administered DAR B and B9 intratracheally 2 hours after instillation of bacteria. Subsequently, bacterial outgrowth was determined 16 hours after infection in whole lung homogenates. Here, both compounds clearly reduced bacterial outgrowth in *K. pneumoniae* and *P. aeruginosa* pneumonia (Fig. 4D and E). Thus, inhalation might become an add-on treatment option beside systemic administration for the new drug to be developed. (*36*) and was recently reported as a means to prevent ventilator associated pneumonia. (*37*) Again, we induced pneumonia with *K. pneumoniae* or *P. aeruginosa*, but this time administered DAR B and B9 (D22) intratracheally 2 hours after instillation of bacteria. Subsequently, bacterial outgrowth was determined 16 hours after infection in whole lung homogenates. Here, both compounds clearly reduced bacterial outgrowth in *K. pneumoniae* and *P. aeruginosa* pneumonia (Fig. 4D and E). Thus, inhalation might become an add-on treatment option beside systemic administration for the new drug to be developed. (*36*) and was recently reported as a means to prevent ventilator associated pneumonia. (*37*) Again, we induced pneumonia with *K. pneumoniae* or *P. aeruginosa*, but this time administered DAR B and B9 intratracheally 2 hours after instillation of bacteria. Subsequently, bacterial outgrowth was determined 16 hours after infection in whole lung homogenates. Here, both compounds clearly reduced bacterial outgrowth in *K. pneumoniae* and *P. aeruginosa* pneumonia (Fig. 4D and E). Thus, inhalation might become an add-on treatment option beside systemic administration for the new drug to be developed.

**Figure 4:**
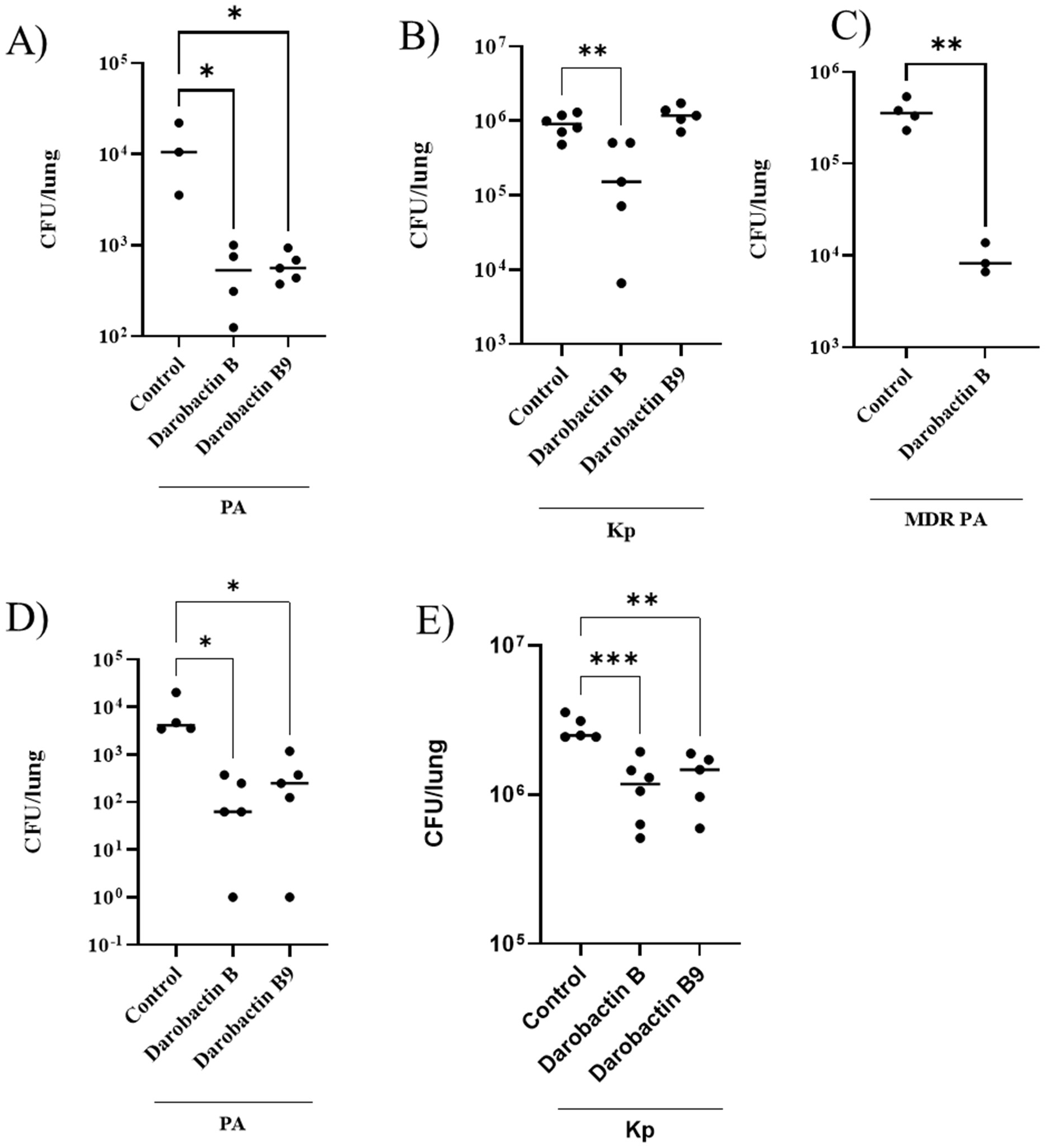
Bacterial outgrowth in *P. aeruginosa* (PA) and *K. pneumoniae* (Kp) pneumonia after systemic or topical administration of darobactins. Mice were intranasally infected with *P. aeruginosa* PA103 (**A**) or *K. pneumoniae* (**B**); 2, 8 and 14 hours after infection darobactin B and B9 (D22) (25 mg/kg) or carrier (0.9% NaCl) were injected intraperitoneally. 24 hours after infection mice were euthanized and bacterial outgrowth in lung homogenates was determined. The experiment was repeated with MDR PA EXT111762 for DAR B (**C**). Similar studies were performed with *P. aeruginosa* PA103 (**D**) and *K. pneumoniae* (**E**) with darobactin B and B9 (D22) (50 mg/kg) administered intratracheally 2 hours after infection. Bacterial outgrowth was determined 16 hours after infection. Two experiments were pooled (n=3-6). Statistical analysis was performed by one-way ANOVA, *p<0.05, **p<0.01, ***p<0.001.

## DISCUSSION

Drug development is a time- and cost-consuming task. HIT molecules discovered in a screening campaign are profiled and optimized to reach the desired properties, e.g. higher efficacy, low toxicity and suitable pharmacokinetics/pharmacodynamics. One way to improve limitations in one or more of these fields is the iterative screening of synthetic derivative libraries. By incorporating the knowledge gained from the previous into the next optimization cycle the sweet spot satisfying the demands is identified.

In this context, antibiotics have to be regarded as a special case. That is because many antibiotic drug candidates, like the darobactins, are complex microbial metabolites, which are, in most cases, difficult to access by synthetic chemistry. Hence, the optimization cycle of medicinal chemistry is often extremely challenging. Although proof of principle for chemical synthesis of the bicyclic peptides was recently achieved (*38, 39*), fermentative access seems to be the more viable solution currently. It is fundamental to identify the best possible starting candidate (Lead) before embarking on a resource-intensive optimization study. Larger scale fermentative production might be a solution for already optimized compounds, but for initially broad SAR-studies a small scale high-throughput production and screening approach would be favorable.

In this study, we aimed to screen a randomized library of ∼16k derivatives that might contribute to faster project progress in Hit-to-Lead initiatives. In an HTS whole cell inhibition approach, we used nanoscale bioreactor capsules (=beads) to screen for anti-Gram-negative activity. We used *E. coli* as a first line indicator strain, but the general strategy to employ growth deficient sensor strains that can be activated ‘on demand’ can be implemented for other pathogens of interest.

In our encapsulation approach, a single cell represents the inoculum of the heterologous producer of target derivative structures, while an auxotrophic, ultra-sensitive and fluorescence-labelled *E. coli* sensor strain was engineered for detection of antibiotic activity. Within the alginate beads, the conditions for microbial growth differ considerably from classical flask or micro-titer-plates. In particular, the availability of nutrients is limited. Hence, encapsulation of producer- and reporter-cells required delicate balancing of inocula to minimize trivial overgrowing effects. This proved to be challenging, since producer cells, even when harboring only an empty vector control, showed longer lag phases upon induction with IPTG. To counteract, a growth deficient sensor strain was used and enabled successful fine-tuning of growth rates. We allowed the single producer cells a head start to form micro-colonies in their beads, before the sensor strain was activated by riboflavin supplementation. To reduce the probability of undesired encapsulation events (*e.g*., more than one producer strain into one bead) the λ has to be in the range of λ=0.3 according to Poissant distribution. (*40, 41*) In turn, many beads might not have carried a producer cell. Yet, co-encapsulation of two different producer clones cannot be excluded completely, but if such a bead is selected based on bioactivity, DNA sequencing identifies these rare events. In this study 67.5% of enriched beads contained single clones.

We selected the natural product darobactin A as template for library generation. DAR A was the first small molecule inhibitor of the Gram-negative specific molecular target BamA. (*11*) Since the discovery of this bacterial metabolite a few years ago, targeted derivatization approaches were undertaken, resulting in several molecules with binding affinity to the β-barrel structure of BamA. (*16, 18*) Additionally, a genome mining-based search for related RiPP structures that have the potential to inhibit the outer membrane protein BamA was sparked. (*12, 16, 42, 43*) Of these, the bicyclic heptameric DAR-like structures still represent the most promising class for antibiotic development. Because the minimal BGC only consists of a precursor and a radical SAM enzyme encoding gene, heterologous expression is feasible and sets the basis for biotechnological optimization (*i.e.*, by codon exchange of the precursor gene to introduce other proteinogenic AAs). Hence, DAR derivatives have been generated to-date, thereunder compounds with stronger activity and broader activity spectrum than DAR A. (*16–18, 44*) Hypothesis-driven bioengineering of the core peptide sequence helped formulate first SAR conclusions for BamA interacting with bicyclic heptapetides. The side chains of AAs, which are not directly involved in the DarE mediated macrocyclization, are promising moieties for optimization efforts. (*16–18*) Sophisticated *in silico* design and selective *in vitro* bioactivity evaluation comprehensively summarized the impact of R2, R4 and R6 modifications as well as interacting effects among them on BamA bonding. (*45*)

However, rational engineering poses the risk of overlooking non-obvious beneficial modifications or combinations. Due to the exponential nature of combinatorial events in randomized peptides (20^X^), already 3 randomized positions translate into 8000 different molecules. To leverage this numbers game into a manageable format, a high-throughput biosynthesis and screening approach was carried out to enrich active variants from a randomized library. The enrichment of HITs from the master library clearly reduced sequence diversity, which on the one hand is guided by DarÉs substrate tolerance, but on the other hand, allows deduction of certain SAR rules by activity screening of the derivatives produced. In the master library, proline was the most abundant AA for all positions, while it was fully eliminated in the validated HITs. This finding reflects the sequence breaking properties of P, which prevent formation of the desired bicyclic heptapeptide, already observed in the single AA exchange study (Figure S5) (*46*). Due to the high-throughput nature of the herein described approach, we could afford to additionally keep position 7 flexible (W or F) and test all daropeptides in a whole cell assay. In fact, almost equal distribution in the master library (*i.e.*, 47.5% W) and in the most active validated HITs (40% W) was observed, indicating F^7^ and W^7^ substitutions perform similar against *E. coli* in our test system. Previous reports showed that for *A. baumannii* activity, W^7^ seems to perform better than F^7^. This is most prominently observed in DAR B9 (D22) compared to the native DAR A or DAR B and was also found in the data presented here: While the newly isolated WNWTKTF shows no activity against *A. baumannii*, WNWTKTW (D38) shows moderate activity in our hand 32-64 µg mL^−1^ and studies done by Seyfert *et al*. (4-64 µg mL^−1^). (*18*) However, for other critical pathogens, such as *P. aeruginosa,* no striking difference between W^7^ and F^7^ was observed. (*24*) For some *E. coli* and *K. pneumoniae* strains F^7^ seems to perform even slightly better than W^7^ (see also Table S3).

It was shown that R^6^ > H^6^ ≈ K^6^ > S^6^ > T^6^ against *A. baumannii* if tryptophan is incorporated at the C-terminus of the heptapeptide. (*18*) Our findings partly agree with this data: by comparing DAR B9 (D22) and WNWTKTW (D38) we saw a 4-16-fold increased MIC against the tested pathogens as a consequence of R^6^/T^6^ exchange. In derivatives with F^7^, we saw the effects less clearly. For *A. baumannii* ATCC19606, arginine (DAR B) replacement by histidine at position 6 (newly isolated WNWTKHF) reduces the activity by only one dilution step. For activity against *E. coli* and *P. aeruginosa,* R^6^ might be better (MIC shift up to 4-fold), while for *K. pneumoniae* the opposite could be observed in our experiments. Similarly, the MICs against *P. aeruginosa* EXT111762 were >4 times lower in the newly isolated WNWTKTF compared to the native DAR A featuring S^6^ instead of T^6^. Taking together, our data indicate that the performance of the AAs at position 6 is influenced by the respective AA side chain at the C-terminus, hinting towards a synergism of position 6 and 7 sidechains towards *A. baumannii* activity. If F^7^, we could observe that R^6^ ≈ H^6^ > T^6^ ≥ S^6^. Furthermore, in the screening, (i) alipathic-hydrophobic sidechains were recovered; however, exhibited low activity against the utilized test strain, and aromatic and acidic AAs were completely eliminated during HTS (Fig. 2C), indicating that incorporation of those is detrimental to activity.

At position 2, asparagine (native to DAR A and B) showed strongest enrichment (11-fold) of all AA residues. While active derivatives with K^2^, H^2,^ and M^2^ were recovered in the HTS, activities appeared to be sub-par to derivatives carrying N^2^ and hence were not selected for MIC determination (Figure 2D). This is in line with literature reports of rational-engineering-based DAR optimization, where most potent molecules exclusively carry N^2^. (*16–18, 45*) In fact, position 2 in the DAR scaffold was subject to extensive rational engineering attempts. For example, DAR derivatives exhibiting Q^2^, which extends the sidechain by one CH_2_ unit while keeping its amidic nature and the other substituents fixed, resulted in slightly improved binding affinity against purified BamA_Ec_; however, activity against all tested strains was reduced by at least one potency in a whole-cell antibacterial activity assay. Thereby, underpinning the importance of whole-cell-testing for accurate SAR interrogation. (*45*)

The insights gained about SAR of bicyclic heptapeptides, together with lessons learned from former peptide-based antibiotic candidates can contribute to future development projects. For example, drugs that have targets in the outer membrane will usually benefit from an overall positive charge. However, the toxicity has to be monitored. The clinical trials of the promising and highly active candidate murepavadin, which is the first example of an outer membrane protein (LptD)-targeting antibiotic class (MIC_90_ of 0.12 µg mL^−1^ against *P. aeruginosa* strains), had to be halted in clinical phase III, due to higher-than-expected acute kidney injuries. (*47*)

Up to now the BamA inhibitors are in preclinics, and to evaluate the potential of the two current frontrunner molecules DAR B and B9, ADMET data were generated to evaluate their potential for antibiotic drug development, before *in vivo* efficacy in murine models was investigated. We followed the recommendations of the drug regulatory agencies in the U.S. (FDA) and Europe (EMA) and tested cyto- and cardiotoxicity, respectively. Very low HepG2 cytotoxicity (IC values out of range) and no hERG ion channel inhibition was observed. The cytochrome P450 (CYP) enzyme family is considered the main route of drug metabolism in humans and inhibition can lead to undesired drug-drug interaction (DDI) and adverse effects. (*35*) Intriguingly, the inhibition assays of two major CYPs with different substrates showed no potential for DDI. Yet, further CYP inhibition and induction assays should be performed in the future to identify enzymes that metabolize DAR as well as identifying the DAR metabolites and their activity. (*48*) The unformulated compounds are hydrophilic and highly soluble in water. Therefore, the compounds are bidirectionally non-permeable in Caco-2 cells, indicating negligible intestinal absorption, rendering oral drug administration currently non-feasible. (*49*) Accordingly, efficacy of two different routes of compound administration for the treatment of *P. aeruginosa* and *K. pneumoniae* mediated pneumonia were evaluated.

In a preclinical model of lung infection, mice were infected with either of the pathogens intranasally. Due to the pathogenicity of the *P. aeruginosa* strain, the initial bacterial load must be limited to not kill the mice. However, a reduction of >log1 was observed for both compounds (in comparison to carrier control, Figure 4). Against *K. pneumoniae* the reduction of CFUs in lung was also significant for all experiments, except that DAR B9 failed to reduce bacterial outgrowth if administered intraperitoneally. Furthermore, the MDR *P. aeruginosa* strain EXT111762 was tested and DAR B significantly reduced CFUs in lung tissue, emphasizing the potential of the compound class against pathogens that show resistance against clinically applied antibiotics.

It must be kept in mind that the treatment of lung infections caused by MDR Gram-negative bacteria is most challenging and will require new approaches. Currently, guidelines do not recommend adjunctive treatment with inhaled antibiotics for MDR bacteria in standard care of pneumonia. (*50*) However, in select cases where systemic therapy fails, adjunct topical antibiotic therapy is an option and was shown to result in improved outcomes. (*51, 52*) Moreover, a recent study demonstrated a reduced rate of ventilator-associated pneumonia after inhaled antibiotic therapy in patients ventilated for at least 3 days, (*37*) which might pave the way for routinely administered topical antibiotics in intensive care units. In patients with cystic fibrosis (CF), inhaled antibiotic treatment is already routinely applied. (*53*) Often, limited treatment options in chronic *P. aeruginosa* infections in CF patients force clinicians to apply alternating regimens (*e.g.*, tobramycin or colistin). (*54*) Thus, additional therapeutic options for inhaled antibiotics, particularly for *P. aeruginosa* are urgently needed to improve clinical practice.

## CONCLUSION

Antimicrobial resistance, especially of the priority one pathogens (*i.e.*, *P. aeruginosa*, carbapenem-resistant; *A. baumannii*, carbapenem-resistant; *Enterobacteriaceae*, carbapenem-resistant, ESBL-producing), is a tremendous threat to human health and novel treatment options are urgently needed. The BAM complex with its major component BamA represents a clinically not addressed target and is of particular interest in this regard. In combination with early discovery pipelines as the recently developed high throughput inhibitor screening platform, (*55*) the herein described optimization workflow will likely help to identify and optimize new antibiotic candidates. The present study underlines the potential of bicyclic BamA-inhibitors for further development into an anti-Gram-negative antibiotic. This bears the potential to address an unmet medical need, *i.e.*, treatment of infections, such as complicated urinary tract infections (cUTI), hospital acquired pneumonia and ventilator acquired pneumonia (HAP and VAP), intrabdominal infections (IAI), and others, caused by MDR Gram-negative bacteria.

## MATERIALS AND METHODS

### General molecular biology methods

For plasmid isolation, the innuPREP plasmid DNA extraction kit (Analytik Jena, Germany) was used according to manual. For general cloning work, primers were synthesized by Eurofins (Eberswalde,Germany), Q5 polymerase, restriction enzymes and T7 ligase (NEB Biolabs, USA) were used according to manual. DNA fragments were analysed and purified on 1 % or 2 % TAE-agarose gels; DNA was recovered from gels using the large fragment recovery kit (ZymoResearch, USA). Matching DNA fragments were fused using self-prepared isothermal assembly mix. (*56*) *E. coli* was transformed by electroporation using standard methodology. Correct assembly was verified by test restriction and/or sequencing (Microsynth AG, Switzerland). Lysogenic-broth “LB” (peptone from casein 10 g/L, yeast extract 5 g/L and NaCl 5 g/L) was used for all cultivation steps during cloning. Kanamycin was used for selection of pRSF-duett-based vectors and T7 transcription was induced by addition of 0.5 mM IPTG to the growth medium.

### UPLC-HRMS analysis

UPLC-HRMS measurements were performed using an instrumental setup consisting of an Agilent Infinity 1290 UPLC system coupled to a DAD detector and a micrOTOFQ II mass spectrometer (Bruker Daltonics, Bremen, Germany) with an electrospray ionization source. For high(er) accuracy measurements a UPLC system of the aforementioned type coupled to DAD and ELSD detectors and a maXis II ESI-qTOF-UHRMS (Bruker Daltonics, Bremen, Germany) was used. The stationary phase of the UPLC system consisted in both cases of an Acquity UPLC BEH C18 1.7 μm (2.1 × 100 mm) column and an Acquity UPLC BEH C18 1.7 μm VanGuard Pre-Column (2.1 × 5 mm; both columns purchased from Waters, Eschborn, Germany). The LC system was operated using the following gradient (A: H2O, 0.1% formic acid; B: MeCN, 0.1 % formic acid): 0 min: 95 %A; 0.80 min: 95 %A; 18.70 min: 4.75 %A; 18.80 min: 0 %A; 23.00 min: 0 %A; 23.10 min: 95 %A; 25.00 min: 95 %A. The flow rate was set to 600 μL/min and the column oven temperature was maintained at 45 °C. A 10 mM sodium formate solution in H_2_O/ iPrOH (1:1) served as internal standard for the calibration of mass spectra. Recorded spectra were analysed with CompassData Analysis 4.2 and 5.3 (Bruker).

### Generation of *E. coli* Bap1^DarR^

The DAR-resistant expression host was generated according to a previously described method. (*13*) by introduction of three point mutations (*i.e.,* 1300A>G, 1334A>C, and 2113G>A) that confer DAR resistance (*11*) into the *E. coli* BAP1 (*57*) *bamA* gene. Briefly, the *bamA* gene containing three pointpoint mutations was amplified from *E. coli* DAR-resistant mutant (*11*) using bamA-recF and bamA-recR primer pair (Table S4). λ-Red recombineering was performed to introduce the linear amplified fragment into *E. coli* BAP1 with the help of pKD46. (*58*) DAR A (32 µg mL^−1^) was supplemented to LB agar for selection of *E. coli* BAP1 Dar^R^. The altered *bamA* gene was confirmed by Sanger sequencing (Microsynth AG).

### Alteration of the darobactin macrocycle size and core peptide length

To test the possibility of introduction or elimination of amino acids in the macrocycles of darobactin, primers were designed to bind in front of the core region of darA, with a 3’ extension covering the core region and another extension, overlapping the reverse primer for isothermal recircularization. Within these primers, the core region was altered to introduce the desired modification. pZW-ADC5 (*13*) was used as a template for PCR and the resulting linear fragment was re-circularized and used to transform E. coli Top10 and positive transformants were selected on LB_Kan_. After corroboration of the correct assembly by sequencing, the resulting plasmids were used to transform *E. coli* Bap1^DarR^, which were subsequently grown in LB_Kan_ and the production of darobactin derivatives was induced and production was continued at 30 °C and 200 rpm. Subsequently, 0.5 mL samples were taken, the cells were removed from the supernatant and the supernatant was desalted and 8-fold concentrated using self-packed C18 stage tips. Such prepared samples were analysed on UPLC-HRMS. To quickly assess the activity of these molecules, a parallelized, competition-based agar assay was utilized: 5 µL of an over-night preculture was spotted on an LB_Kan/IPTG_ agar plate and allowed to grow at 30 °C for 3 days. Thereafter, 10 mL of 60 °C LB_Kan_ was inoculated with 50 µL of *E. coli* MG1655 BamA6 and used to overlay the colony plate. After another over-night incubation at 37 °C, presence or absence of an antibiotic halo was recorded. Constructs and primers used to generate them can be found in Table S4.

### Single amino acid exchange in DarA

To exchange each amino acid of the darobactin scaffold with each other proteinogenic amino acid, the same strategy as before was used to create and analyse the different constructs. However, instead of introducing targeted mutations, each codon of the core amino acids of DarA was randomised by introduction of a NNK codon in each amino acid position at a time. After recircularization, clones were picked and sequenced to obtain each amino acid variant. In few cases, this randomised approach was not sufficient to cover all exchanges. Then, primers with the specific desired modification were used to clone the specific construct directly. Constructs and primers used to generate them can be found in Table S4.

### Generation of E. coli bamB::aac(3)IV

Darobactin-sensitive strain was generated by knocking out another member of the BAM complex, *bamB,* from *E. coli* bamA6 (previously known as *yaeT6* (Ruiz et al., 2006). The work started with amplification of apramycin resistance cassette (Sequence S1) by Q5 polymerase (New England Biolabs) using the primer pair bamB-strep-F and bamB-strep-R (Table S4). The resistance cassette was introduced to replace *bamB* by λ-Red recombineering. (*58*)

pKD46, a heat-sensitive plasmid carrying λ-Red genes, was introduced into *E. coli* bamA6 by electroporation, resulting *E. coli* bamA6-pKD46. This strain was then grown in LB medium with carbenicillin (50 µg mL^−1^) at 30°C for an hour, before being supplemented with 20 mM L-arabinose and further incubated. After the culture reached the OD_600_ of ≈0.6, it was washed three times with ice-cold 10% glycerol. The previously amplified apramycin resistance cassette (∼100 ng) was added into 50 µL of electrocompetent *E. coli* bamA6-pKD46 and electroporated. After electroporation, the cells were incubated for 1 hour for cell recovery, then plated on LB agar with apramycin (50 µg mL^−1^) and incubated at 37°C to promote the loss pf pKD46. The knockout of *bamB* was confirmed by PCR (bamB-testF2 and dbamBtest-R primer pair) and sequencing of the strain *E. coli bamB::aac*(*3*)IV using bamB-testF2 primer (Table S4).

The DAR-sensitive strain *E. coli bamB::aac*(*3*)IV was then made riboflavin auxotroph by replacement of *ribC* by *ribM*. The procedure was performed adapted from pKO3 plasmid system (Link et al., 1997). The plasmid pIN_ribM was prepared from an overnight culture grown in LB at 30 °C with 25 µg mL^−1^ chloramphenicol and introduced into *E. coli bamB::aac*(*3*)IV by electroporation. Directly after the electric shock, 1 mL SOC medium with 10 µM riboflavin was added, and the cells were allowed to recover for 1 hour at 30°C. Subsequently, the cells were plated on LB agar (supplemented with 10 µM riboflavin and 10 µg mL^−1^ chloramphenicol) and incubated overnight at 43°C in the dark. Large colonies were picked and resuspended in 1 mL LB with 10 µM riboflavin, serially diluted, and immediately plated on LB agar with 5% (w/v) sucrose and 10 µM riboflavin and incubated at 43°C. The loss of the pIN_ribM plasmid was tested by incubating replica plates at 30°C. Gene replacement was corroborated by PCR.

### Cloning a focussed randomised library for high throughput screening

To avoid homology related issues during library preparation, the darobactin BGC from pZW-ADC5 was recloned into pRSF-duett, omitting the second T7 expression cassette. Therefore, the BGC was amplified in two fragments and introduced to pRSF-duett using the *Nco*I and *Avr*II restriction sites, generating the vector pNB04. Subsequently, DarA was randomised by two separate PCRs with pNB04 as template using reverse primers in which the codons for N^2^, S^4^ and S^6^ were exchanged to NNK, combined with a fixed codon for F^7^ and W^7^, respectively. Furthermore, the K^5^ AAA codon was changed to AAG, to eliminate the *Hin*dIII restriction site in the DAR core region in both reverse primers. To eliminate misalignment issues and to in general improve DNA yields for subsequent cloning steps another PCR was performed using the previously generated randomised fragments as templates. Randomized PCR fragments and pNB04 were digested using *Mlu*I and *Hin*dIII and the restriction products were gel purified and ligated into the pNB04 backbone using T7 ligase. 20 x 1 µL of each ligation mixture was used to transform 50 µL of E. coli Top10 electrocompetent cells. Subsequently, each transformation reaction was spread on an LB_kan_ plate and incubated at 37 °C. After over-night incubation, the bacterial lawns were scraped from the plates, resuspended in 40 mL LB-medium and plasmids were isolated, generating the respective WXWXKXF and WXWXKXW libraries. DNA concentration was determined, and both libraries were combined to equal concentration, finalizing the focussed, randomised master library. From this master library, 2 x 3 PCR reactions were performed with 0.33 µL of master library as a template using primers outfitted with Illumina sequencing adapters. PCR reactions were gel purified and 3 reactions each were pooled and submitted to Illumina sequencing (Göttingen Genomics Laboratory (G2L), Göttingen, Germany) for library fitness and diversity analysis. The final expression library was prepared, transforming *E. coli* Bap1^DarR^ with 20 x 1 µL of the master library and plating each transformation reaction on LB_kan_. After over-night incubation at 37 °C, the lawns were recovered from the plates, resuspended in 20 mL LB_kan_, diluted 1:1 with sterile glycerol and stored at −80 °C. Constructs and primers used to generate them can be found in Table S4.

### Bioinformatic library analysis

The Illumina amplicon sequencing reads (see above) were analyzed as follows. First, the paired-end short reads were merged into a single extended fragment using FLASH (v1.2.11, -M 200). (*59*) To screen for the W^1^-X^2^-W^3^-X^4^-K^5^-X^6^-F^7^ and the W^1^-X^2^-W^3^-X^4^-K^5^-X^6^-W^7^ heptapeptides, two regular expressions were created, that match all possible codon combinations that translate to KETELSITDKALNELSNKPKIPEITAWXWXKXF and KETE…WXWXKXW, respectively, with X being all possible NNK codons. The regular expressions were than applied to the merged sequencing reads using an in-house Perl script. The matched sequences were then translated into amino acid sequences to compute coverage and fidelity.

### Parallelized analysis of enriched darobactin producers

First, colonies from the spotting plates were replated on fresh LB_Kan_ plates, from which 2 mL 96 deepwell master blocks containing 1 mL LB_Kan_ were inoculated, each well with one of the plated colonies. After overnight incubation at 37 °C at 200 rpm, 200 µL were transferred to 96-well plates, where the cells were precipitated by centrifugation. The supernatant was discarded and cell pellets were sent for plate sequencing using the T7 forward primer. Additionally, 50 µL per well were used to inoculate fresh 2 mL 96 deepwell masterblocks containing LB_Kan/IPTG_ for µ-scale darobactin production. Remaining 750 µL were diluted 1:1 with sterile glycerol and stored at −80 °C. For µ-scale darobactin production, incubation of the 96 deepwell plates took place at 30 °C and 200 rpm and was continued for 3 days. Subsequently, production was quenched by addition of 1:1 MeOH and the parallelized cultures were dried in-vacuo using a speedvac. After drying, the material was redissolved in 50 µL H_2_O (20x concentration), insoluble cell debris was removed by centrifugation and 1µL of the extracts was analysed by UPLC-HRMS, while remaining extract was used for activity testing.

### Large scale production and purification of darobactin derivatives

For production of tangible amounts of compounds, 15 x 1 L of LB_Kan_ in 2 L Erlenmeyer flasks were inoculated with 5 mL of derivative producer preculture and incubated at 28 °C at 180 rpm. At OD600 0.3-0.5, production was induced by addition of IPTG and production continued for 3 days. For the isolation, the same procedure as described in was used. (*44*) In brief, cells were separated from the supernatant and the supernatant was extracted using Amberlite XAD16N resin. The resin was washed with H2O and adsorbed material was eluted with 80% MeOH + 0.1 % formic acid. MeOH was then evaporated, resulting in an aqueous extract, which was then loaded to a SP Sepharose strong cation exchange column. Darobactin derivatives were eluted with 50 mM ammonium acetate buffers from pH 5 to pH 11. Darobactin derivatives then underwent final purification on a HPLC using a suited MeCN/H2O + 0.1 % FA gradient and were weighted out in glass vials.

### NMR spectroscopy

NMR spectra were recorded in D_2_O at 298 K on an Avance Neo 700 MHz NMR spectrometer equipped with a 5 mm CryoProbe Prodigy TCI (^1^H, ^15^N, ^13^C Z-GRD; ^1^H: 700.28 MHz, ^13^C: 176.09 MHz; Bruker BioSpin GmbH, Ettlingen, Germany). ^1^H spectra were referenced to the residual solvent signal (d = 4.79 ppm). (*60*) For ^13^C measurements 3-(trimethylsilyl)propionic-2,2,3,3-d_4_ acid sodium salt (TSPA, d = 1.7 ppm) (*61*) was used as external standard. NOESY, ROESY, and TOCSY spectra were additionally measured with H_2_O suppression. Analysis of NMR spectra was accomplished using the software TopSpin 3.6.0 and TopSpin 4.3.0 (Bruker BioSpin GmbH, Rheinstetten, Germany). Chemical shifts are given in ppm and coupling constants in Hz. Molecular models for analysis of NOESY and ROESY spectra were obtained using the web-based tool “MolView”. (*62*)

### Genome sequencing EXT111762

The clinical isolate *Pseudomonas aeruginosa* was isolated from a cystic fibrosis patient in 2006. (23) The genetic material of strain EXT111672 was sequenced by macrogen using illumina paired end whole genome sequencing resulting in a total of 22.998.364 reads with a length of 151 base pairs. The reads were assembled to a draft genome (96 contigs; N50: 402990) of an estimated size of 6.3 kb and GC content of 66.5%. Database alignment of the 16S rRNA gene sequences, MLST (acsA, aroE, guaA, mutL, nuoD, ppsA, and trpE) and ANI identified the isolate as *Pseudomonas aeruginosa*. The draft genome is deposited at Genbank (ID to be added after acceptance). AMRfinder (*63*) and CART (*64*) were used to annotate resistance determinants and results are summarized in the supplemental files.

### MIC testing

The minimum inhibitory concentration of the purified compounds were determined as previously described. (*16, 24*) Briefly, overnight cultures of the test strains were cultivated at 37°C and 180rpm in Mueller Hinton II broth. For the BamA inhibitor sensitive *E. coli* MG1655 *bamB::aac*(*3*)*-IV*, we added 50 µg mL^−1^ apramycin and kanamycin to the culture medium. Before the bacterial cultures were distributed to the 96 well assay plates, the cell density of the respective culture was adjusted to 5 × 10^5^ cells mL^−1^. All compounds were dissolved in water and tested in a triplicated 12-point dilution series. In addition, dilution series of ceftazidime, ciprofloxacin and gentamicin were used as positive and quality control for each strain on each assay plate. Bacterial suspension without test compound or positive control was used as negative control, e.g. growth control. Medium background was averaged from 5 replicates. Assay plates were incubated for 18h at 180rpm and 85% relative humidity before the turbidity of all wells was determined with a microplate spectrophotometer at 600 nm (LUMIstar® Omega BMG Labtech). Growth inhibition was calculated relative to the absorption of the controls.

### Time-kill-curve

Bacteria killing kinetics in contact with DAR B were prepared as described before with only slight modifications. (*65*) The overnight cultures of *Pseudomonas aeruginosa* ATCC27853 and EXT111762 were adjusted to McFarland Standard 1 and further diluted to 5 × 10^5^ cells mL^−1^ in cation adjusted Mueller Hinton II broth (BD). A volume of 5 mL was distributed to each cultivation flask and supplemented with either 0.5x, 1x, 4x or 8x of the previously determined MIC of DAR B (8 µg ml^−1^ for ATCC27853 and 2 µg mL^−1^ for EXT111762). Additionally, a growth control without any supplementation was inoculated. Each strain-treatment combination was replicated in three separated culture flasks. The cultures were incubated at 37°C and 180 rpm and sampled after 0h, 0.5h, 1h, 4h, 8h, 24h and 48h. At each sampling time, 20 µL culture volume was removed from each flask and subject to a 6-point 1:10 dilution series in U-bottom microtiter plates. Form each dilution (10^−1^ to 10^−6^ dilution rel. to the initial culture) 10 µL were spotted on Mueller Hinton II agar plates. After 18h (ATCC27853) or 24h (EXT111762) the CFUs were counted and the concentration calculated according to:

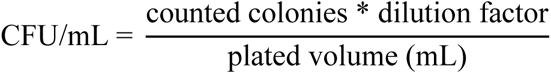

Counting was carried out using a stereomicroscope if 10-100 colonies were grown out of the spotted aliquot. At least 2 dilutions were counted per treatment and replicate. The upper limit of detection was defined as >100 CFU in the 10^−6^ diluted aliquot. Accordingly, the lower limit of detection was reached if no CFU was observed in the 10^−1^ aliquot.

### Early ADMET

Early ADMET study of DAR B and DAR B9 was performed by Eurofins Discovery.

### Partition Coefficient

The total amount of compound was determined as the peak area of the principal peak in a calibration standard (100 µM) containing organic solvent (methanol/water, 60/40, v/v). The amount of compound in buffer was determined as the combined, volume-corrected, and weighted areas of the corresponding peaks in the aqueous phases of three organic-aqueous samples of different composition. An automated weighting system was used to ensure the preferred use of raw data from those samples with well quantifiable peak signals. The amount of compound in organic solvent was calculated by subtraction. Subsequently, *LogD* was calculated as the Log_10_ of the amount of compound in the organic phase divided by the amount of compound in the aqueous phase.

### Fluorescein assessment for Permeability assays

Fluorescein was used as the cell monolayer integrity marker. Fluorescein permeability assessment (in the A-B direction at pH 7.4 on both sides) was performed after the permeability assay for the test compound. The cell monolayer that had a fluorescein permeability of less than 1.5 × 10^−6^ cm s^−1^ for Caco-2 and MDR1-MDCKII cells and 2.5 × 10^−6^ cm s^−1^ for MDCKII cells was considered intact, and the permeability result of the test compound from intact cell monolayer is reported.

### Caco-2 Permeability

The apparent permeability coefficient (P_app_) of the test compound was calculated as follows:

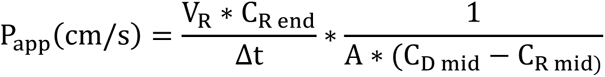

whereby V_R_ is the volume of the receiver chamber. C_R,end_ is the concentration of the test compound in the receiver chamber at the end time point, Δt is the incubation time and A is the surface area of the cell monolayer. C_D,mid_ is the calculated mid-point concentration of the test compound in the donor side, which is the mean value of the donor concentration at time 0 minute and the donor concentration at the end time point. C_R,mid_ is the mid-point concentration of the test compound in the receiver side, which is one half of the receiver concentration at the end time point. Concentrations of the test compound were expressed as peak areas of the test compound.

### Cytochrome P450 Inhibition (HPLC-UV/VIS and HPLC-MS/MS detection)

Peak areas corresponding to the metabolite of each substrate were recorded. The percentage of control activity was then calculated by comparing the peak area obtained in the presence of the test compound to that obtained in the absence of the test compound. Subsequently, the percent inhibition was calculated by subtracting the percent control activity from 100 for each compound. IC_50_ values (concentration causing a half-maximal inhibition of control values) were determined by non-linear regression analysis of the concentration-response curve using Hill equation curve fitting.

### Cell Viability

The percentage of control is calculated using the following equation. The percentage of inhibition is calculated by subtracting the percent of control from 100. The IC_50_ value (concentration causing a half-maximal inhibition of the control value) is determined by non-linear regression analysis of the concentration-response curve using the Hill equation.

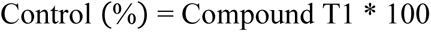

Compound is the individual reading in the presence of the test compound. T1 is the mean reading in the absence of the test compound.

### hERG Potassium Channel Assay - Qube APC

Electrophysiological assays were conducted to profile darobactin B for activity on the ion channel target CHO-hERG using the Qube electrophysiological platform. Darobactin B compound was tested in the presence of 0.1% Pluronic F-68 Non-Ionic Surfactant and at room temperature. After whole cell configuration is achieved, the cell is held at −80mV. The cell is depolarized to +40mV for 500ms and then to −80mV over a 100ms ramp to elicit the hERG tail current. This paradigm is delivered once every 8s to monitor the current amplitude. The peak current amplitude was calculated before and after compound addition and the amount of inhibition was assessed by dividing the Test compound current amplitude by the Control current amplitude. Control data is the mean hERG current amplitude collected 15 seconds at the end of the control period; Test compound data is the mean hERG current amplitude collected 15 seconds at the end of test concentration application for each concentration. All data were filtered for seal quality, seal drop, and current amplitude. Verapamil was used as reference standards, as an integral part of each assay to ensure the validity of the results obtained.

### *In vivo* efficacy

C57BL/6N wildtype mice were purchased from Charles River Laboratories at the age of 8 - 12 weeks. Animal experiments were conducted according to the legal regulations of the German Animal Welfare Act (Tierschutzgesetz) and approved by the regional authorities of the State of Hesse (Regierungspräsidium Giessen). *P. aeruginosa* (strain ATCC 29260), or *K. pneumoniae* (strain ATCC 700603) was cultured in Luria broth (LB) medium (Roth) at 37°C with aeration (for *P. aeruginosa*) and harvested at mid-log phase. Inoculum was determined by serial dilutions on blood agar plates, and mice were infected with 1 × 10^5^ colony forming units (CFU) of *P. aeruginosa* or 1 × 10^7^ CFU of *K. pneumoniae* instilled in 50µl NaCl 0,9% intranasally. (*66*) Before inoculation, mice were short-term anesthetized by inhalation of isoflurane. Darobactin B and B9 or NaCl 0.9% (= control) were administered intraperitoneally (à 25mg/kg in 100µl) or intratracheally (à 50mg/kg in 70µl) at indicated time points. At indicated time points mice were euthanized, lungs were removed and placed in 5 ml NaCl 0.9%. The lung tissues were dissected using the gentleMACS dissociator (Miltenyi Biotec). Next these suspensions were filtered through 100 µm cell filters. Subsequently homogenates were serially diluted in NaCl 0.9% on blood agar plates to determine bacterial outgrowth.

## List of Supplementary Materials

Figures S1 to S109 including NMR data

Tables S1 to S7

Sequence S1

## Acknowledgments

The authors thank Mona Abdullahi for compound purification, Kirsten Bommersheim for MIC assays, Christoph Hartwig for chemical analytical support and Heike Hausmann and the NMR facility of the Justus-Liebig-University for conducting the NMR measurements.

## Author contributions

Conceptualization: TFS

Methodology: NB, MM, SS, SR, ZGW

Investigation: NB, ZGW, IDMK, JF, MM, UtM

Visualization: NB, ZGW, JF, UtM, MM

Funding acquisition: TFS, SH, UlM

Project administration: TFS, UlM

Supervision: TFS, SH, UlM,

Writing – original draft: NB, ZGW, MM, UlM, TFS

Writing – review & editing: NB, ZGW, MM, UlM, TFS

## Data and materials availability

The experimental data that support the findings of this study is shown in the article and its supplementary materials.

## References and Notes

1. U. Hofer, The cost of antimicrobial resistance. Nat. Rev. Microbiol. 17, 3 (2019).

2. World Health Organization, Global antimicrobial resistance and use surveillance system (GLASS) report 2022 (Geneva, 2022).

3. P. V. Jonas, Olga B.; Irwin, Alec; Berthe, Franck Cesar Jean; Le Gall, Francois G.; Marquez, Drug-resistant infections : a threat to our economic future (Vol. 2) : final report. HNP/Agriculture Glob. Antimicrob. Resist. Initiat. Washington, D.C. World Bank Group. (2017), doi:10.1596/26707.

4. C. J. Murray, K. S. Ikuta, F. Sharara, L. Swetschinski, G. Robles Aguilar, A. Gray, C. Han, C. Bisignano, P. Rao, E. Wool, S. C. Johnson, A. J. Browne, M. G. Chipeta, F. Fell, S. Hackett, G. Haines-Woodhouse, B. H. Kashef Hamadani, E. A. P. Kumaran, B. McManigal, R. Agarwal, S. Akech, S. Albertson, J. Amuasi, J. Andrews, A. Aravkin, E. Ashley, F. Bailey, S. Baker, B. Basnyat, A. Bekker, R. Bender, A. Bethou, J. Bielicki, S. Boonkasidecha, J. Bukosia, C. Carvalheiro, C. Castañeda-Orjuela, V. Chansamouth, S. Chaurasia, S. Chiurchiù, F. Chowdhury, A. J. Cook, B. Cooper, T. R. Cressey, E. Criollo-Mora, M. Cunningham, S. Darboe, N. P. J. Day, M. De Luca, K. Dokova, A. Dramowski, S. J. Dunachie, T. Eckmanns, D. Eibach, A. Emami, N. Feasey, N. Fisher-Pearson, K. Forrest, D. Garrett, P. Gastmeier, A. Z. Giref, R. C. Greer, V. Gupta, S. Haller, A. Haselbeck, S. I. Hay, M. Holm, S. Hopkins, K. C. Iregbu, J. Jacobs, D. Jarovsky, F. Javanmardi, M. Khorana, N. Kissoon, E. Kobeissi, T. Kostyanev, F. Krapp, R. Krumkamp, A. Kumar, H. H. Kyu, C. Lim, D. Limmathurotsakul, M. J. Loftus, M. Lunn, J. Ma, N. Mturi, T. Munera-Huertas, P. Musicha, M. M. Mussi-Pinhata, T. Nakamura, R. Nanavati, S. Nangia, P. Newton, C. Ngoun, A. Novotney, D. Nwakanma, C. W. Obiero, A. Olivas-Martinez, P. Olliaro, E. Ooko, E. Ortiz-Brizuela, A. Y. Peleg, C. Perrone, N. Plakkal, A. Ponce-de-Leon, M. Raad, T. Ramdin, A. Riddell, T. Roberts, J. V. Robotham, A. Roca, K. E. Rudd, N. Russell, J. Schnall, J. A. G. Scott, M. Shivamallappa, J. Sifuentes-Osornio, N. Steenkeste, A. J. Stewardson, T. Stoeva, N. Tasak, A. Thaiprakong, G. Thwaites, C. Turner, P. Turner, H. R. van Doorn, S. Velaphi, A. Vongpradith, H. Vu, T. Walsh, S. Waner, T. Wangrangsimakul, T. Wozniak, P. Zheng, B. Sartorius, A. D. Lopez, A. Stergachis, C. Moore, C. Dolecek, M. Naghavi, Global burden of bacterial antimicrobial resistance in 2019: a systematic analysis. Lancet 399, 629–655 (2022).

5. E. Tacconelli, E. Carrara, A. Savoldi, S. Harbarth, M. Mendelson, D. L. Monnet, C. Pulcini, G. Kahlmeter, J. Kluytmans, Y. Carmeli, M. Ouellette, K. Outterson, J. Patel, M. Cavaleri, E. M. Cox, C. R. Houchens, M. L. Grayson, P. Hansen, N. Singh, U. Theuretzbacher, N. Magrini, A. O. Aboderin, S. S. Al-Abri, N. Awang Jalil, N. Benzonana, S. Bhattacharya, A. J. Brink, F. R. Burkert, O. Cars, G. Cornaglia, O. J. Dyar, A. W. Friedrich, A. C. Gales, S. Gandra, C. G. Giske, D. A. Goff, H. Goossens, T. Gottlieb, M. Guzman Blanco, W. Hryniewicz, D. Kattula, T. Jinks, S. S. Kanj, L. Kerr, M. P. Kieny, Y. S. Kim, R. S. Kozlov, J. Labarca, R. Laxminarayan, K. Leder, L. Leibovici, G. Levy-Hara, J. Littman, S. Malhotra-Kumar, V. Manchanda, L. Moja, B. Ndoye, A. Pan, D. L. Paterson, M. Paul, H. Qiu, P. Ramon-Pardo, J. Rodríguez-Baño, M. Sanguinetti, S. Sengupta, M. Sharland, M. Si-Mehand, L. L. Silver, W. Song, M. Steinbakk, J. Thomsen, G. E. Thwaites, J. W. van der Meer, N. Van Kinh, S. Vega, M. V. Villegas, A. Wechsler-Fördös, H. F. L. Wertheim, E. Wesangula, N. Woodford, F. O. Yilmaz, A. Zorzet, Discovery, research, and development of new antibiotics: the WHO priority list of antibiotic-resistant bacteria and tuberculosis. Lancet Infect. Dis. 18, 318–327 (2018).

6. M. Lakemeyer, W. Zhao, F. A. Mandl, P. Hammann, S. A. Sieber, Thinking Outside the Box—Novel Antibacterials To Tackle the Resistance Crisis. Angew. Chemie Int. Ed. 57, 14440–14475 (2018).

7. J. A. Al-Tawfiq, H. Momattin, A. Y. Al-Ali, K. Eljaaly, R. Tirupathi, M. B. Haradwala, S. Areti, S. Alhumaid, A. A. Rabaan, A. Al Mutair, P. Schlagenhauf, Antibiotics in the pipeline: a literature review (2017–2020). Infection 50, 553–564 (2022).

8. K. Lewis, The Science of Antibiotic Discovery. Cell 181, 29–45 (2020).

9. M. Hutchings, A. Truman, B. Wilkinson, Antibiotics: past, present and future. Curr. Opin. Microbiol. 51, 72–80 (2019).

10. D. J. Newman, G. M. Cragg, Natural Products as Sources of New Drugs over the Nearly Four Decades from 01/1981 to 09/2019. J. Nat. Prod. 83, 770–803 (2020).

11. Y. Imai, K. J. Meyer, A. Iinishi, Q. Favre-Godal, R. Green, S. Manuse, M. Caboni, M. Mori, S. Niles, M. Ghiglieri, C. Honrao, X. Ma, J. J. Guo, A. Makriyannis, L. Linares-Otoya, N. Böhringer, Z. G. Wuisan, H. Kaur, R. Wu, A. Mateus, A. Typas, M. M. Savitski, J. L. Espinoza, A. O’Rourke, K. E. Nelson, S. Hiller, N. Noinaj, T. F. Schäberle, A. D’Onofrio, K. Lewis, A new antibiotic selectively kills Gram-negative pathogens. Nature 576, 459–464 (2019).

12. H. Nguyen, I. D. Made Kresna, N. Böhringer, J. Ruel, E. D. La Mora, J. C. Kramer, K. Lewis, Y. Nicolet, T. F. Schäberle, K. Yokoyama, Characterization of a Radical SAM Oxygenase for the Ether Crosslinking in Darobactin Biosynthesis. J. Am. Chem. Soc. 144, 18876–18886 (2022).

13. Z. G. Wuisan, I. D. M. Kresna, N. Böhringer, K. Lewis, T. F. Schäberle, Optimization of heterologous Darobactin A expression and identification of the minimal biosynthetic gene cluster. Metab. Eng. 66, 123–136 (2021).

14. M. P. Bos, V. Robert, J. Tommassen, Biogenesis of the gram-negative bacterial outer membrane. Annu. Rev. Microbiol. 61, 191–214 (2007).

15. Y. Gu, H. Li, H. Dong, Y. Zeng, Z. Zhang, N. G. Paterson, P. J. Stansfeld, Z. Wang, Y. Zhang, W. Wang, C. Dong, Structural basis of outer membrane protein insertion by the BAM complex. Nature 531, 64–69 (2016).

16. C. E. Seyfert, C. Porten, B. Yuan, S. Deckarm, F. Panter, C. D. Bader, J. Coetzee, F. Deschner, K. H. M. E. Tehrani, P. G. Higgins, H. Seifert, T. C. Marlovits, J. Herrmann, R. Müller, Darobactins Exhibiting Superior Antibiotic Activity by Cryo-EM Structure Guided Biosynthetic Engineering**. Angew. Chemie - Int. Ed. 62 (2023), doi:10.1002/anie.202214094.

17. N. Böhringer, R. Green, Y. Liu, U. Mettal, M. Marner, M. Modaresi Seyed, P. Jakob Roman, Z. Wuisan, T. Maier, A. Iinishi, S. Hiller, K. Lewis, T. F. Schäberle, Mutasynthetic Production and Antimicrobial Characterization of Darobactin Analogs. Microbiol. Spectr. 9, e01535–21 (2021).

18. S. Groß, F. Panter, D. Pogorevc, C. E. Seyfert, S. Deckarm, C. D. Bader, J. Herrmann, R. Müller, Improved broad-spectrum antibiotics against Gram-negative pathogensviadarobactin biosynthetic pathway engineering. Chem. Sci. 12, 11882–11893 (2021).

19. S. Schmitt, M. Montalbán-López, D. Peterhoff, J. Deng, R. Wagner, M. Held, O. P. Kuipers, S. Panke, Analysis of modular bioengineered antimicrobial lanthipeptides at nanoliter scale. Nat. Chem. Biol. 15, 437–443 (2019).

20. N. J. Guido, S. Handerson, E. M. Joseph, D. Leake, L. A. Kung, Determination of a screening metric for high diversity DNA libraries. PLoS One 11, 1–18 (2016).

21. C. L. Hagan, D. Kahne, The reconstituted Escherichia coli Bam complex catalyzes multiple rounds of β-barrel assembly. Biochemistry 50, 7444–7446 (2011).

22. N. Ruiz, T. Wu, D. Kahne, T. J. Silhavy, Probing the Barrier Function of the Outer Membrane with Chemical Conditionality. ACS Chem. Biol. 1, 385–395 (2006).

23. M. Marner, L. Kolberg, J. Horst, N. Böhringer, J. Hübner, I. D. M. Kresna, Y. Liu, Antimicrobial Activity of Ceftazidime-Avibactam, Ceftolozane-Multidrug-Resistant Pseudomonas aeruginosa Isolates from. Microbiol. Spectr. (2023), doi:10.1128/spectrum.04437-22.

24. D. Wang, C. Seeve, L. S. Pierson, E. A. Pierson, Transcriptome profiling reveals links between ParS/ParR, MexEF-OprN, and quorum sensing in the regulation of adaptation and virulence in Pseudomonas aeruginosa. BMC Genomics 14, 618 (2013).

25. H. Ogasawara, S. Shinohara, K. Yamamoto, A. Ishihama, Novel regulation targets of the metal-response BasS-BasR two-component system of Escherichia coli. Microbiol. (United Kingdom) 158, 1482–1492 (2012).

26. L. Fernández, H. Jenssen, M. Bains, I. Wiegand, W. J. Gooderham, R. E. W. Hancock, The two-component system CprRS senses cationic peptides and triggers adaptive resistance in Pseudomonas aeruginosa independently of ParRS. Antimicrob. Agents Chemother. 56, 6212–6222 (2012).

27. L. J. V Piddock, Multidrug-resistance efflux pumpsnot just for resistance. Nat. Rev. Microbiol. 4, 629–636 (2006).

28. N. Masuda, E. Sakagawa, S. Ohya, N. Gotoh, H. Tsujimoto, T. Nishino, Substrate specificities of MexAB-OprM, MexCD-OprJ, and MexXY-OprM efflux pumps in Pseudomonas aeruginosa. Antimicrob. Agents Chemother. 44, 3322–3327 (2000).

29. Z. Beharry, T. Palzkill, Functional analysis of active site residues of the fosfomycin resistance enzyme FosA from Pseudomonas aeruginosa. J. Biol. Chem. 280, 17786–17791 (2005).

30. B. M. D., T. M. A., R. J. D., B. C. R., G. Ioannis, H. A. M., C. Emilia, P. Fabio, D. J. P., P.-W. K. M., H. Shozeb, B. R. A., Deciphering the Evolution of Cephalosporin Resistance to Ceftolozane-Tazobactam in Pseudomonas aeruginosa. MBio 9, 10.1128/mbio.02085-18 (2018).

31. P. A. White, H. W. Stokes, K. L. Bunny, R. M. Hall, Characterisation of a chloramphenicol acetyltransferase determinant found in the chromosome of Pseudomonas aeruginosa. FEMS Microbiol. Lett. 175, 27–35 (1999).

32. T. Takashi, S. Eiko, S. Mie, Detection of gyrA Mutations among 335Pseudomonas aeruginosa Strains Isolated in Japan and Their Susceptibilities to Fluoroquinolones. Antimicrob. Agents Chemother. 43, 406–409 (1999).

33. N. A. Genthe, J. B. Thoden, H. M. Holden, Structure of the Escherichia coli ArnA N-formyltransferase domain in complex with N5-formyltetrahydrofolate and UDP-Ara4N. Protein Sci. 25, 1555–1562 (2016).

34. U. M. Zanger, M. Schwab, Cytochrome P450 enzymes in drug metabolism: Regulation of gene expression, enzyme activities, and impact of genetic variation. Pharmacol. Ther. 138, 103–141 (2013).

35. B. S. Quon, C. H. Goss, B. W. Ramsey, Inhaled antibiotics for lower airway infections. Ann. Am. Thorac. Soc. 11, 425–434 (2014).

36. S. Ehrmann, F. Barbier, J. Demiselle, J.-P. Quenot, J.-E. Herbrecht, D. Roux, J.-C. Lacherade, M. Landais, P. Seguin, D. Schnell, A. Veinstein, P. Gouin, S. Lasocki, Q. Lu, G. Beduneau, M. Ferrandiere, G. Plantefève, C. Dahyot-Fizelier, N. Chebib, E. Mercier, N. Heuzé-Vourc’h, R. Respaud, N. Gregoire, D. Garot, M.-A. Nay, F. Meziani, P. Andreu, R. Clere-Jehl, N. Zucman, M.-A. Azaïs, M. Saint-Martin, C. S. Gandonnière, D. Benzekri, H. Merdji, E. Tavernier, Inhaled Amikacin to Prevent Ventilator-Associated Pneumonia. N. Engl. J. Med. (2023), doi:10.1056/nejmoa2310307.

37. M. Nesic, D. B. Ryffel, J. Maturano, M. Shevlin, S. R. Pollack, D. R. J. Gauthier, P. Trigo-Mouriño, L.-K. Zhang, D. M. Schultz, J. M. McCabe Dunn, L.-C. Campeau, N. R. Patel, D. A. Petrone, D. Sarlah, Total Synthesis of Darobactin A. J. Am. Chem. Soc. 144, 14026–14030 (2022).

38. Y.-C. Lin, F. Schneider, K. J. Eberle, D. Chiodi, H. Nakamura, S. H. Reisberg, J. Chen, M. Saito, P. S. Baran, Atroposelective Total Synthesis of Darobactin A. J. Am. Chem. Soc. 144, 14458–14462 (2022).

39. D. J. Collins, A. Neild, A. deMello, A. Q. Liu, Y. Ai, The Poisson distribution and beyond: Methods for microfluidic droplet production and single cell encapsulation. Lab Chip 15, 3439–3459 (2015).

40. M. Oberpaul, S. Brinkmann, M. Marner, S. Mihajlovic, B. Leis, M. A. Patras, C. Hartwig, A. Vilcinskas, P. E. Hammann, T. F. Schäberle, M. Spohn, J. Glaeser, Combination of high-throughput microfluidics and FACS technologies to leverage the numbers game in natural product discovery. Microb. Biotechnol. 15, 415–430 (2022).

41. K. A. Muñoz, P. J. Hergenrother, Computational discovery of dynobactin antibiotics. Nat. Microbiol. 7, 1512–1513 (2022).

42. R. D. Miller, A. Iinishi, S. M. Modaresi, B. K. Yoo, T. D. Curtis, P. J. Lariviere, L. Liang, S. Son, S. Nicolau, R. Bargabos, M. Morrissette, M. F. Gates, N. Pitt, R. P. Jakob, P. Rath, T. Maier, A. G. Malyutin, J. T. Kaiser, S. Niles, B. Karavas, M. Ghiglieri, S. E. J. Bowman, D. C. Rees, S. Hiller, K. Lewis, Computational identification of a systemic antibiotic for gram-negative bacteria (2022).

43. N. Böhringer, J. C. Kramer, E. de la Mora, L. Padva, Z. G. Wuisan, Y. Liu, M. Kurz, M. Marner, H. Nguyen, P. Amara, K. Yokoyama, Y. Nicolet, U. Mettal, T. F. Schäberle, Genome- and metabolome-guided discovery of marine BamA inhibitors revealed a dedicated darobactin halogenase. Cell Chem. Biol. 30, 943–952.e7 (2023).

44. I. Nilsson, A. Sääf, P. Whitley, G. Gafvelin, C. Waller, G. von Heijne, Proline-induced disruption of a transmembrane α-helix in its natural environment11Edited by F. Cohen. J. Mol. Biol. 284, 1165–1175 (1998).

45. H. Kaur, R. P. Jakob, J. K. Marzinek, R. Green, Y. Imai, J. R. Bolla, E. Agustoni, C. V. Robinson, P. J. Bond, K. Lewis, T. Maier, S. Hiller, The antibiotic darobactin mimics a β-strand to inhibit outer membrane insertase. Nature 593, 125–129 (2021).

46. EMA/CHMP/ICH, ICH Harmonised Guideline – Drug Interaction Studies (M12) - Step 2b. Eur. Med. Agency 31, 1–69 (2022).

47. H. Yamashita, S., Tanaka, Y., Endoh, Y., Taki, Y., Sakane, T., Nadai, T., & Sezaki, 1997_Analysis of drug permeation across Caco-2 monolayer.pdf Pharm. Res. 14, 486–491 (1997).

48. P. D. Tamma, S. L. Aitken, R. A. Bonomo, A. J. Mathers, D. van Duin, C. J. Clancy, Infectious Diseases Society of America 2023 Guidance on the Treatment of Antimicrobial Resistant Gram-Negative Infections. Clin. Infect. Dis., 1–53 (2023).

49. A. L. H. Kwa, C. S. Loh, J. G. H. Low, A. Kurup, V. H. Tam, Nebulized colistin in the treatment of pneumonia due to multidrug-resistant Acinetobacter baumannii and Pseudomonas aeruginosa. Clin. Infect. Dis. 41, 754–757 (2005).

50. D. H. Hamer, Treatment of nosocomial pneumonia and tracheobronchitis caused by multidrug-resistant Pseudomonas aeruginosa with aerosolized colistin. Am. J. Respir. Crit. Care Med. 162, 328–330 (2000).

51. C. Castellani, A. J. A. Duff, S. C. Bell, H. G. M. Heijerman, A. Munck, F. Ratjen, I. Sermet-Gaudelus, K. W. Southern, J. Barben, P. A. Flume, P. Hodková, N. Kashirskaya, M. N. Kirszenbaum, S. Madge, H. Oxley, B. Plant, S. J. Schwarzenberg, A. R. Smyth, G. Taccetti, T. O. F. Wagner, S. P. Wolfe, P. Drevinek, ECFS best practice guidelines: the 2018 revision. J. Cyst. Fibros. 17, 153–178 (2018).

52. E. C. Dasenbrook, M. W. Konstan, D. R. VanDevanter, Association between the introduction of a new cystic fibrosis inhaled antibiotic class and change in prevalence of patients receiving multiple inhaled antibiotic classes. J. Cyst. Fibros. 14, 370–375 (2015).

53. P. Rath, A. Hermann, R. Schaefer, E. Agustoni, J. M. Vonach, M. Siegrist, C. Miscenic, A. Tschumi, D. Roth, C. Bieniossek, S. Hiller, High-throughput screening of BAM inhibitors in native membrane environment. Nat. Commun. 14 (2023), doi:10.1038/s41467-023-41445-w.

54. D. G. Gibson, L. Young, R.-Y. Chuang, J. C. Venter, C. A. Hutchison, H. O. Smith, Enzymatic assembly of DNA molecules up to several hundred kilobases. Nat. Methods 6, 343–345 (2009).

55. B. A. Pfeifer, S. J. Admiraal, H. Gramajo, D. E. Cane, C. Khosla, Biosynthesis of complex polyketides in a metabolically engineered strain of E. coli. Science (80-.). 291, 1790–1792 (2001).

56. K. A. Datsenko, B. L. Wanner, One-step inactivation of chromosomal genes in Escherichia coli K-12 using PCR products. Proc. Natl. Acad. Sci. U. S. A. 97, 6640–6645 (2000).

57. T. Magoč, S. L. Salzberg, FLASH: fast length adjustment of short reads to improve genome assemblies. Bioinformatics 27, 2957–2963 (2011).

58. H. E. Gottlieb, V. Kotlyar, A. Nudelman, NMR Chemical Shifts of Common Laboratory Solvents as Trace Impurities. J. Org. Chem. 62, 7512–7515 (1997).

59. R. Hollenstein, H.-O. Kalinowski, S. Berger and S. Braun. 13C NMR-Spektroskopie. Georg Thieme Verlag, Stuttgart 1984. pp. 685. DM98. Magn. Reson. Chem. 23, 589 (1985).

60. Molview.org.

61. M. Feldgarden, V. Brover, D. H. Haft, A. B. Prasad, D. J. Slotta, I. Tolstoy, G. H. Tyson, S. Zhao, C. H. Hsu, P. F. McDermott, D. A. Tadesse, C. Morales, M. Simmons, G. Tillman, J. Wasilenko, J. P. Folster, W. Klimke, Validating the AMRFINder tool and resistance gene database by using antimicrobial resistance genotype-phenotype correlations in a collection of isolates. Antimicrob. Agents Chemother. 63 (2019), doi:10.1128/AAC.00483-19.

62. B. P. Alcock, W. Huynh, R. Chalil, K. W. Smith, A. R. Raphenya, M. A. Wlodarski, A. Edalatmand, A. Petkau, S. A. Syed, K. K. Tsang, S. J. C. Baker, M. Dave, M. C. McCarthy, K. M. Mukiri, J. A. Nasir, B. Golbon, H. Imtiaz, X. Jiang, K. Kaur, M. Kwong, Z. C. Liang, K. C. Niu, P. Shan, J. Y. J. Yang, K. L. Gray, G. R. Hoad, B. Jia, T. Bhando, L. A. Carfrae, M. A. Farha, S. French, R. Gordzevich, K. Rachwalski, M. M. Tu, E. Bordeleau, D. Dooley, E. Griffiths, H. L. Zubyk, E. D. Brown, F. Maguire, R. G. Beiko, W. W. L. Hsiao, F. S. L. Brinkman, G. Van Domselaar, A. G. McArthur, CARD 2023: expanded curation, support for machine learning, and resistome prediction at the Comprehensive Antibiotic Resistance Database. Nucleic Acids Res. 51, D690–D699 (2023).

63. W. Wang, Y. Li, X. Guo, A mouse model of Citrobacter rodentium oral infection and evaluation of innate and adaptive immune responses. STAR Protoc. 1 (2020), doi:10.1016/j.xpro.2020.100218.

64. U. Matt, J. M. Warszawska, M. Bauer, W. Dietl, I. Mesteri, B. Doninger, I. Haslinger, G. Schabbauer, T. Perkmann, C. J. Binder, S. Reingruber, P. Petzelbauer, S. Knapp, Bβ15–42 Protects against Acid-induced Acute Lung Injury and Secondary Pseudomonas Pneumonia In Vivo. Am. J. Respir. Crit. Care Med. 180, 1208–1217 (2009).

