## Supporting Information for "Nanoliter-scale selection of optimized bioengineered peptide antibiotics that rescue mice with bacterial lung infection"

Table S1 Minimum inhibitory concentrations of darobactin A against different *E. coli* MG1655 knockdown strains. Recombination was carried out using the lambda red system. Values given in µg/mL. Strain *E. coli* MG1655 bamB::aac(3)-IV was selected as ultra-sensitive strain for HTS of the master library, confirmatory screening of sorted events and later MIC determination of purified compounds.

| | **Darobactin A** |
|---|---|
| ***E.coli* MG1655 wt** | 8-4 |
| ***E.coli* MG1655 bamA6** | 0.25-0.125 |
| ***E.coli* MG1655 *bamB::aac(3)IV*** | >0.031 |

..................Column Break..................

Table S2. Antibiogram of Pseudomonas aeruginosa EXT111762 determined by microbroth-dilution assays.

| **EXT111762** | |
|---|---|
| | MIC [µg/mL] |
| Cefiderocol (FDC) | 16-8 |
| Ceftazidime (CEF) | 32-16 |
| Cefotaxime (CTX) | >64 |
| Imipenem (IMP) | 64 |
| Meropenem (MER) | 64 |
| Ciprofloxacin (CIP) | 0.5 – 0.25 |
| Tetracycline (TET) | 1 |
| Colistin (COL) | 0.25 |
| Tobramycin (TOB) | 0.5 |
| Gentamycin (GEN) | 0.5 |
| Chloramphenicol (CHL) | 8 |
| Linezolid (LIN) | >64 |
| Azithromycin (AZN) | 32 |
| Rifampicin (RIF) | 16 |
| Darobactin A (DAA) | >64 |
| Darobactin B (DAB) | 2-1 |
| Darobactin B9 (DAB9) | 2-1 |

Table S3. Minimum inhibitory concentrations of darobactin B, B9 and standard antibiotics ceftazidime (CZA), ciprofloxacin (CIP) and gentamicin (GEN). Each MIC determination was carried out in triplicate. Values are given in µg/mL. E.c: *Escherichia coli*, P. a: *Pseudomonas aeruginosa*, K.p: *Klebsiella pneumoniae*, S.e: *Salmonella enteritidis*; E.clo: *Enterobacter cloacae*, S.mar: *Serratia marcescens*, A.h: *Aeromonas hydrophila*, M.c: *Moraxella catarrhalis*, P.m: *Proteus mirabilis*, A.b: *Acinetobacter baumannii*.

| | | **DAR B** | **DAR B9** | **DAR A** | **CAZ** | **CIP** | **GEN** |
|---|---|---|---|---|---|---|---|
| *E.c* | ATCC 35218 | 1 | 1 | 8-4 | 0.25-0.125 | 0.008 | 2 |
| | ATCC 25922 | 1-0.5 | 1 | | 0.5 -0.25 | 0.008 | 1-0.5 |
| | ATCC25922 ΔTolC | 0.5-0.25 | 0.25 | | 0.5-0.25 | 0.004-0.001 | 0.5 |
| | NRZ14408 | 1-0.5 | 2-1 | 4 | 32-16 | >0.5 | >64 |
| | K0416 | 1-0.5 | 1 | 4 | 64 | >0.5 | 4 |
| | Survcare052 | 1 | 2 | 8-4 | >64 | >0.5 | 0.5 |
| *P.a* | PA01 | 1-0.5 | 1 | 4-2 | 2 | 0.25 | 2 |
| | PA103 | 8 | 4 | | 8 | 0.06 | 2 |
| | ATCC27853 | 8 | 8 | >64 | 4 | 0.125 | 2 |
| | EXT111762 | 2-1 | 2-1 | | 32-16 | 0.5 | 0.5 |
| *K.p* | DSM30104 | 2-1 | 4-2 | 4 | 0.06 | 0.06-0.03 | 0.125-0.06 |
| | ATCC 700603 | 2-1 | 4-2 | | >64 | >0.5 | 64 |
| *S.e* | ATCC 13076 | 0.5 | 0.5-0.25 | 8 | 1 | 0.008 | 0.25-0.125 |
| *E.clo* | RKI 146/09 | 1 | 2 | | >64 | 0.015 | 2 |
| *S.mar* | RKI 184_11 | 4 | 4-2 | | 16 | >0.5 | 4 |
| *A.h* | ATCC7966 | 16 | 16 | | 0.25-0.125 | 0.001 | 0.25 |
| *M.c* | ATCC 25238 | >64 | >64 | | 8 | 0.25-0.125 | 0.25 |
| *P.m* | ATCC 137050 | 64 | 64 | | 0.125 | 0.015 | 1 |
| *A. b* | ATCC 19606 | 32 | 8 | >64 | 16 | >0.5 | 32 |

Table S4. Constructs used in this study and primers used to generate them.

| | | |
|---|---|---|
| **Heptapeptide and macrocycle extension and contraction** | | |
| Construct | Modification | Primer sequence 5' -> 3' |
| pDK-2 | AWNWSKSF | aactggtcaaaaagcttctaaagcttatccca |
| | | tagaagctttttgaccagttccacgcggccgtgatctcagggatcttagg |
| pDK-3 | RWNWSKSF | tggaactggtcaaaaagcttttaaagcttatcc |
| | | aagctttttgaccagttccaacgggccgtgatctcagggatcttagg |
| pDK-4 | WNWSKSFA | taaagcttatcccatcaggttattttattttcctgaaaaaaca |
| | | acctgatgggataagctttacgcgaagctttttgaccagttccaggcc |
| pDK-5 | WNWSKSWA | taaagcttatcccatcaggttattttattttcctgaaaaaac |
| | | aataacctgatgggataagctttacgcccagctttttgaccagttccaggc |
| pDK-6 | WANWSKSF | aactggtcaaaaagcttctaaagcttatccca |
| | | agaagctttttgaccagttcgcccaggccgtgatctcaggg |
| pDK-7 | WNAWSKSF | tggtcaaaaagcttctaaagcttatcccatca |
| | | ctttagaagctttttgaccacgcgttccaggccgtgatctcagg |
| pDK-8 | WNWASKSF | tcaaaaagcttctaaagcttatcccatcaggttattttattttc |
| | | aagctttagaagctttttgacgcccagttccaggccgtgatctc |
| pDK-9 | WNWSAKSF | aaaagcttctaaagcttatcccatcaggttattttattttcc |
| | | gataagctttagaagcttttcgctgaccagttccaggccgtga |
| pDK-10 | WNWSKS- | tggaactggtcaaaaagcDDRtaaagcttatcccatcaggttattttattttcctgaaaaaac |
| | | acctgatgggataagctttaYHHgctttttgaccagttccaggccg |
| pDK-11 | WNWSK-- | tggaactggtcaaaaRWGttctaaagcttatcccatcaggttattttattttcctgaaaaaac |
| | | acctgatgggataagctttagaaCWYttttgaccagttccaggccgtg |
| pDK-12 | W - WSKSF | tggtcaaaaagcttttaaagcttatcccatcagg |
| | | ataagctttaaaagctttttgaccaccaggccgtgatctcaggga |
| pDK-13 | WNW- KSF | aaaagcttctaaagcttatcccatcaggttattttattttcc |
| | | tgggataagctttagaagcttgaccagttccaggccgtga |
| **Singe Amino acid exchange** | | |
| Construct | Modification | Primer sequence 5' -> 3' |
| pDK-W1A | ANWSKSF | NNNaactggtcaaaaagcttctaaagcttatcccatcaggttattttattttcctgaaaaaac |
| | | acctgatgggataagctttagaagctttttgaccagttNNNggccgtgatctcagggatcttagg |
| pDK-W1C | CNWSKSF | NNNaactggtcaaaaagcttctaaagcttatcccatcaggttattttattttcctgaaaaaac |
| | | acctgatgggataagctttagaagctttttgaccagttNNNggccgtgatctcagggatcttagg |
| pDK-W1D | DNWSKSF | NNNaactggtcaaaaagcttctaaagcttatcccatcaggttattttattttcctgaaaaaac |
| | | acctgatgggataagctttagaagctttttgaccagttNNNggccgtgatctcagggatcttagg |
| pDK-W1E | ENWSKSF | NNNaactggtcaaaaagcttctaaagcttatcccatcaggttattttattttcctgaaaaaac |
| | | acctgatgggataagctttagaagctttttgaccagttNNNggccgtgatctcagggatcttagg |
| pDK-W1F | FNWSKSF | NNNaactggtcaaaaagcttctaaagcttatcccatcaggttattttattttcctgaaaaaac |
| | | acctgatgggataagctttagaagctttttgaccagttNNNggccgtgatctcagggatcttagg |
| pDK-W1G | GNWSKSF | aactggtcaaaaagcttctaaagcttatccca |
| | | tagaagctttttgaccagttgccggccgtgatctcagggatcttagg |
| pDK-W1H | HNWSKSF | NNNaactggtcaaaaagcttctaaagcttatcccatcaggttattttattttcctgaaaaaac |
| | | acctgatgggataagctttagaagctttttgaccagttNNNggccgtgatctcagggatcttagg |

| | | |
|---|---|---|
| pDK-W1I | INWSKSF | NNNaactggtcaaaaagcttctaaagcttatcccatcaggttattttattttcctgaaaaaac |
| | | acctgatgggataagctttagaagctttttgaccagttNNNggccgtgatctcagggatcttagg |
| pDK-W1K | KNWSKSF | NNNaactggtcaaaaagcttctaaagcttatcccatcaggttattttattttcctgaaaaaac |
| | | acctgatgggataagctttagaagctttttgaccagttNNNggccgtgatctcagggatcttagg |
| pDK-W1L | LNWSKSF | NNNaactggtcaaaaagcttctaaagcttatcccatcaggttattttattttcctgaaaaaac |
| | | acctgatgggataagctttagaagctttttgaccagttNNNggccgtgatctcagggatcttagg |
| pDK-W1M | MNWSKSF | aactggtcaaaaagcttctaaagcttatccca |
| | | tagaagctttttgaccagttcatggccgtgatctcagggatcttagg |
| pDK-W1N | NNWSKSF | aactggtcaaaaagcttctaaagcttatccca |
| | | tagaagctttttgaccagttgttggccgtgatctcagggatcttagg |
| pDK-W1P | PNWSKSF | NNNaactggtcaaaaagcttctaaagcttatcccatcaggttattttattttcctgaaaaaac |
| | | acctgatgggataagctttagaagctttttgaccagttNNNggccgtgatctcagggatcttagg |
| pDK-W1Q | QNWSKSF | NNNaactggtcaaaaagcttctaaagcttatcccatcaggttattttattttcctgaaaaaac |
| | | acctgatgggataagctttagaagctttttgaccagttNNNggccgtgatctcagggatcttagg |
| pDK-W1R | RNWSKSF | NNNaactggtcaaaaagcttctaaagcttatcccatcaggttattttattttcctgaaaaaac |
| | | acctgatgggataagctttagaagctttttgaccagttNNNggccgtgatctcagggatcttagg |
| pDK-W1S | SNWSKSF | NNNaactggtcaaaaagcttctaaagcttatcccatcaggttattttattttcctgaaaaaac |
| | | acctgatgggataagctttagaagctttttgaccagttNNNggccgtgatctcagggatcttagg |
| pDK-W1T | TNWSKSF | NNNaactggtcaaaaagcttctaaagcttatcccatcaggttattttattttcctgaaaaaac |
| | | acctgatgggataagctttagaagctttttgaccagttNNNggccgtgatctcagggatcttagg |
| pDK-W1V | VNWSKSF | NNNaactggtcaaaaagcttctaaagcttatcccatcaggttattttattttcctgaaaaaac |
| | | acctgatgggataagctttagaagctttttgaccagttNNNggccgtgatctcagggatcttagg |
| pDK-W1Y | YNWSKSF | NNNaactggtcaaaaagcttctaaagcttatcccatcaggttattttattttcctgaaaaaac |
| | | acctgatgggataagctttagaagctttttgaccagttNNNggccgtgatctcagggatcttagg |
| pDK-N2A | WAWSKSF | tggNNNtggtcaaaaagcttctaaagcttatcccatcaggttattttattttcctgaaaaaac |
| | | acctgatgggataagctttagaagctttttgaccaNNNccaggccgtgatctcaggg |
| pDK-N2C | WCWSKSF | tggNNNtggtcaaaaagcttctaaagcttatcccatcaggttattttattttcctgaaaaaac |
| | | acctgatgggataagctttagaagctttttgaccaNNNccaggccgtgatctcaggg |
| pDK-N2D | WDWSKSF | tggtcaaaaagcttctaaagcttatcccatca |
| | | ctttagaagctttttgaccaatcccaggccgtgatctcaggga |
| pDK-N2E | WEWSKSF | tggNNNtggtcaaaaagcttctaaagcttatcccatcaggttattttattttcctgaaaaaac |
| | | acctgatgggataagctttagaagctttttgaccaNNNccaggccgtgatctcaggg |
| pDK-N2F | WFWSKSF | tggtcaaaaagcttctaaagcttatcccatca |
| | | ctttagaagctttttgaccaaaaccaggccgtgatctcaggga |
| pDK-N2G | WGWSKSF | tggtcaaaaagcttctaaagcttatcccatca |
| | | ctttagaagctttttgaccagccccaggccgtgatctcaggga |
| pDK-N2H | WHWSKSF | tggNNNtggtcaaaaagcttctaaagcttatcccatcaggttattttattttcctgaaaaaac |
| | | acctgatgggataagctttagaagctttttgaccaNNNccaggccgtgatctcaggg |
| pDK-N2I | WIWSKSF | tggNNNtggtcaaaaagcttctaaagcttatcccatcaggttattttattttcctgaaaaaac |
| | | acctgatgggataagctttagaagctttttgaccaNNNccaggccgtgatctcaggg |
| pDK-N2K | WKWSKSF | tggNNNtggtcaaaaagcttctaaagcttatcccatcaggttattttattttcctgaaaaaac |
| | | acctgatgggataagctttagaagctttttgaccaNNNccaggccgtgatctcaggg |
| pDK-N2L | WLWSKSF | tggNNNtggtcaaaaagcttctaaagcttatcccatcaggttattttattttcctgaaaaaac |

| | | |
|---|---|---|
| | | acctgatgggataagctttagaagctttttgaccaNNNccaggccgtgatctcaggg |
| pDK-N2M | WMWSKSF | tggtcaaaaagcttctaaagcttatcccatca |
| | | tagaagctttttgaccagttcatggccgtgatctcagggatcttagg |
| pDK-N2P | WPWSKSF | tggNNNtggtcaaaaagcttctaaagcttatcccatcaggttattttattttcctgaaaaaac |
| | | acctgatgggataagctttagaagctttttgaccaNNNccaggccgtgatctcaggg |
| pDK-N2Q | WQWSKSF | tggtcaaaaagcttctaaagcttatcccatca |
| | | ctttagaagctttttgaccactgccaggccgtgatctcaggga |
| pDK-N2R | WRWSKSF | tggNNNtggtcaaaaagcttctaaagcttatcccatcaggttattttattttcctgaaaaaac |
| | | acctgatgggataagctttagaagctttttgaccaNNNccaggccgtgatctcaggg |
| pDK-N2S | WSWSKSF | tggNNNtggtcaaaaagcttctaaagcttatcccatcaggttattttattttcctgaaaaaac |
| | | acctgatgggataagctttagaagctttttgaccaNNNccaggccgtgatctcaggg |
| pDK-N2T | WTWSKSF | tggNNNtggtcaaaaagcttctaaagcttatcccatcaggttattttattttcctgaaaaaac |
| | | acctgatgggataagctttagaagctttttgaccaNNNccaggccgtgatctcaggg |
| pDK-N2V | WVWSKSF | tggtcaaaaagcttctaaagcttatcccatca |
| | | ctttagaagctttttgaccacacccaggccgtgatctcaggga |
| pDK-N2W | WWWSKSF | tggtcaaaaagcttctaaagcttatcccatca |
| | | ctttagaagctttttgaccaccaccaggccgtgatctcaggga |
| pDK-N2Y | WYWSKSF | tggNNNtggtcaaaaagcttctaaagcttatcccatcaggttattttattttcctgaaaaaac |
| | | acctgatgggataagctttagaagctttttgaccaNNNccaggccgtgatctcaggg |
| pDK-W3A | WNASKSF | tggaacNNNtcaaaaagcttctaaagcttatcccatcaggttattttattttcctgaaaaaac |
| | | acctgatgggataagctttagaagctttttgaNNNgttccaggccgtgatctcagg |
| pDK-W3C | WNCSKSF | tggaacNNNtcaaaaagcttctaaagcttatcccatcaggttattttattttcctgaaaaaac |
| | | acctgatgggataagctttagaagctttttgaNNNgttccaggccgtgatctcagg |
| pDK-W3D | WNDSKSF | tcaaaaagcttctaaagcttatcccatcaggttattttattttc |
| | | aagctttagaagctttttgaatcgttccaggccgtgatctcaggg |
| pDK-W3E | WNESKSF | tggaacNNNtcaaaaagcttctaaagcttatcccatcaggttattttattttcctgaaaaaac |
| | | acctgatgggataagctttagaagctttttgaNNNgttccaggccgtgatctcagg |
| pDK-W3F | WNFSKSF | tggaacNNNtcaaaaagcttctaaagcttatcccatcaggttattttattttcctgaaaaaac |
| | | acctgatgggataagctttagaagctttttgaNNNgttccaggccgtgatctcagg |
| pDK-W3G | WNGSKSF | tcaaaaagcttctaaagcttatcccatcaggttattttattttc |
| | | aagctttagaagctttttgagccgttccaggccgtgatctcaggg |
| pDK-W3H | WNHSKSF | tggaacNNNtcaaaaagcttctaaagcttatcccatcaggttattttattttcctgaaaaaac |
| | | acctgatgggataagctttagaagctttttgaNNNgttccaggccgtgatctcagg |
| pDK-W3I | WNISKSF | tggaacNNNtcaaaaagcttctaaagcttatcccatcaggttattttattttcctgaaaaaac |
| | | acctgatgggataagctttagaagctttttgaNNNgttccaggccgtgatctcagg |
| pDK-W3K | WNKSKSF | tggaacNNNtcaaaaagcttctaaagcttatcccatcaggttattttattttcctgaaaaaac |
| | | acctgatgggataagctttagaagctttttgaNNNgttccaggccgtgatctcagg |
| pDK-W3L | WNLSKSF | tggaacNNNtcaaaaagcttctaaagcttatcccatcaggttattttattttcctgaaaaaac |
| | | acctgatgggataagctttagaagctttttgaNNNgttccaggccgtgatctcagg |
| pDK-W3M | WNMSKSF | agatcacggcctggaacatgtcaaaaagcttctaaagcttatcccatcaggt |
| | | catgttccaggccgtgatctcaggg |
| pDK-W3N | WNNSKSF | tggaacNNNtcaaaaagcttctaaagcttatcccatcaggttattttattttcctgaaaaaac |
| | | acctgatgggataagctttagaagctttttgaNNNgttccaggccgtgatctcagg |

| | | |
|---|---|---|
| pDK-W3P | WNPSKSF | tggaacNNNtcaaaaagcttctaaagcttatcccatcaggttattttattttcctgaaaaaac |
| | | acctgatgggataagctttagaagctttttgaNNNgttccaggccgtgatctcagg |
| pDK-W3Q | WNQSKSF | tggaacNNNtcaaaaagcttctaaagcttatcccatcaggttattttattttcctgaaaaaac |
| | | acctgatgggataagctttagaagctttttgaNNNgttccaggccgtgatctcagg |
| pDK-W3R | WNRSKSF | tggaacNNNtcaaaaagcttctaaagcttatcccatcaggttattttattttcctgaaaaaac |
| | | acctgatgggataagctttagaagctttttgaNNNgttccaggccgtgatctcagg |
| pDK-W3S | WNSSKSF | tggaacNNNtcaaaaagcttctaaagcttatcccatcaggttattttattttcctgaaaaaac |
| | | acctgatgggataagctttagaagctttttgaNNNgttccaggccgtgatctcagg |
| pDK-W3T | WNTSKSF | tggaacNNNtcaaaaagcttctaaagcttatcccatcaggttattttattttcctgaaaaaac |
| | | acctgatgggataagctttagaagctttttgaNNNgttccaggccgtgatctcagg |
| pDK-W3V | WNVSKSF | tggaacNNNtcaaaaagcttctaaagcttatcccatcaggttattttattttcctgaaaaaac |
| | | acctgatgggataagctttagaagctttttgaNNNgttccaggccgtgatctcagg |
| pDK-W3Y | WNYSKSF | tggaacNNNtcaaaaagcttctaaagcttatcccatcaggttattttattttcctgaaaaaac |
| | | acctgatgggataagctttagaagctttttgaNNNgttccaggccgtgatctcagg |
| pDK-S4A | WNWAKSF | tggaactggNNNaaaagcttctaaagcttatcccatcaggttattttattttcctgaaaaaac |
| | | acctgatgggataagctttagaagcttttNNNccagttccaggccgtgatctc |
| pDK-S4C | WNWCKSF | tggaactggNNNaaaagcttctaaagcttatcccatcaggttattttattttcctgaaaaaac |
| | | acctgatgggataagctttagaagcttttNNNccagttccaggccgtgatctc |
| pDK-S4D | WNWDKSF | tggaactggNNNaaaagcttctaaagcttatcccatcaggttattttattttcctgaaaaaac |
| | | acctgatgggataagctttagaagcttttNNNccagttccaggccgtgatctc |
| pDK-S4E | WNWEKSF | tggaactggNNNaaaagcttctaaagcttatcccatcaggttattttattttcctgaaaaaac |
| | | acctgatgggataagctttagaagcttttNNNccagttccaggccgtgatctc |
| pDK-S4F | WNWFKSF | aaaagcttctaaagcttatcccatcaggttattttattttcc |
| | | gataagctttagaagcttttaaaccagttccaggccgtgatctca |
| pDK-S4G | WNWGKSF | tggaactggNNNaaaagcttctaaagcttatcccatcaggttattttattttcctgaaaaaac |
| | | acctgatgggataagctttagaagcttttNNNccagttccaggccgtgatctc |
| pDK-S4H | WNWHKSF | tggaactggNNNaaaagcttctaaagcttatcccatcaggttattttattttcctgaaaaaac |
| | | acctgatgggataagctttagaagcttttNNNccagttccaggccgtgatctc |
| pDK-S4I | WNWIKSF | tggaactggNNNaaaagcttctaaagcttatcccatcaggttattttattttcctgaaaaaac |
| | | acctgatgggataagctttagaagcttttNNNccagttccaggccgtgatctc |
| pDK-S4K | WNWKKSF | tggaactggWWKaaaagcttctaaagcttatcccatcaggttattttattttcctgaaaaaac |
| | | acctgatgggataagctttagaagcttttMWWccagttccaggccgtgatctc |
| pDK-S4L | WNWLKSF | tggaactggNNNaaaagcttctaaagcttatcccatcaggttattttattttcctgaaaaaac |
| | | acctgatgggataagctttagaagcttttNNNccagttccaggccgtgatctc |
| pDK-S4M | WNWMKSF | tggaactggWWKaaaagcttctaaagcttatcccatcaggttattttattttcctgaaaaaac |
| | | acctgatgggataagctttagaagcttttMWWccagttccaggccgtgatctc |
| pDK-S4N | WNWNKSF | tggaactggNNNaaaagcttctaaagcttatcccatcaggttattttattttcctgaaaaaac |
| | | acctgatgggataagctttagaagcttttNNNccagttccaggccgtgatctc |
| pDK-S4P | WNWPKSF | tggaactggNNNaaaagcttctaaagcttatcccatcaggttattttattttcctgaaaaaac |
| | | acctgatgggataagctttagaagcttttNNNccagttccaggccgtgatctc |
| pDK-S4Q | WNWQKSF | tggaactggNNNaaaagcttctaaagcttatcccatcaggttattttattttcctgaaaaac |
| | | acctgatgggataagctttagaagcttttNNNccagttccaggccgtgatctc |
| pDK-S4R | WNWRKSF | tggaactggNNNaaaagcttctaaagcttatcccatcaggttattttattttcctgaaaaaac |

| | | |
|---|---|---|
| | | acctgatgggataagctttagaagcttttNNNccagttccaggccgtgatctc |
| pDK-S4T | WNWTKSF | tggaactggNNNaaaagcttctaaagcttatcccatcaggttattttattttcctgaaaaaac |
| | | acctgatgggataagctttagaagcttttNNNccagttccaggccgtgatctc |
| pDK-S4V | WNWVKSF | tggaactggNNNaaaagcttctaaagcttatcccatcaggttattttattttcctgaaaaaac |
| | | acctgatgggataagctttagaagcttttNNNccagttccaggccgtgatctc |
| pDK-S4W | WNWWKSF | tggaactggNNNaaaagcttctaaagcttatcccatcaggttattttattttcctgaaaaaac |
| | | acctgatgggataagctttagaagcttttNNNccagttccaggccgtgatctc |
| pDK-S4Y | WNWYKSF | tggaactggNNNaaaagcttctaaagcttatcccatcaggttattttattttcctgaaaaaac |
| | | acctgatgggataagctttagaagcttttNNNccagttccaggccgtgatctc |
| pDK-K5A | WNWSASF | tggaactggtcaNNNagcttctaaagcttatcccatcaggttattttattttcctgaaaaaac |
| | | acctgatgggataagctttagaagctNNNtgaccagttccaggccgtga |
| pDK-K5C | WNWSCSF | tggaactggtcaNNNagcttctaaagcttatcccatcaggttattttattttcctgaaaaaac |
| | | acctgatgggataagctttagaagctNNNtgaccagttccaggccgtga |
| pDK-K5D | WNWSDSF | tggaactggtcaNNNagcttctaaagcttatcccatcaggttattttattttcctgaaaaaac |
| | | acctgatgggataagctttagaagctNNNtgaccagttccaggccgtga |
| pDK-K5E | WNWSESF | agcttctaaagcttatcccatcaggttattttattttcc |
| | | tgggataagctttagaagctttctgaccagttccaggccgtga |
| pDK-K5F | WNWSFSF | tggaactggtcaNNNagcttctaaagcttatcccatcaggttattttattttcctgaaaaaac |
| | | acctgatgggataagctttagaagctNNNtgaccagttccaggccgtga |
| pDK-K5G | WNWSGSF | tggaactggtcaNNNagcttctaaagcttatcccatcaggttattttattttcctgaaaaaac |
| | | acctgatgggataagctttagaagctNNNtgaccagttccaggccgtga |
| pDK-K5H | WNWSHSF | tggaactggtcaNNNagcttctaaagcttatcccatcaggttattttattttcctgaaaaaac |
| | | acctgatgggataagctttagaagctNNNtgaccagttccaggccgtga |
| pDK-K5I | WNWSISF | tggaactggtcaNNNagcttctaaagcttatcccatcaggttattttattttcctgaaaaaac |
| | | acctgatgggataagctttagaagctNNNtgaccagttccaggccgtga |
| pDK-K5L | WNWSLSF | tggaactggtcaNNNagcttctaaagcttatcccatcaggttattttattttcctgaaaaaac |
| | | acctgatgggataagctttagaagctNNNtgaccagttccaggccgtga |
| pDK-K5M | WNWSMSF | tggaactggtcaNNNagcttctaaagcttatcccatcaggttattttattttcctgaaaaaac |
| | | acctgatgggataagctttagaagctNNNtgaccagttccaggccgtga |
| pDK-K5N | WNWSNSF | tggaactggtcaNNNagcttctaaagcttatcccatcaggttattttattttcctgaaaaaac |
| | | acctgatgggataagctttagaagctNNNtgaccagttccaggccgtga |
| pDK-K5P | WNWSPSF | tggaactggtcaNNNagcttctaaagcttatcccatcaggttattttattttcctgaaaaaac |
| | | acctgatgggataagctttagaagctNNNtgaccagttccaggccgtga |
| pDK-K5Q | WNWSQSF | tggaactggtcaNNNagcttctaaagcttatcccatcaggttattttattttcctgaaaaaac |
| | | acctgatgggataagctttagaagctNNNtgaccagttccaggccgtga |
| pDK-K5R | WNWSRSF | tggaactggtcaNNNagcttctaaagcttatcccatcaggttattttattttcctgaaaaaac |
| | | acctgatgggataagctttagaagctNNNtgaccagttccaggccgtga |
| pDK-K5S | WNWSSSF | tggaactggtcaNNNagcttctaaagcttatcccatcaggttattttattttcctgaaaaaac |
| | | acctgatgggataagctttagaagctNNNtgaccagttccaggccgtga |
| pDK-K5T | WNWSTSF | tggaactggtcaNNNagcttctaaagcttatcccatcaggttattttattttcctgaaaaaac |
| | | acctgatgggataagctttagaagctNNNtgaccagttccaggccgtga |
| pDK-K5V | WNWSVSF | agcttctaaagcttatcccatcaggttattttattttcc |
| | | tgggataagctttagaagctcactgaccagttccaggccgtga |

| | | |
|---|---|---|
| pDK-K5W | WNWSWSF | tggaactggtcaNNNagcttctaaagcttatcccatcaggttattttattttcctgaaaaaac |
| | | acctgatgggataagctttagaagctNNNtgaccagttccaggccgtga |
| pDK-K5Y | WNWSYSF | tggaactggtcaNNNagcttctaaagcttatcccatcaggttattttattttcctgaaaaaac |
| | | acctgatgggataagctttagaagctNNNtgaccagttccaggccgtga |
| pDK-S6A | WNWSKAF | tggaactggtcaaaaNNNttctaaagcttatcccatcaggttattttattttcctgaaaaaac |
| | | acctgatgggataagctttagaaNNNttttgaccagttccaggccgtg |
| pDK-S6C | WNWSKCF | tggaactggtcaaaaNNNttctaaagcttatcccatcaggttattttattttcctgaaaaaac |
| | | acctgatgggataagctttagaaNNNttttgaccagttccaggccgtg |
| pDK-S6D | WNWSKDF | tggaactggtcaaaaNNNttctaaagcttatcccatcaggttattttattttcctgaaaaaac |
| | | acctgatgggataagctttagaaNNNttttgaccagttccaggccgtg |
| pDK-S6E | WNWSKEF | ttctaaagcttatcccatcaggttattttattttcctgaaaaaac |
| | | tgatgggataagctttagaattcttttgaccagttccaggccgtga |
| pDK-S6F | WNWSKFF | tggaactggtcaaaaNNNttctaaagcttatcccatcaggttattttattttcctgaaaaaac |
| | | acctgatgggataagctttagaaNNNttttgaccagttccaggccgtg |
| pDK-S6G | WNWSKGF | tggaactggtcaaaaNNNttctaaagcttatcccatcaggttattttattttcctgaaaaaac |
| | | acctgatgggataagctttagaaNNNttttgaccagttccaggccgtg |
| pDK-S6H | WNWSKHF | tggaactggtcaaaaNNNttctaaagcttatcccatcaggttattttattttcctgaaaaaac |
| | | acctgatgggataagctttagaaNNNttttgaccagttccaggccgtg |
| pDK-S6I | WNWSKIF | tggaactggtcaaaaNNNttctaaagcttatcccatcaggttattttattttcctgaaaaaac |
| | | acctgatgggataagctttagaaNNNttttgaccagttccaggccgtg |
| pDK-S6K | WNWSKKF | tggaactggtcaaaaNNNttctaaagcttatcccatcaggttattttattttcctgaaaaaac |
| | | acctgatgggataagctttagaaNNNttttgaccagttccaggccgtg |
| pDK-S6L | WNWSKLF | tggaactggtcaaaaNNNttctaaagcttatcccatcaggttattttattttcctgaaaaaac |
| | | acctgatgggataagctttagaaNNNttttgaccagttccaggccgtg |
| pDK-S6M | WNWSKMF | ttctaaagcttatcccatcaggttattttattttcctgaaaaaac |
| | | tgatgggataagctttagaacatttttgaccagttccaggccgtga |
| pDK-S6N | WNWSKNF | tggaactggtcaaaaNNNttctaaagcttatcccatcaggttattttattttcctgaaaaaac |
| | | acctgatgggataagctttagaaNNNttttgaccagttccaggccgtg |
| pDK-S6P | WNWSKPF | tggaactggtcaaaaNNNttctaaagcttatcccatcaggttattttattttcctgaaaaaac |
| | | acctgatgggataagctttagaaNNNttttgaccagttccaggccgtg |
| pDK-S6Q | WNWSKQF | tggaactggtcaaaaNNNttctaaagcttatcccatcaggttattttattttcctgaaaaaac |
| | | acctgatgggataagctttagaaNNNttttgaccagttccaggccgtg |
| pDK-S6R | WNWSKRF | tggaactggtcaaaaNNNttctaaagcttatcccatcaggttattttattttcctgaaaaaac |
| | | acctgatgggataagctttagaaNNNttttgaccagttccaggccgtg |
| pDK-S6T | WNWSKTF | tggaactggtcaaaaNNNttctaaagcttatcccatcaggttattttattttcctgaaaaaac |
| | | acctgatgggataagctttagaaNNNttttgaccagttccaggccgtg |
| pDK-S6V | WNWSKVF | tggaactggtcaaaaNNNttctaaagcttatcccatcaggttattttattttcctgaaaaaac |
| | | acctgatgggataagctttagaaNNNttttgaccagttccaggccgtg |
| pDK-S6W | WNWSKWF | tggaactggtcaaaaNNNttctaaagcttatcccatcaggttattttattttcctgaaaaaac |
| | | acctgatgggataagctttagaaNNNttttgaccagttccaggccgtg |
| pDK-S6Y | WNWSKYF | tggaactggtcaaaaNNNttctaaagcttatcccatcaggttattttattttcctgaaaaaac |
| | | acctgatgggataagctttagaaNNNttttgaccagttccaggccgtg |
| pDK-F7A | WNWSKSA | tggaactggtcaaaaagcDDRtaaagcttatcccatcaggttattttattttcctgaaaaaac |

| | | |
|---|---|---|
| | | acctgatgggataagctttaYHHgctttttgaccagttccaggccg |
| pDK-F7C | WNWSKSC | tggaactggtcaaaaagcDDRtaaagcttatcccatcaggttattttattttcctgaaaaaac |
| | | acctgatgggataagctttaYHHgctttttgaccagttccaggccg |
| pDK-F7D | WNWSKSD | tggaactggtcaaaaagcDDRtaaagcttatcccatcaggttattttattttcctgaaaaaac |
| | | acctgatgggataagctttaYHHgctttttgaccagttccaggccg |
| pDK-F7E | WNWSKSE | tggaactggtcaaaaagcDDRtaaagcttatcccatcaggttattttattttcctgaaaaaac |
| | | acctgatgggataagctttaYHHgctttttgaccagttccaggccg |
| pDK-F7G | WNWSKSG | tggaactggtcaaaaagcDDRtaaagcttatcccatcaggttattttattttcctgaaaaaac |
| | | acctgatgggataagctttaYHHgctttttgaccagttccaggccg |
| pDK-F7H | WNWSKSH | tggaactggtcaaaaagcDDRtaaagcttatcccatcaggttattttattttcctgaaaaaac |
| | | acctgatgggataagctttaYHHgctttttgaccagttccaggccg |
| pDK-F7I | WNWSKSI | tggaactggtcaaaaagcDDRtaaagcttatcccatcaggttattttattttcctgaaaaaac |
| | | acctgatgggataagctttaYHHgctttttgaccagttccaggccg |
| pDK-F7K | WNWSKSK | tggaactggtcaaaaagcDDRtaaagcttatcccatcaggttattttattttcctgaaaaaac |
| | | acctgatgggataagctttaYHHgctttttgaccagttccaggccg |
| pDK-F7L | WNWSKSL | tggaactggtcaaaaagcDDRtaaagcttatcccatcaggttattttattttcctgaaaaaac |
| | | acctgatgggataagctttaYHHgctttttgaccagttccaggccg |
| pDK-F7M | WNWSKSM | taaagcttatcccatcaggttattttattttcctgaaaaaaca |
| | | acctgatgggataagctttacatgctttttgaccagttccaggccg |
| pDK-F7N | WNWSKSN | tggaactggtcaaaaagcDDRtaaagcttatcccatcaggttattttattttcctgaaaaaac |
| | | acctgatgggataagctttaYHHgctttttgaccagttccaggccg |
| pDK-F7P | WNWSKSP | tggaactggtcaaaaagcDDRtaaagcttatcccatcaggttattttattttcctgaaaaaac |
| | | acctgatgggataagctttaYHHgctttttgaccagttccaggccg |
| pDK-F7Q | WNWSKSQ | tggaactggtcaaaaagcDDRtaaagcttatcccatcaggttattttattttcctgaaaaaac |
| | | acctgatgggataagctttaYHHgctttttgaccagttccaggccg |
| pDK-F7R | WNWSKSR | tggaactggtcaaaaagcDDRtaaagcttatcccatcaggttattttattttcctgaaaaaac |
| | | acctgatgggataagctttaYHHgctttttgaccagttccaggccg |
| pDK-F7S | WNWSKSS | tggaactggtcaaaaagcDDRtaaagcttatcccatcaggttattttattttcctgaaaaaac |
| | | acctgatgggataagctttaYHHgctttttgaccagttccaggccg |
| pDK-F7T | WNWSKST | tggaactggtcaaaaagcDDRtaaagcttatcccatcaggttattttattttcctgaaaaaac |
| | | acctgatgggataagctttaYHHgctttttgaccagttccaggccg |
| pDK-F7V | WNWSKSV | tggaactggtcaaaaagcDDRtaaagcttatcccatcaggttattttattttcctgaaaaaac |
| | | acctgatgggataagctttaYHHgctttttgaccagttccaggccg |
| pDK-F7W | WNWSKSW | taaagcttatcccatcaggttattttattttcctgaaaaaaca |
| | | acctgatgggataagctttaccagctttttgaccagttccaggccg |
| pDK-F7Y | WNWSKSY | tggaactggtcaaaaagcDDRtaaagcttatcccatcaggttattttattttcctgaaaaaac |
| | | acctgatgggataagctttaYHHgctttttgaccagttccaggccg |
| | verification sequencing | ccataccgcgaaaggttttgcg<br>aacgtttcatggattctgagatgttaatagcattcat |
| ***ΔbamB* strain creation** | | |
| Construct | | Primer sequence 5' -> 3' |
| bamB-strep-F | | cctgtccaggagccgttttcaaagtgaacgacagagacgaaggctggagctgcttcgaag |
| bamB-strep-R | | cagatgaaaattaataatttgtccatctgagagggacccggatccgtcgacctgcagttc |

| | | |
|---|---|---|
| bamB-testF2 | | caaagtgaacgacagagacg |
| dbamBtest-R | | caaggtgcgcgtagtgcatg |
| **Focussed randomized library** | | |
| Construct | amplified fragment | Primer sequence 5' -> 3' |
| pNB04 | darA-darB | gtttaactttaataaggagatataccatgcataataccttaaatga |
| | | tgctgctgacctgaaactgggtaac |
| pNB04 | darC-darE | acccagtttcaggtcagcagcaaag |
| | | tgctcagcggtggcagcagcttacgccgcgatggtttgtt |
| rand_Phe7 | darA | ccactgacgcgttgcgcgag |
| | | ggaccgccgcaagctttagaaMNNcttMNNccaMNNccaggccgtgatctcagggatct |
| rand_Trp7 | darA | ccactgacgcgttgcgcgag |
| | | ggaccgccgcaagctttaccaMNNcttMNNccaMNNccaggccgtgatctcagggatct |
| rand_lib_Phe7 | randomized darA-Phe7 | ccactgacgcgttgcgcgag<br>ggaccgccgcaagcttta |
| rand_lib_Trp7 | randomized darA-Trp7 | ccactgacgcgttgcgcgag<br>ggaccgccgcaagcttta |
| amplicon_seq | master_library (randomized darA-Phe7/Trp7 pooled) | tcgtcggcagcgtcagatgtgtataagagacaggcatcattcaaagagactgaactctc<br>gtctcgtgggctcggagatgtgtataagagacagtgaacaacttgattgtttatcccaatgg |

Table S5. Bicyclic heptapeptides produced by heterologous expression of the darobactin biosynthetic gene cluster. The residues of the first member of this family, darobactin A, are indicated by DAR A; All seven DarA precursor amino acids were exchanged one by one to all other canonical ones, + indicates that the respective heptapeptide was detected by UPLC-HRMS; - bicyclic heptapeptides were not detected. Clones synthesizing the green-colored heptapeptides produced a halo in the competition assays against E. coli MG1655 *bamB::aac(3)*IV. Clones producing the blue-colored ones did not inhibit the test strain.

| Description | | $W_1$ | $N_2$ | $W_3$ | $S_4$ | $K_5$ | $S_6$ | $F_7$ |
|---|---|---|---|---|---|---|---|---|
| Nucleophilic | C | - | - | - | + | - | + | + |
| | S | - | + | - | DAR A | - | DAR A | + |
| | T | - | + | - | + | - | + | + |
| Hydrophobic | I | - | + | - | + | - | + | + |
| | L | - | + | - | + | - | + | + |
| | M | - | + | - | + | - | + | - |
| | P | - | - | - | - | - | - | + |
| | V | - | + | - | + | - | + | + |
| Amide | N | - | DAR A | - | + | - | + | + |
| | Q | - | + | - | + | - | + | + |
| Basic | H | - | + | - | + | - | + | + |
| | K | - | + | - | + | DAR A | + | + |
| | R | - | + | - | + | + | + | + |
| Small | A | - | + | - | + | - | + | + |
| | G | - | - | - | + | - | + | + |
| Acidic | D | - | - | - | + | - | + | + |
| | E | - | + | - | + | - | + | + |
| Aromatic | F | - | - | - | + | - | + | DAR A |
| | Y | - | + | + | + | - | + | + |
| | W | DAR A | + | DAR A | - | - | - | + |

Sequence S1. Sequence of apramycin resistance gene with oriT flanked with FRT sites  (5' -> 3')

GATCCGTCGACCTGCAGTTCGAAGTTCCTATTCTCTAGAAAGTATAGGAACTTCGAAGTTCCCGCCAGCCTCGC
AGAGCAGGATTCCCGTTGAGCACCGCCAGGTGCGAATAAGGGACAGTGAAGAAGGAACACCCGCTCGCGGG
TGGGCCTACTTCACCTATCCTGCCCGGCTGACGCCGTTGGATACACCAAGGAAAGTCTACACGAACCCTTTGGC
AAAATCCTGTATATCGTGCGAAAAAGGATGGATATACCGAAAAAATCGCTATAATGACCCCGAAGCAGGGTTA
TGCAGCGGAAAATGCAGCTCACGGTAACTGATGCCGTATTTGCAGTACCAGCGTACGGCCCACAGAATGATGT
CACGCTGAAAATGCCGGCCTTTGAATGGGTTCATGTGCAGCTCCATCAGCAAAAGGGGATGATAAGTTTATCA
CCACCGACTATTTGCAACAGTGCCGTTGATCGTGCTATGATCGACTGATGTCATCAGCGGTGGAGTGCAATGTC
GTGCAATACGAATGGCGAAAAGCCGAGCTCATCGGTCAGCTTCTCAACCTTGGGGTTACCCCCGGCGGTGTGC
TGCTGGTCCACAGCTCCTTCCGTAGCGTCCGGCCCCTCGAAGATGGGCCACTTGGACTGATCGAGGCCCTGCG
TGCTGCGCTGGGTCCGGGAGGGACGCTCGTCATGCCCTCGTGGTCAGGTCTGGACGACGAGCCGTTCGATCCT
GCCACGTCGCCCGTTACACCGGACCTTGGAGTTGTCTCTGACACATTCTGGCGCCTGCCAAATGTAAAGCGCA
GCGCCCATCCATTTGCCTTTGCGGCAGCGGGGCCACAGGCAGAGCAGATCATCTCTGATCCATTGCCCCTGCCA
CCTCACTCGCCTGCAAGCCCGGTCGCCCGTGTCCATGAACTCGATGGGCAGGTACTTCTCCTCGGCGTGGGAC
ACGATGCCAACACGACGCTGCATCTTGCCGAGTTGATGGCAAAGGTTCCCTATGGGGTGCCGAGACACTGCAC
CATTCTTCAGGATGGCAAGTTGGTACGCGTCGATTATCTCGAGAATGACCACTGCTGTGAGCGCTTTGCCTTGG
CGGACAGGTGGCTCAAGGAGAAGAGCCTTCAGAAGGAAGGTCCAGTCGGTCATGCCTTTGCTCGGTTGATCC
GCTCCCGCGACATTGTGGCGACAGCCCTGGGTCAACTGGGCCGAGATCCGTTGATCTTCCTGCATCCGCCAGA
GGCGGGATGCGAAGAATGCGATGCCGCTCGCCAGTCGATTGGCTGAGCTCATAAGTTCCTATTCCGAAGTTCC
TATTCTCTAGAAAGTATAGGAACTTCGAAGCAGCTCCAGCCT

Figure S1: Alignment of sequencing result of E. coli bamA6ΔbamB to apramycin resistance cassette

```
apramycin       AGGCTGGAGCTGCTTCGAAGTTCCTATACTTTCTAGAGAATAGGAACTTCGGAATAGGAA    60
sequencing      -----------------------------------------------------------G    1

apramycin       CTTATGAGCTCAGCCAATCGACTGGCGAGCGGCATCGCATTCTTCGCATCCCGCCTCTGG    120
sequencing      CTTATGAGCTCAGCCAATCGACTGGCGAGCGGCATCGCATTCTTCGCATCCCGCCTCTGG    61
                ************************************************************

apramycin       CGGATGCAGGAAGATCAACGGATCTCGGCCCAGTTGACCCAGGGCTGTCGCCACAATGTC    180
sequencing      CGGATGCAGGAAGATCAACGGATCTCGGCCCAGTTGACCCAGGGCTGTCGCCACAATGTC    121
                ************************************************************

apramycin       GCGGGAGCGGATCAACCGAGCAAAGGCATGACCGACTGGACCTTCCTTCTGAAGGCTCTT    240
sequencing      GCGGGAGCGGATCAACCGAGCAAAGGCATGACCGACTGGACCTTCCTTCTGAAGGCTCTT    181
                ************************************************************

apramycin       CTCCTTGAGCCACCTGTCCGCCAAGGCAAAGCGCTCACAGCAGTGGTCATTCTCGAGATA    300
sequencing      CTCCTTGAGCCACCTGTCCGCCAAGGCAAAGCGCTCACAGCAGTGGTCATTCTCGAGATA    241
                ************************************************************

apramycin       ATCGACGCGTACCAACTTGCCATCCTGAAGAATGGTGCAGTGTCTCGGCACCCCATAGGG    360
sequencing      ATCGACGCGTACCAACTTGCCATCCTGAAGAATGGTGCAGTGTCTCGGCACCCCATAGGG    301
                ************************************************************

apramycin       AACCTTTGCCATCAACTCGGCAAGATGCAGCGTCGTGTTGGCATCGTGTCCCACGCCGAG    420
sequencing      AACCTTTGCCATCAACTCGGCAAGATGCAGCGTCGTGTTGGCATCGTGTCCCACGCCGAG    361
                ************************************************************

apramycin       GAGAAGTACCTGCCCATCGAGTTCATGGACACGGGCGACCGGGCTTGCAGGCGAGTGAGG    480
sequencing      GAGAAGTACCTGCCCATCGAGTTCATGGACACGGGCGACCGGGCTTGCAGGCGAGTGAGG    421
                ************************************************************

apramycin       TGGCAGGGGCAATGGATCAGAGATGATCTGCTCTGCCTGTGGCCCCGCTGCCGCAAAGGC    540
sequencing      TGGCAGGGGCAATGGATCAGAGATGATCTGCTCTGCCTGTGGCCCCGCTGCCGCAAAGGC    481
                ************************************************************

apramycin       AAATGGATGGGCGCTGCGCTTTACATTTGGCAGGCGCCAGAATGTGTCAGAGACAACTCC    600
sequencing      AAATGGATGGGCGCTGCGCTTTACATTTGGCAGGCGCCAGAATGTGTCAGAGACAACTCC    541
                ************************************************************

apramycin       AAGGTCCGGTGTAACGGGCGACGTGGCAGGATCGAACGGCTCGTCGTCCAGACCTGACCA    660
sequencing      AAGGTCCGGTGTAACGGGCGACGTGGCAGGATCGAACGGCTCGTCGTCCAGACCTGACCA    601
                ************************************************************

apramycin       CGAGGGCATGACGAGCGTCCCTCCCGGACCCAGCGCAGCACGCAGGGCCTCGATCAGTCC    720
sequencing      CGAGGGCATGACGAGCGTCCCTCCCGGACCCAGCGCAGCACGCAGGGCCTCGATCAGTCC    661
                ************************************************************

apramycin       AAGTGGCCCATCTTCGAGGGGCCGGACGCTACGGA-AGGAGCTGTGGACCAGCAGCACAC    779
sequencing      AAGTGGCCCATCTTCGAGGGGCCGGACGCTACGGAAAGGAGCTGTGGACCAGCAGCACAC    721
                *********************************** ************************

apramycin       CGCCGGGGGTAACCCCAAGGTTGAGAAGCTGACCGATGAGCTCGGCTTTTCGCCATTCGT    839
sequencing      CGCCCGGGGTAACCCCAAGGTTGAGAAGCTGACCGATGAGCTCGGCTTTTCGCCATTCGT    781
                **** *******************************************************

apramycin       ATTGCACGACATTGCACTCCACCGCTGATGACATCAGTCGATCATAGCACGATCAACGGC    899
sequencing      ATTGCACGACATTGCACTCCACCGCTGATGACATCAGTCGATCATAGCACGATCAACGGC    841
                ************************************************************

apramycin       ACTGTTGCAAATAGTC-GGTGGTGATAAACTTATCATCCCCTTTTGCTGATGGAGCTGCA    958
sequencing      ACTGTTGCAAATAGTCCGGTGGTGATAAACTTATCATCCCCTTTTGCTTATGGAGCTGCA    901
                **************** ******************************* ***********

apramycin       CATGAACCCATTCAAAGGCCGGCATTTTCAGCGTGACATCATTCTGTGGGCCGTACGCTG    1018
sequencing      CATGAACCCATTCAAAGGCCGGCATTTTCAGCGTGACATCATTCTGTGGGCCGTACGCTG    961
                ************************************************************

apramycin       GTACTGCAAATACGGCATCAGTTACCGTGAGCTGCATTTTCCGCTGCATAACCCTGCTTC    1078
sequencing      GTACTGCAAATACGGCATCAGTTACCGTGAGCTGCATTTCCGCTGCAAACCC---TGCTT    1018
                *************************************** *      *   *      * 

apramycin       GGGGTCATTATAGCGATTTTTTCGGTATATCCATCCTTTTTCGCACGATATACAGGATTT    1138
sequencing      CGGGTCATTATAGCGATTTTTTCGGTWWTCCTCCTTTTCCA----CGATWTAC--GGATT    1072
                 *************************    *     **       **** ***  *  **

apramycin       TGCCAAAGGGTTCGTGTAGACTTTCCTTGGTGTATCCAACGGCGTCAGCCGGGCAGGATA    1198
sequencing      TGCCAAAGGTTC--TGTAAMCTTTCCTG--------------------------------    1098
                ********* *   ****  *******

apramycin       GGTGAAGTAGGCCCACCCGCGAGCGGGTGTTCCTTCTTCACTGTCCCTTATTCGCACCTG    1258
sequencing      ------------------------------------------------------------    1098

apramycin       GCGGTGCTCAACGGGAATCCTGCTCTGCGAGGCTGGCGGGAACTTCGAAGTTCCTATACT    1318
sequencing      ------------------------------------------------------------    1098

apramycin       TTCTAGAGAATAGGAACTTCGAACTGCAGGTCGACGGATC        1358
sequencing      ----------------------------------------        1098
```

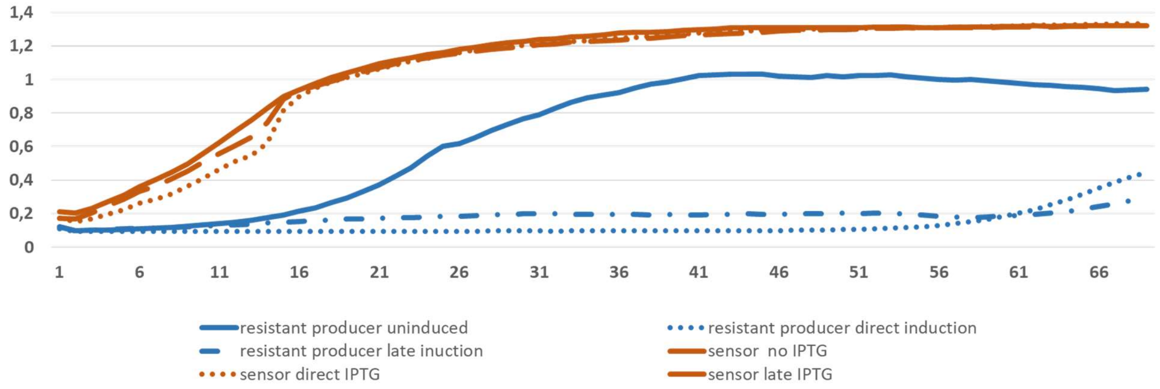

Figure S2 Comparison of growth phenotypes of sensor strain and resistant producer strain in 96 well plate: Blue: resistant producer (*E. coli* BAP1-daroR-pZW-ADC5/pRSF: Darobactin BGC from *P. khanii* HGB1456, T7-lac promoter ); Brown: sensor strain (*E. coli* MG1655-BamA6-ΔBamB) Solid line no induction: Dotted line direct induction with 0.5 mM IPTG; Dashed line induction after 3.5 h with 0.5 mM IPTG.Strains were inoculated to approximately $OD_{600}$ 0.2 in 200 ul LB Kan50 and incubated at 30 °C in TECAN infinite 200Pro plate reader for 70 h while shaking.

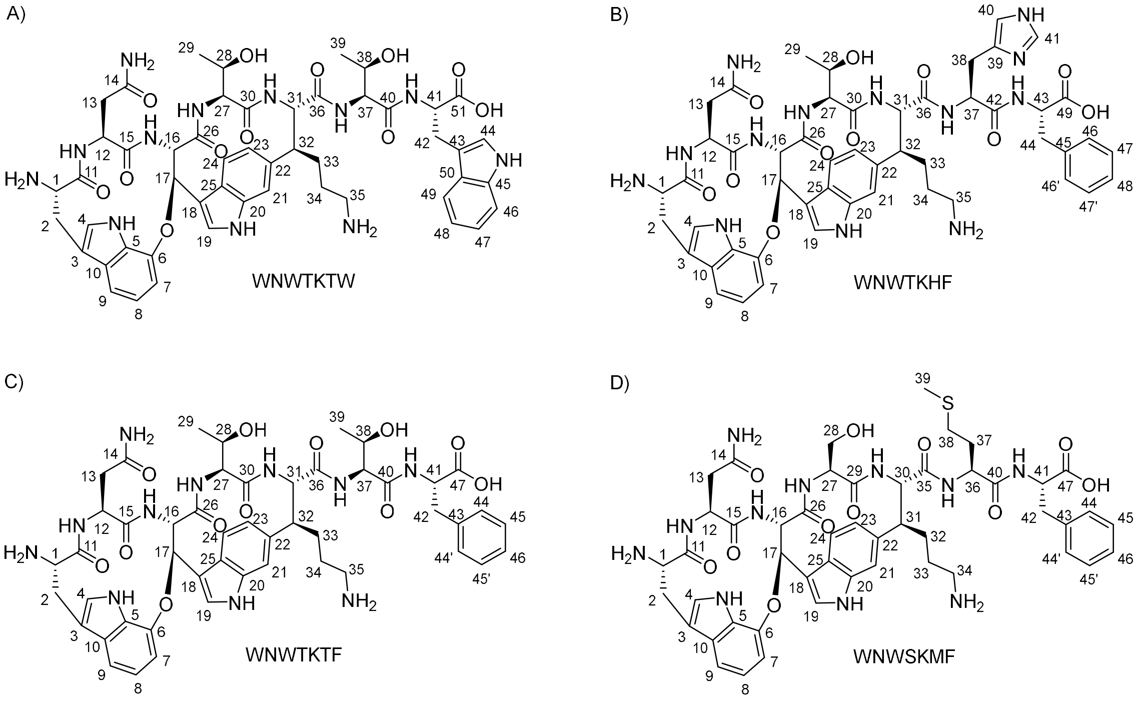

Figure S3: Structures of A) WNWTKTW, B) WNWTKHF, C) WNWTKTF, and D) WNWSKMF, including atom numbering.

Table S6: $^{1}H$ (700 MHz) and $^{13}C$ (175 MHz) NMR data of WNWTKTW ($D_2O$; δ in ppm). For $^{13}C$ measurements 3-(trimethylsilyl)propionic-2,2,3,3-$d_4$ acid sodium salt (TSPA) was used as external standard. The following abbreviations are used in this table: mult.: multiplicity, int.: integral, obs.: obscured, $C_q$: quaternary carbon atom. For atom numbering, cf. Figure S3.

| Position | $\delta_C$ [ppm], Type | $\delta_H$ [ppm], mult. (*J* in Hz), int.[a] | Position | $\delta_C$ [ppm], Type | $\delta_H$ [ppm], mult. (*J* in Hz), int.[a] |
|---|---|---|---|---|---|
| 1 | 59.1, CH | 4.05, dd (7.5, 10.6), 1H | 27 | 62.6, CH | 3.71, d (6.5), 1H |
| 2 | 30.8, $CH_2$ | 3.57, dd (7.1, 13.3), 1H<br>3.32, m, 1H[b] | 28 | 72.5, CH | 3.40, p (6.4), 1H |
| 3 | 112.6, $C_q$ | | 29 | 22.9, $CH_3$ | 0.80, d (6.3), 3H |
| 4 | 129.20 / 129.19,[c] CH | 7.37, s, 1H | 30 | 172.5, $C_q$ | |
| 5 | 133.5, $C_q$ | | 31 | 64.8, CH | 4.00, d (10.6), 1H |
| 6 | 149.6, $C_q$ | | 32 | 52.3, CH | 2.94, td (3.4, 11.0), 1H |
| 7 | 113.5, CH | 7.26, m, 1H[d] | 33 | 30.0, $CH_2$ | 1.43, m, 1H[h]<br>1.31, m, 1H |
| 8 | 124.7, CH | 7.20, t (7.6), 1H | 34 | 29.9, $CH_2$ | 1.69, m, 1H[i]<br>1.50, m, 1H[i] |
| 9 | 118.3, CH | 7.24, m, 1H[d] | 35 | 43.6, $CH_2$ | 2.73, m, 1H<br>2.60, m, 1H |
| 10 | 133.6, $C_q$ | | 36 | 176.4, $C_q$ | |
| 11 | 172.7, $C_q$ | | 37 | 63.3, CH | 4.37, d (3.4), 1H |
| 12 | 55.3, CH | 3.34, m, 1H[b] | 38 | 71.0, CH | 4.32, m, 1H |
| 13 | 43.3, $CH_2$ | 2.17, m, 2H[e] | 39 | 23.3, $CH_3$ | 1.06, d (6.4), 3H |
| 14 | 178.2, $C_q$ | | 40 | 174.7, $C_q$ | |
| 15 | 173.0, $C_q$ | | 41 | 60.1, CH | 4.62, t (5.2), 1H |
| 16 | 67.9, CH | 4.72, d (9.0), 1H | 42 | 32.0, $CH_2$ | 3.33, m, 1H[b] |
| 17 | 81.4, CH | 6.21, d (9.0), 1H | 43 | 114.4, $C_q$ | |
| 18 | 116.1, $C_q$ | | 44 | 129.20 / 129.19,[c] CH | 7.31, s, 1H |
| 19 | 128.9, CH | 7.87, s, 1H | 45 | 140.8, $C_q$ | |
| 20 | 141.6, $C_q$ | | 46 | 116.6, CH | 7.67, d (8.1), 1H |
| 21 | 115.0, CH | 7.38, m, 1H[f] | 47 | 126.7, CH | 7.39, m, 1H[f] |
| 22 | 137.4, $C_q$ | | 48 | 123.8, CH | 7.24, m, 1H[d] |
| 23 | 129.34 / 129.32,[g] CH | 6.91, d (8.2), 1H | 49 | 123.5, CH | 7.76, d (7.9), 1H |
| 24 | 121.9, CH | 7.44, d (8.1), 1H | 50 | 132.1, $C_q$ | |
| 25 | 129.34 / 129.32,[g] $C_q$ | | 51 | 182.1, $C_q$ | |
| 26 | 172.6, $C_q$ | | | | |

[a] The $^{1}$H shifts of multiplets were extracted from the HSQC spectrum. [b] The observed integral for this signal was 4H due to the overlap of the proton signals for H-2, H-12, and H-42. Thus, for each of these positions an integral of 1H was assigned. [c] Due to signal overlap of the 2D correlation signals, the $^{13}$C shifts for C-4, and C-44 cannot be distinguished. [d] The observed integral for this signal was 3H due to the overlap of the proton signals for H-7, H-9, and H-48. Thus, for each of these positions an integral of 1H was assigned. [e] The observed integral for this signal was 3H due to the overlay of the proton signal for H-13 with an impurity. The expected integral for H-13 is 2H. [f] The observed integral for this signal was 2H due to the overlap of the proton signals for H-21, and H-47. Thus, for each of these positions an integral of 1H was assigned. [g] Due to signal overlap of the 2D correlation signals, the $^{13}$C shifts for C-23, and C-25 cannot be distinguished. [h] The observed integral for this signal was 2H due to the overlap of the proton signal for H-33 with an impurity. For H-33 two signals with an integral of 1H each are expected. [i] The observed integral for this signal was 2H due to the overlap of the proton signal for H 34 with an impurity. For H 34 two signals with an integral of 1H each are expected.

**WNWTKTW**

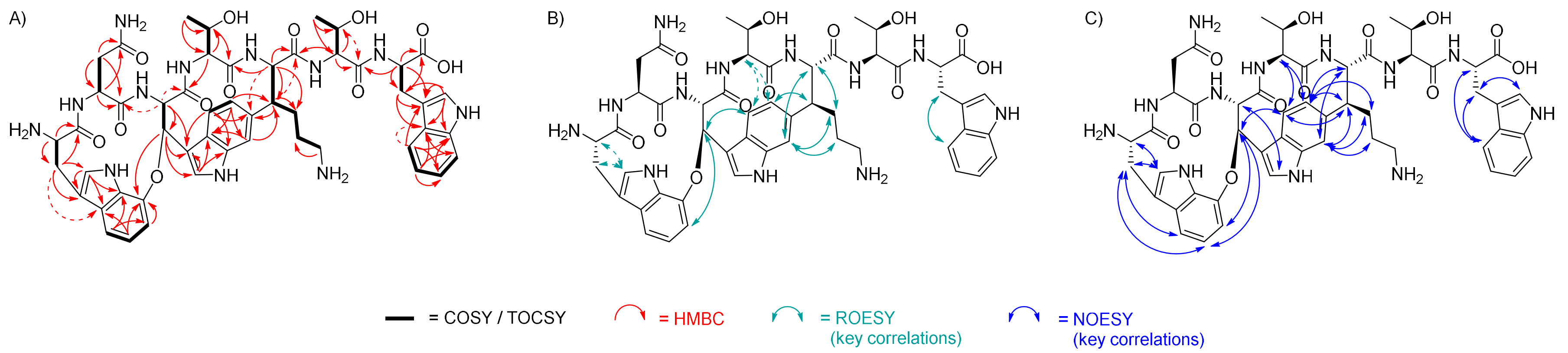

Figure S4: Key COSY/TOCSY (A, bold bonds), key HMBC (A, red arrows), key ROESY (B, turquoise arrows), and key NOESY (C, blue arrows) correlations of **WNWTKTW**. Dashed arrows indicate weak correlations.

Table S7: $^{1}H$ (700 MHz) and $^{13}C$ (175 MHz) NMR data of WNWTKHF ($D_2O$; δ in ppm). For $^{13}C$ measurements 3-(trimethylsilyl)propionic-2,2,3,3-$d_4$ acid sodium salt (TSPA) was used as external standard. The following abbreviations are used in this table: mult.: multiplicity, int.: integral, obs.: obscured, $C_q$: quaternary carbon atom. For atom numbering, cf. Figure S3.

| Position | $\delta_C$ [ppm], Type | $\delta_H$ [ppm], mult. (*J* in Hz), int.[a] | Position | $\delta_C$ [ppm], Type | $\delta_H$ [ppm], mult. (*J* in Hz), int.[a] |
|---|---|---|---|---|---|
| 1 | 59.1, CH | 4.04, dd (7.5, 10.5), 1H | 26 | 172.72 / 172.66,[f] $C_q$ | |
| 2 | 30.8, $CH_2$ | 3.57, dd (7.3, 13.6), 1H<br>3.32, m, 1H[b] | 27 | 62.5, CH | 3.70, d (6.2), 1H |
| 3 | 112.6, $C_q$ | | 28 | 72.5, CH | 3.36, p (obs),[i] 1H[b] |
| 4 | 129.2, CH | 7.36, s, 1H | 29 | 22.8, $CH_3$ | 0.72, d (6.2), 3H |
| 5 | 133.57 / 133.50,[c] $C_q$ | | 30 | 172.4, $C_q$ | |
| 6 | 149.6, $C_q$ | | 31 | 64.5, CH | 4.15, d (10.5), 1H |
| 7 | 113.5, CH | 7.26, d (obs; 7.6), 1H[d] | 32 | 52.6, CH | 2.99, td (obs; 3.1, 11.3),[j] 1H |
| 8 | 124.7, CH | 7.20, m (obs),[e] 1H | 33 | 30.2, $CH_2$ | 2.03, m, 1H<br>1.59, m, 1H |
| 9 | 118.3, CH | 7.25, d (obs; 7.9) 1H[d] | 34 | 30.0, $CH_2$ | 1.88, m, 1H<br>1.72, m, 1H |
| 10 | 133.57 / 133.50,[c] $C_q$ | | 35 | 43.9, $CH_2$ | 2.97, pseudo-t (7.4), 2H |
| 11 | 172.72 / 172.66,[f] $C_q$ | | 36 | 175.8, $C_q$ | |
| 12 | 55.3, CH | 3.33, m, 1H[b] | 37 | 56.9, CH | 4.69, m, 1H[g] |
| 13 | 43.3, $CH_2$ | 2.17, m, 2H | 38 | 31.1, $CH_2$ | 3.20, m, 1H[k]<br>3.10, dd (8.1, 15.5), 1H |
| 14 | 178.2, $C_q$ | | 39 | 132.7, $C_q$ | |
| 15 | 173.0, $C_q$ | | 40 | 121.7, CH | 7.18, s, 1H |
| 16 | 67.8, CH | 4.71, m, 1H[g] | 41 | 138.2, CH | 8.50, s,[l] 1H |
| 17 | 81.4, CH | 6.20, d (9.0), 1H | 42 | 175.1, $C_q$ | |
| 18 | 116.1, $C_q$ | | 43 | 61.0, CH | 4.49, dd (5.2, 7.2),[m] 1H |
| 19 | 128.9, CH | 7.86, s, 1H | 44 | 41.7, $CH_2$ | 3.22, m, 1H[k]<br>3.05, dd (8.0, 14.0), 1H |
| 20 | 141.6, $C_q$ | | 45 | 142.0, $C_q$ | |
| 21 | 115.0, CH | 7.45, s, 1H | 46, 46’ | 134.0, CH | 7.31, d (7.8), 2H |
| 22 | 137.4, $C_q$ | | 47, 47’ | 133.2, CH | 7.43, m, 2H[h] |
| 23 | 129.4, CH | 6.93, d (8.3), 1H | 48 | 131.5, CH | 7.37, m (obs),[n] 1H |
| 24 | 121.9, CH | 7.44, m, 1H[h] | 49 | 181.8, $C_q$ | |
| 25 | 129.3, $C_q$ | | | | |

[a] The $^{1}$H shifts of multiplets were extracted from the HSQC spectrum. [b] The observed integral for this signal was 3H due to the overlap of the proton signals for H-2, H-12, and H-28. Thus, for each of these positions an integral of 1H was assigned. [c] Due to signal overlap of the 2D correlation signals, the $^{13}$C shifts for C-5, and C-10 cannot be distinguished. [d] According to the HSQC data, the peaks for H-7, and H-9 were interpreted as partially overlapping doublets. Due to the overlap, the observed integral for this signal was 2H, so that an integral of 1H each was assigned for H-7, and H-9. [e] Even though the $^{1}$H signal of H8 appears on first glance to be a doublet, the HSQC data indicates-, that a part of this peak is obscured by the neighboring signal. Therefore, the $^{1}$H signal of H-8 was interpreted as multiplet. [f] Due to signal overlap of the 2D correlation signals, the $^{13}$C shifts for C-11, and C-26 cannot be distinguished. [g] The observed integral for this signal was 2H due to the overlap of the proton signals for H-16, and H-37. Thus, for each of these positions an integral of 1H was assigned. [h] The observed integral for this signal was 3H due to the overlap of the proton signals for H-24, H-47, and H-47’. Consequently, for H-24 an integral of 1H was assigned, while for H-47 / H-47’ an integral of 2H was assigned. [i] The pentet (p) for H-28 is partly obscured in the spectrum of **WNWTKHF**, but can be inferred from comparison with the spectrum of **WNWTKTW** (Table **S3-1, Figure SXX-8**). [j] The triplet of doublet (td) for H-32 is partly obscured in the spectrum of **WNWTKHF**, but can be inferred from comparison with the spectrum of **WNWTKTW** (Table **S3-1, Figure SXX-8**). Coupling constants were calculated for the visible part of the multiplet. [k] The observed integral for this signal was 2H due to the overlap of the proton signals for H-38, and H-44. Thus, for each of these positions an integral of 1H was assigned. [l] The $^{1}$H signal multiplicity for H-41 was interpreted as singlet, but hints of a not well resolved triplet ($J$ = 1.1) structure can be recognized upon closer inspection. [m] The $^{1}$H signal multiplicity for H-43 was interpreted as doublet of doublet (dd). Yet, a closer inspection hints towards a ddd structure, which is not completely resolved. [n] Even though the $^{1}$H signal of H48 appears on first glance to be a doublet, the HSQC data indicates-, that a part of this peak is obscured by the neighboring signal. Therefore, the $^{1}$H signal of H-48 was interpreted as multiplet.

**WNWTKHF**

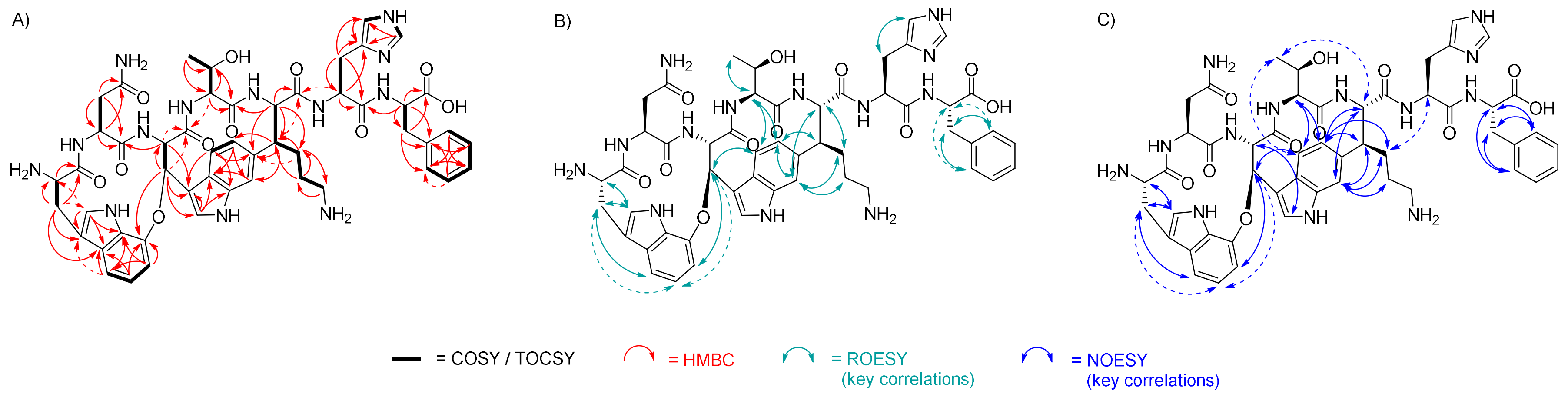

**Figure S5-3:** Key COSY/TOCSY (A, bold bonds), key HMBC (A, red arrows), key ROESY (B, turquoise arrows), and key NOESY (C, blue arrows) correlations of **WNWTKHF**. Dashed arrows indicate weak correlations.

Table S8: $^{1}H$ (700 MHz) and $^{13}C$ (175 MHz) NMR data of WNWTKTF ($D_2O$; δ in ppm). For $^{13}C$ measurements 3-(trimethylsilyl)propionic-2,2,3,3-$d_4$ acid sodium salt (TSPA) was used as external standard. The following abbreviations are used in this table: mult.: multiplicity, int.: integral, obs.: obscured, $C_q$: quaternary carbon atom. For atom numbering, cf. Figure S3.

| Position | $\delta_C$ [ppm], Type | $\delta_H$ [ppm], mult. (*J* in Hz), int.[a] | Position | $\delta_C$ [ppm], Type | $\delta_H$ [ppm], mult. (*J* in Hz), int.[a] |
|---|---|---|---|---|---|
| 1 | 59.2, CH | 4.04, pseudo-t (8.9), 1H | 25 | 129.3, $C_q$ | |
| 2 | 30.8, $CH_2$ | 3.56, dd (6.8, 12.6), 1H<br>3.32, m, 1H[b] | 26 | 172.6, $C_q$ | |
| 3 | 112.6, $C_q$ | | 27 | 62.6, CH | 3.74, d (6.1), 1H |
| 4 | 129.2, CH | 7.36, s (obs), 1H[c] | 28 | 72.5, CH | 3.41, p (5.9), 1H |
| 5 | 133.58 / 133.50,[d] $C_q$ | | 29 | 22.9, $CH_3$ | 0.80, d (6.1), 3H |
| 6 | 149.6, $C_q$ | | 30 | 172.5, $C_q$ | |
| 7 | 113.5, CH | 7.26, d (obs; 7.4),[e] 1H | 31 | 64.7, CH | 4.28, d (10.5), 1H |
| 8 | 124.7, CH | 7.20, t (6.9), 1H | 32 | 52.5, CH | 3.06, m,[h] 1H |
| 9 | 118.3, CH | 7.25, d (obs; 8.6),[e] 1H | 33 | 30.3, $CH_2$ | 2.05, m, 1H[i]<br>1.67, m, 1H[i] |
| 10 | 133.58 / 133.50,[d] $C_q$ | | 34 | 30.1, $CH_2$ | 1.90, m, 1H[j]<br>1.74, m, 1H[j] |
| 11 | 172.8, $C_q$ | | 35 | 43.9, $CH_2$ | 2.99, br t (6.1), 2H[k] |
| 12 | 55.3, CH | 3.34, m, 1H[b] | 36 | 176.5, $C_q$ | |
| 13 | 43.3, $CH_2$ | 2.17, m, 2H[f] | 37 | 63.7, CH | 4.33, d (3.4), 1H |
| 14 | 178.2, $C_q$ | | 38 | 71.3, CH | 4.22, m, 1H |
| 15 | 173.0, $C_q$ | | 39 | 23.3, $CH_3$ | 1.08, d (6.2), 3H |
| 16 | 67.8, CH | 4.71, d (8.8), 1H[g] | 40 | 175.0, $C_q$ | |
| 17 | 81.4, CH | 6.21, d (8.8), 1H | 41 | 60.8, CH | 4.50, pseudo-t (5.8), 1H |
| 18 | 116.1, $C_q$ | | 42 | 42.1, $CH_2$ | 3.19, dd (4.5, 13.6), 1H[l]<br>3.06, m,[h] 1H |
| 19 | 128.9, CH | 7.86, s, 1H | 43 | 142.0, $C_q$ | |
| 20 | 141.7, $C_q$ | | 44, 44’ | 134.1, CH | 7.31, d (7.1), 2H |
| 21 | 115.0, CH | 7.48, s, 1H | 45, 45’ | 133.1, CH | 7.42, t (7.1), 2H |
| 22 | 137.5, $C_q$ | | 46 | 131.4, CH | 7.37, m (obs),[m] 1H |
| 23 | 129.4, CH | 6.95, d (7.9), 1H | 47 | 181.9, $C_q$ | |
| 24 | 121.9, CH | 7.45, d (8.0), 1H | | | |

[a] The $^{1}H$ shifts of multiplets were extracted from the HSQC spectrum. [b] The observed integral for this signal was 2H due to the overlap of the proton signals for H-2, and H-12. Thus, for each of these positions an integral of 1H was assigned. [c] The observed integral for this signal was 2H due to partial overlap with the signal for H-46. The expected integral for H-4 is 1H. [d] Due to signal overlap of the 2D correlation signals, the $^{13}C$ shifts for C-5, and C-10 cannot be distinguished.

[e] According to the HSQC data, the peaks for H-7, and H-9 were interpreted as partially overlapping doublets. Due to the overlap, the observed integral for this signal was 2H, so that an integral of 1H each was assigned for H-7, and H-9. [f] The observed integral for this signal was 3H due to overlap with an impurity. The expected integral for H-13 is 2H. [g] The observed integral for this signal was 2H due to overlap with the $H_2O$ signal. The expected integral for H-16 is 1H. [h] Even though the peak appears on first glance to be a doublet of doublet (dd, $J$ = 7.9 Hz and 12.9 Hz), it was interpreted as multiplet, because it corresponds to the overlapping signals of H-32, and H-42. [i] The observed integral for the peak at 2.05 ppm was 4H, while for the peak at 1.67 ppm an integral of 2H was observed. For H-33 an integral of 1H is expected for each of the peaks. Deviation from the expected value is due to overlap with an impurity. [j] The observed integral for the peak at 1.90 ppm was 3H, while for the peak at 1.74 ppm an integral of 2H was observed. For H-34 an integral of 1H is expected for each of the peaks. Deviation from the expected value is due to overlap with an impurity. [k] The observed integral for this signal was 3H due to overlap with an impurity. The expected integral for H-35 is 2H. [l] The observed integral for this signal was 2H due to overlap with an impurity. For H-42 two signals with an integral of 1H each are expected. [m] Even though the $^{1}$H signal of H-46 appears on first glance to be a singlet, the HSQC data indicates, that a part of this peak is obscured by the neighboring signal. Therefore, the $^{1}$H signal of H-46 was interpreted as multiplet.

**WNWTKTF**

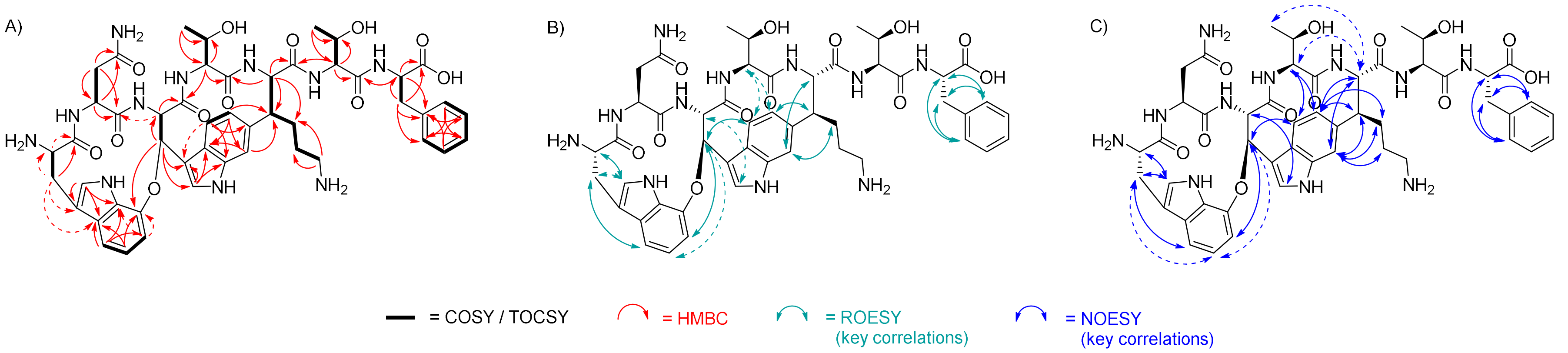

Figure S6: Key COSY/TOCSY (A, bold bonds), key HMBC (A, red arrows), key ROESY (B, turquoise arrows), and key NOESY (C, blue arrows) correlations of WNWTKTF. Dashed arrows indicate weak correlations.

Table S9: $^{1}H$ (700 MHz) and $^{13}C$ (175 MHz) NMR data of WNWSKMF ($D_2O$; $\delta$ in ppm). For $^{13}C$ measurements 3-(trimethylsilyl)propionic-2,2,3,3-d4 acid sodium salt (TSPA) was used as external standard. The following abbreviations are used in this table: mult.: multiplicity, int.: integral, obs.: obscured, $C_q$: quaternary carbon atom, n.o.: not observed, n.a.: not assigned. For atom numbering, cf. Figure S3.

| Position | $\delta_C$ [ppm], Type | $\delta_H$ [ppm], mult. ($J$ in Hz), int.[a] | Position | $\delta_C$ [ppm], Type | $\delta_H$ [ppm], mult. ($J$ in Hz), int.[a] |
|---|---|---|---|---|---|
| 1 | 59.1, CH | 4.04, dd (7.5, 10.8), 1H | 25 | 129.4, $C_q$ | |
| 2 | 30.8, $CH_2$ | 3.56, dd (7.3, 13.5), 1H<br>3.31, m, 1H[b] | 26 | 172.4, $C_q$ | |
| 3 | 112.6, $C_q$ | | 27 | 58.5, CH | 3.94, t (6.7), 1H |
| 4 | 129.2, CH | 7.35, s, 1H | 28 | 66.4, $CH_2$ | 3.20, m, 1H[e]<br>3.12, m, 1H |
| 5 | n.o.[c] | | 29 | 172.6, $C_q$ | |
| 6 | 149.6, $C_q$ | | 30 | 64.7, CH | 4.20, d (10.5), 1H |
| 7 | 113.4, CH | 7.25, d (obs; 8.7),[d] 1H | 31 | 52.4, CH | 3.06, m[f] |
| 8 | 124.7, CH | 7.19, t (7.6), 1H | 32 | 30.4, $CH_2$ | 2.06, m (obs)[g] |
| 9 | 118.2, CH | 7.23, d (obs; 8.8),[d] 1H | 33 | 30.1, $CH_2$ | 1.90, m (obs)[g]<br>1.74, m (obs)[g] |
| 10 | 133.5, $C_q$ | | 34 | 43.9, $CH_2$ | 3.00, t (6.5), 2H[h] |
| 11 | 172.7, $C_q$ | | 35 | 176.2, $C_q$ | |
| 12 | 55.3, CH | 3.32, m, 1H[b] | 36 | 57.5, CH | 4.45, dd (5.0, 9.6), 1H |
| 13 | 43.5, $CH_2$ | 2.19, dd (6.8, 14.0), 1H<br>2.14, dd (6.9, 14.2), 1H | 37 | n.a.[i] | n.a.[i] |
| 14 | 178.3, $C_q$ | | 38 | 33.9, $CH_2$ | 2.50, ddd (5.5, 7.8, 13.5), 1H<br>2.36, m, 1H[j] |
| 15 | 172.9, $C_q$ | | 39 | 18.6, $CH_3$ | 2.02, s, 3H |
| 16 | 67.8, CH | 4.68, d (8.9), 1H | 40 | 176.7, $C_q$ | |
| 17 | 81.3, CH | 6.19, d (8.9), 1H | 41 | 60.6, CH | 4.49, dd (5.5, 7.2), 1H |
| 18 | 116.1, $C_q$ | | 42 | 42.0, $CH_2$ | 3.22, m, 1H[e]<br>3.05, m[f] |
| 19 | 128.9, CH | 7.85, s, 1H | 43 | 142.0, $C_q$ | |
| 20 | 141.6, $C_q$ | | 44, 44’ | 134.1, CH | 7.29, d (7.6), 2H |
| 21 | 115.1, CH | 7.48, s, 1H | 45, 45’ | 133.1, CH | 7.42, t (7.2), 2H |
| 22 | 137.5, $C_q$ | | 46 | 131.5, CH | 7.36, m (obs),[k] 1H |
| 23 | 129.5, CH | 6.96, d (8.2), 1H | 47 | 181.7, $C_q$ | |
| 24 | 121.8, CH | 7.44, d (8.3), 1H | | | |

[a] The $^{1}$H shifts of multiplets were extracted from the HSQC spectrum. Due to the background signals, integration had to be performed with baseline correction to obtain rational integrals. [b] The observed integral for this signal was 2H due to the overlap of the proton signals for H-2, and H-12. Thus, for each of these positions an integral of 1H was assigned. [c] The quaternary carbon atom C-5 is not observed in the $^{13}$C spectrum. This might be due to the low sample amount. Likewise, signal overlap with C-10 is possible. [d] According to the HSQC data, the peaks for H-7, and H-9 were interpreted as partially overlapping doublets. Due to the overlap, the observed integral for this signal was 2H, so that an integral of 1H each was assigned for H-7, and H-9. [e] The observed integral for this signal was 2H due to the overlap of the proton signals for H-28, and H-42. Thus, for each of these positions an integral of 1H was assigned. [f] Even though the peak appears on first glance to be a doublet of doublet (dd, $J$ = 7.4 Hz and 13.7 Hz), it was interpreted as multiplet, because according to the HSQC spectrum it corresponds to the overlapping signals of H-31, and H-42. Still, the observed integral for this signal was 1H. Deviation from the expected value of 2H might be due to the chosen integration method. [g] Integrals for H-32, and H-33 cannot be provided as the respective proton signals overlap heavily with background signals. [h] The observed integral for this signal was 3H due to overlap with an impurity. The expected integral for H-34 is 2H. [i] Even though COSY correlations indicate the presence of H-37, it was not possible to unambiguously assign the $^{1}$H and $^{13}$C shifts for this position due to overlapping background signals. [j] The observed integral for this signal was 2H due to overlap with an impurity. For H-38 two signals with an integral of 1H each are expected. [k] Even though the $^{1}$H signal of H-46 appears on first glance to be a doublet, the HSQC data indicates, that a part of this peak is obscured by the neighboring signal. Therefore, the $^{1}$H signal of H-46 was interpreted as multiplet.

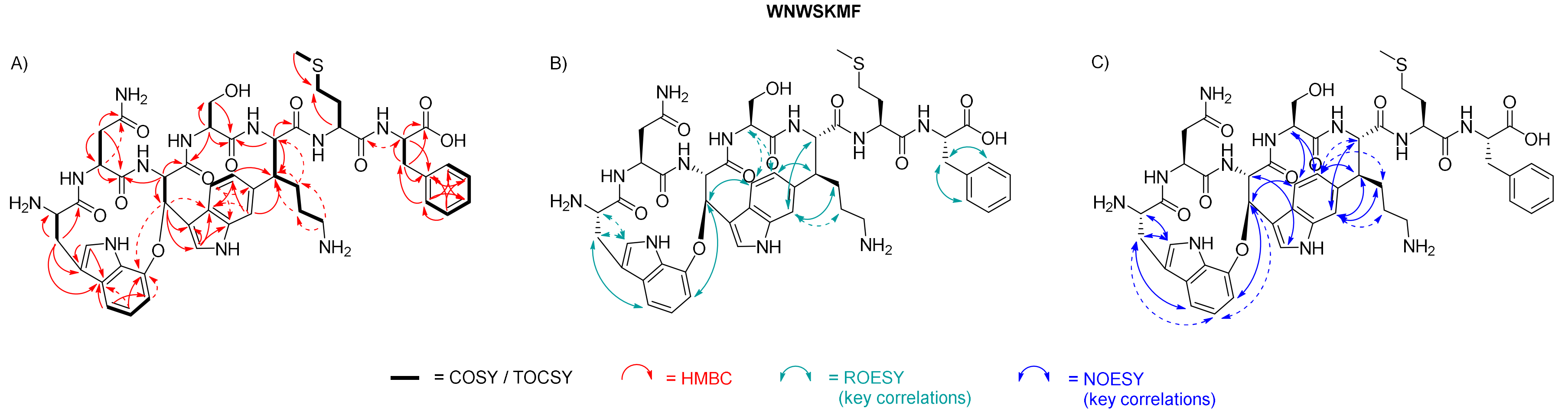

**Figure S7:** Key COSY/TOCSY (A, bold bonds), key HMBC (A, red arrows), key ROESY (B, turquoise arrows), and key NOESY (C, blue arrows) correlations of **WNWSKMF**. Dashed arrows indicate weak correlations.

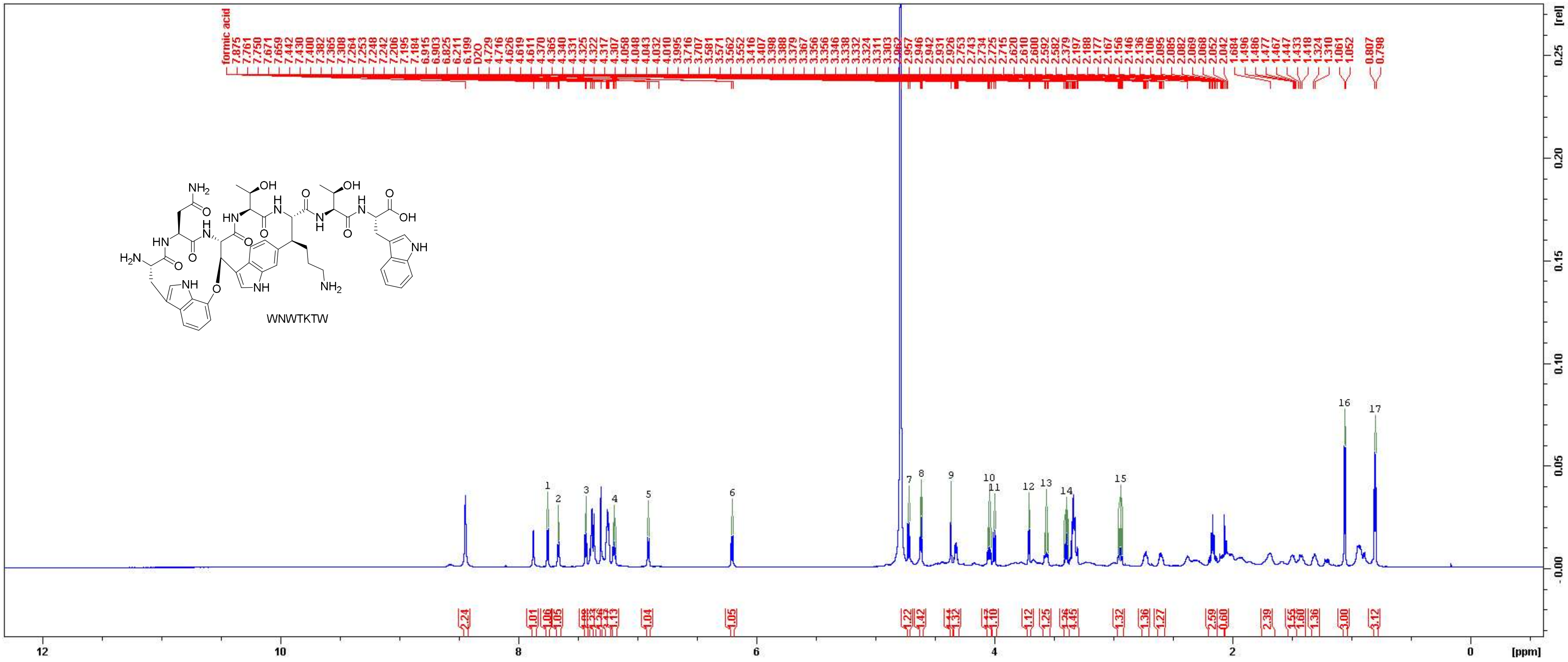

Figure S8: $^{1}H$ NMR spectrum of **WNWTKTW** in $D_2O$ (700 MHz).

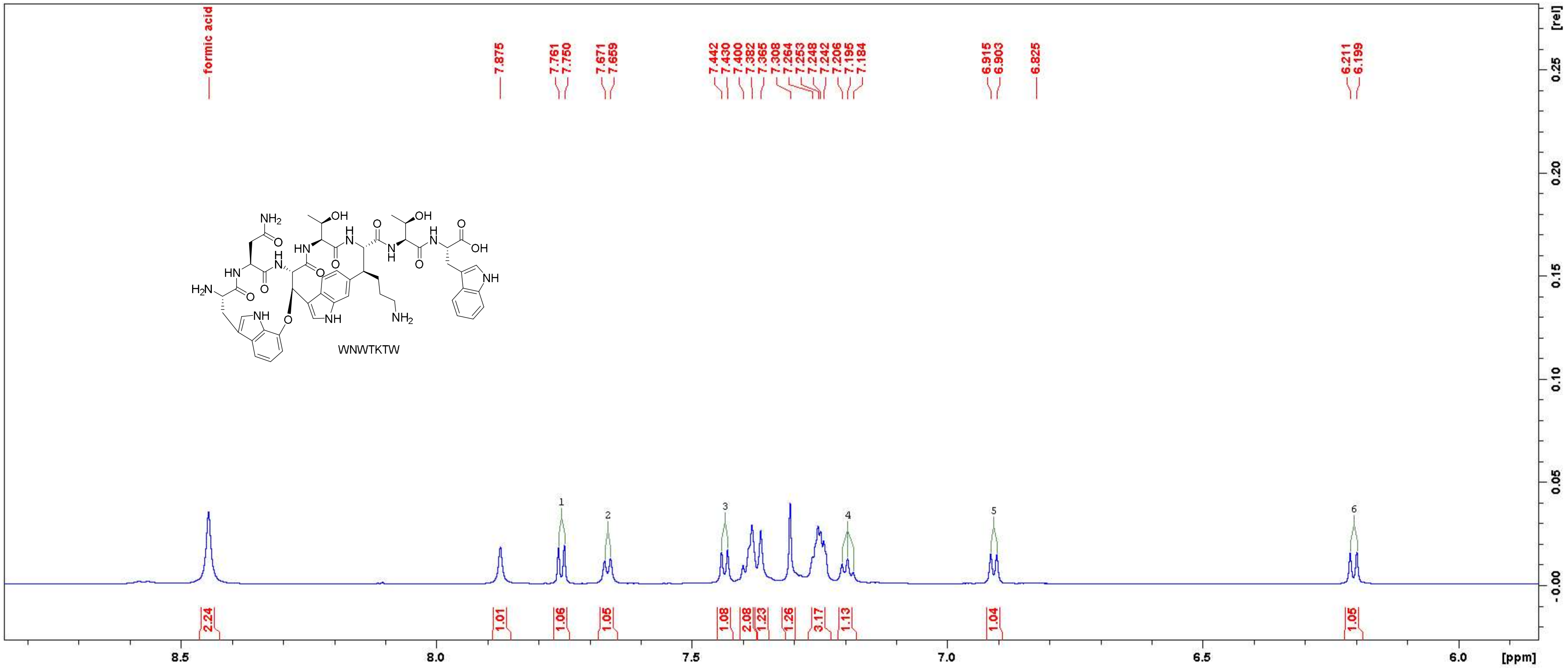

Figure S9: $^{1}$H NMR spectrum of WNWTKTW in $D_2O$ (700 MHz). Close-up in the region between 5.9 ppm and 8.8 ppm.

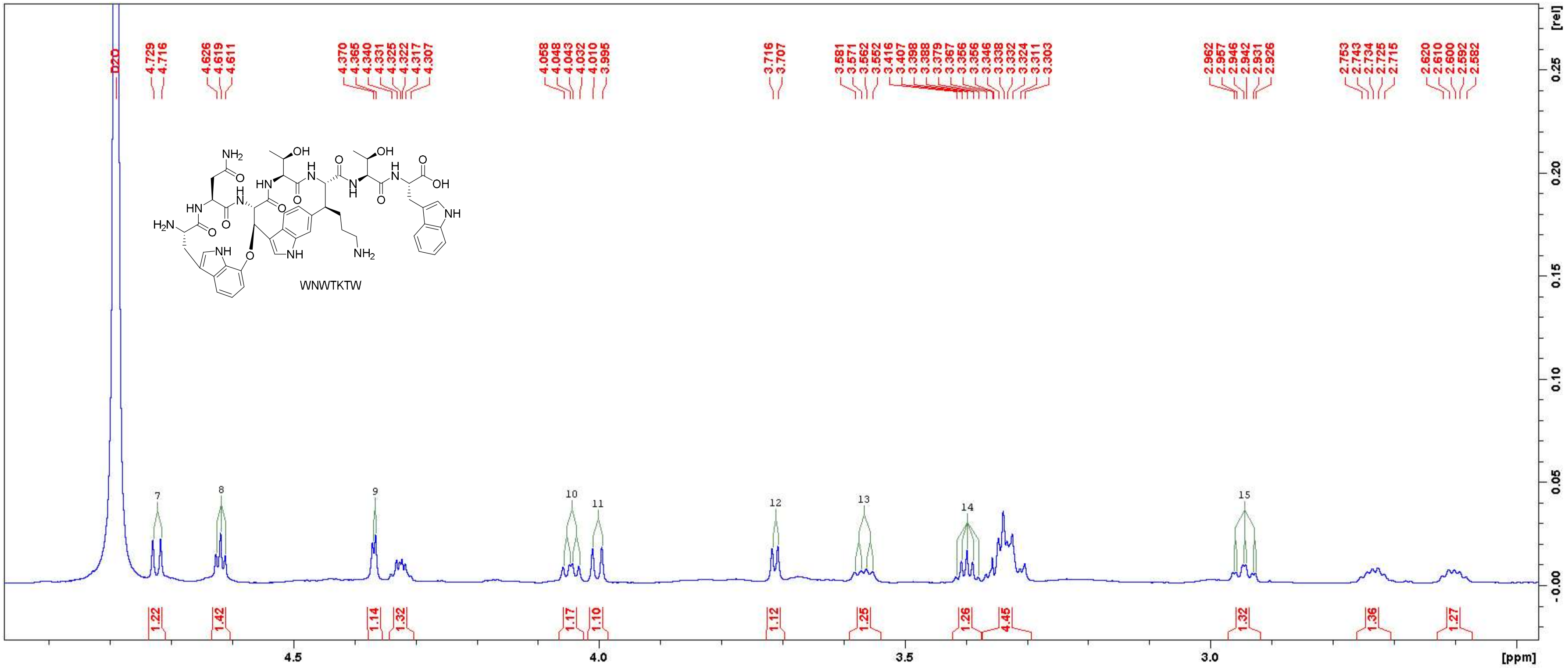

Figure S10: $^{1}H$ NMR spectrum of WNWTKTW in $D_2O$ (700 MHz). Close-up in the region between 2.5 ppm and 4.9 ppm.

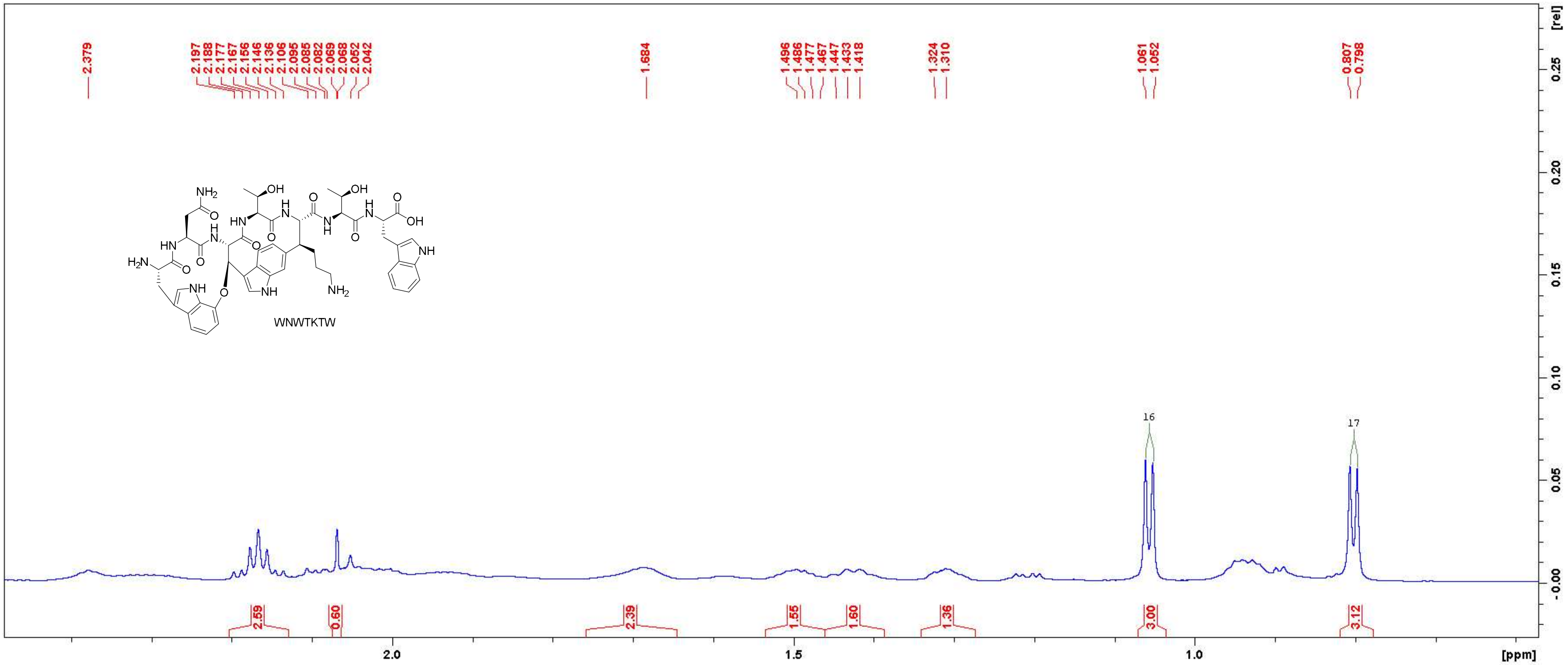

Figure S11: [1]H NMR spectrum of WNWTKTW in $D_2O$ (700 MHz). Close-up in the region between 0.6 ppm and 2.4 ppm.

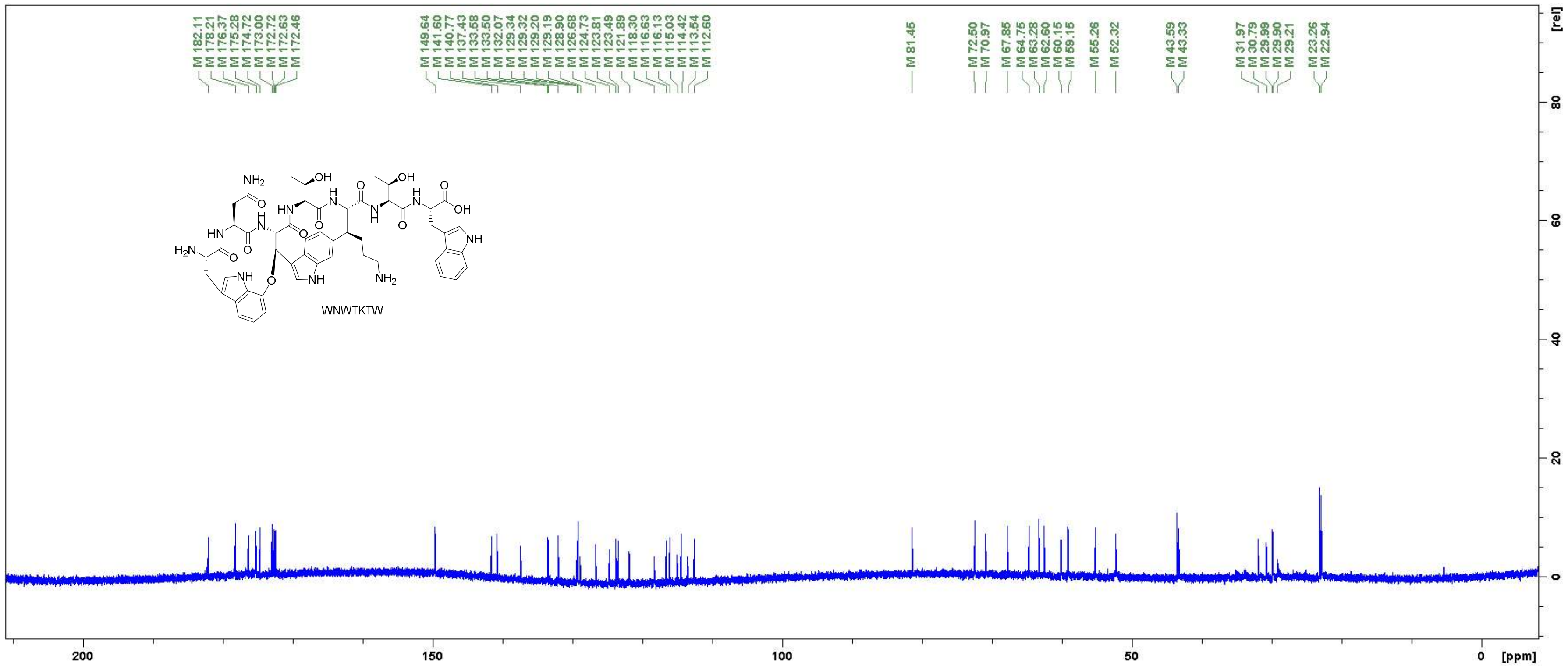

Figure S12: $^{13}C$ NMR spectrum of WNWTKTW in $D_2O$ (175 MHz).

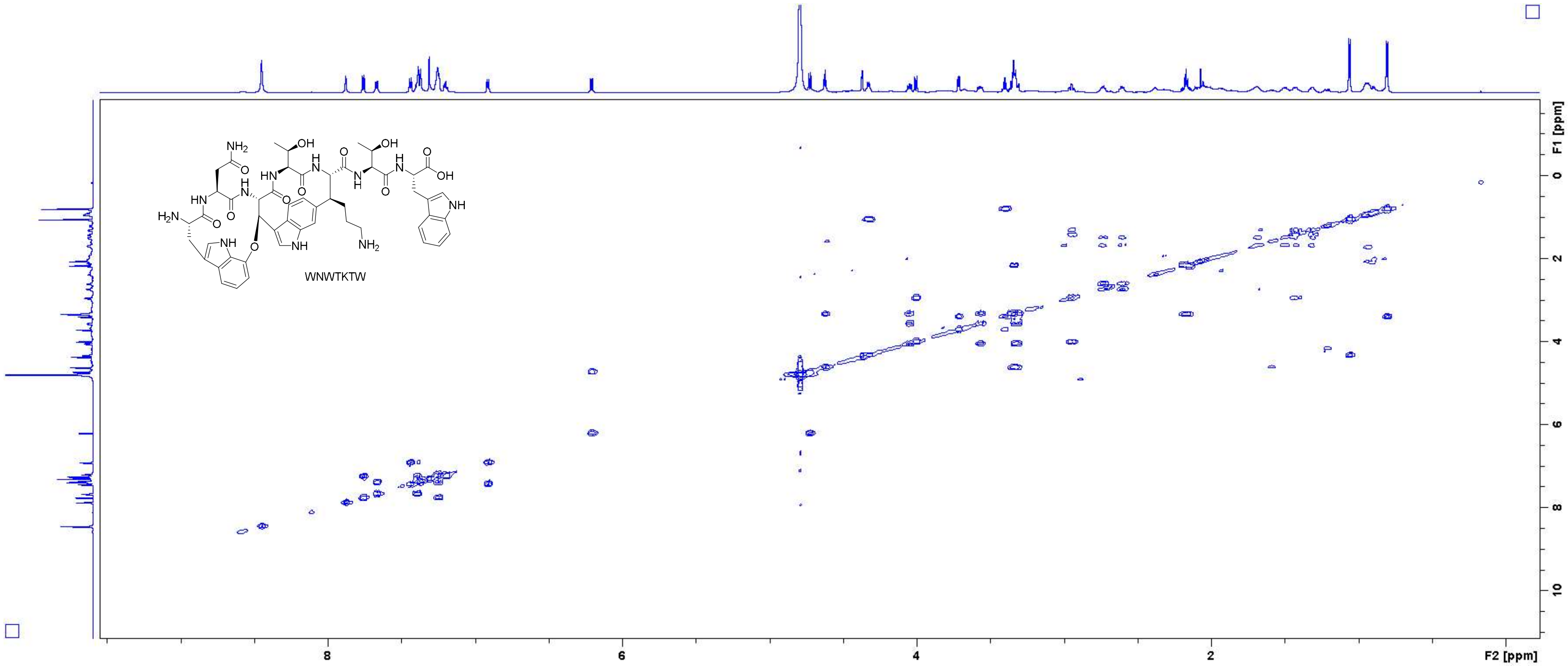

Figure S13: COSY spectrum of WNWTKTW in $D_2O$ (700 MHz).

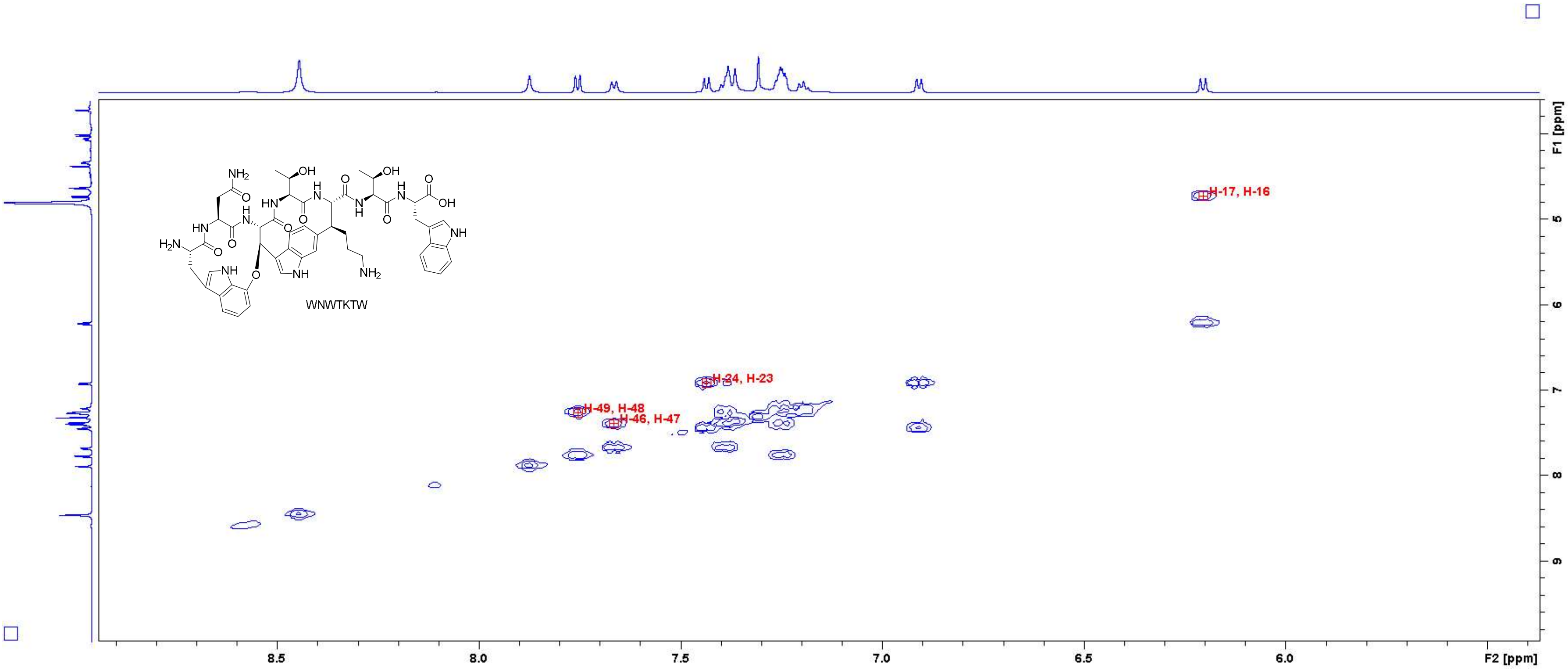

Figure S14: COSY spectrum of WNWTKTW in $D_2O$ (700 MHz). Close-up in the region of 5.4 – 8.9 ppm (F2 axis) and 3.8 – 9.8 ppm (F1 axis) with peak assignments. For atom numbering, cf. Figure S3.

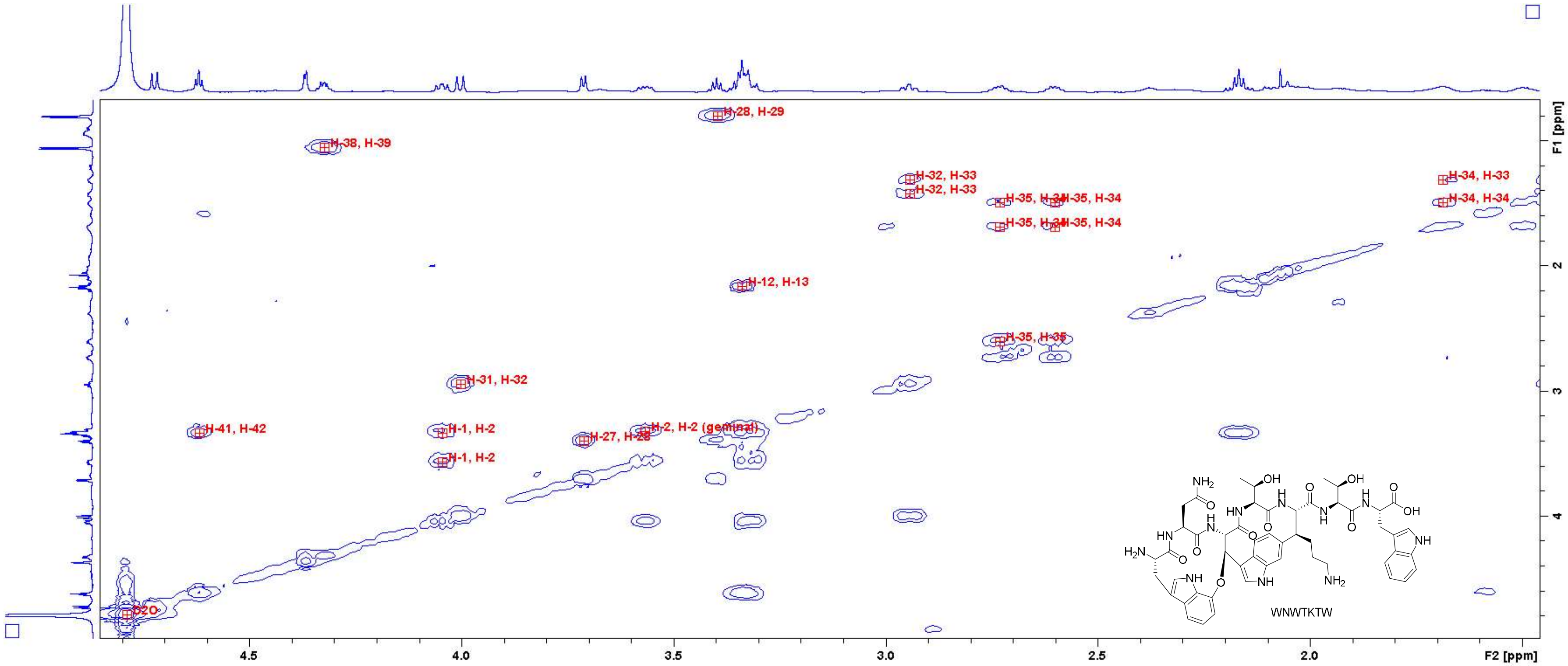

Figure S15: COSY spectrum of WNWTKTW in D2O (700 MHz). Close-up in the region of 1.5 – 4.8 ppm (F2 axis) and 0.8 – 4.8 ppm (F1 axis) with peak assignments. For atom numbering, cf. Figure S3.

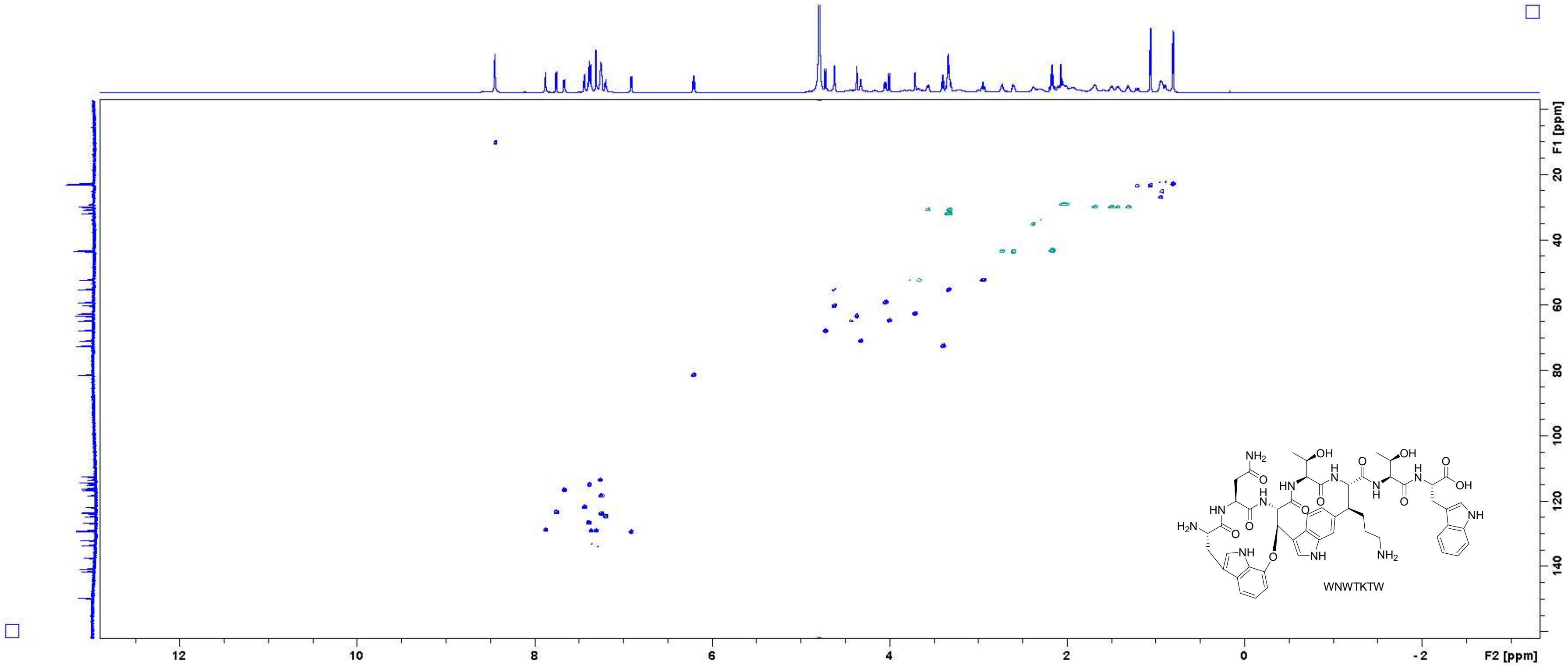

Figure S16: HSQC spectrum of WNWTKTW in $D_2O$ (700 MHz).

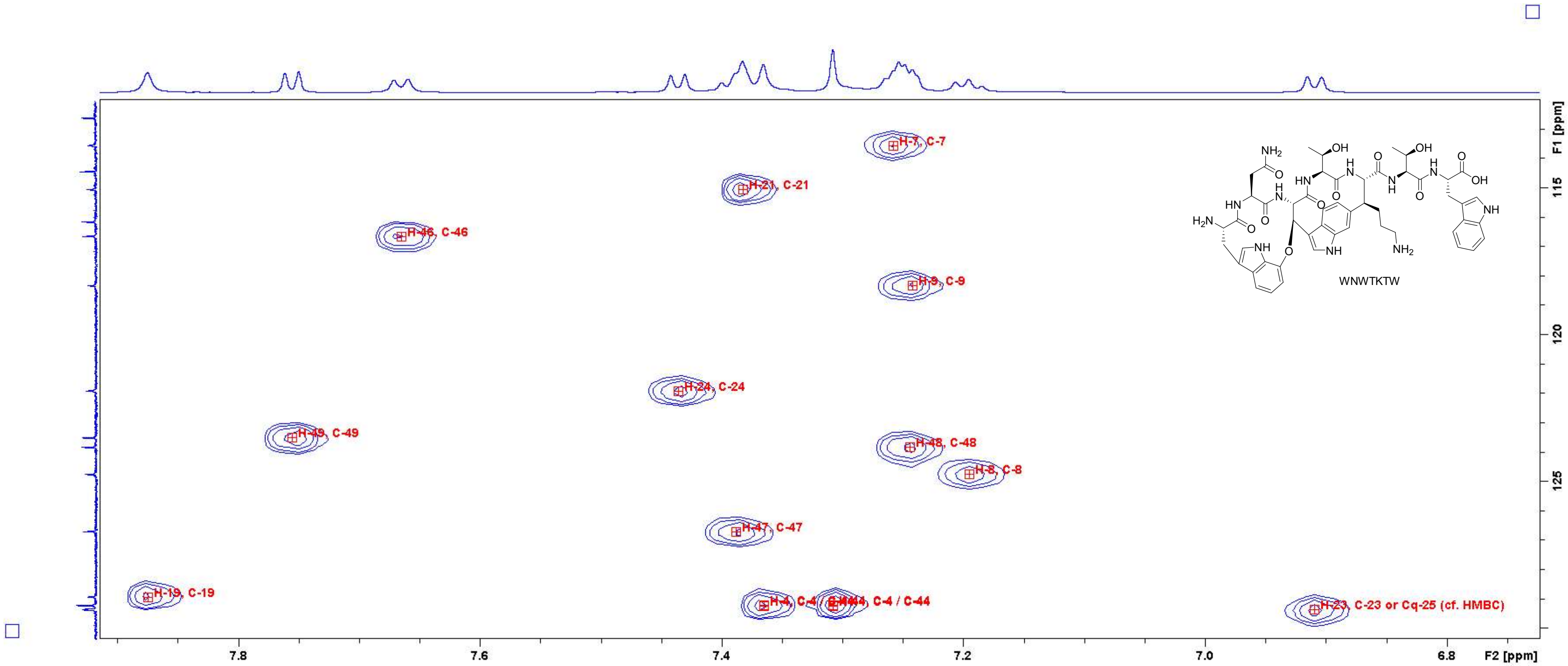

Figure S17: HSQC spectrum of **WNWTKTW** in $D_2O$ (700 MHz). Close-up in the region of 6.75 – 7.90 ppm (F2 axis) and 113 – 130 ppm (F1 axis) with peak assignments. For atom numbering, cf. Figure S3.

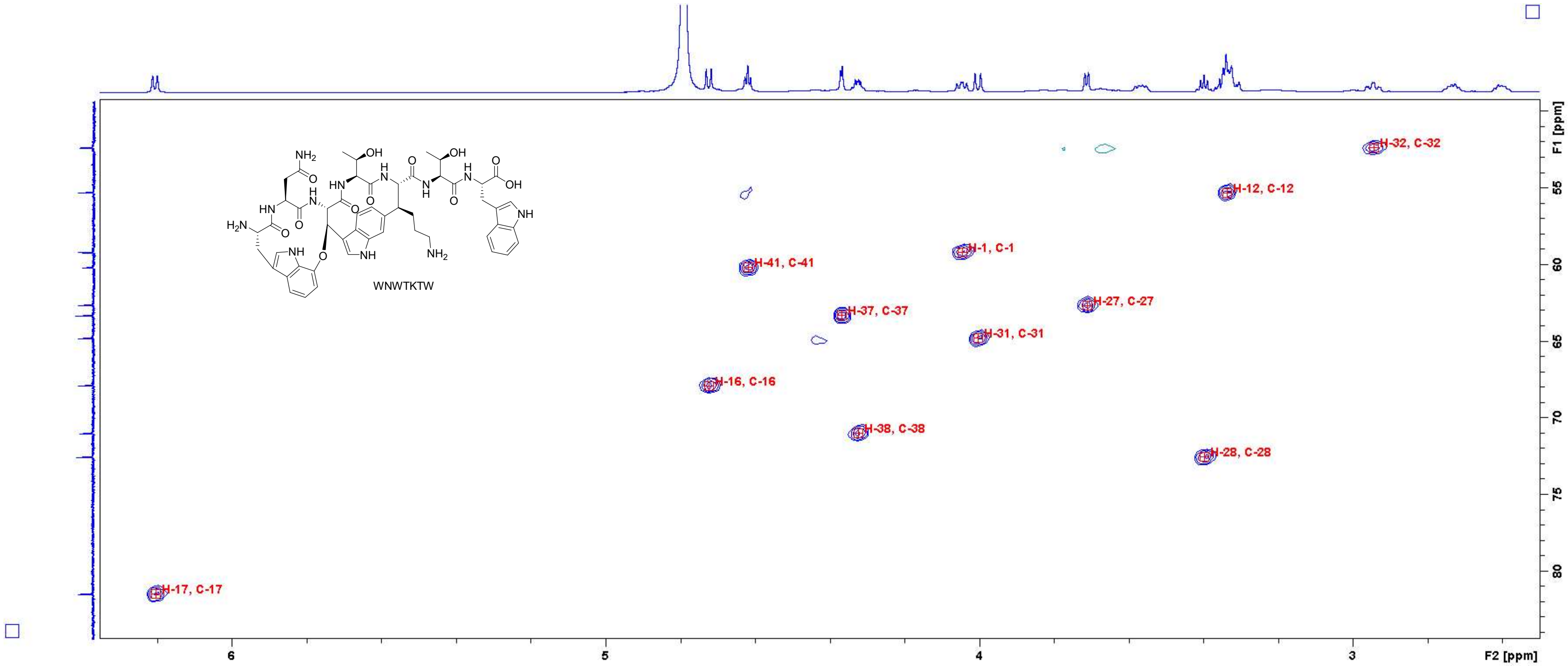

**Figure S18-16:** HSQC spectrum of **WNWTKTW** in $D_2O$ (700 MHz). Close-up in the region of 2.6 – 6.2 ppm (F2 axis) and 50 – 84 ppm (F1 axis) with peak assignments. For atom numbering, cf. Figure S3-1.

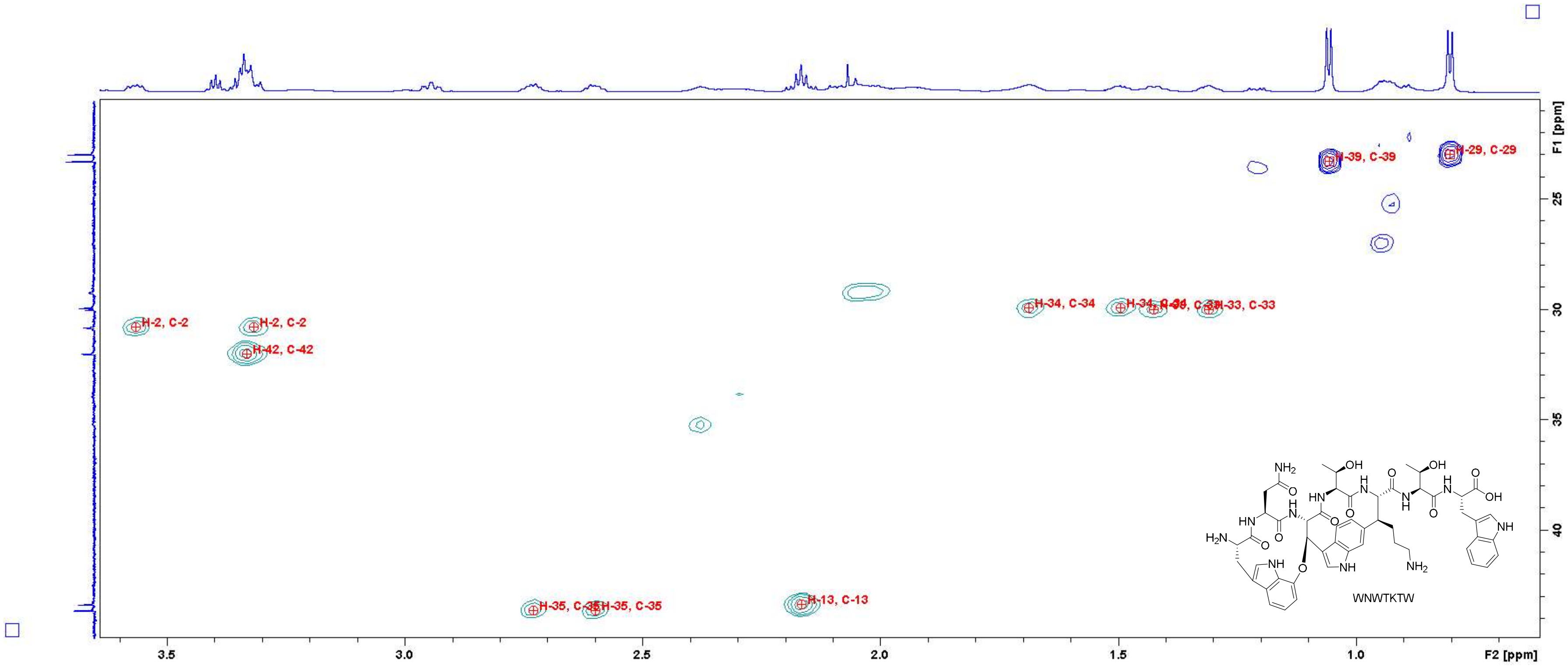

**Figure S19:** HSQC spectrum of **WNWTKTW** in $D_2O$ (700 MHz). Close-up in the region of 0.7 – 3.6 ppm (F2 axis) and 21 – 44 ppm (F1 axis) with peak assignments. For atom numbering, cf. Figure S3.

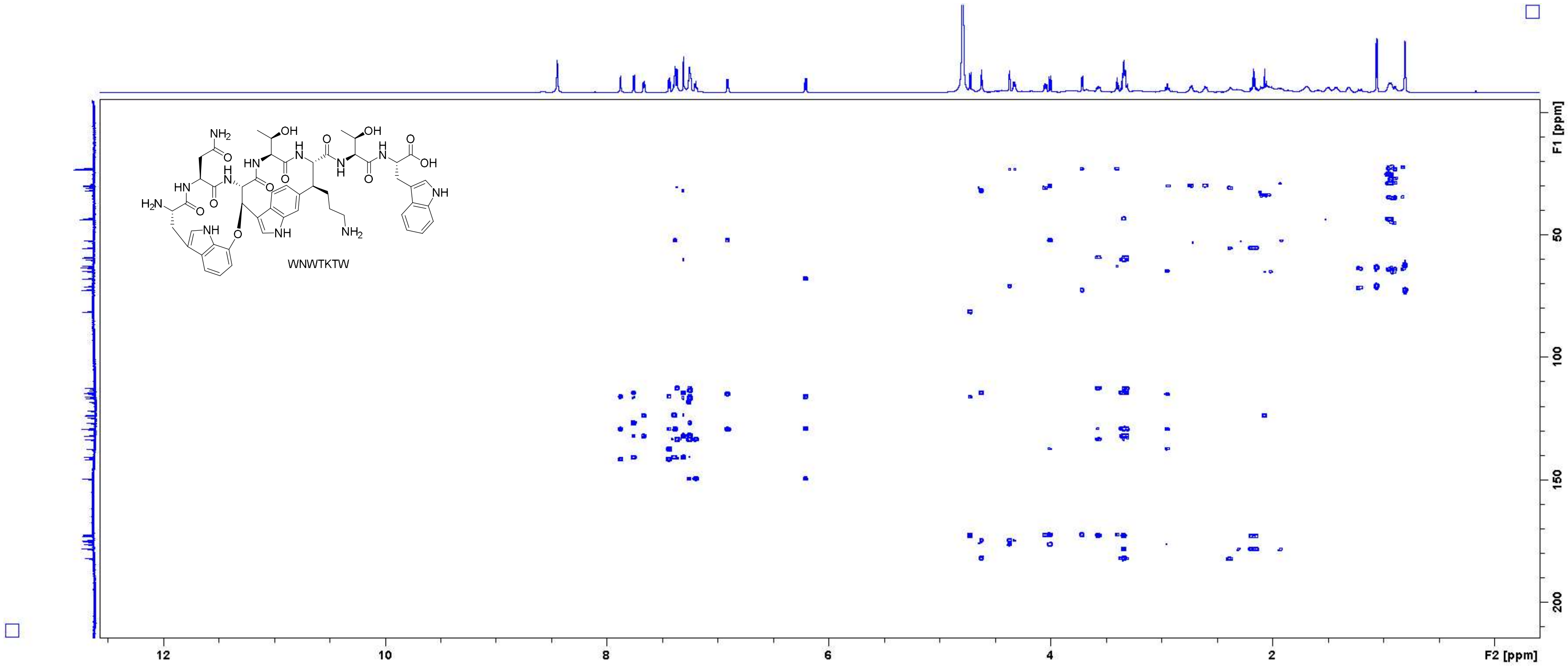

Figure S20: HMBC spectrum of WNWTKTW in $D_2O$ (700 MHz).

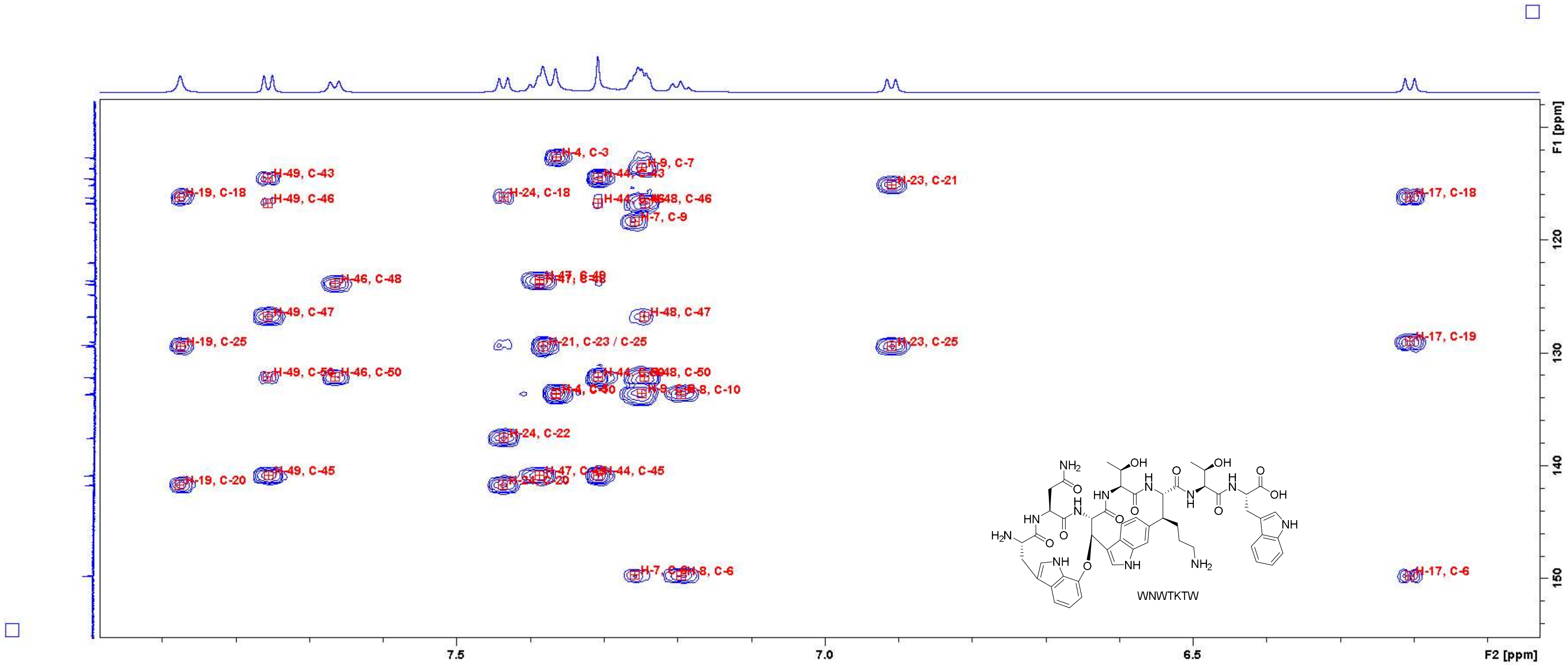

Figure S21: HMBC spectrum of **WNWTKTW** in $D_2O$ (700 MHz). Close-up in the region of 6.1 – 7.9 ppm (F2 axis) and 108 – 154 ppm (F1 axis) with peak assignments. For atom numbering, cf. Figure S3.

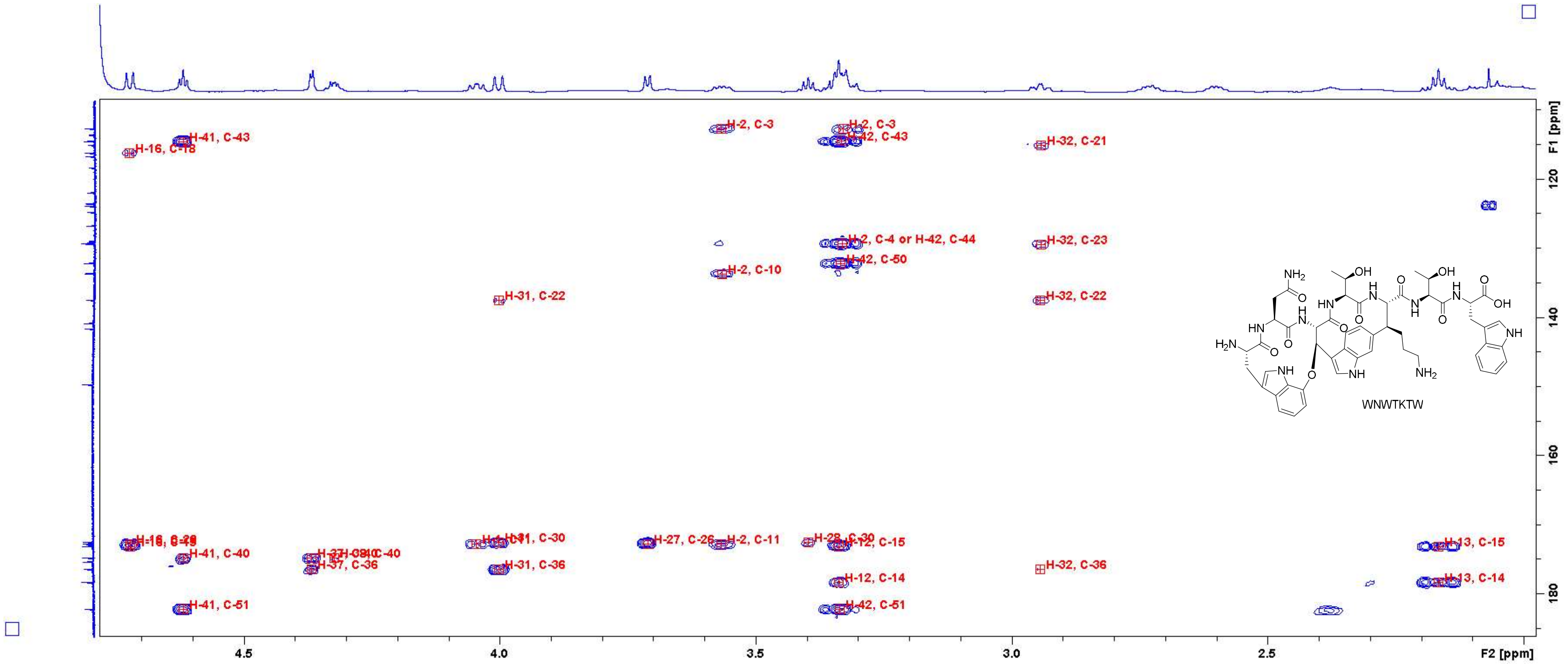

Figure S22: HMBC spectrum of **WNWTKTW** in $D_2O$ (700 MHz). Close-up in the region of 2.0 – 4.7 ppm (F2 axis) and 110 – 185 ppm (F1 axis) with peak assignments. For atom numbering, cf. Figure S3.

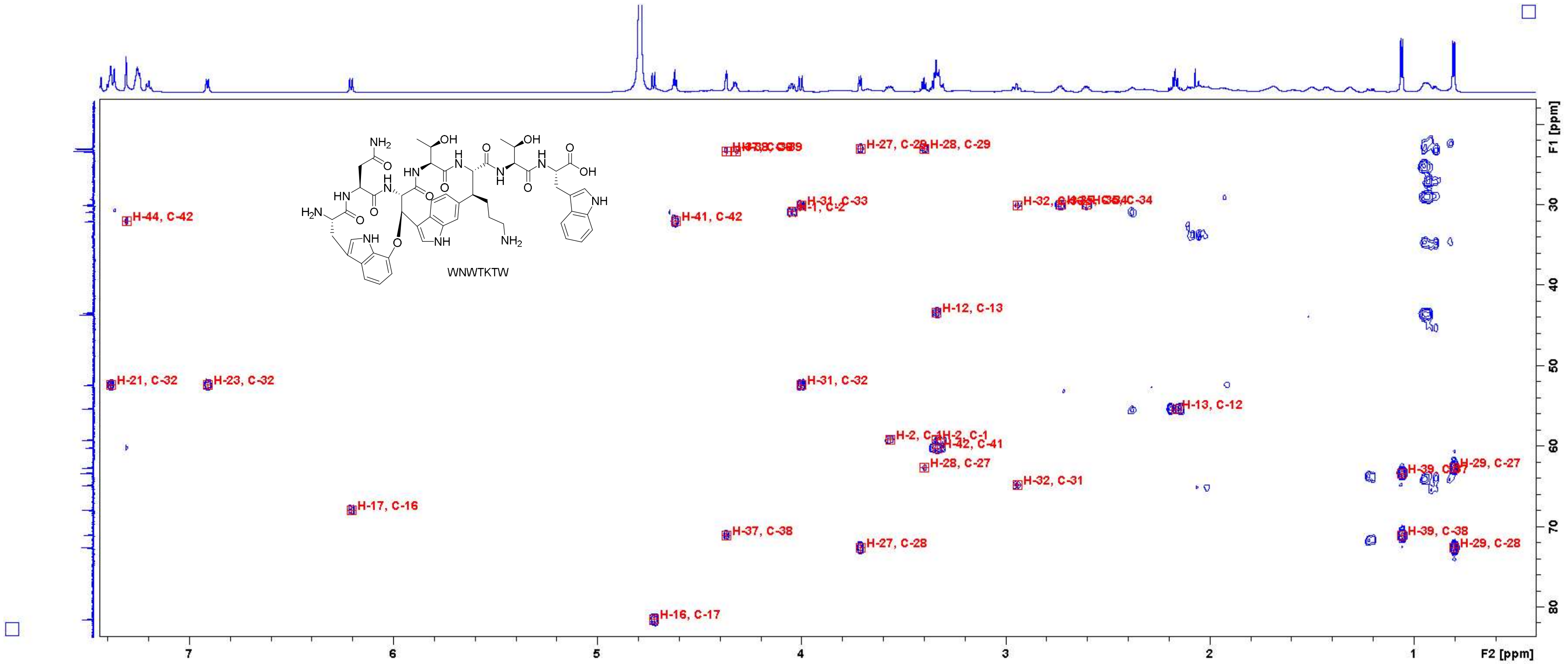

**Figure S23:** HMBC spectrum of **WNWTKTW** in $D_2O$ (700 MHz). Close-up in the region of 0.6 – 7.4 ppm (F2 axis) and 18 – 82 ppm (F1 axis) with peak assignments. For atom numbering, cf. Figure S3.

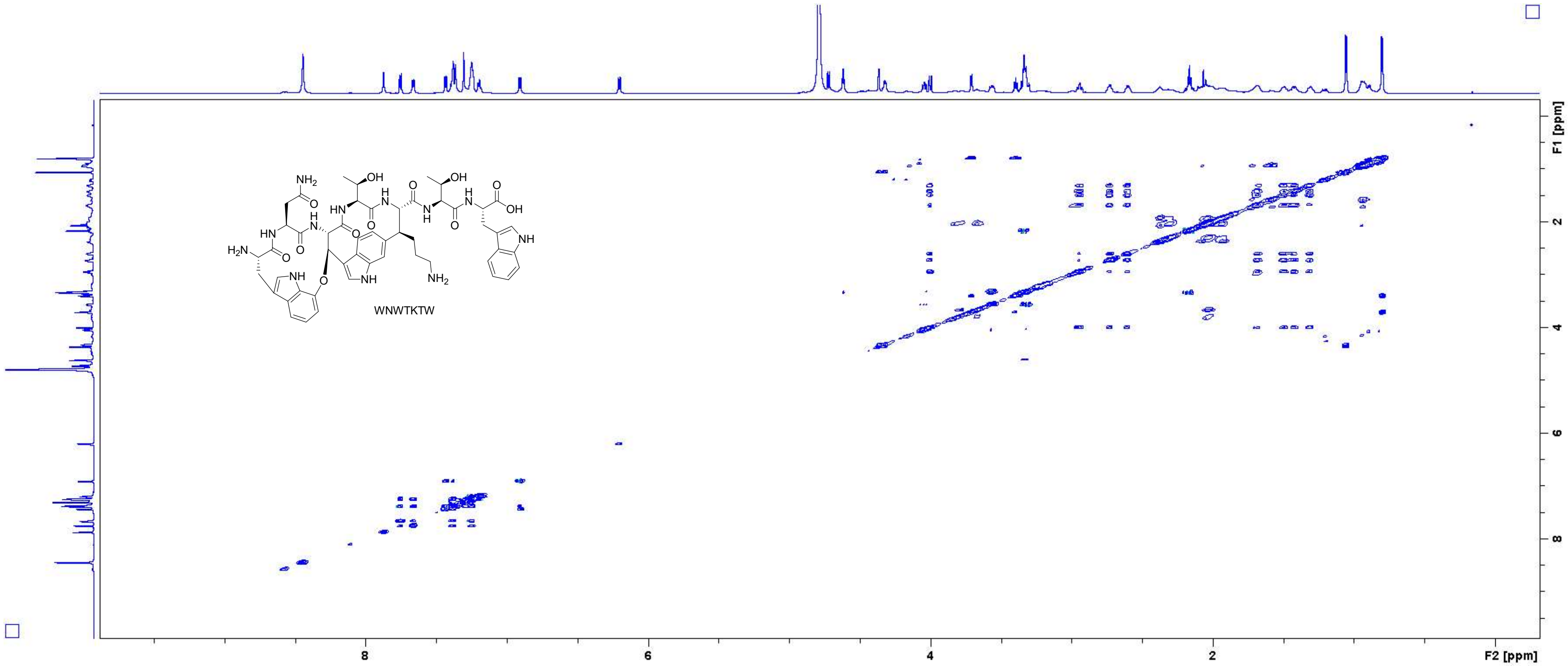

Figure S24: TOCSY spectrum of **WNWTKTW** in $D_2O$ (700 MHz, measured with $H_2O$ suppression).

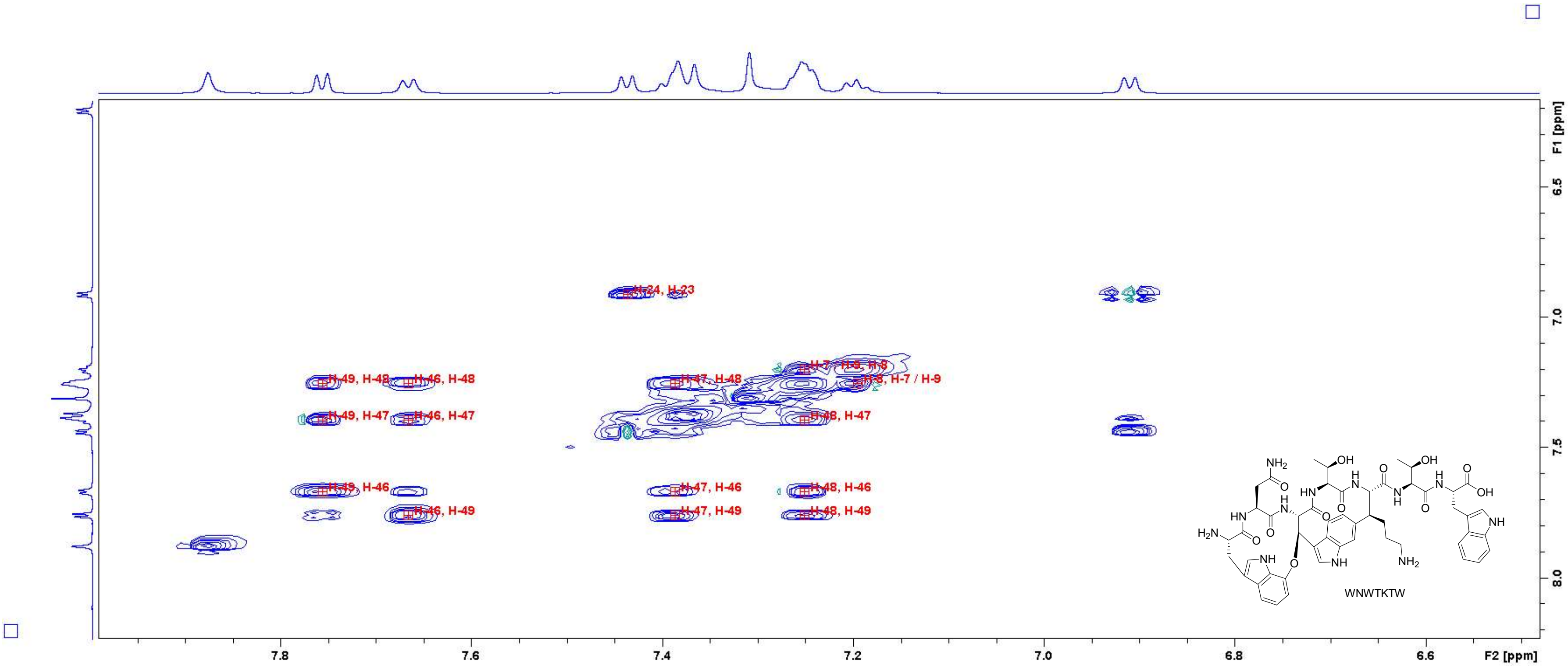

Figure S25: TOCSY spectrum of WNWTKTW in $D_2O$ (700 MHz, measured with $H_2O$ suppression). Close-up in the region of 6.50 – 7.95 ppm (F2 axis) and 6.2 – 8.2 ppm (F1 axis) with peak assignments. For atom numbering, cf. Figure S3.

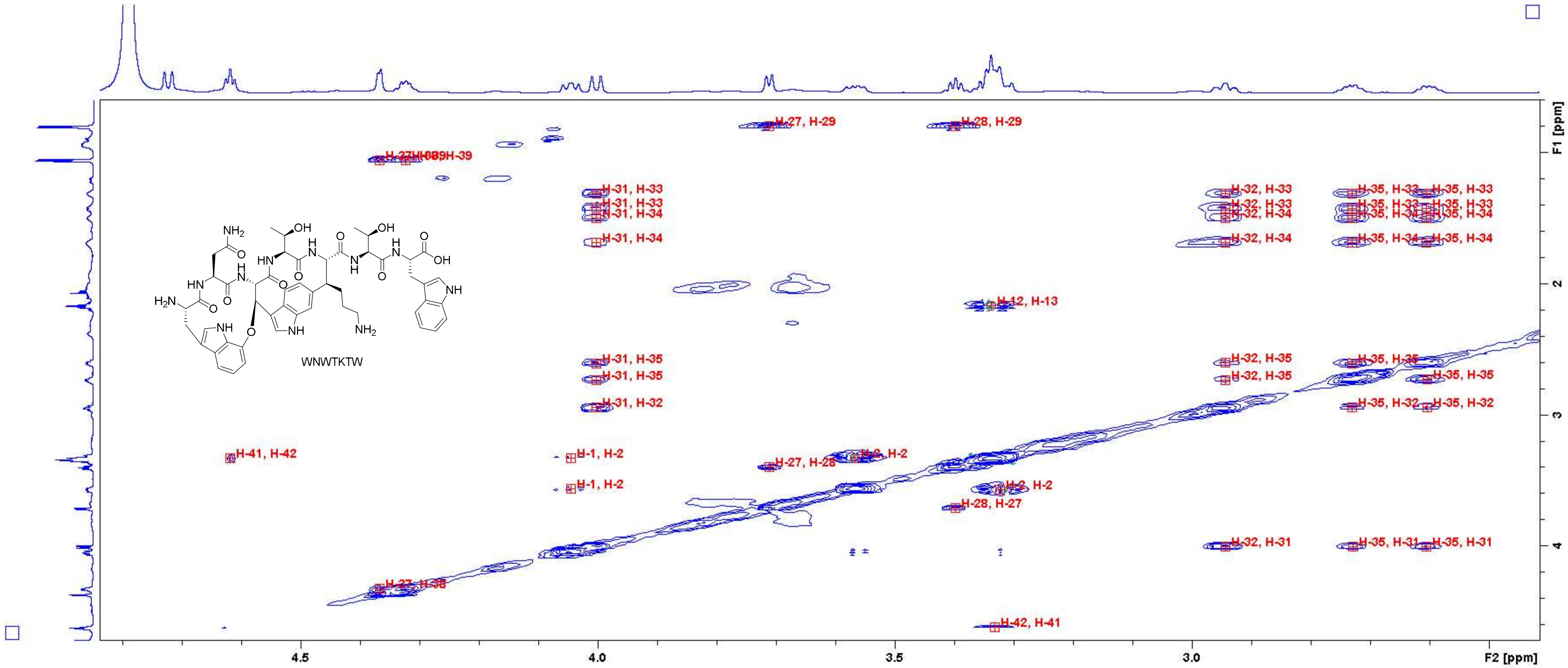

Figure S26: TOCSY spectrum of WNWTKTW in $D_2O$ (700 MHz, measured with $H_2O$ suppression). Close-up in the region of 2.5 – 4.8 ppm (F2 axis) and 0.6 – 4.6 ppm (F1 axis) with peak assignments. For atom numbering, cf. Figure S3.

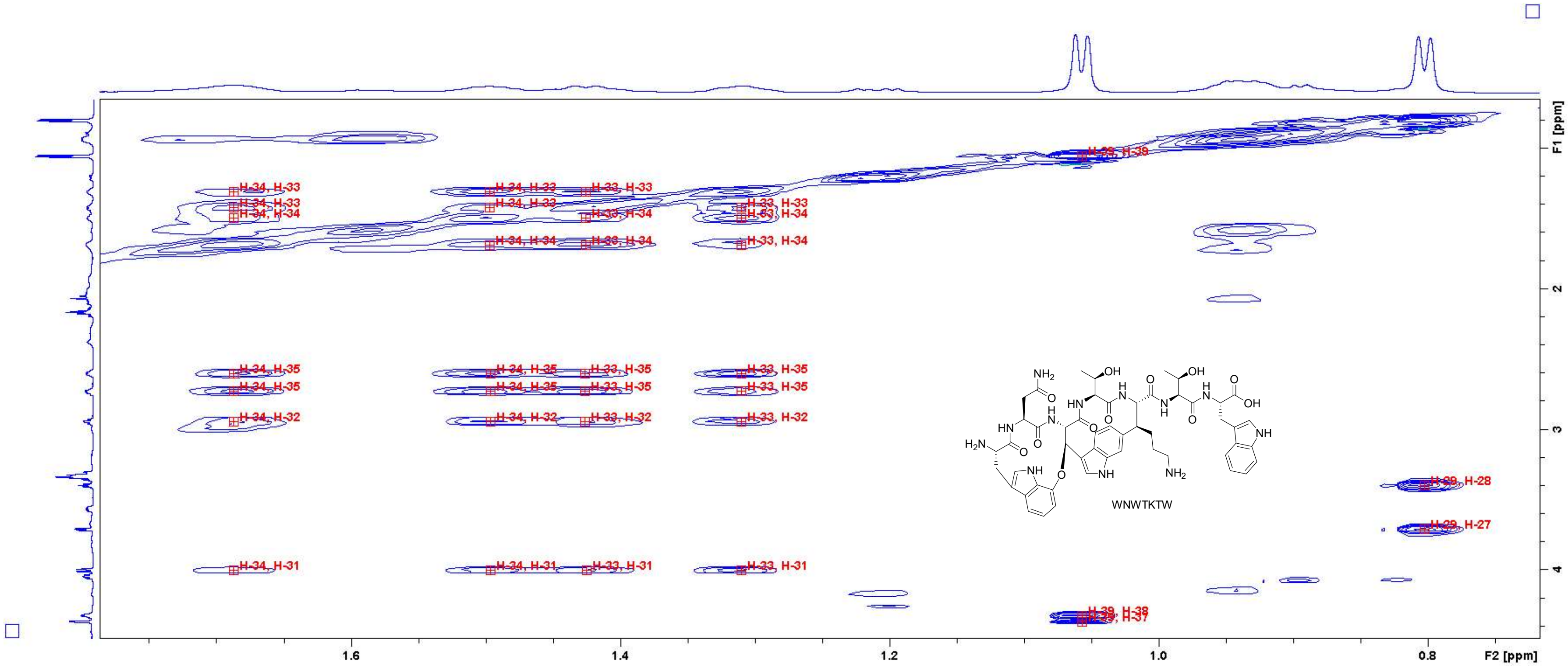

Figure S27: TOCSY spectrum of **WNWTKTW** in $D_2O$ (700 MHz, measured with $H_2O$ suppression). Close-up in the region of 0.75 – 1.75 ppm (F2 axis) and 0.8 – 4.4 ppm (F1 axis) with peak assignments. For atom numbering, cf. Figure S3.

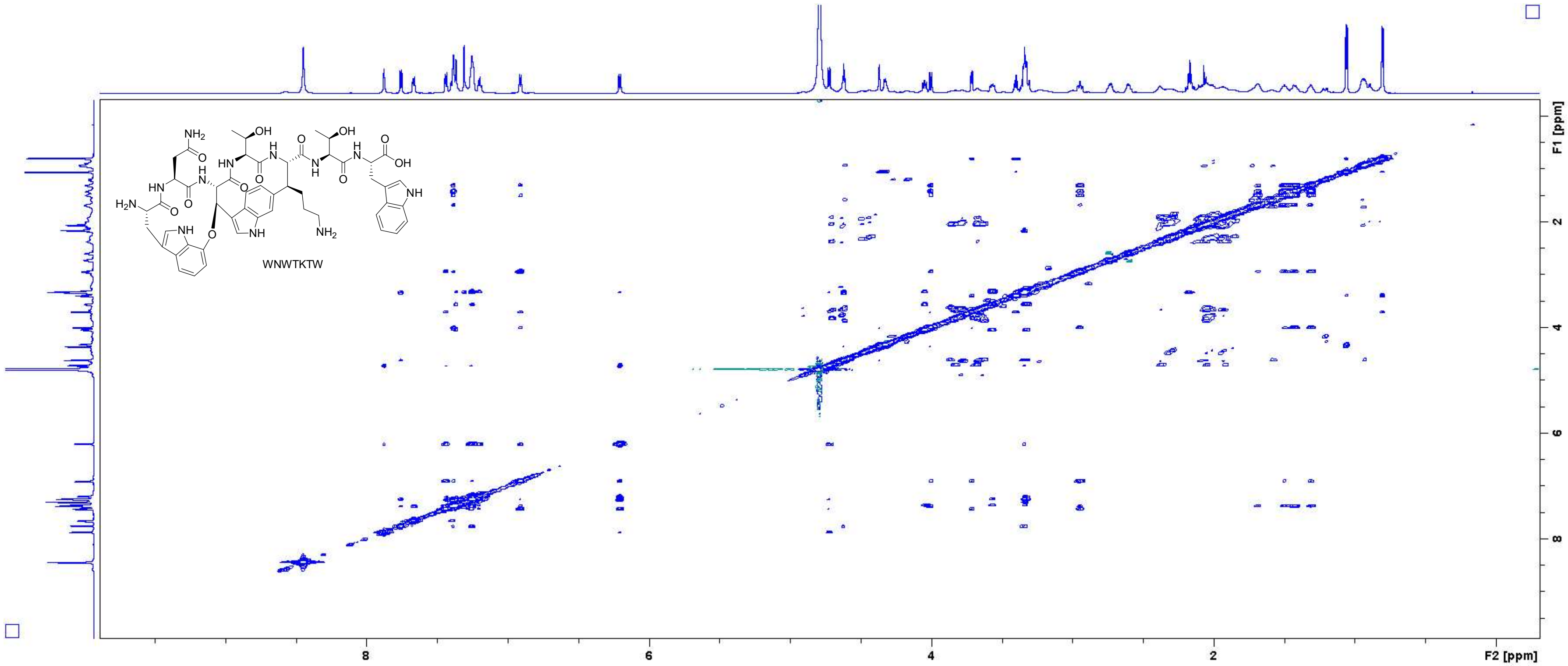

Figure S28: NOESY spectrum of WNWTKTW in $D_2O$ (700 MHz).

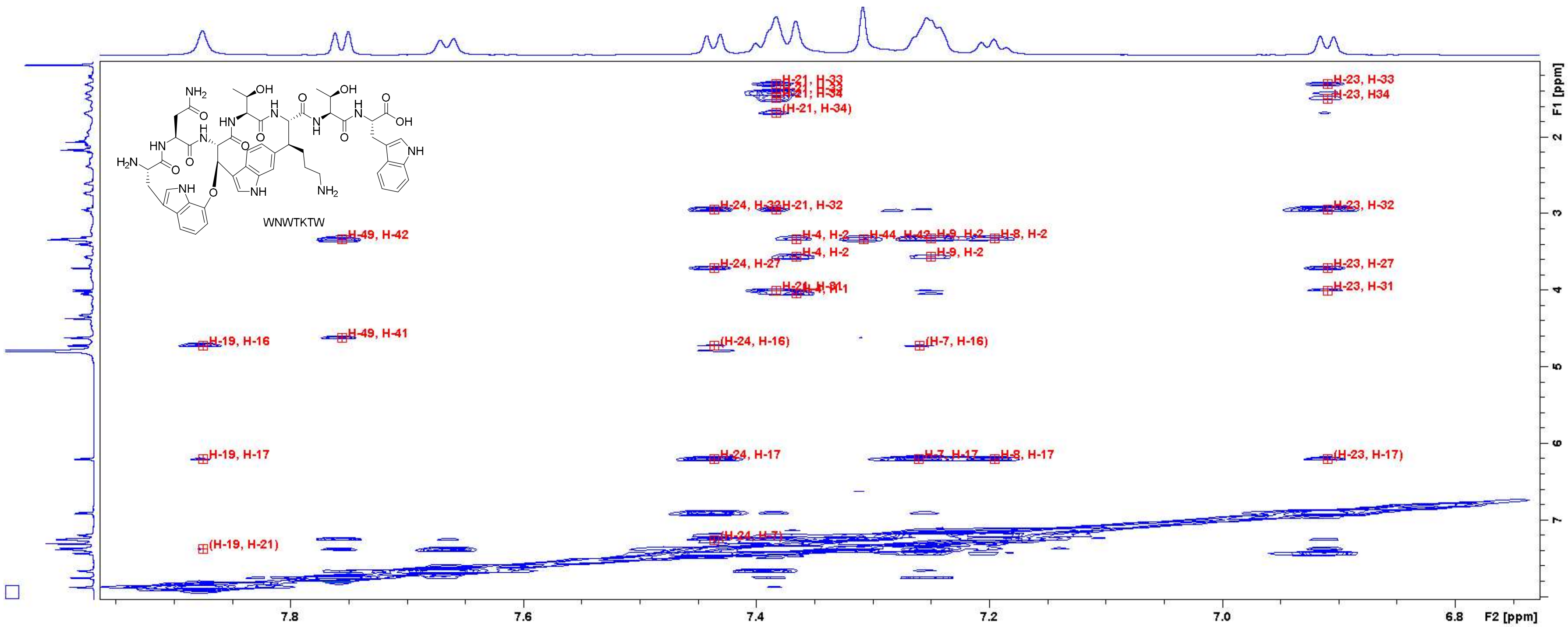

Figure S29: NOESY spectrum of **WNWTKTW** in $D_2O$ (700 MHz). Close-up in the region of 6.75 – 7.95 ppm (F2 axis) and 1.2 – 8.0 ppm (F1 axis) with peak assignments. For atom numbering, cf. Figure S3.

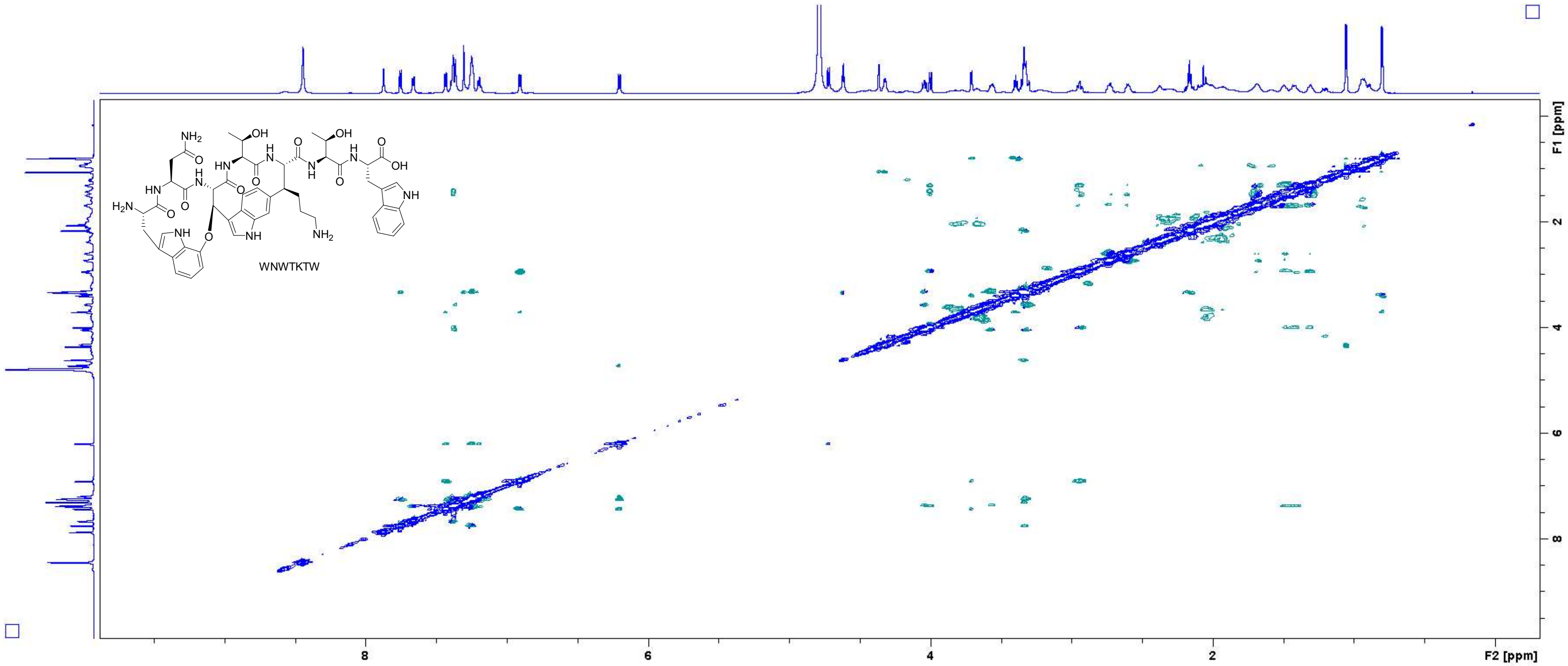

Figure S30: ROESY spectrum of WNWTKTW in $D_2O$ (700 MHz, measured with $H_2O$ suppression).

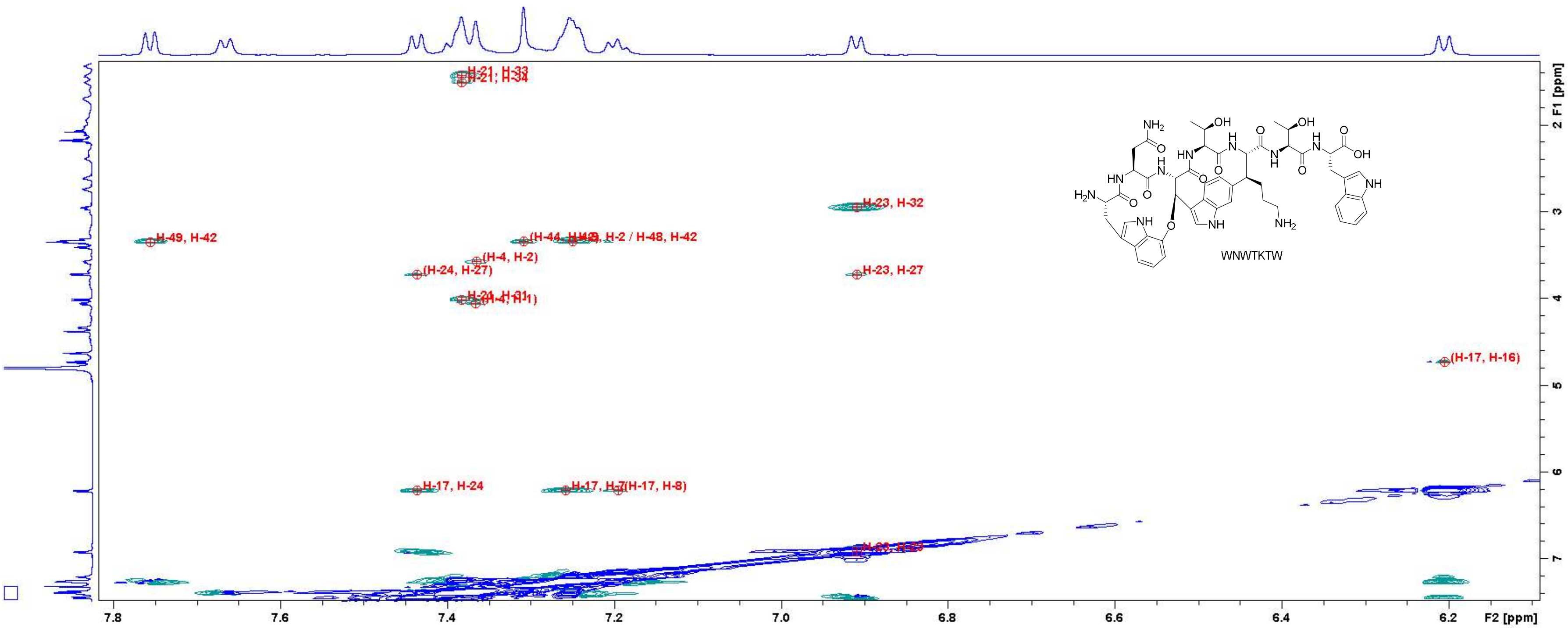

Figure S31: ROESY spectrum of WNWTKTW in $D_2O$ (700 MHz, measured with $H_2O$ suppression). Close-up in the region of 6.15 – 7.80 ppm (F2 axis) and 1.4 – 7.4 ppm (F1 axis) with peak assignments. For atom numbering, cf. Figure S3.

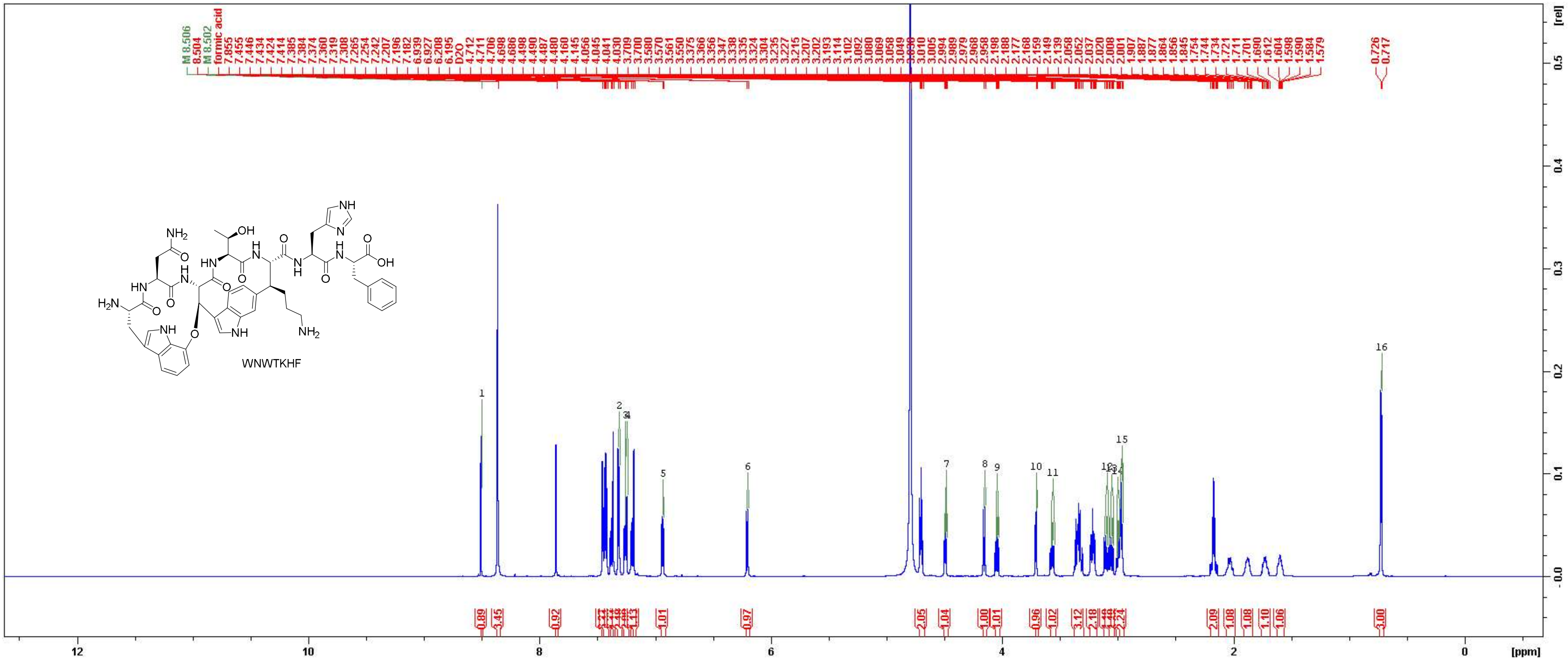

Figure S32: $^{1}$H NMR spectrum of WNWTKHF in $D_2O$ (700 MHz).

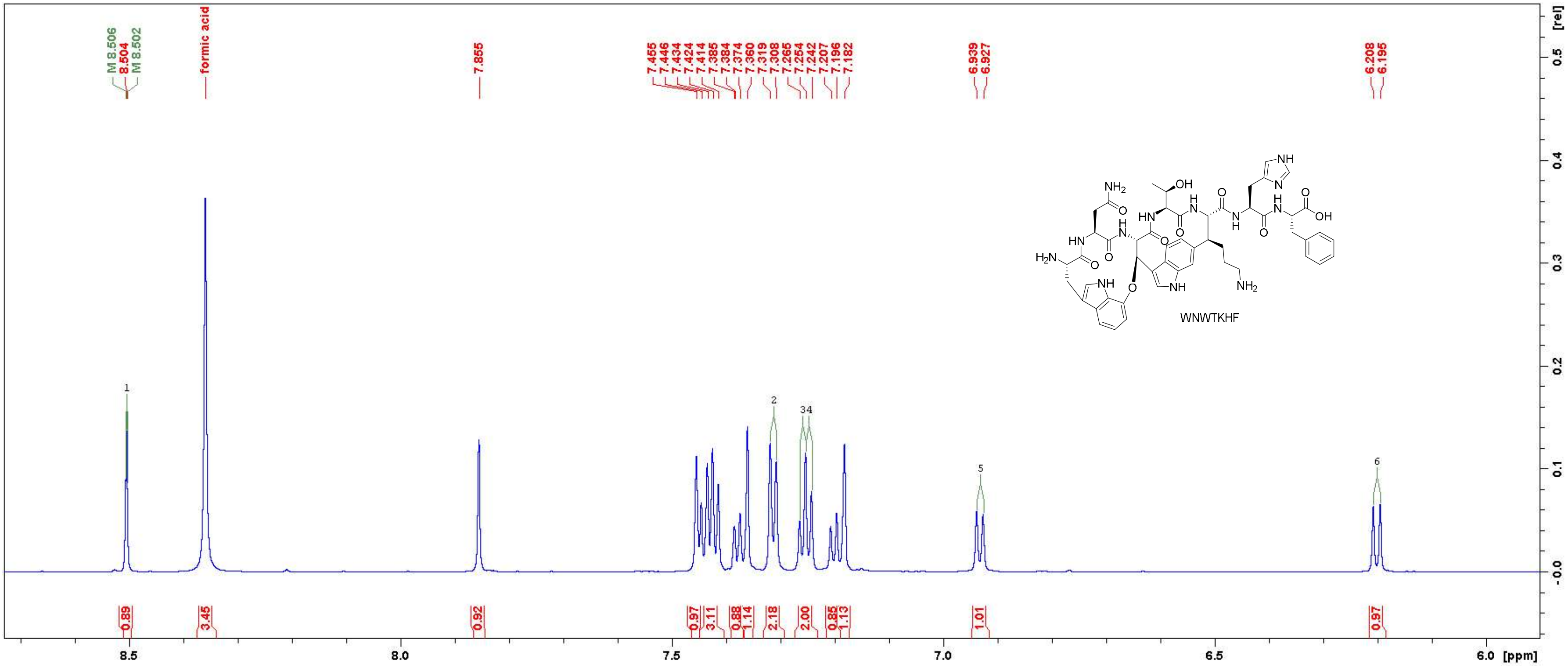

Figure S33: $^{1}H$ NMR spectrum of WNWTKHF in $D_2O$ (700 MHz). Close-up in the region between 6.0 ppm and 8.7 ppm.

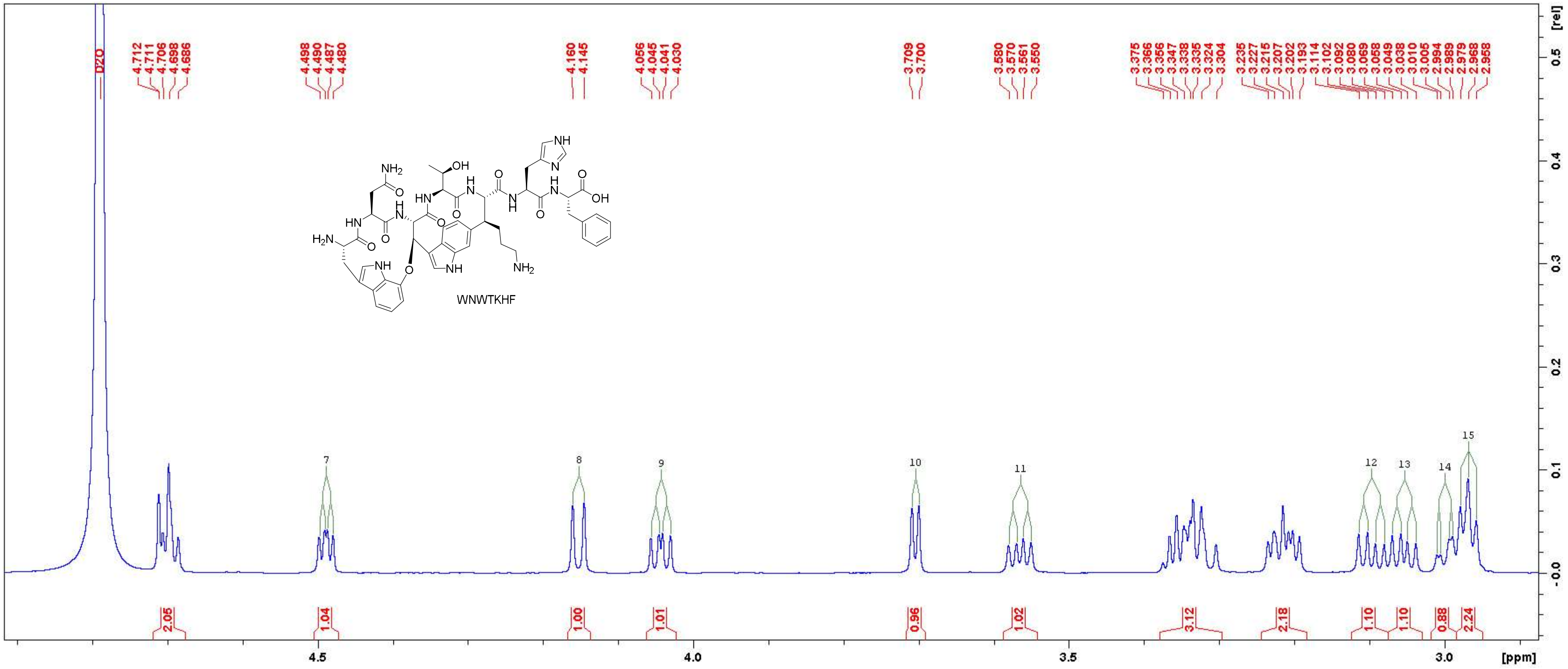

Figure S34: $^1$H NMR spectrum of WNWTKHF in $D_2O$ (700 MHz). Close-up in the region between 2.9 ppm and 4.9 ppm.

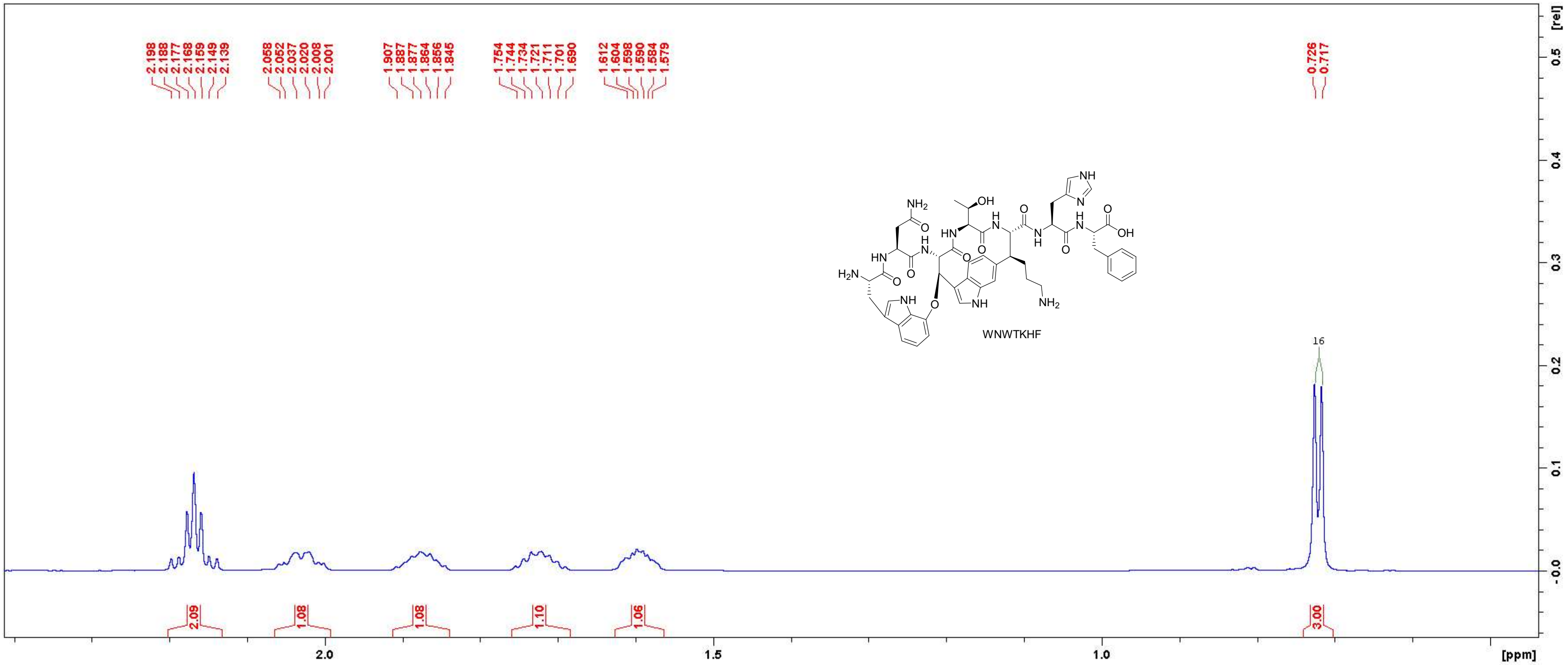

Figure S35: $^{1}H$ NMR spectrum of WNWTKHF in $D_2O$ (700 MHz). Close-up in the region between 0.5 ppm and 2.4 ppm.

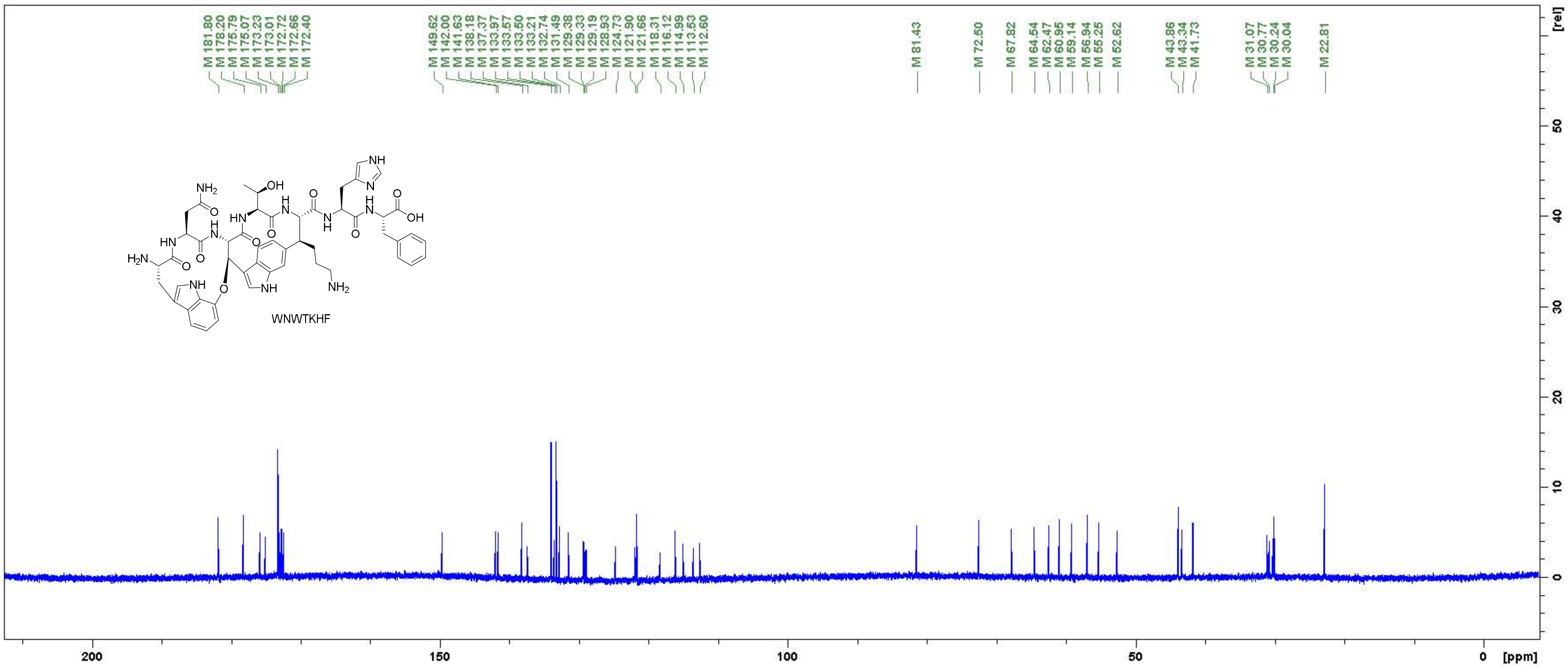

Figure S36: $^{13}$C NMR spectrum of WNWTKHF in $D_2O$ (175 MHz).

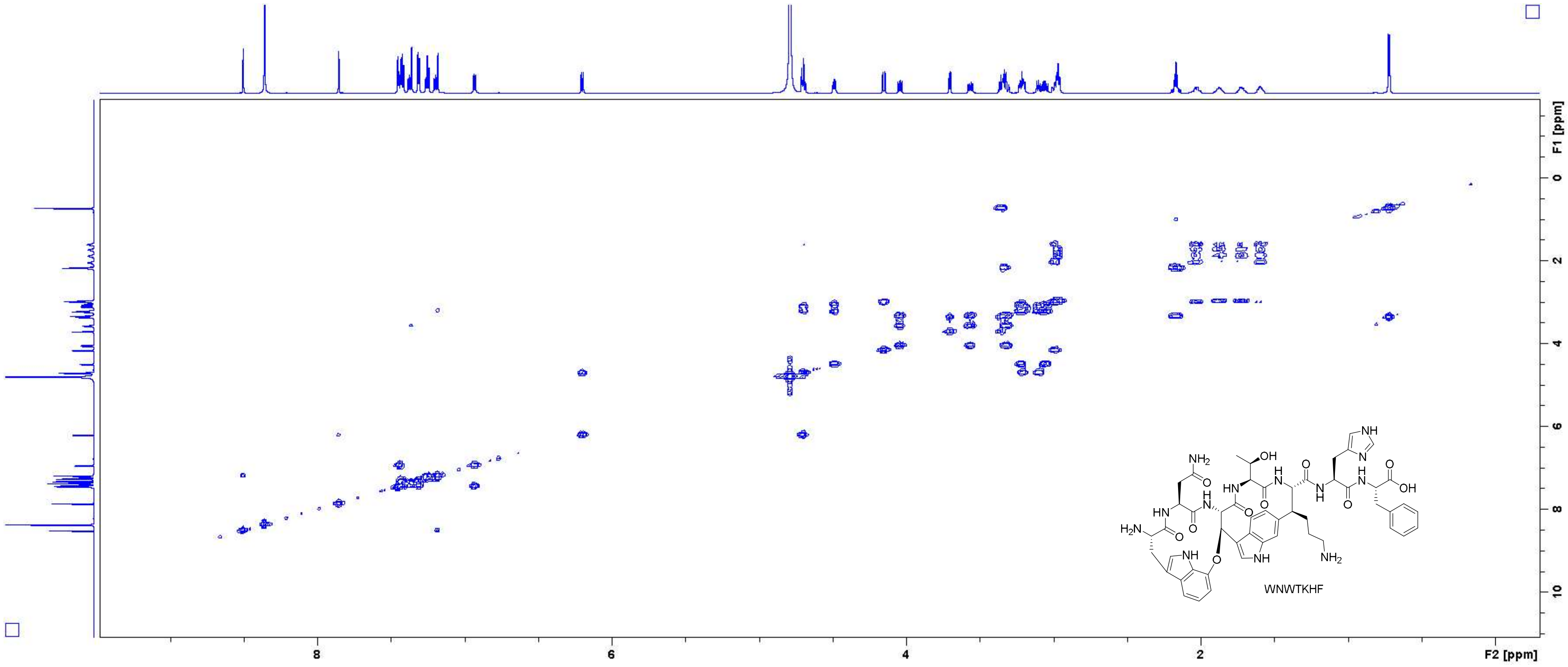

Figure S37: COSY spectrum of WNWTKHF in $D_2O$ (700 MHz).

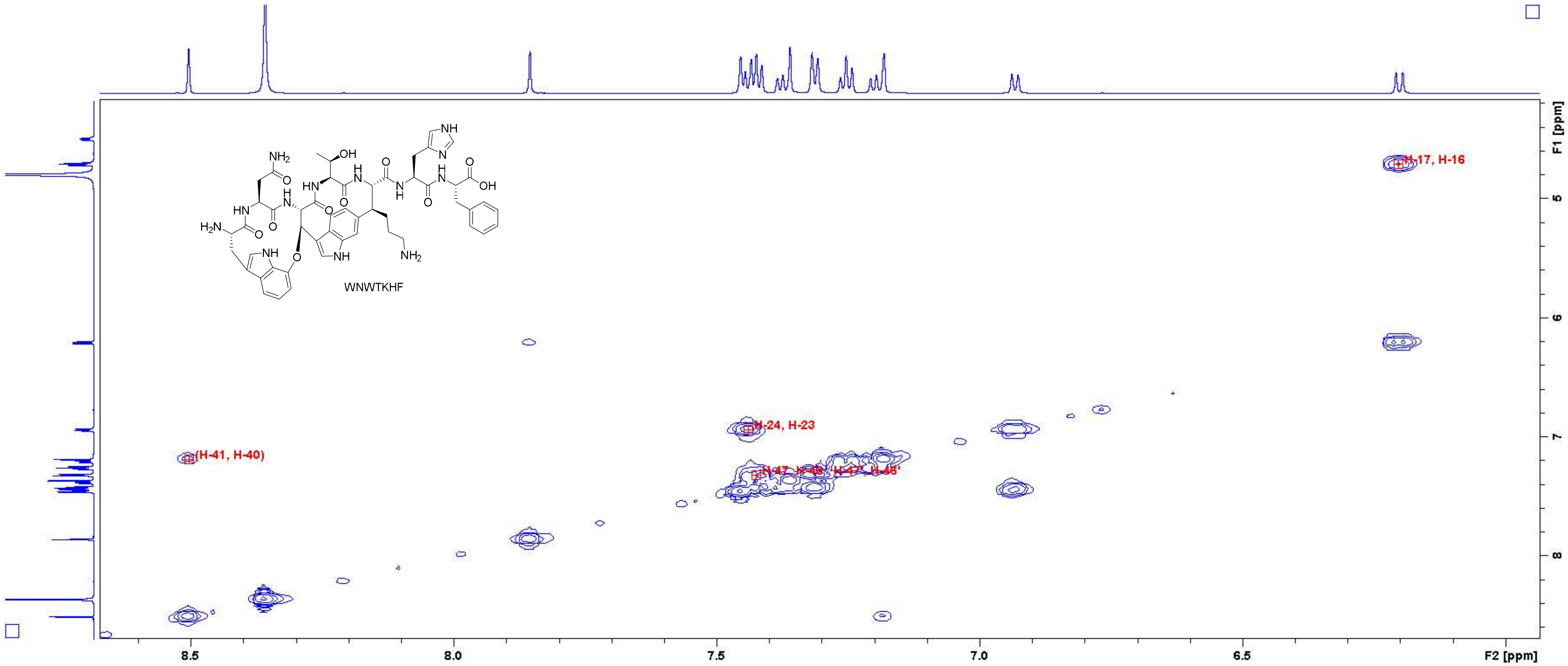

Figure S38: COSY spectrum of WNWTKHF in $D_2O$ (700 MHz). Close-up in the region of 6.0 – 8.6 ppm (F2 axis) and 4.2 – 8.6 ppm (F1 axis) with peak assignments. For atom numbering, cf. Figure S3.

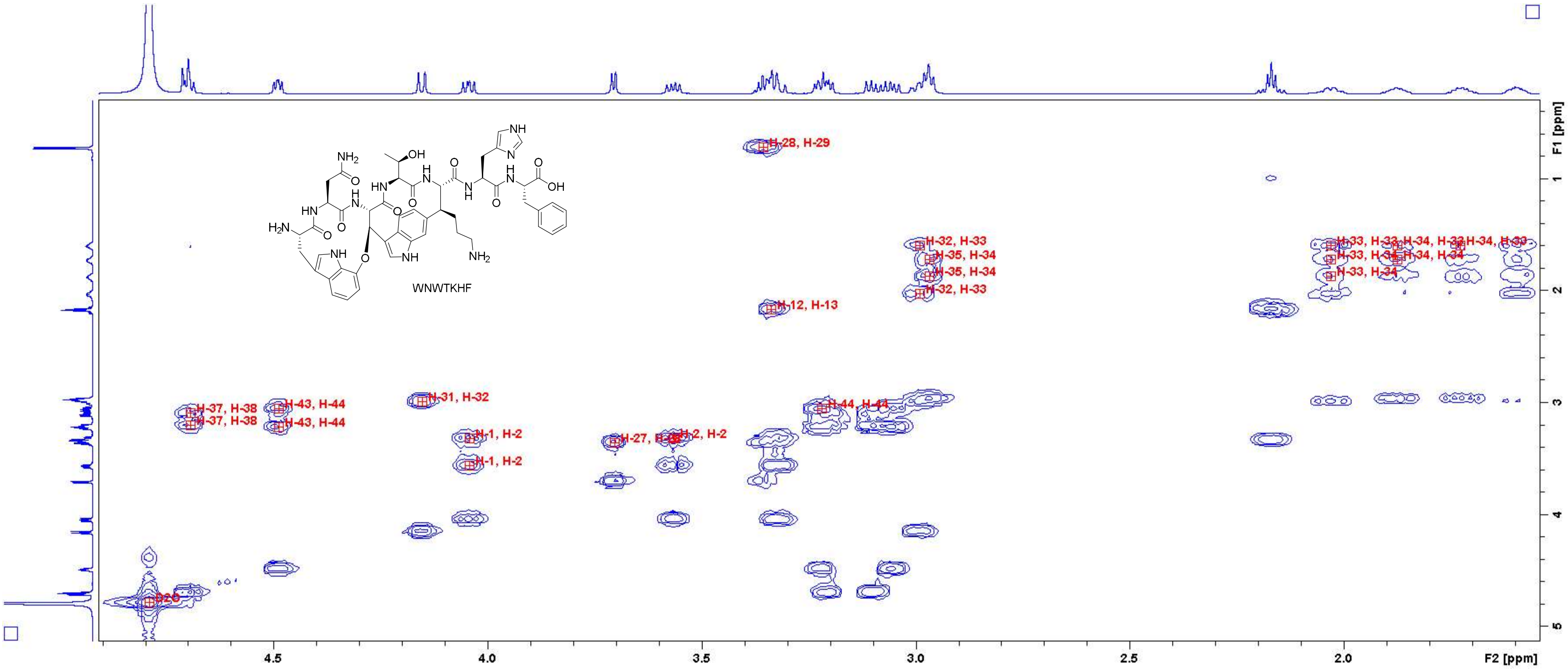

Figure S39: COSY spectrum of WNWTKHF in $D_2O$ (700 MHz). Close-up in the region of 1.6 – 4.9 ppm (F2 axis) and 0.4 – 5.0 ppm (F1 axis) with peak assignments. For atom numbering, cf. Figure S3.

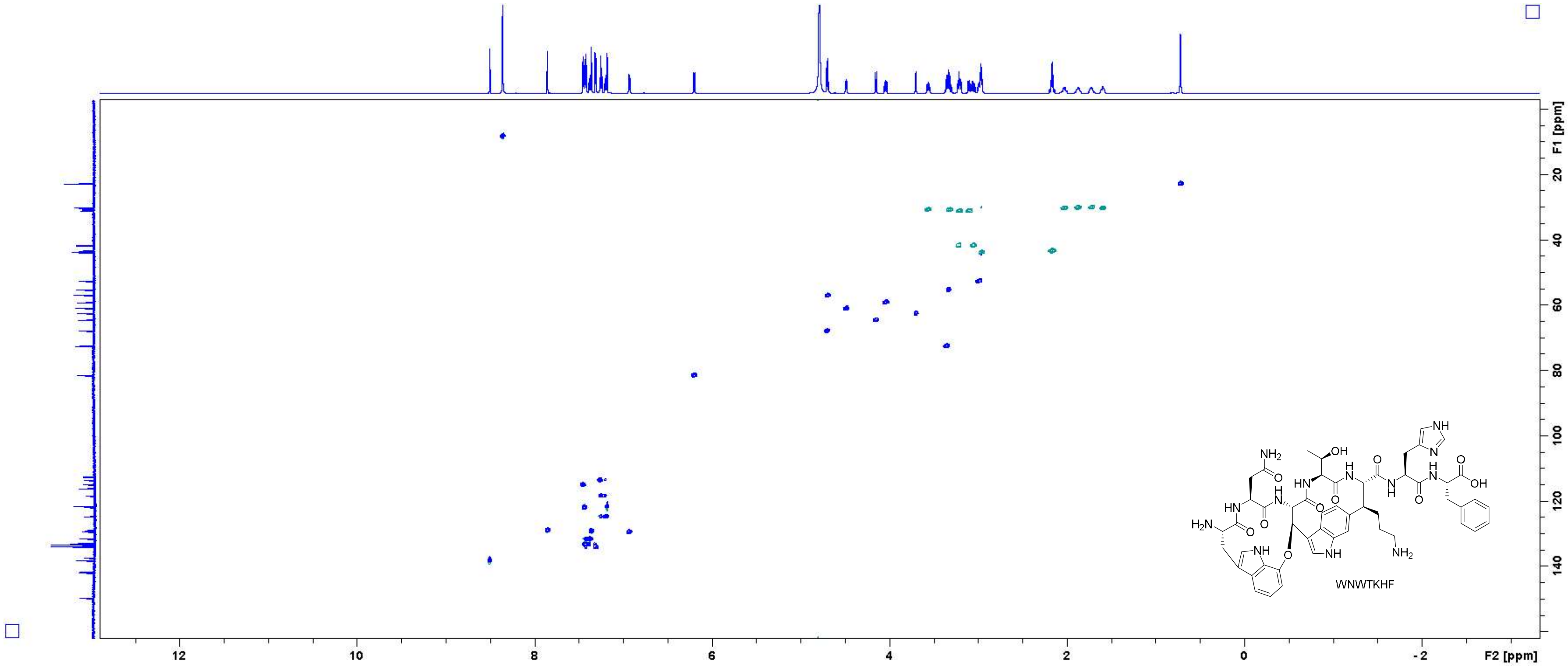

Figure S40: HSQC spectrum of WNWTKHF in $D_2O$ (700 MHz).

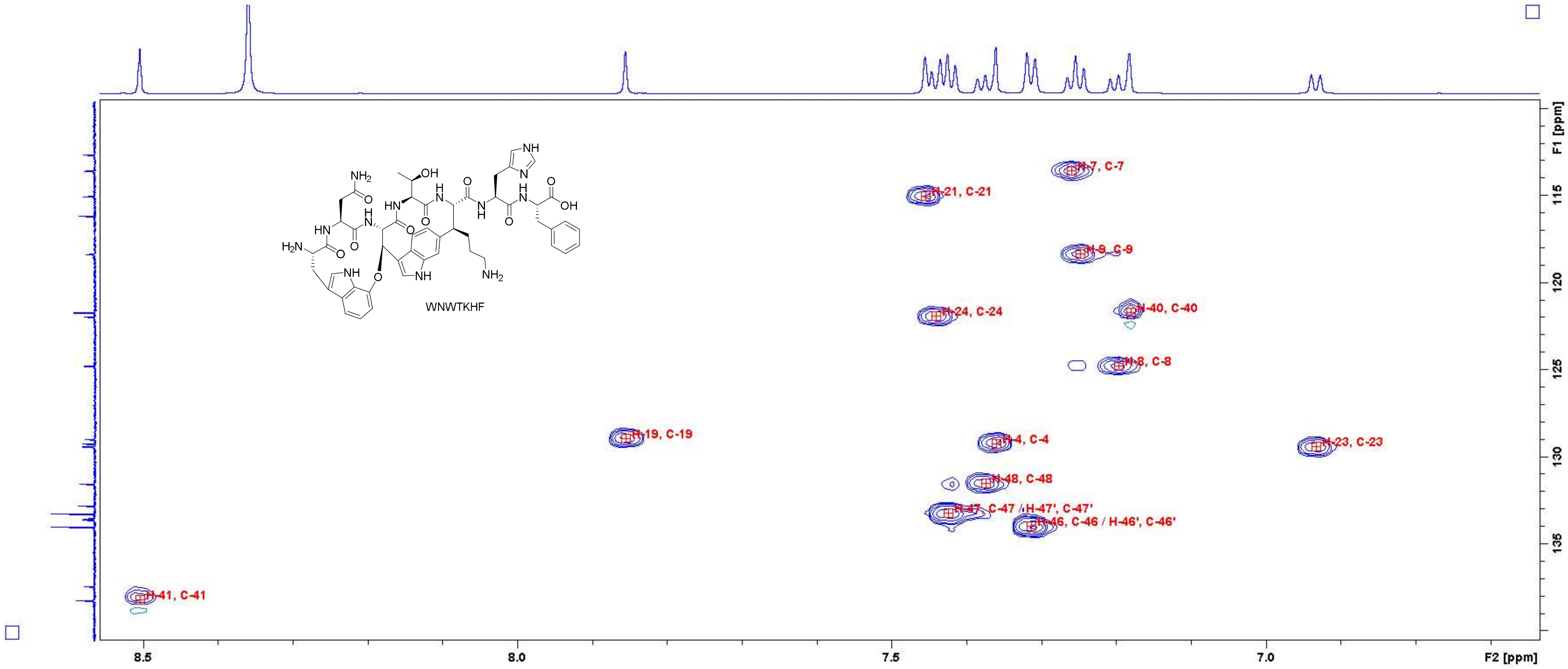

Figure S41: HSQC spectrum of WNWTKHF in $D_2O$ (700 MHz). Close-up in the region of 6.7 – 8.5 ppm (F2 axis) and 110 – 140 ppm (F1 axis) with peak assignments. For atom numbering, cf. Figure S3.

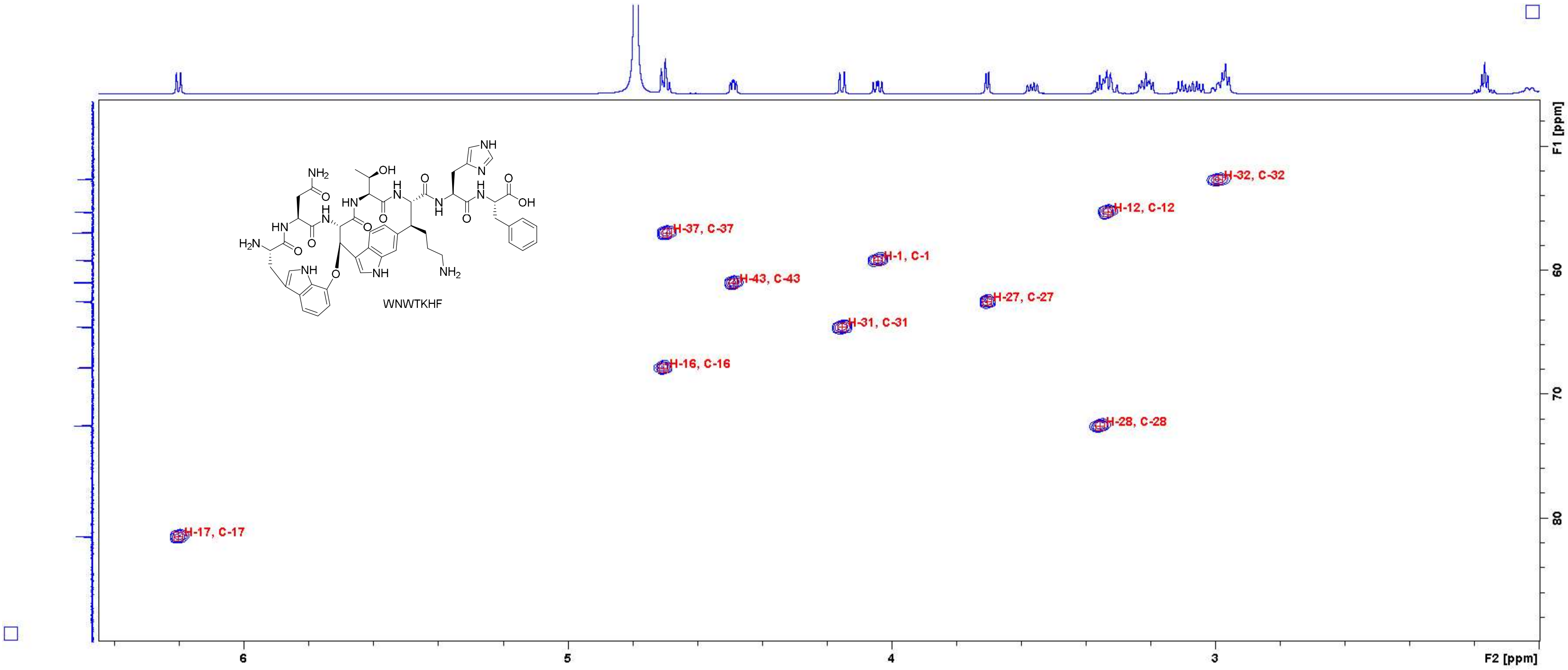

**Figure S42:** HSQC spectrum of **WNWTKHF** in $D_2O$ (700 MHz). Close-up in the region of 2.0 – 6.4 ppm (F2 axis) and 48 – 88 ppm (F1 axis) with peak assignments. For atom numbering, cf. Figure S3.

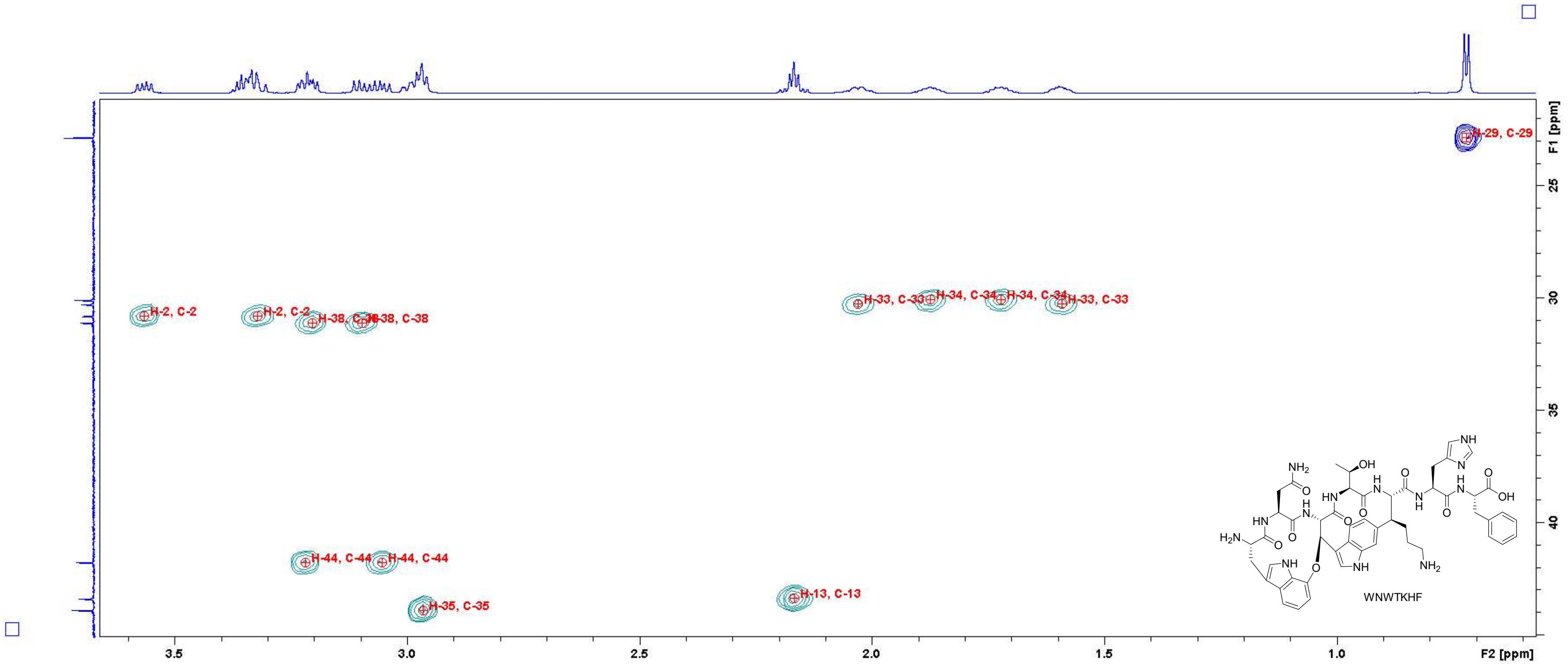

**Figure S43:** HSQC spectrum of **WNWTKHF** in $D_2O$ (700 MHz). Close-up in the region of 0.6 – 3.6 ppm (F2 axis) and 22 – 45 ppm (F1 axis) with peak assignments. For atom numbering, cf. Figure S3.

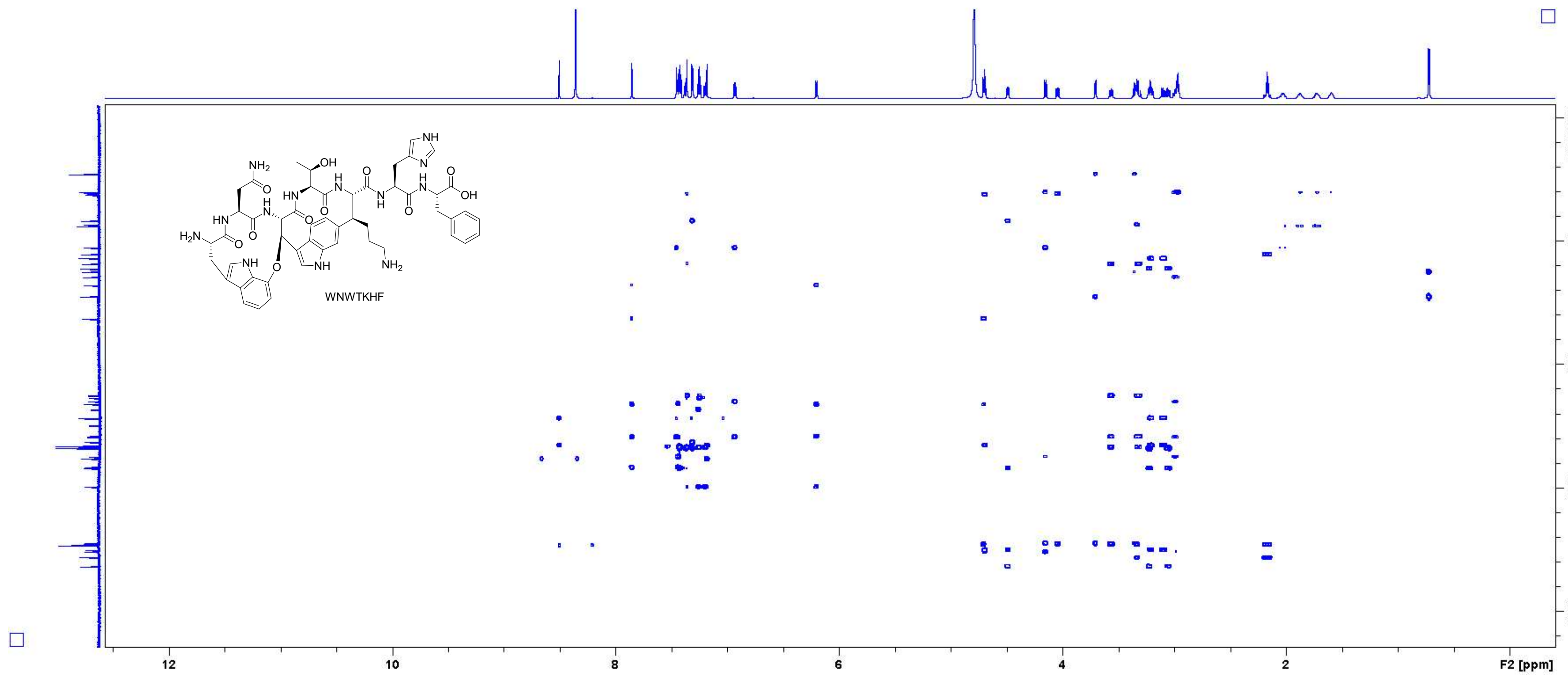

Figure S44: HMBC spectrum of WNWTKHF in $D_2O$ (700 MHz).

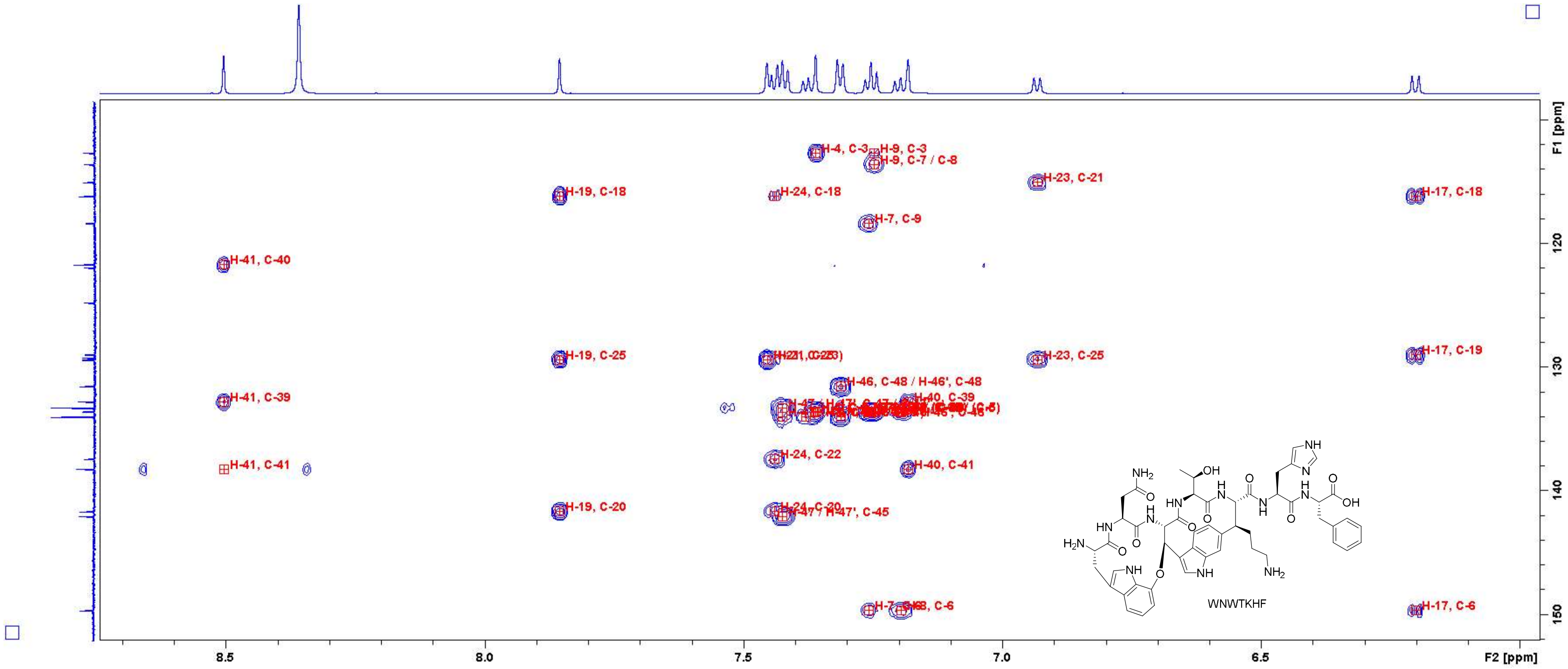

Figure S45: HMBC spectrum of WNWTKHF in D₂O (700 MHz). Close-up in the region of 6.0 – 8.7 ppm (F2 axis) and 110 – 152 ppm (F1 axis) with peak assignments. For atom numbering, cf. Figure S3.

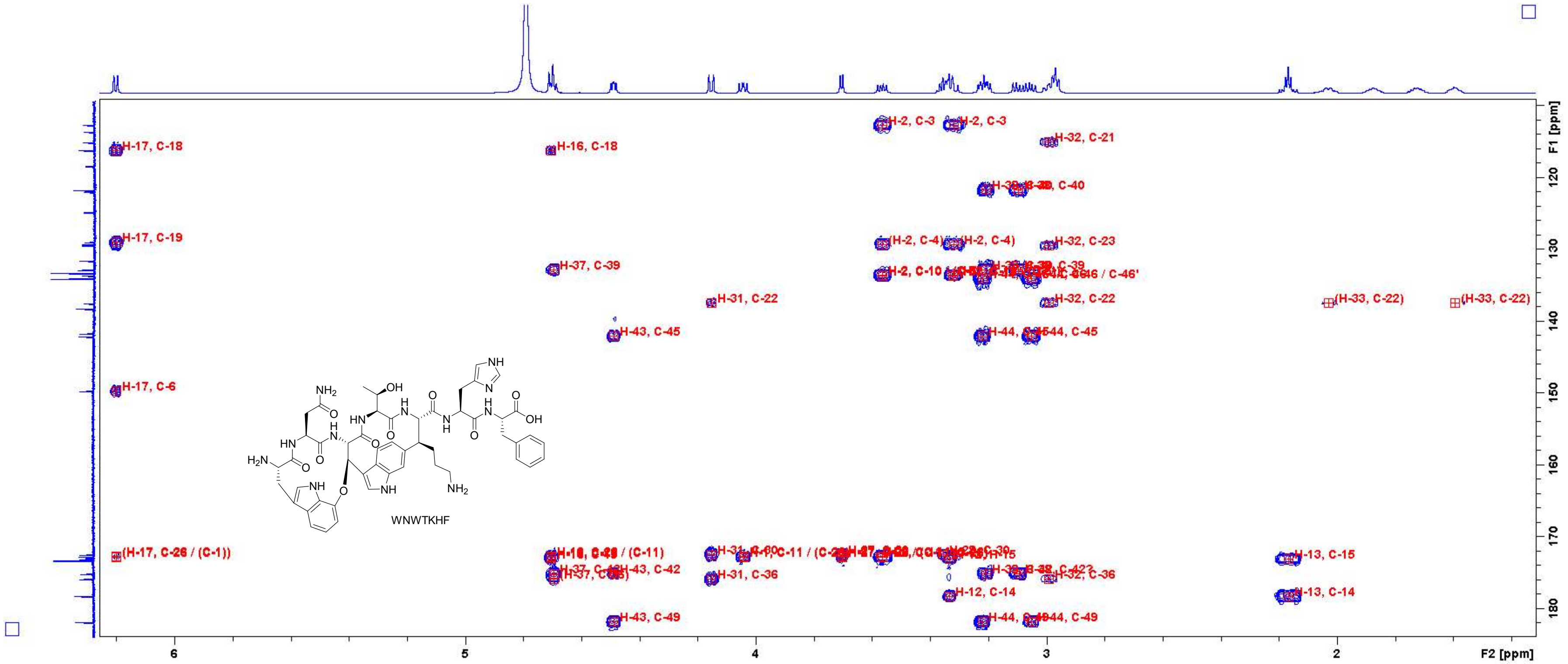

Figure S46: HMBC spectrum of WNWTKHF in $D_2O$ (700 MHz). Close-up in the region of 1.4 – 6.2 ppm (F2 axis) and 110 – 182 ppm (F1 axis) with peak assignments. For atom numbering, cf. Figure S3.

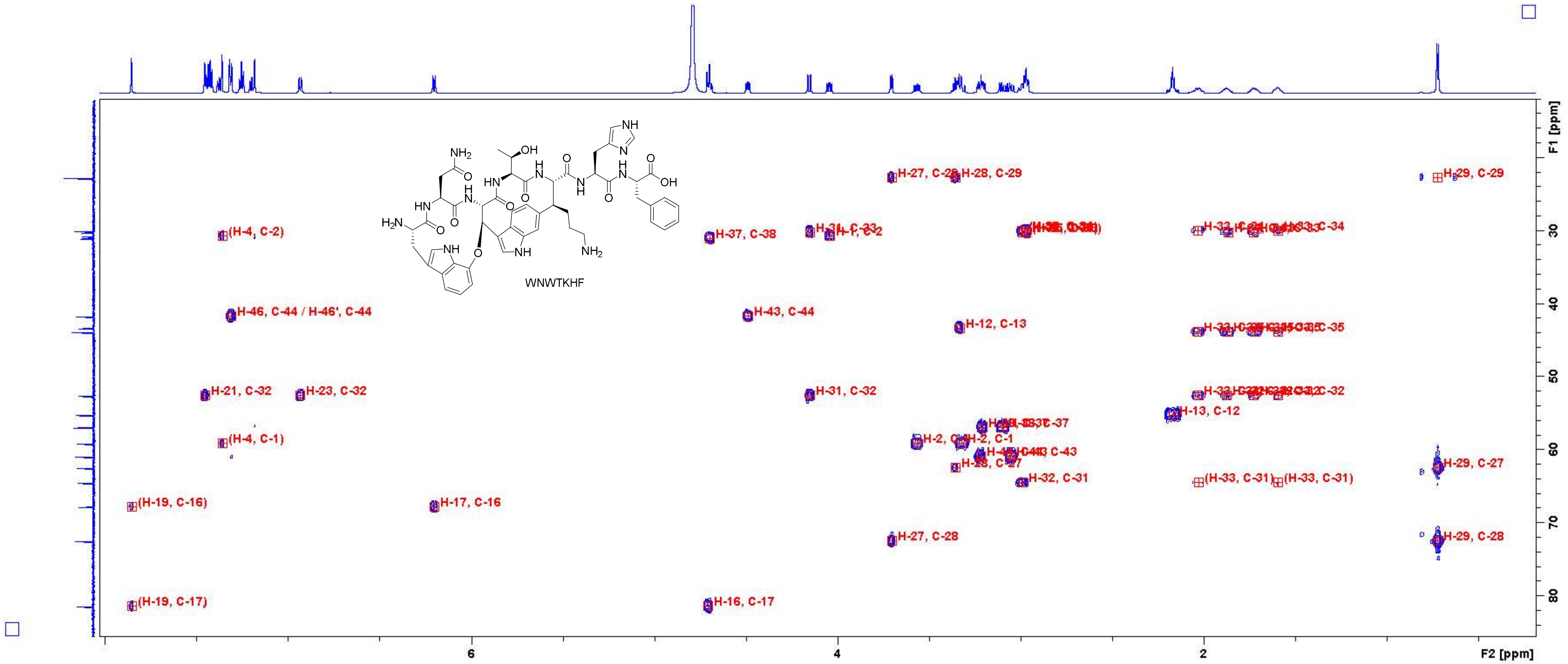

**Figure S47:** HMBC spectrum of **WNWTKHF** in $D_2O$ (700 MHz). Close-up in the region of 0.5 – 8.0 ppm (F2 axis) and 12 – 84 ppm (F1 axis) with peak assignments. For atom numbering, cf. Figure S3.

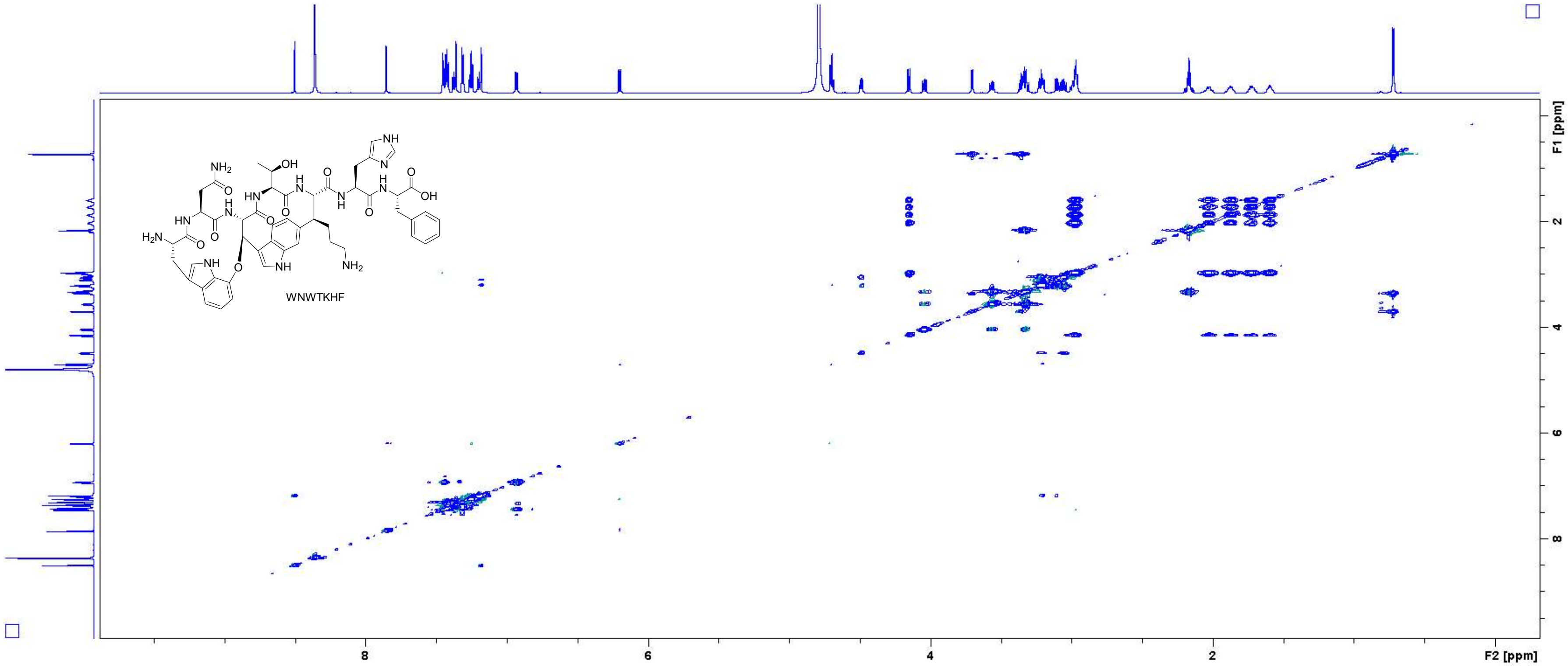

Figure S48: TOCSY spectrum of **WNWTKHF** in $D_2O$ (700 MHz, measured with $H_2O$ suppression).

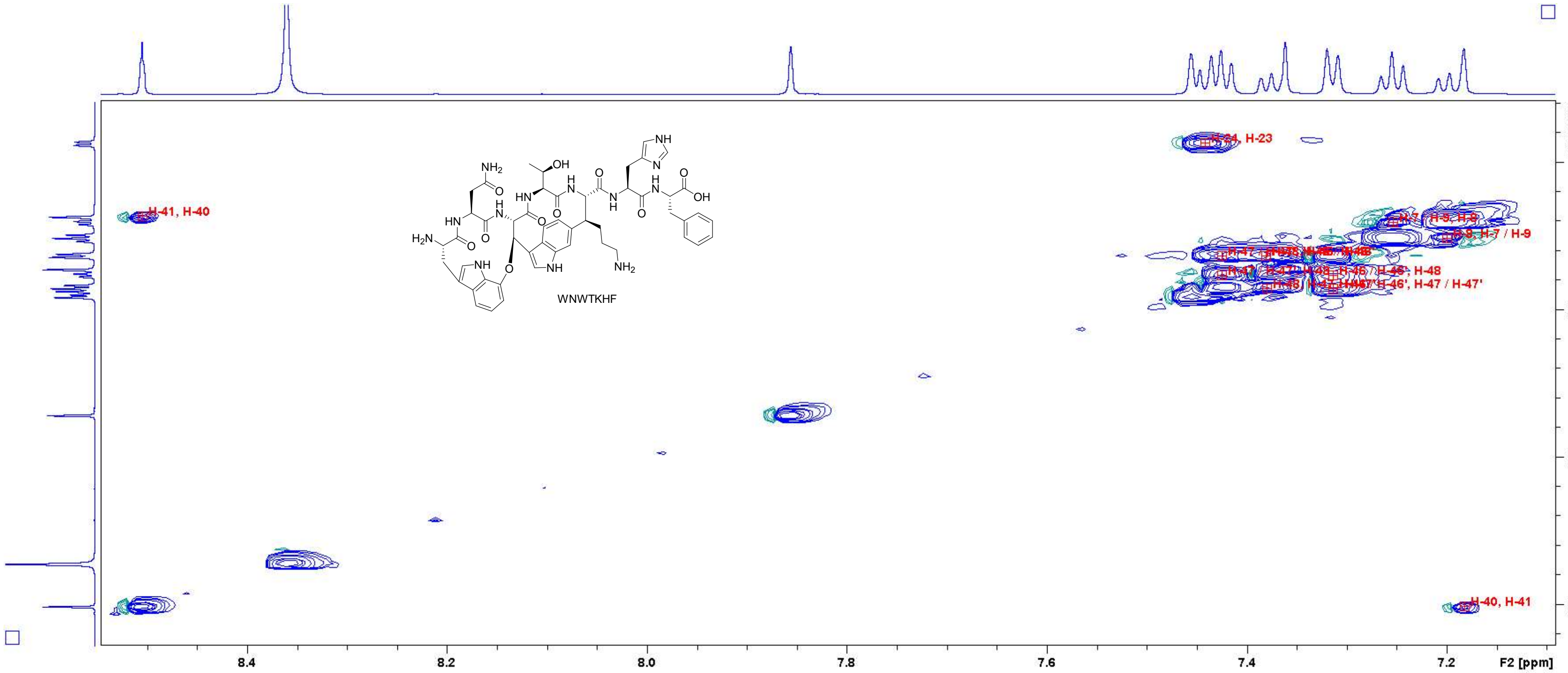

Figure S49: TOCSY spectrum of WNWTKHF in $D_2O$ (700 MHz, measured with $H_2O$ suppression). Close-up in the region of 7.1 – 8.5 ppm (F2 axis) and 6.8 – 8.6 ppm (F1 axis) with peak assignments. For atom numbering, cf. Figure S3.

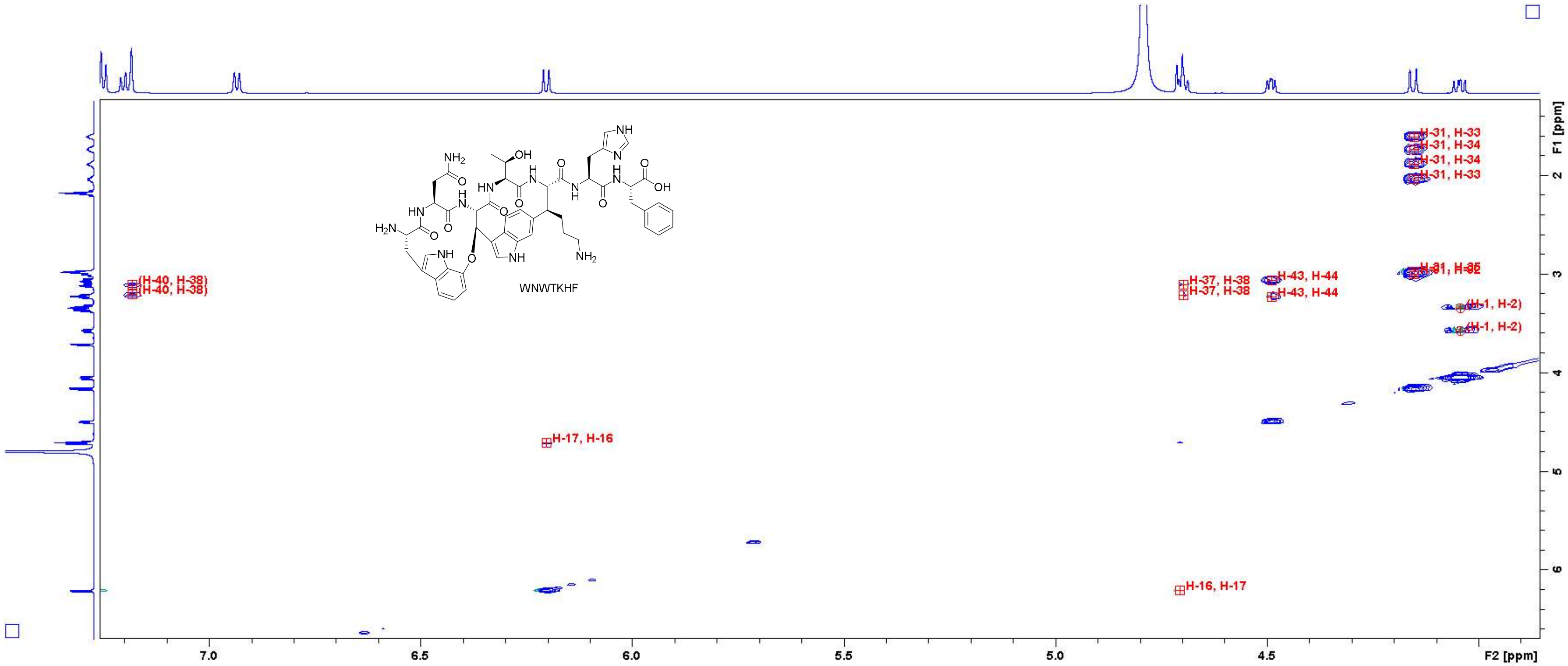

Figure S50: TOCSY spectrum of WNWTKHF in $D_2O$ (700 MHz, measured with $H_2O$ suppression). Close-up in the region of 3.9 – 7.2 ppm (F2 axis) and 1.4 – 6.6 ppm (F1 axis) with peak assignments. For atom numbering, cf. Figure S3.

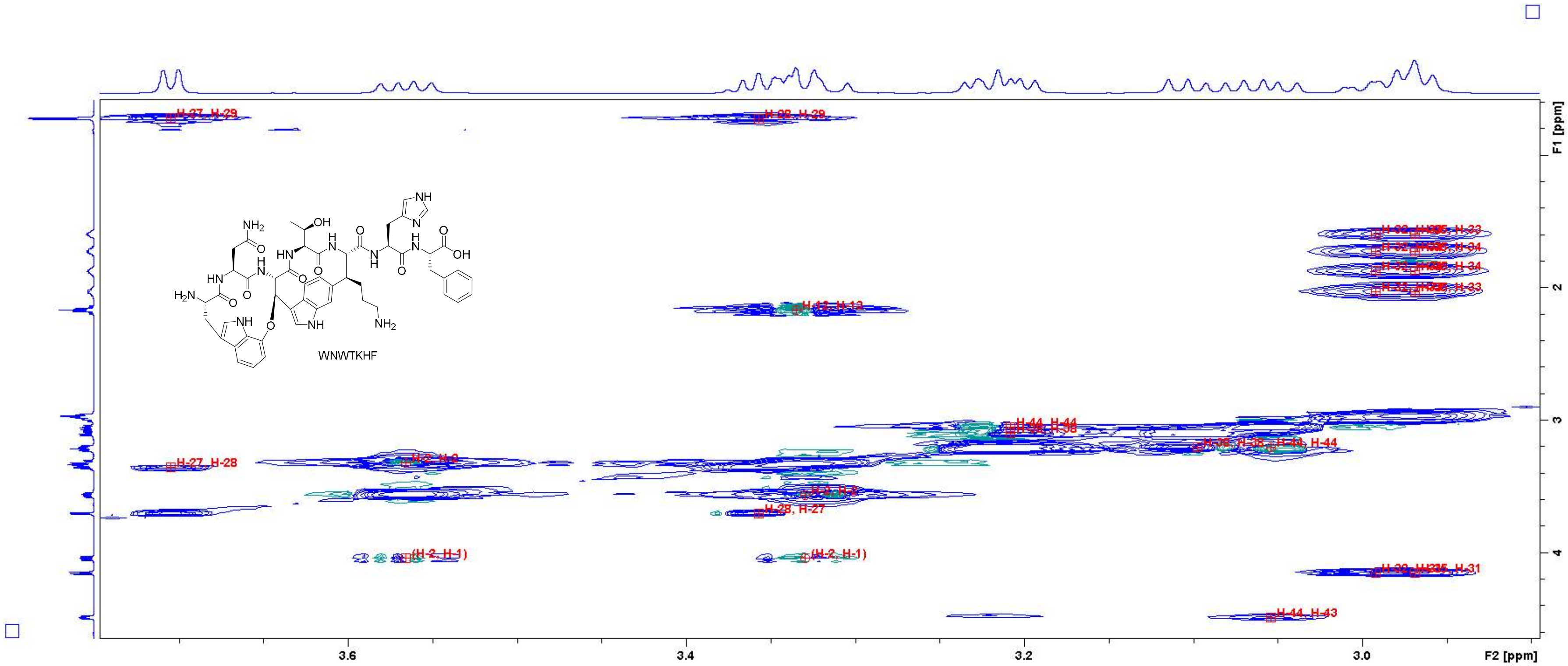

Figure S51: TOCSY spectrum of WNWTKHF in $D_2O$ (700 MHz, measured with $H_2O$ suppression). Close-up in the region of 2.9 – 3.7 ppm (F2 axis) and 0.6 – 4.6 ppm (F1 axis) with peak assignments. For atom numbering, cf. Figure S3.

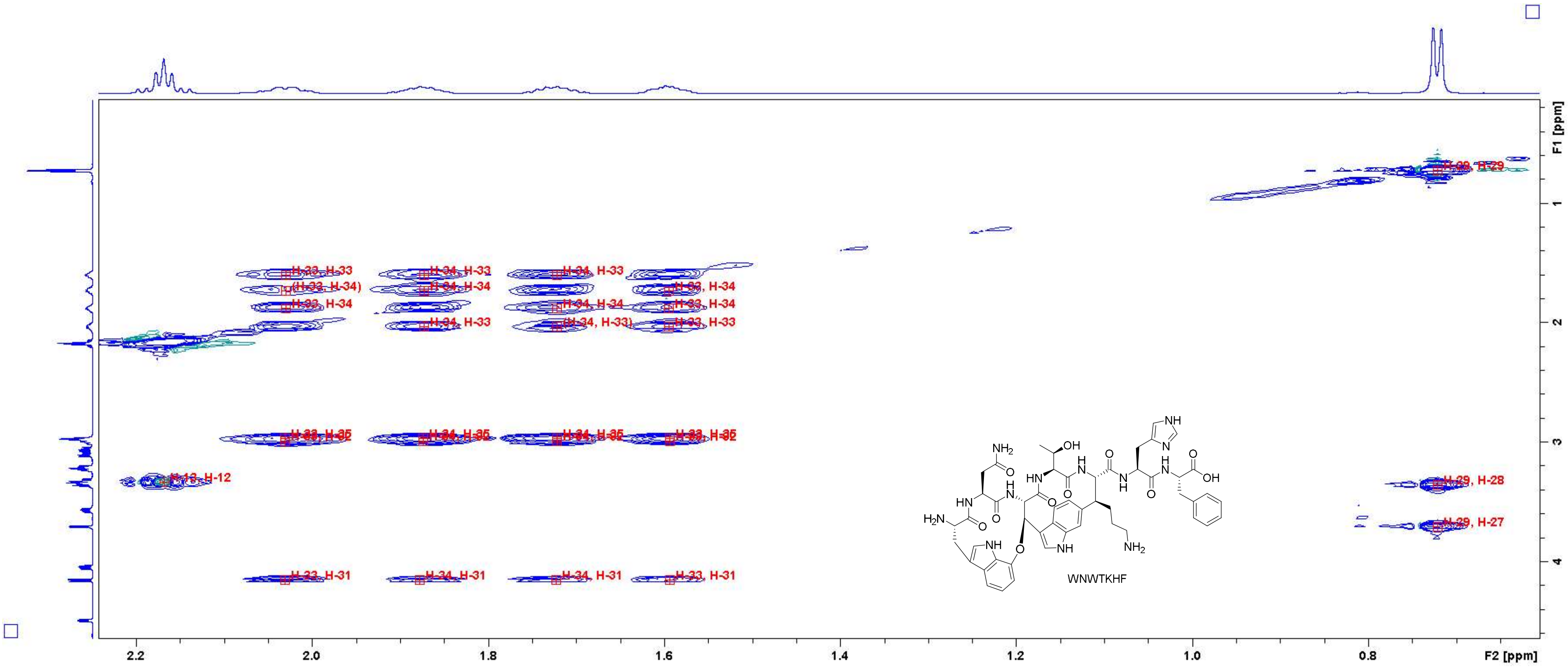

Figure S52: TOCSY spectrum of WNWTKHF in $D_2O$ (700 MHz, measured with $H_2O$ suppression). Close-up in the region of 0.65 – 2.20 ppm (F2 axis) and 0.2 – 4.6 ppm (F1 axis) with peak assignments. For atom numbering, cf. Figure S3.

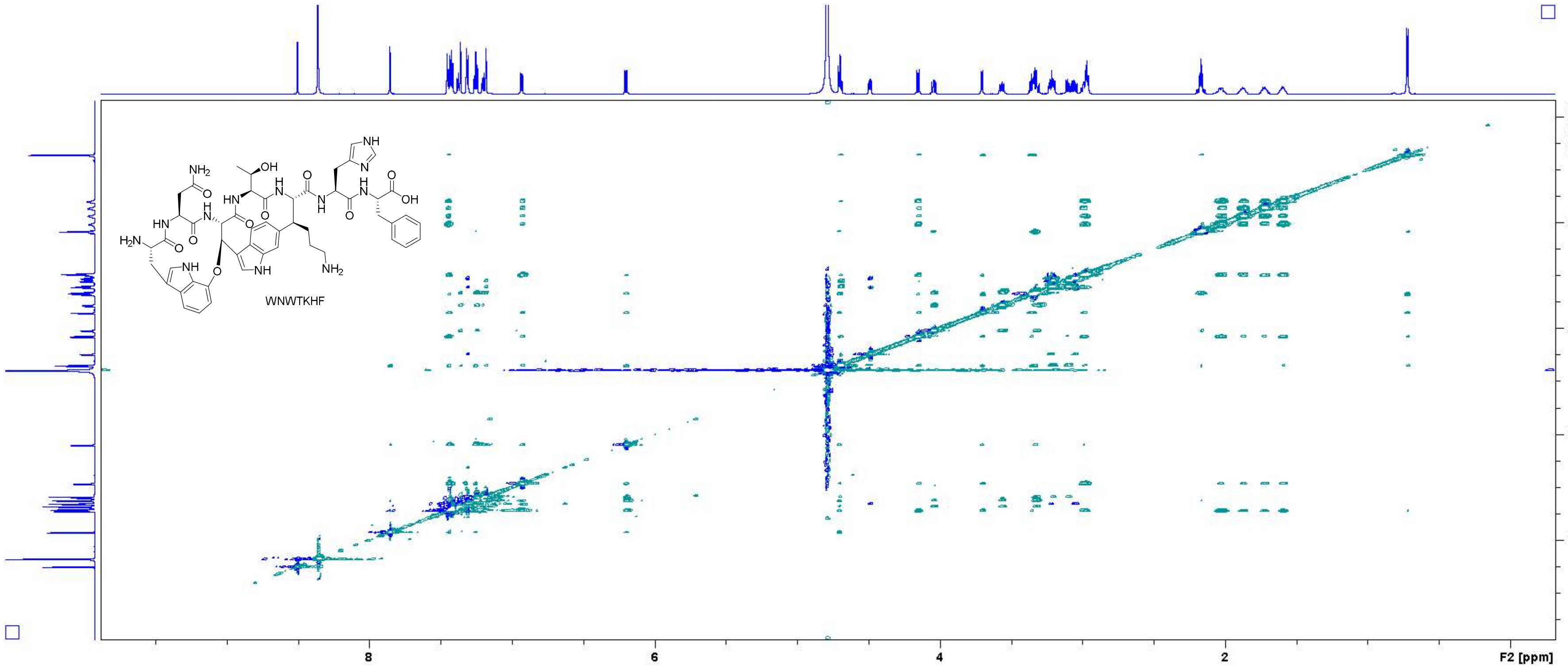

Figure S53: NOESY spectrum of WNWTKHF in $D_2O$ (700 MHz).

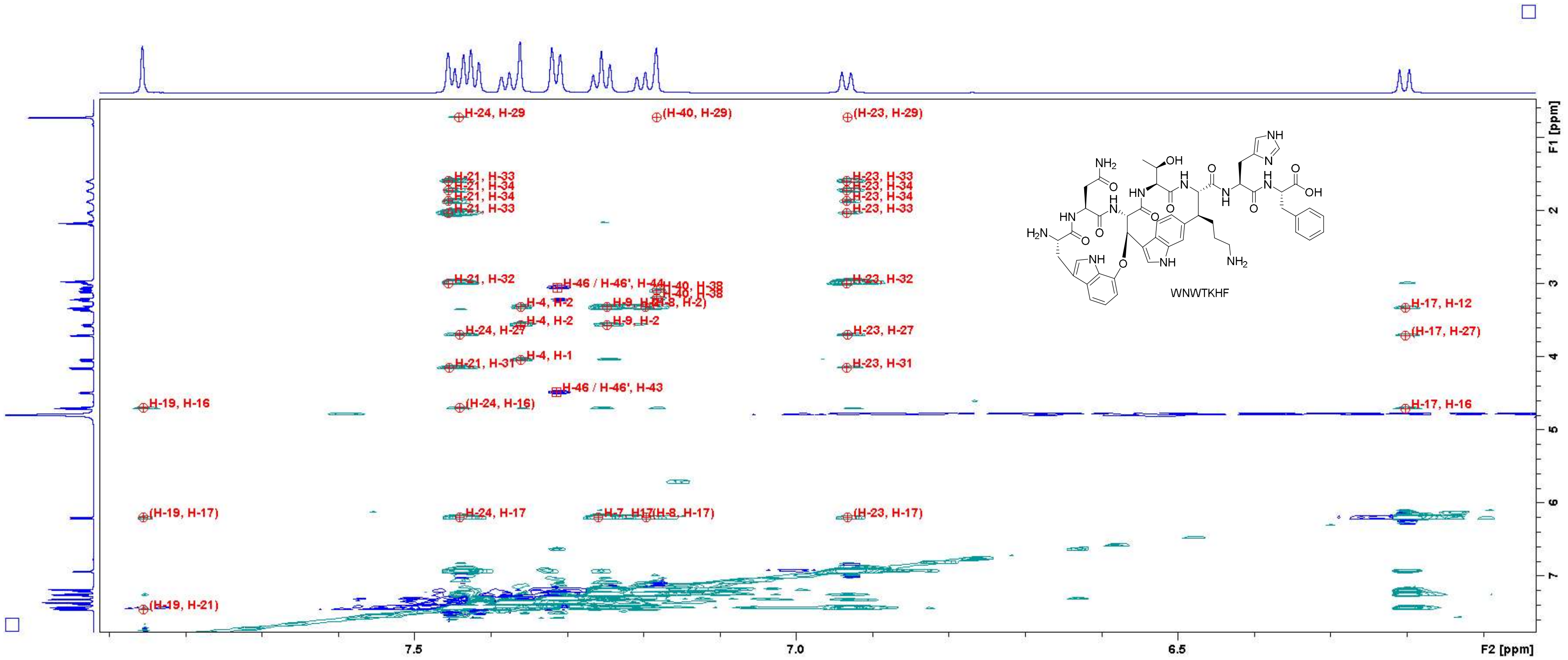

Figure S54: NOESY spectrum of WNWTKHF in D$_2$O (700 MHz). Close-up in the region of 6.1 – 7.9 ppm (F2 axis) and 0.6 – 7.6 ppm (F1 axis) with peak assignments. For atom numbering, cf. Figure S3.

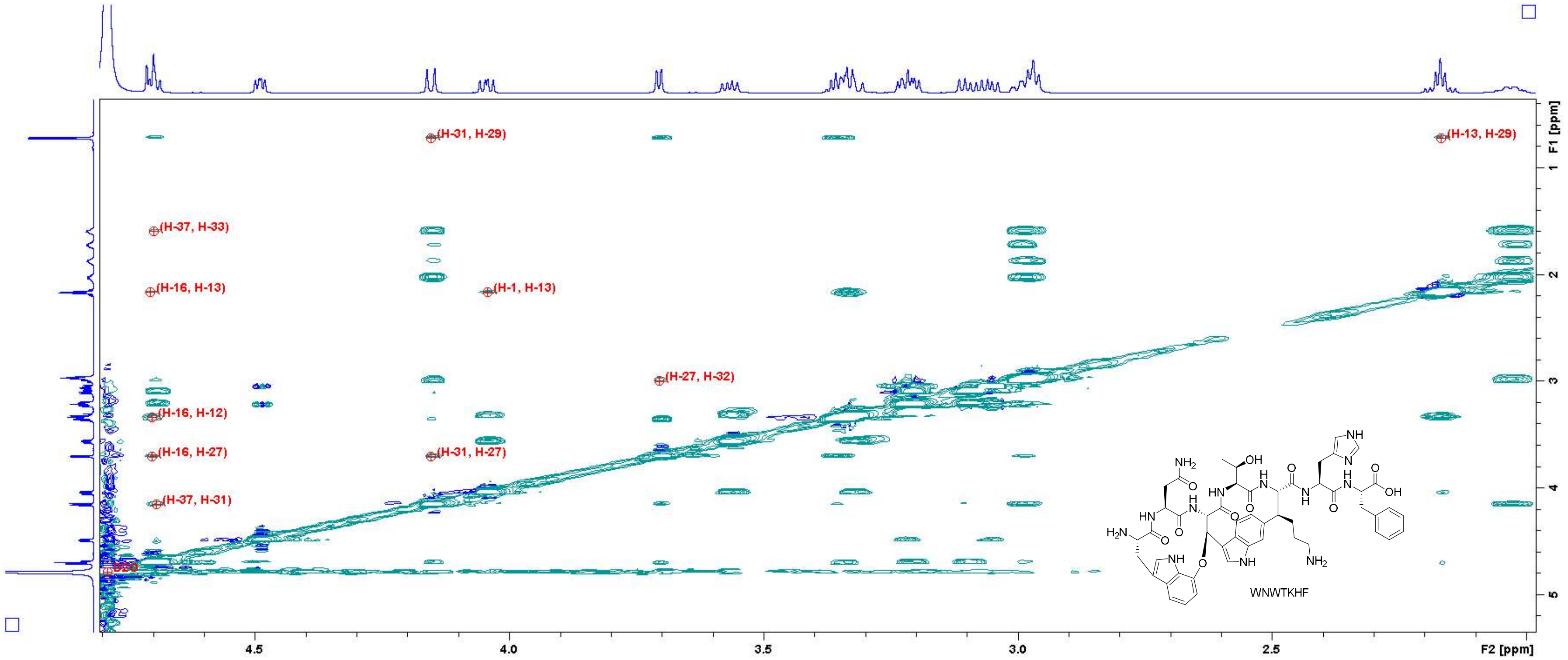

Figure S55: NOESY spectrum of WNWTKHF in $D_2O$ (700 MHz). Close-up in the region of 2.0 – 4.8 ppm (F2 axis) and 0.4 – 5.2 ppm (F1 axis) with peak assignments. For atom numbering, cf. Figure S3.

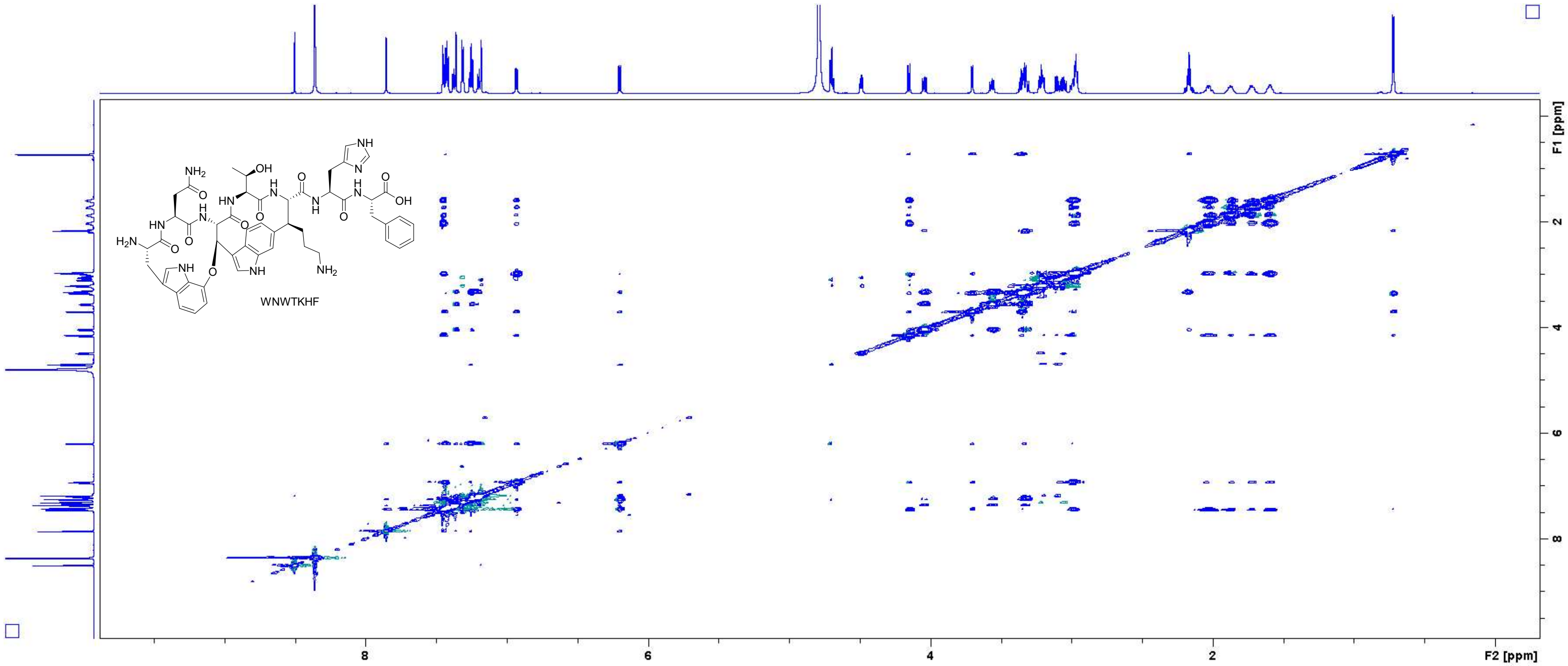

**Figure S56**: NOESY spectrum of **WNWTKHF** in $D_2O$ (700 MHz, measured with $H_2O$ suppression).

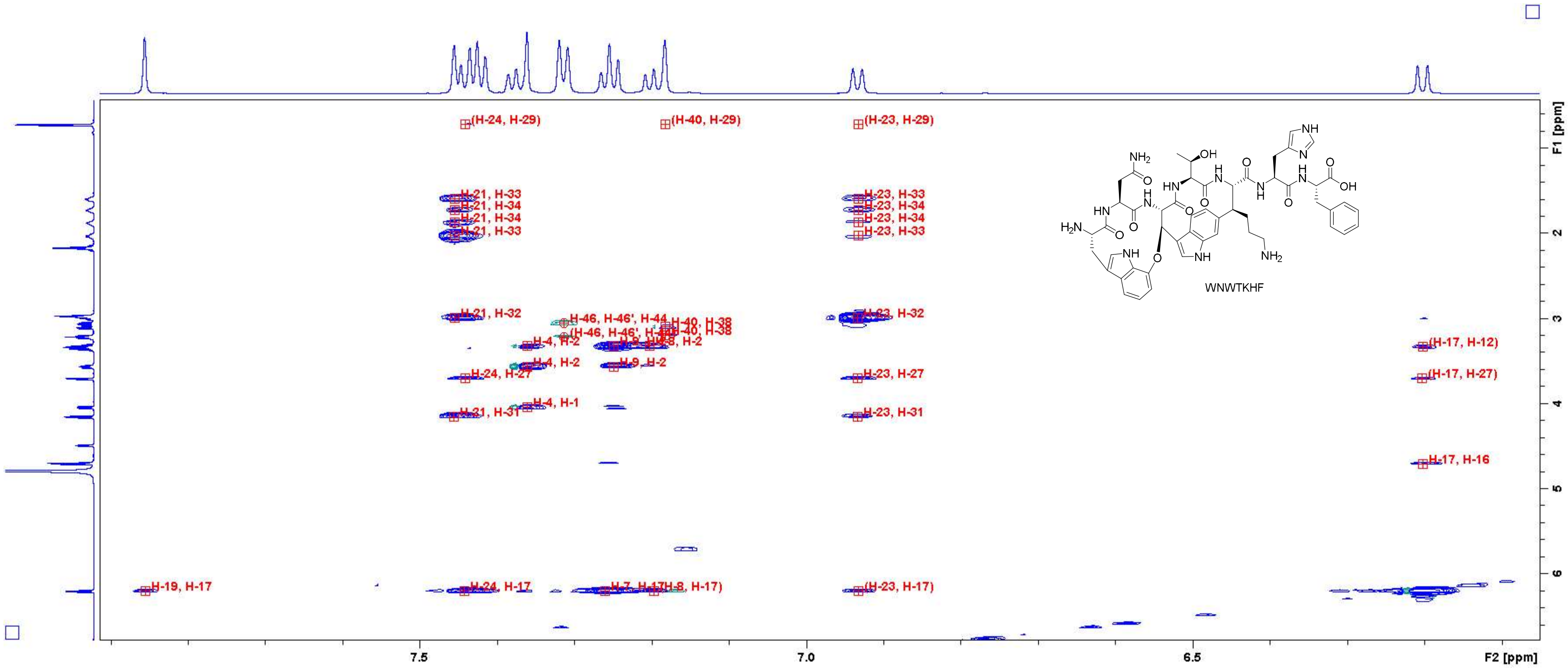

Figure S57: NOESY spectrum of WNWTKHF in $D_2O$ (700 MHz, measured with $H_2O$ suppression). Close-up in the region of 6.1 – 7.9 ppm (F2 axis) and 0.6 – 6.6 ppm (F1 axis) with peak assignments. For atom numbering, cf. Figure S3.

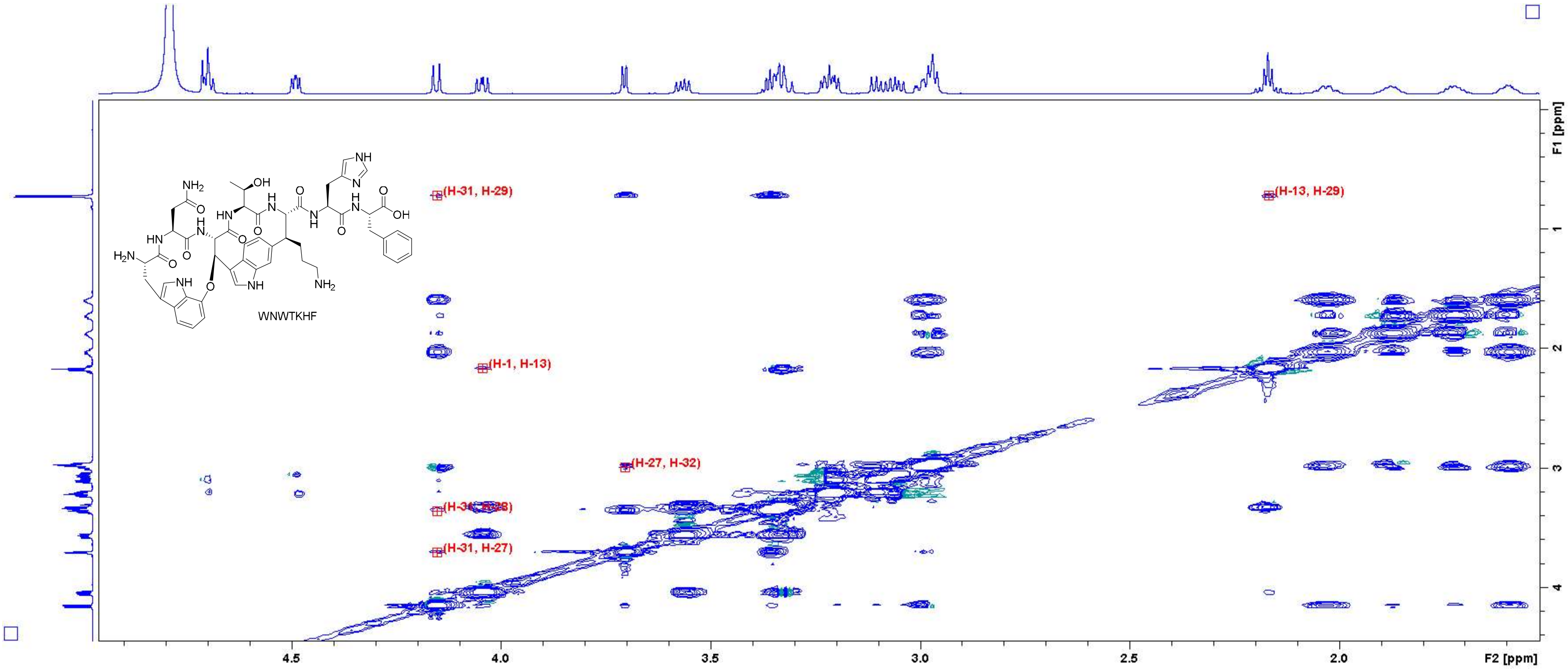

Figure S58: NOESY spectrum of WNWTKHF in $D_2O$ (700 MHz, measured with $H_2O$ suppression). Close-up in the region of 1.6 – 4.9 ppm (F2 axis) and 0.0 – 4.4 ppm (F1 axis) with peak assignments. For atom numbering, cf. Figure S3.

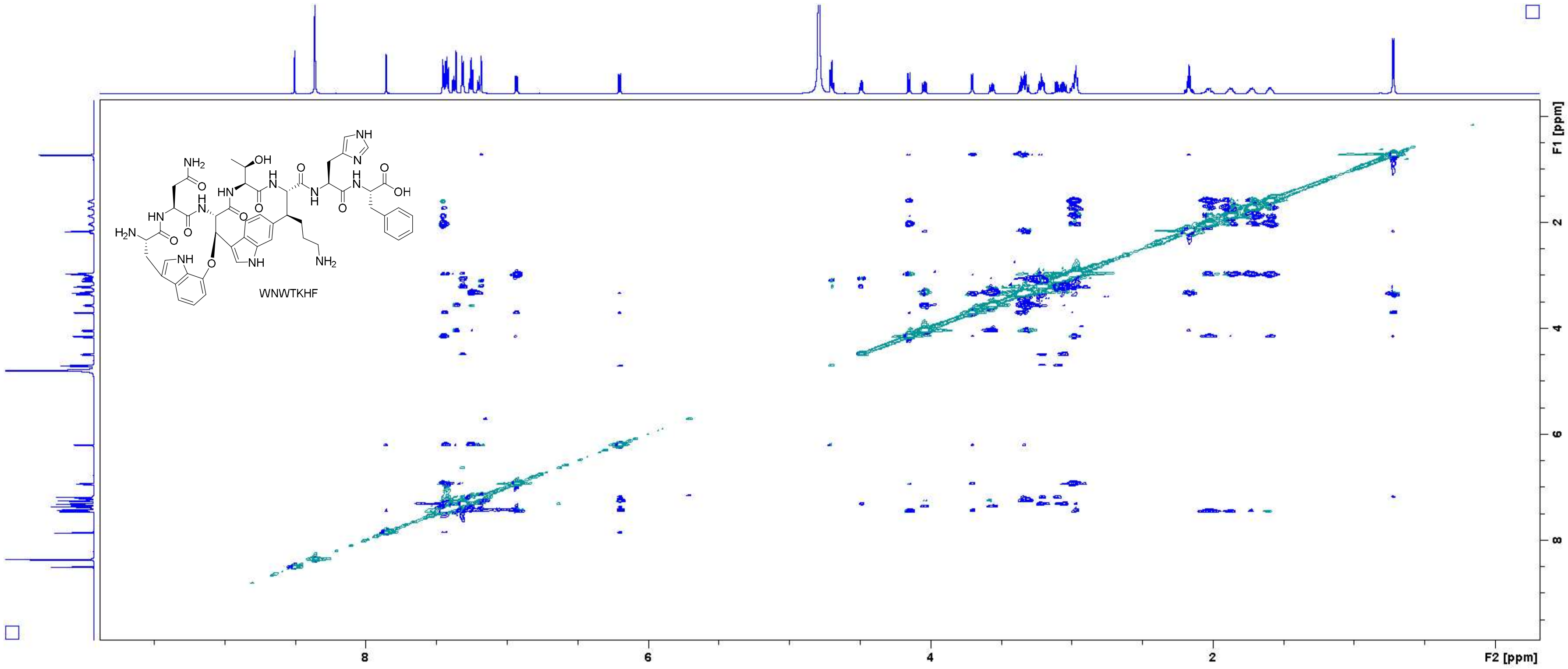

Figure S59: ROESY spectrum of WNWTKHF in $D_2O$ (700 MHz, measured with $H_2O$ suppression).

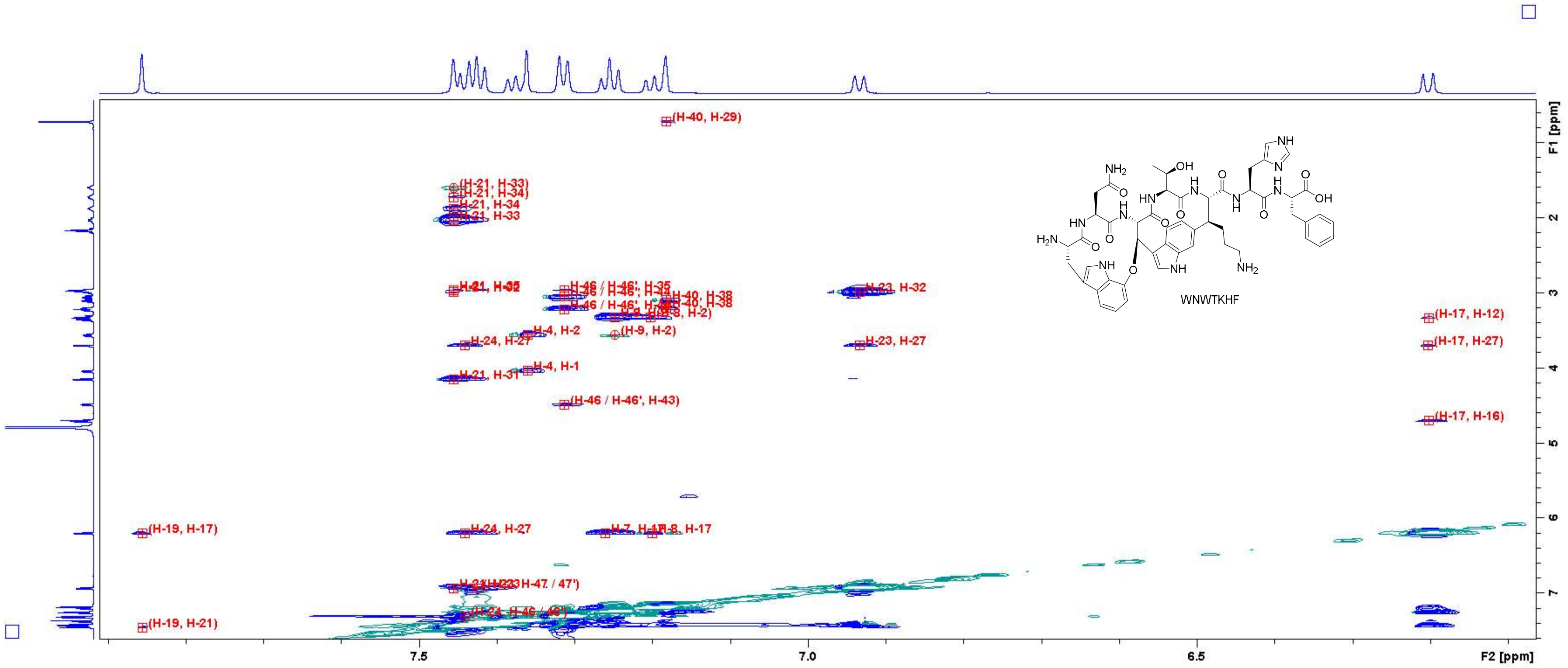

Figure S60: ROESY spectrum of WNWTKHF in $D_2O$ (700 MHz, measured with $H_2O$ suppression). Close-up in the region of 6.1 – 7.9 ppm (F2 axis) and 0.6 – 7.6 ppm (F1 axis) with peak assignments. For atom numbering, cf. Figure S3.

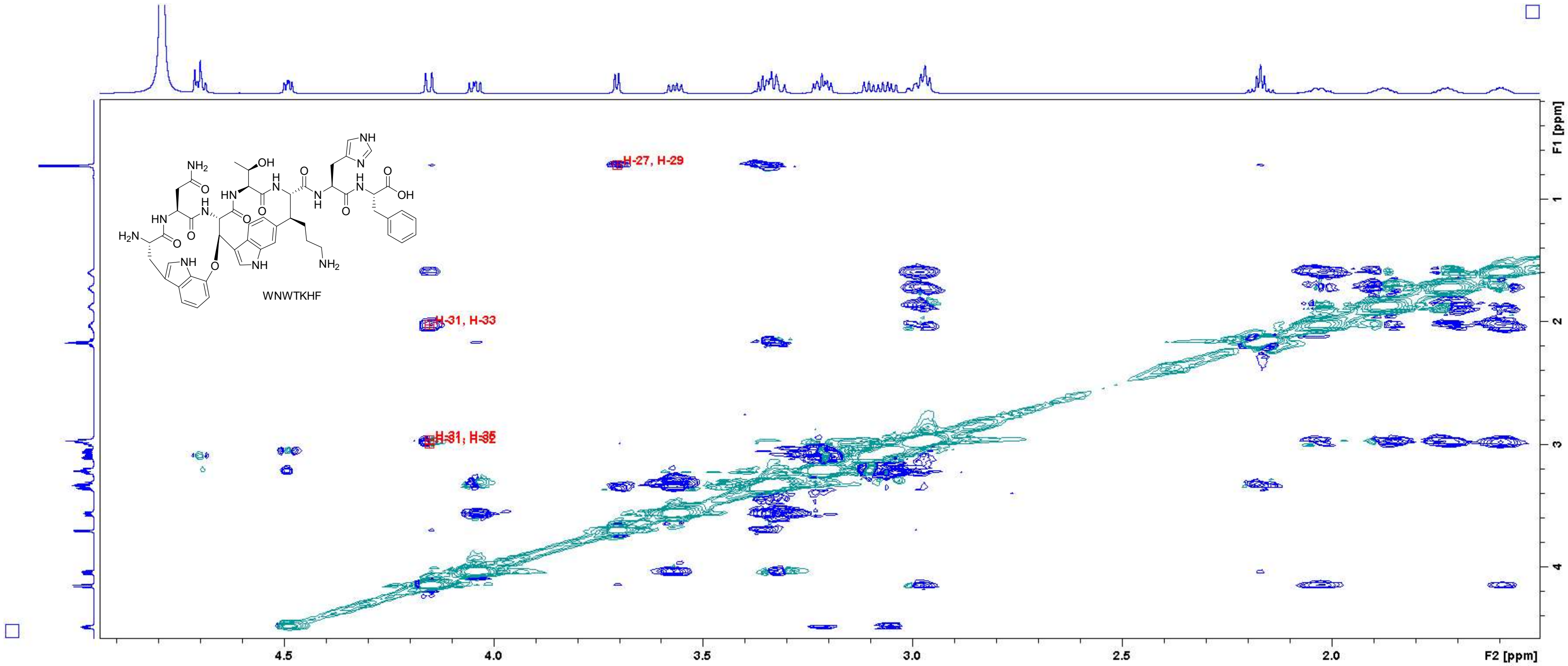

Figure S61: ROESY spectrum of **WNWTKHF** in $D_2O$ (700 MHz, measured with $H_2O$ suppression). Close-up in the region of 1.6 – 4.9 ppm (F2 axis) and 0.2 – 4.4 ppm (F1 axis) with peak assignments. For atom numbering, cf. Figure S3.

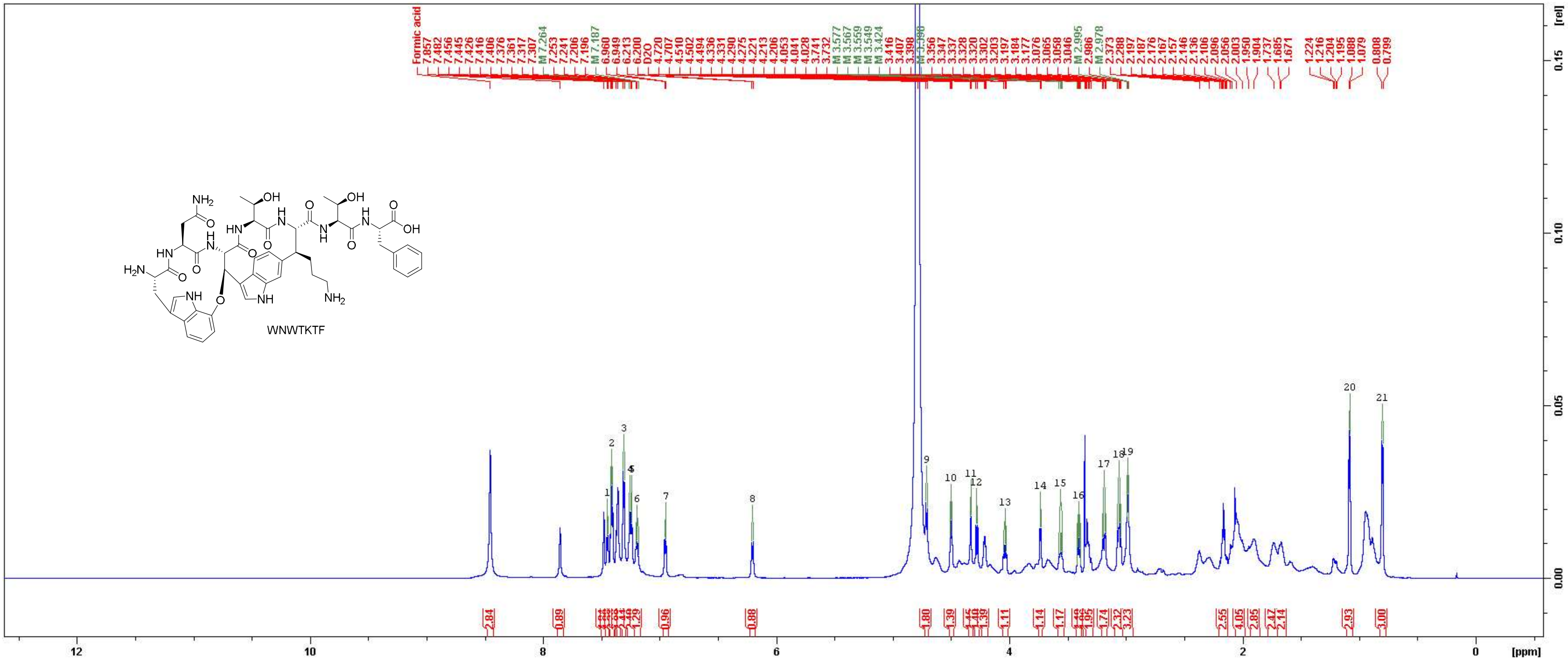

Figure S62: $^{1}H$ NMR spectrum of WNWTKTF in $D_2O$ (700 MHz).

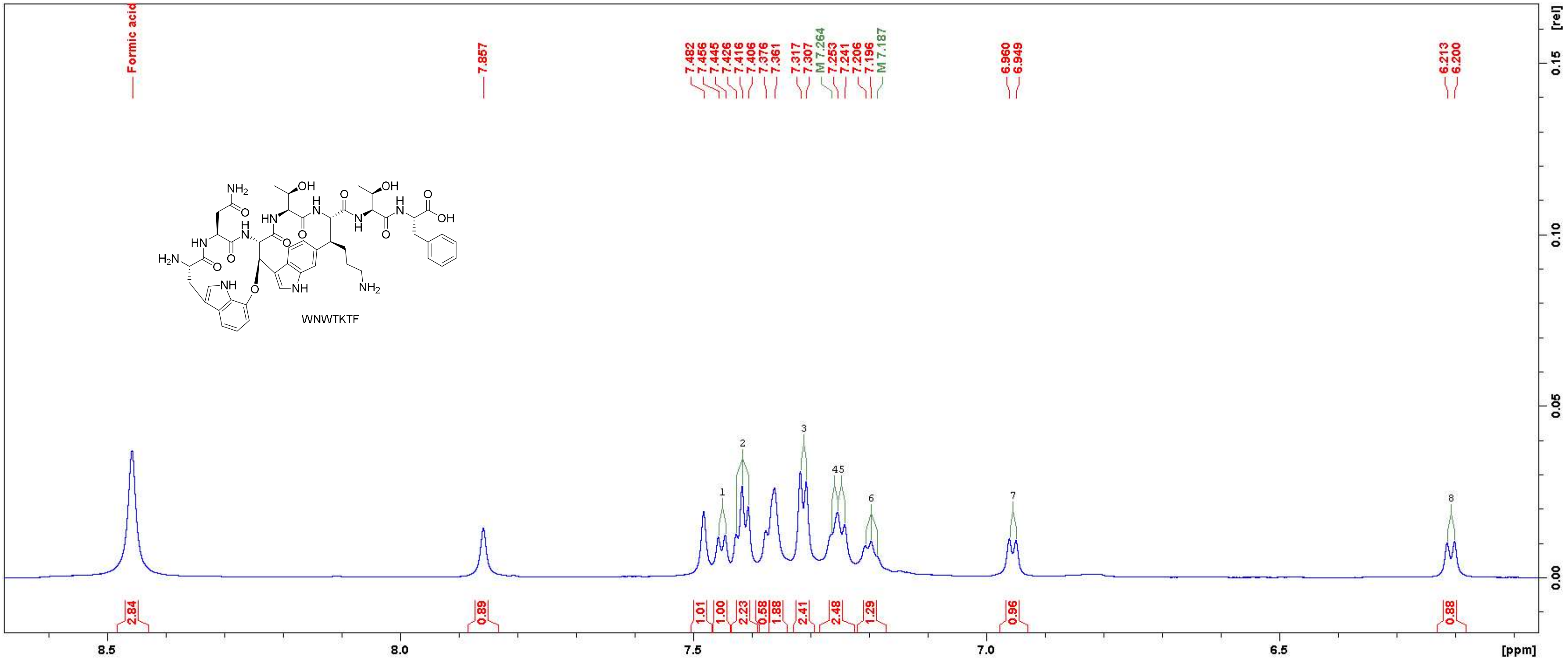

Figure S63: [1]H NMR spectrum of WNWTKTF in $D_2O$ (700 MHz). Close-up in the region between 6.1 ppm and 8.6 ppm.

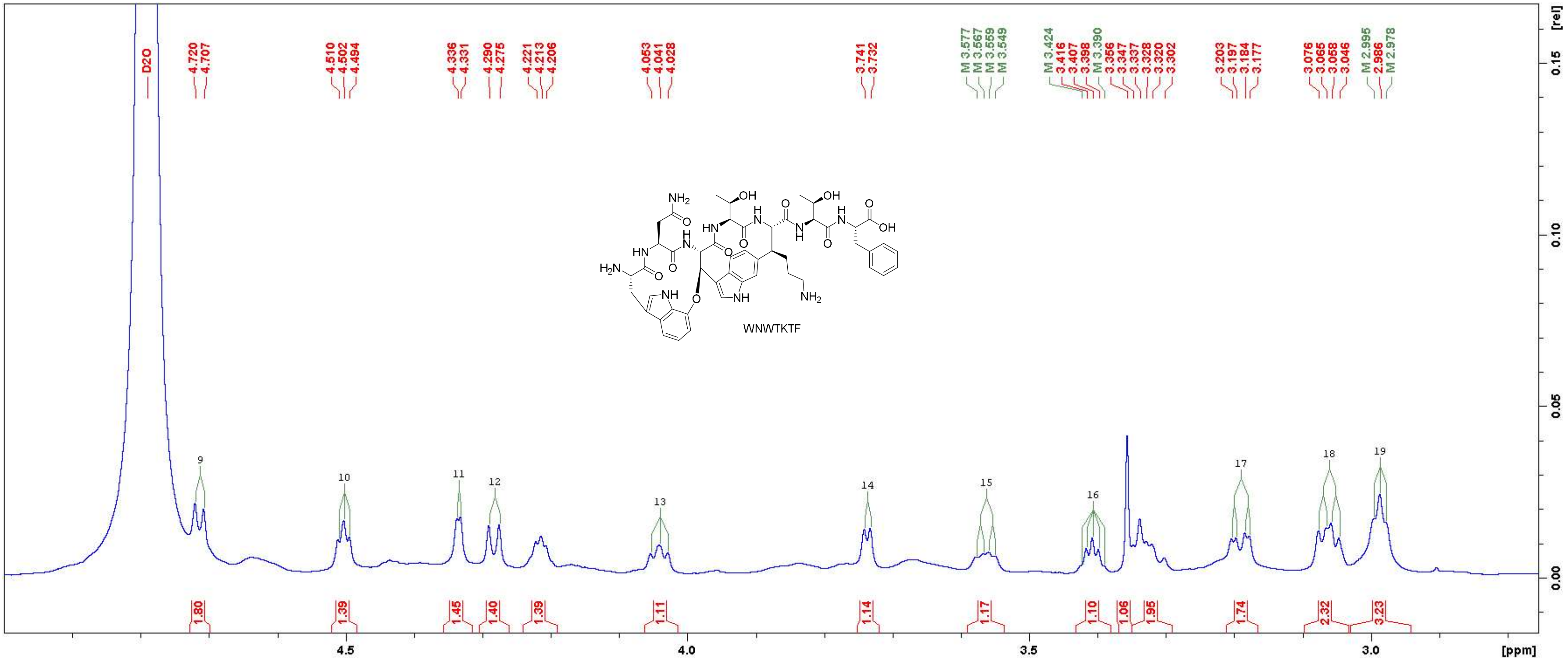

Figure S64: $^{1}$H NMR spectrum of WNWTKTF in $D_2O$ (700 MHz). Close-up in the region between 2.8 ppm and 4.9 ppm.

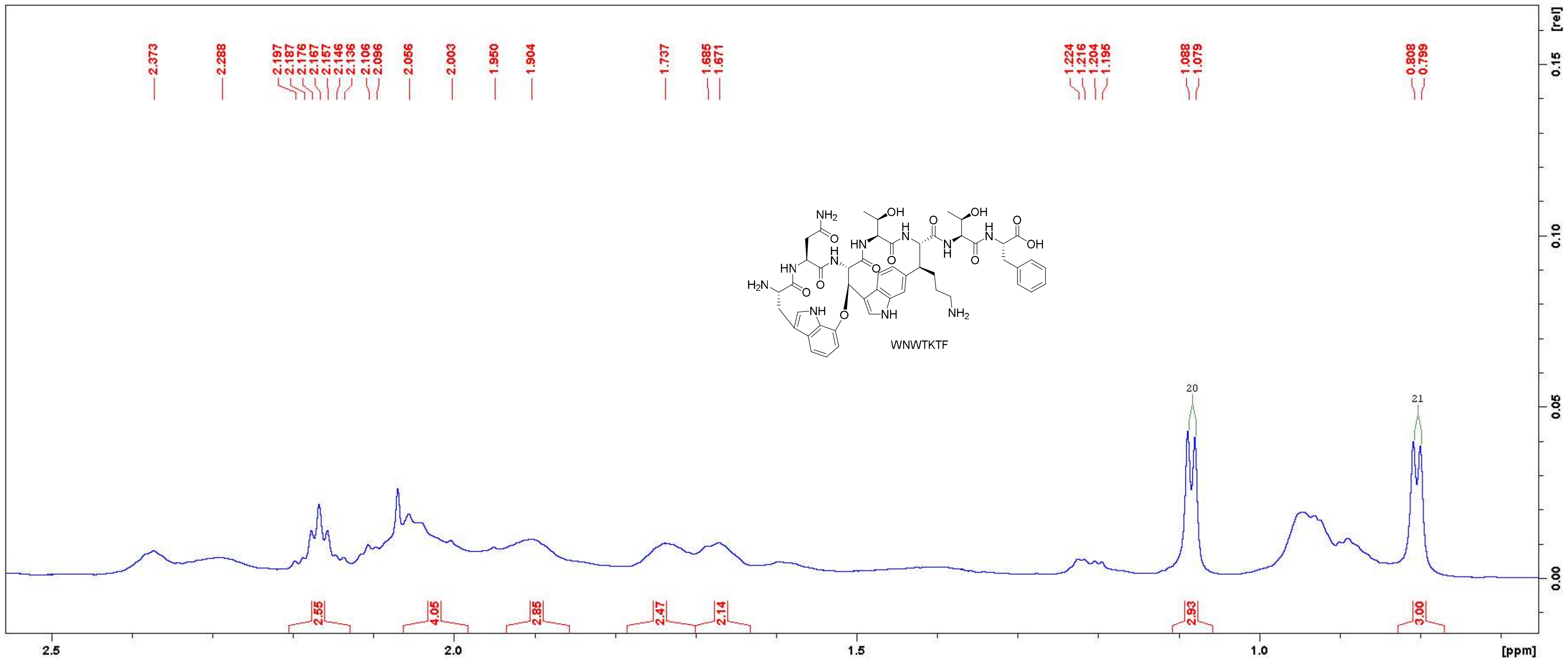

Figure S65: $^{1}H$ NMR spectrum of WNWTKTF in $D_2O$ (700 MHz). Close-up in the region between 0.7 ppm and 2.5 ppm.

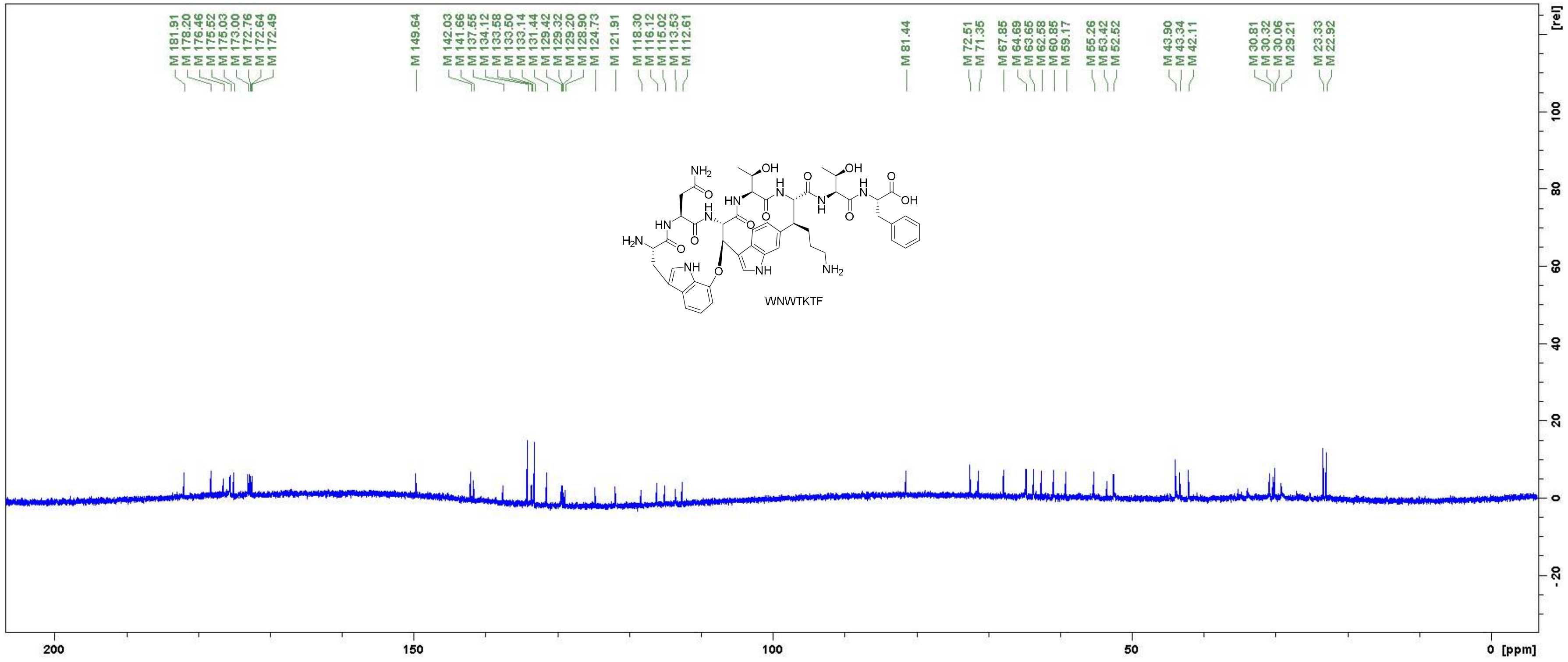

Figure S66: $^{13}C$ NMR spectrum of WNWTKTF in $D_2O$ (175 MHz).

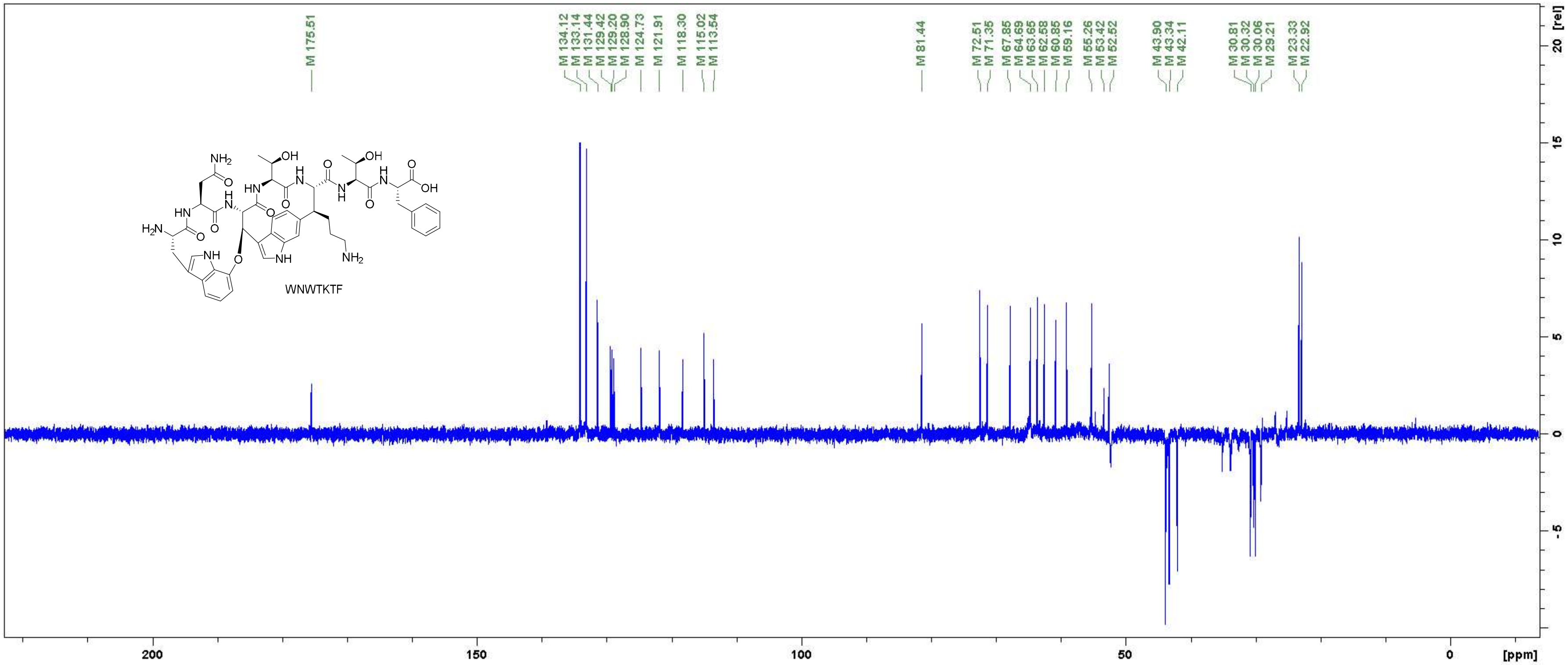

Figure S67: DEPT 135 spectrum of WNWTKTF in $D_2O$ (175 MHz).

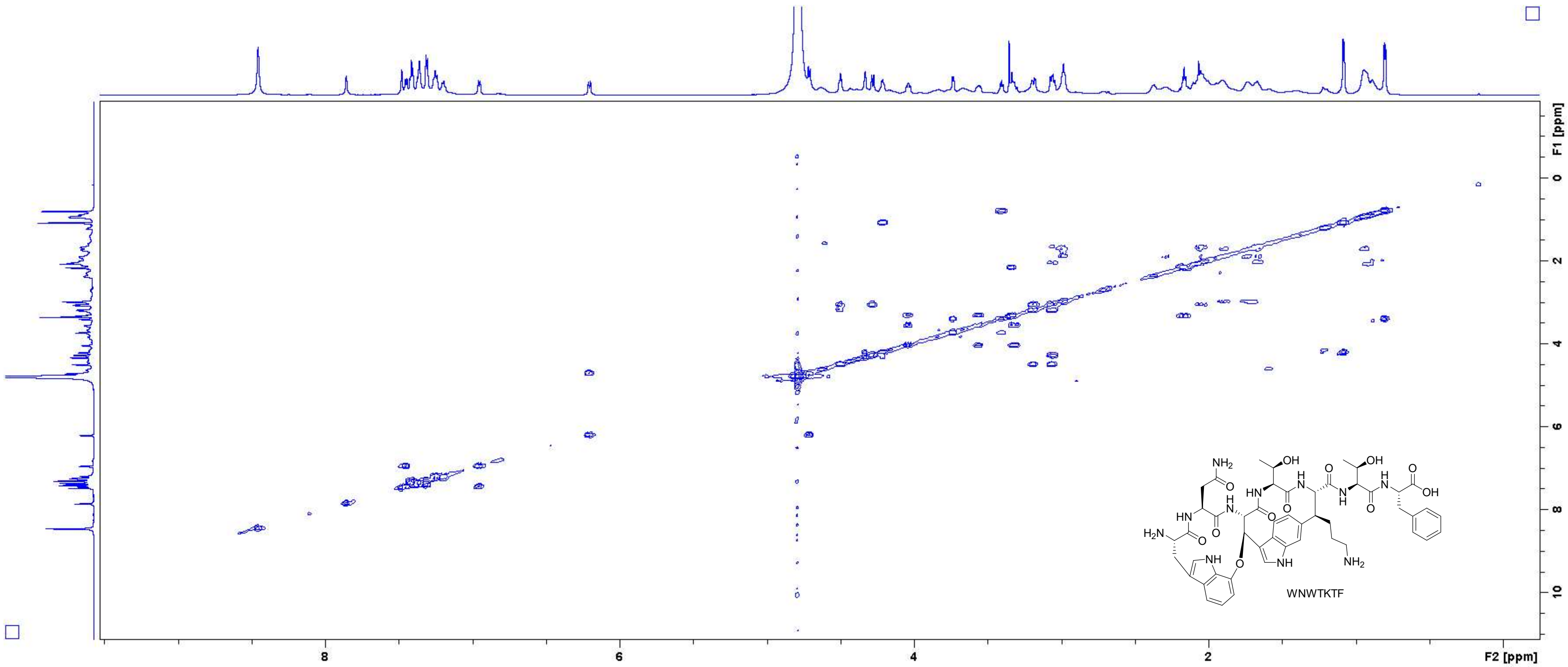

**Figure S68:** COSY spectrum of **WNWTKTF** in $D_2O$ (700 MHz).

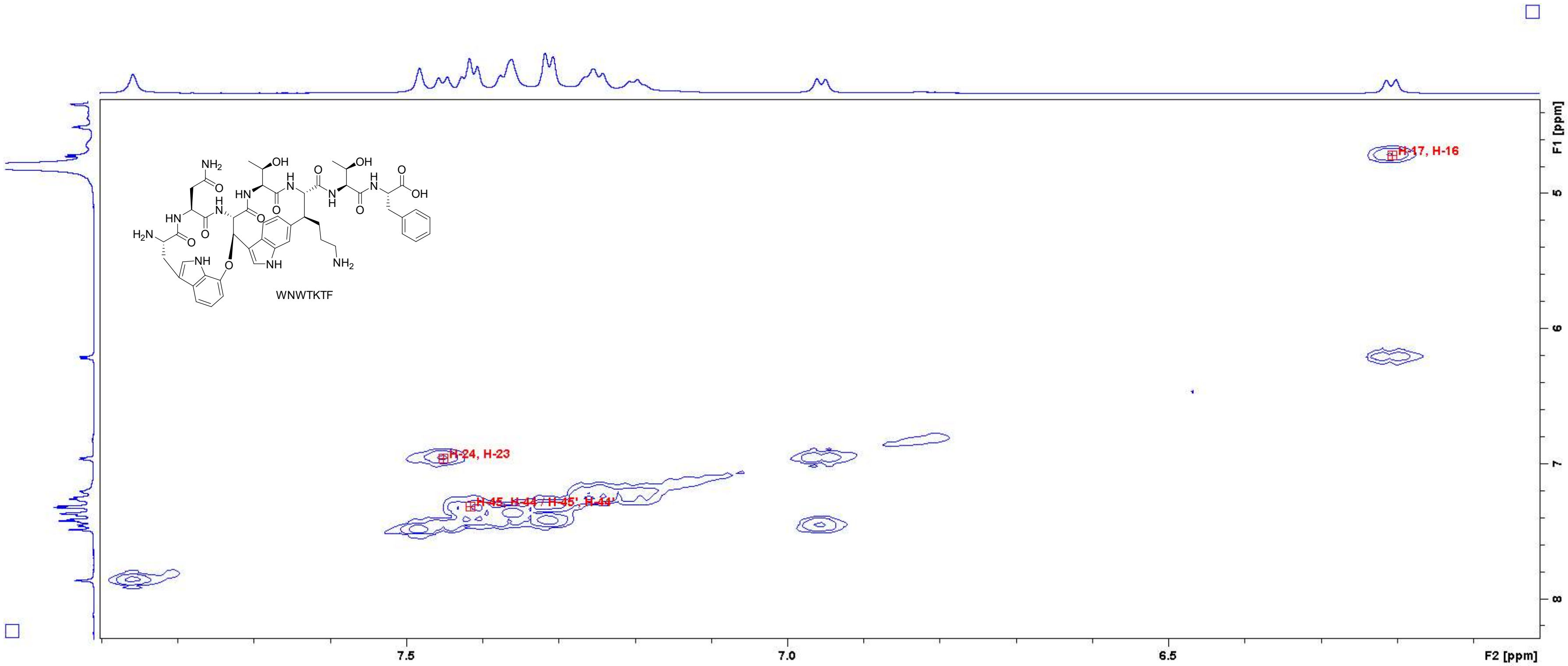

Figure S69: COSY spectrum of WNWTKTF in $D_2O$ (700 MHz). Close-up in the region of 6.1 – 7.9 ppm (F2 axis) and 4.4 – 8.2 ppm (F1 axis) with peak assignments. For atom numbering, cf. Figure S3.

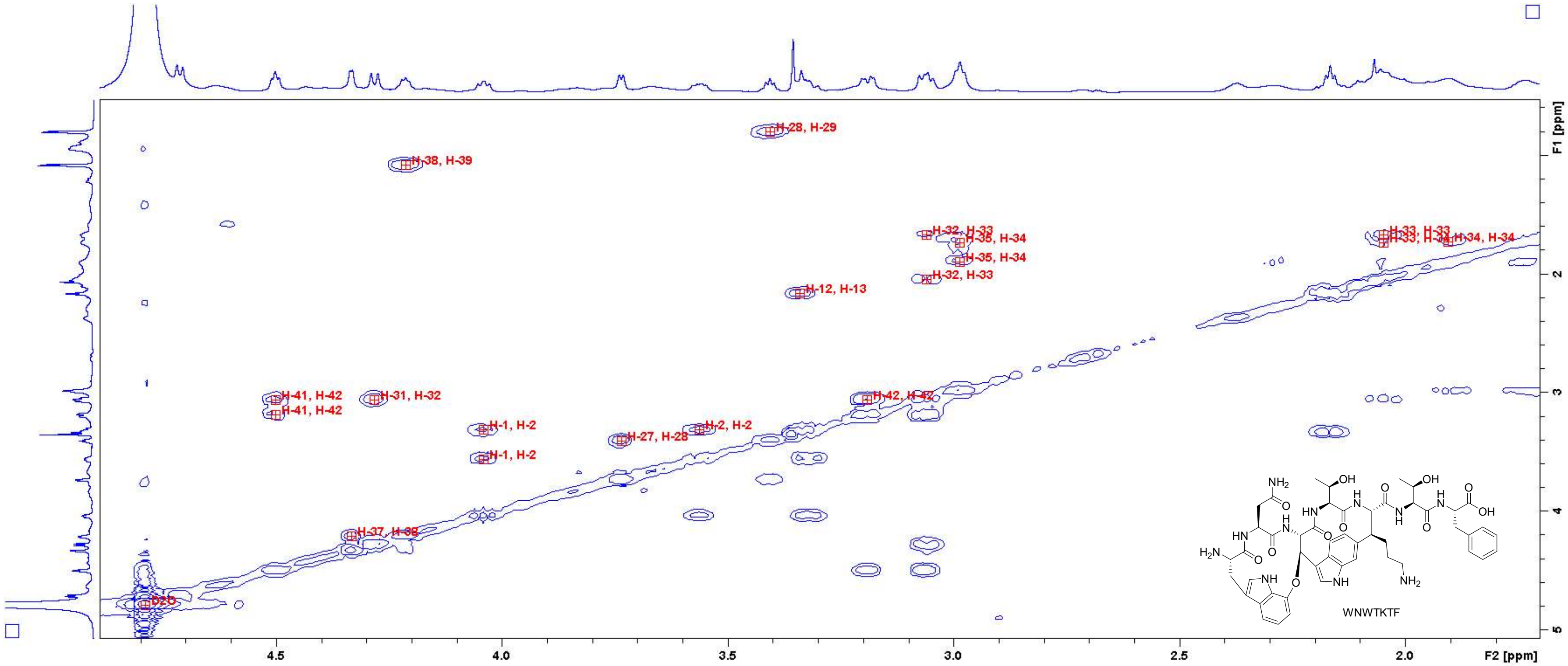

**Figure S70:** COSY spectrum of **WNWTKTF** in $D_2O$ (700 MHz). Close-up in the region of 1.8 – 4.8 ppm (F2 axis) and 0.6 – 5.0 ppm (F1 axis) with peak assignments. For atom numbering, cf. Figure S3.

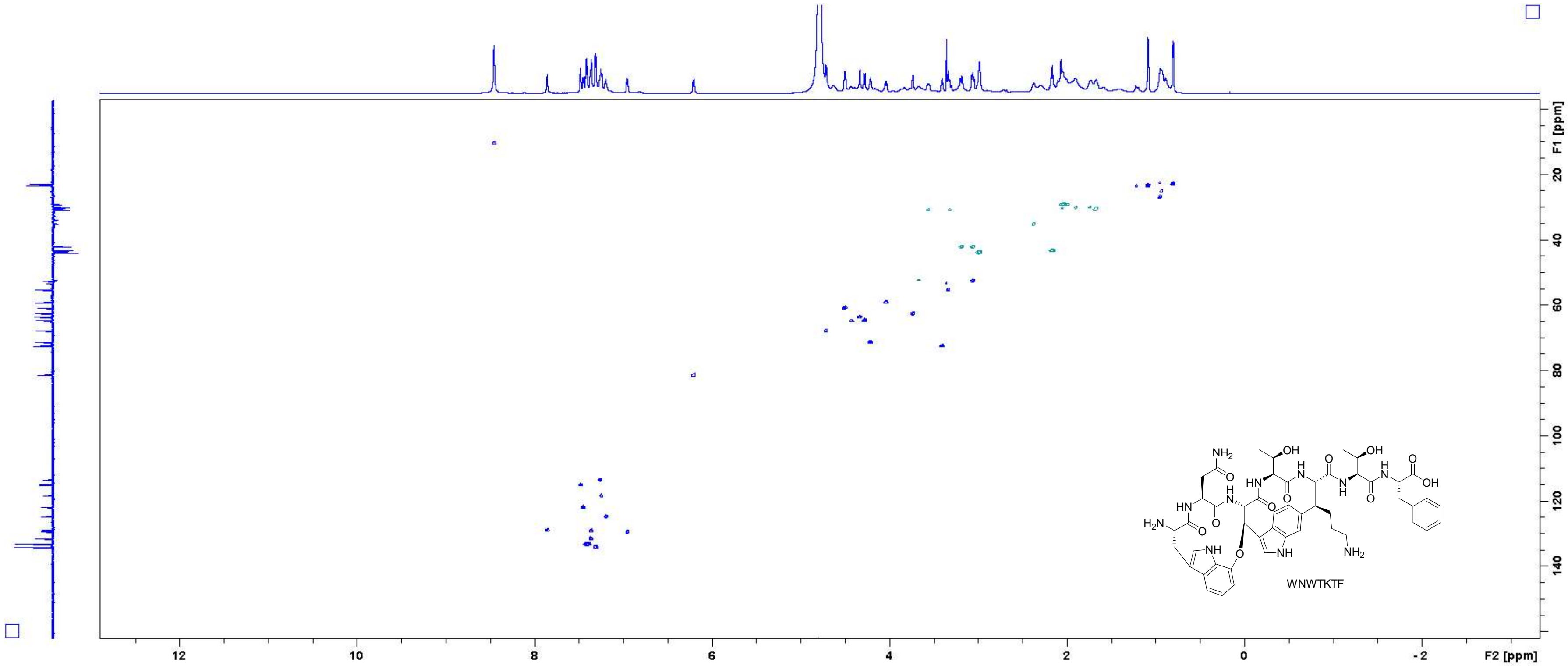

Figure S71: HSQC spectrum of WNWTKTF in $D_2O$ (700 MHz).

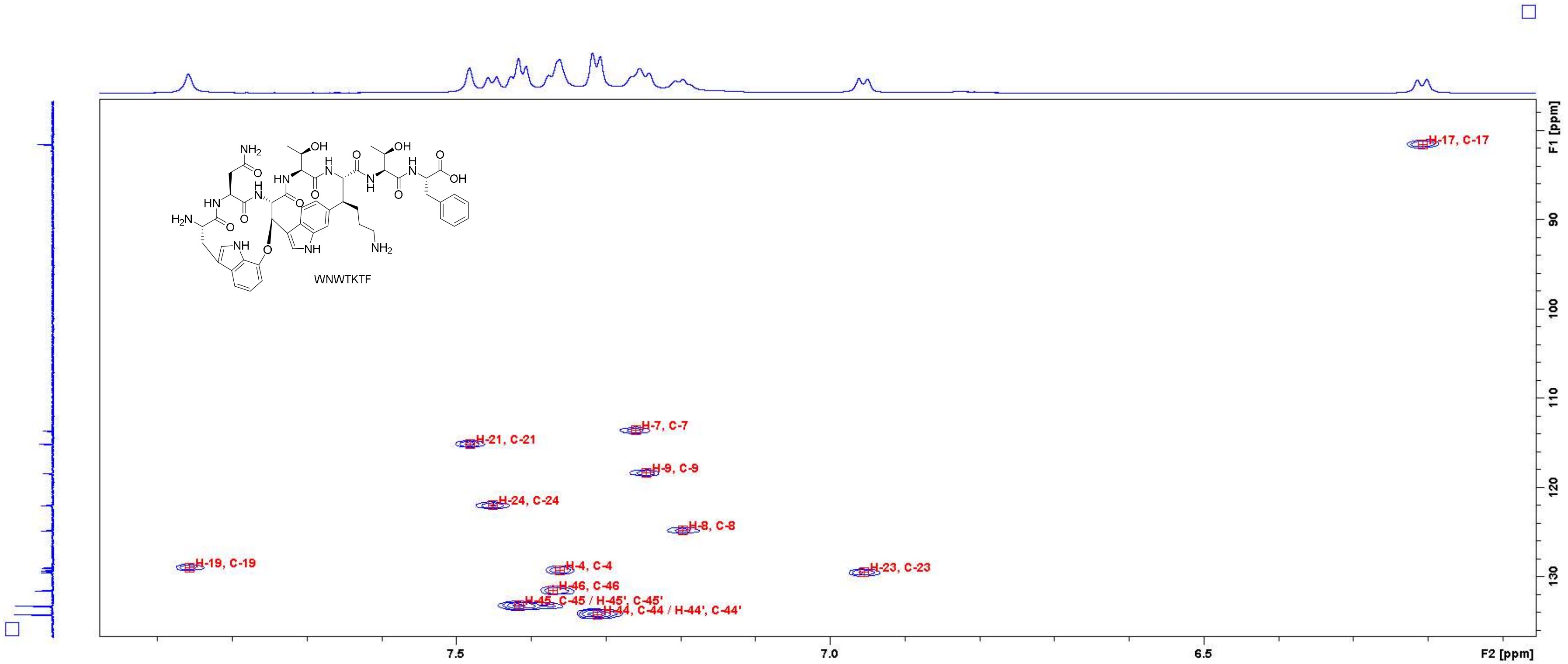

Figure S72: HSQC spectrum of WNWTKTF in $D_2O$ (700 MHz). Close-up in the region of 6.1 – 7.9 ppm (F2 axis) and 78 – 136 ppm (F1 axis) with peak assignments. For atom numbering, cf. Figure S3.

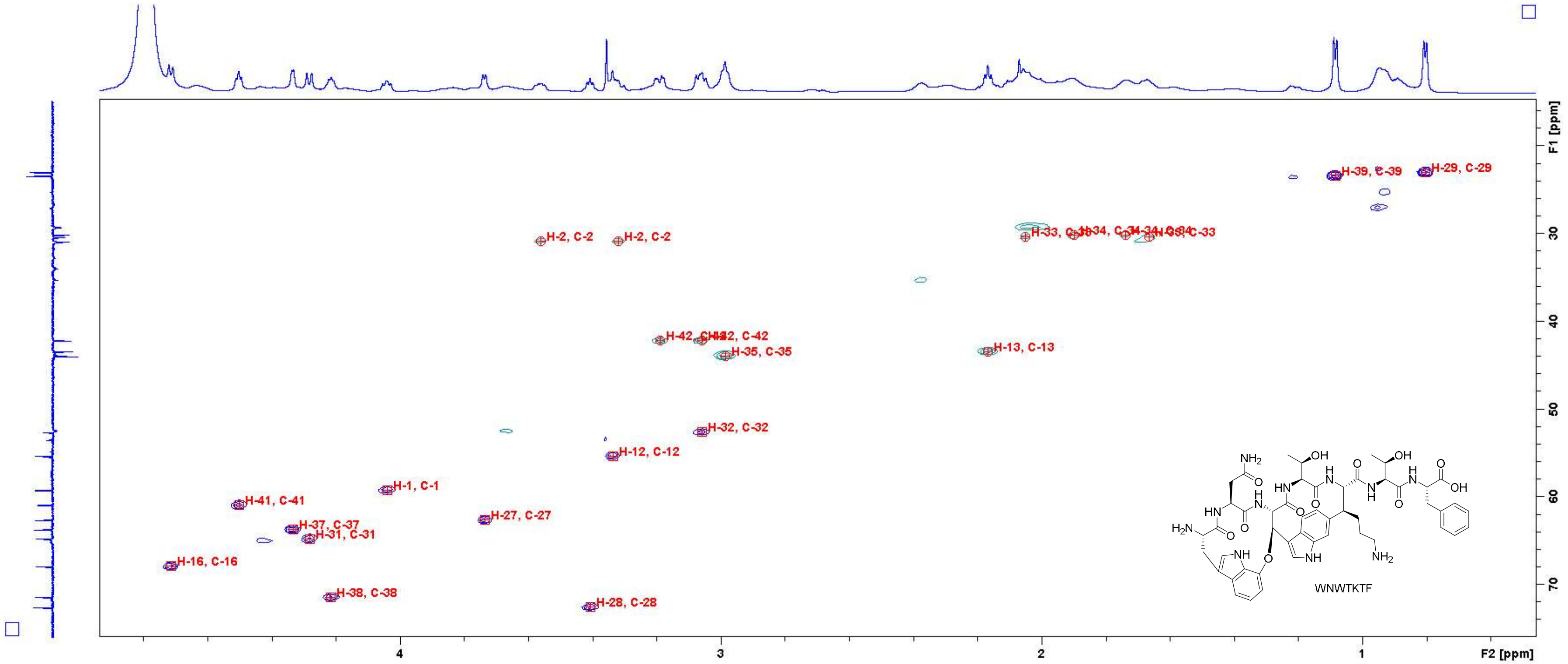

Figure S73: HSQC spectrum of WNWTKTF in $D_2O$ (700 MHz). Close-up in the region of 0.6 – 4.8 ppm (F2 axis) and 16 – 74 ppm (F1 axis) with peak assignments. For atom numbering, cf. Figure S3.

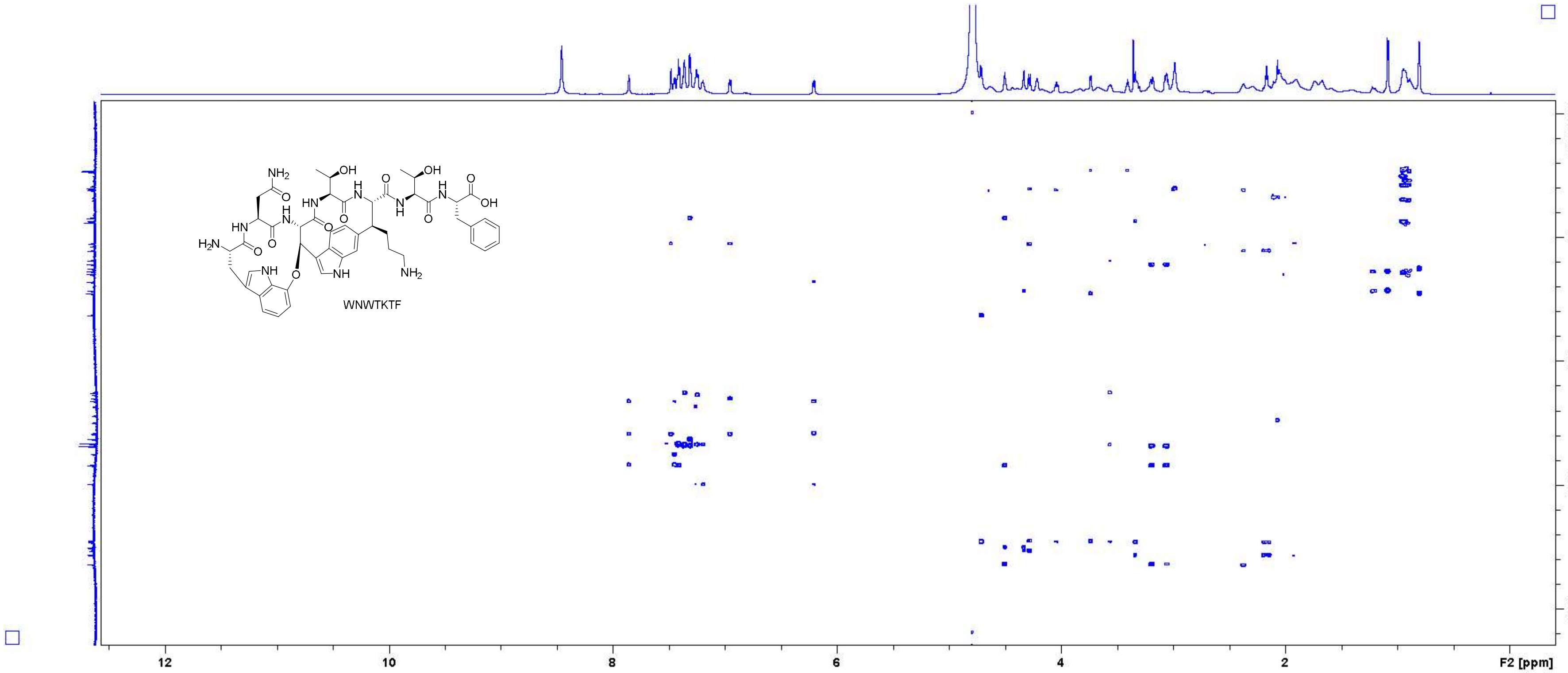

Figure S74: HMBC spectrum of **WNWTKTF** in $D_2O$ (700 MHz).

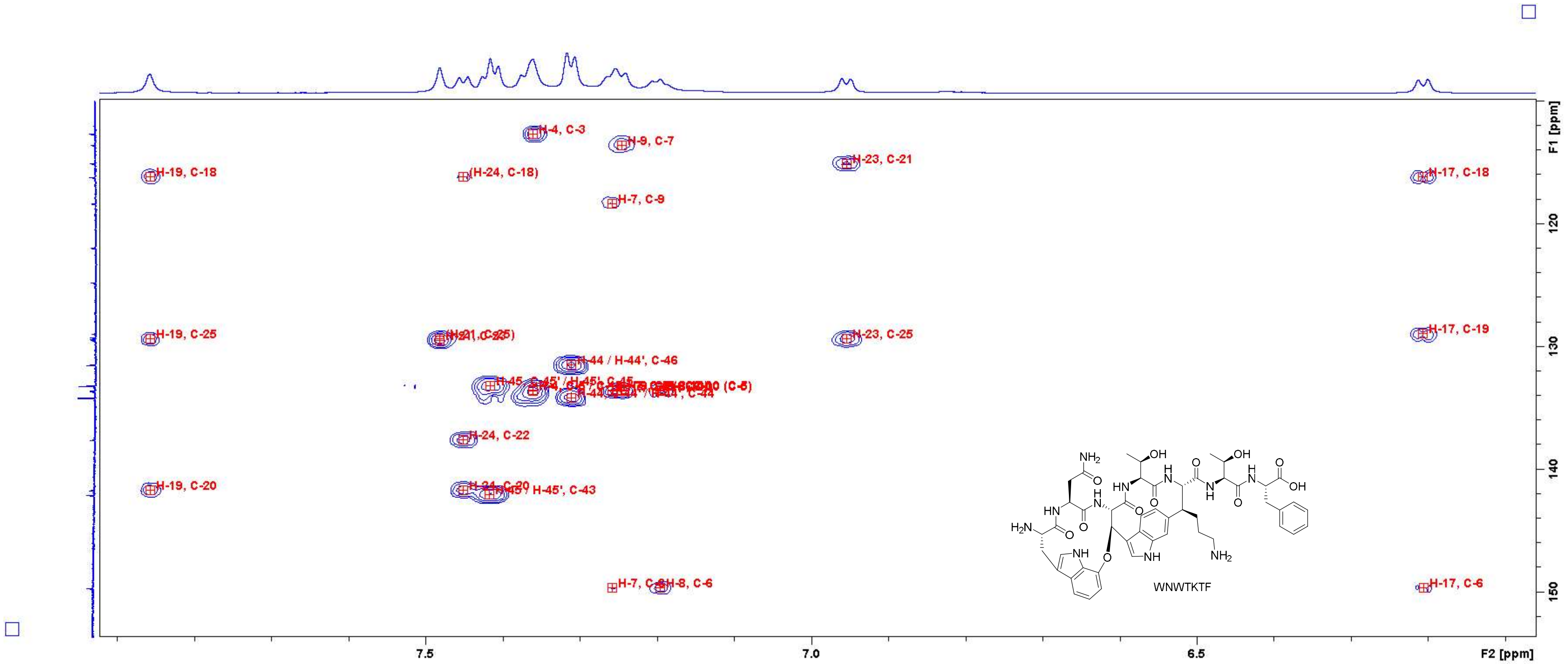

Figure S75: HMBC spectrum of WNWTKTF in $D_2O$ (700 MHz). Close-up in the region of 6.1 – 7.9 ppm (F2 axis) and 110 – 152 ppm (F1 axis) with peak assignments. For atom numbering, cf. Figure S3.

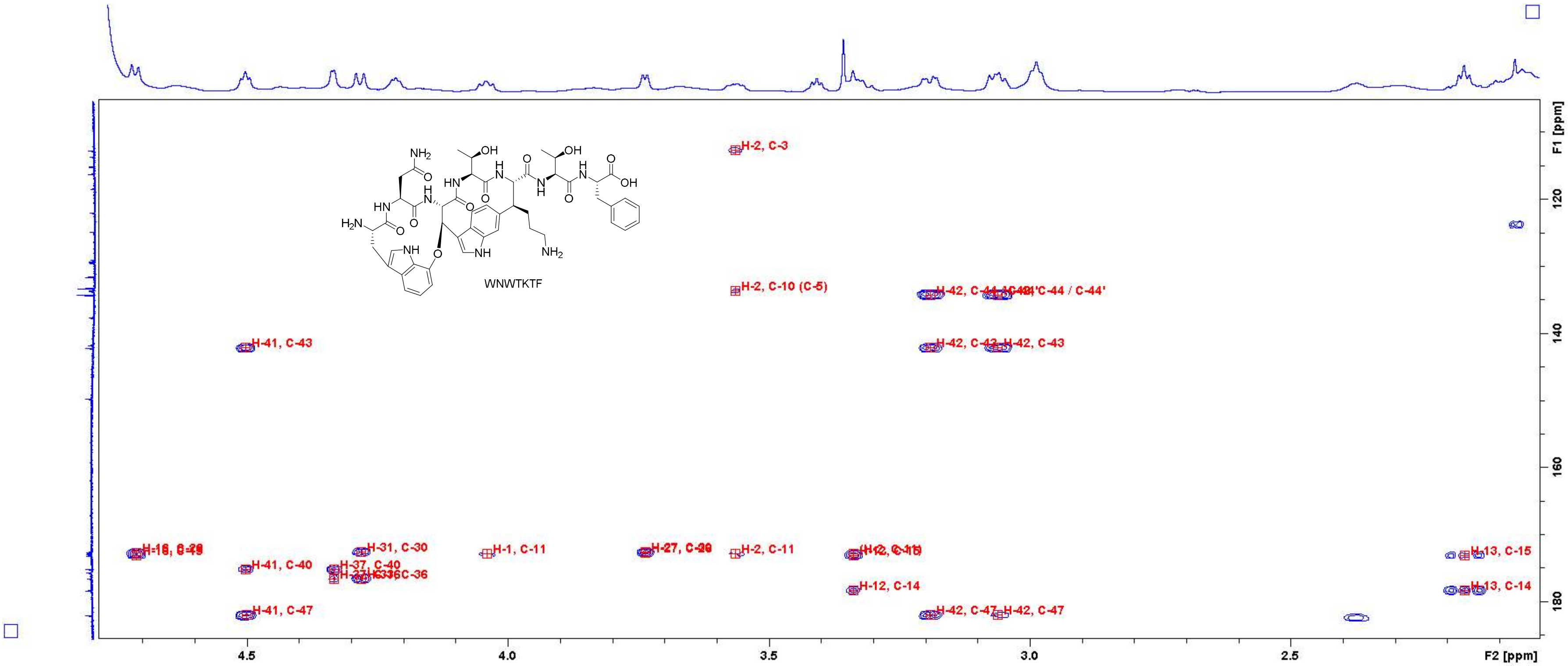

**Figure S76:** HMBC spectrum of **WNWTKTF** in $D_2O$ (700 MHz). Close-up in the region of 2.1 – 4.7 ppm (F2 axis) and 110 – 185 ppm (F1 axis) with peak assignments. For atom numbering, cf. Figure S3.

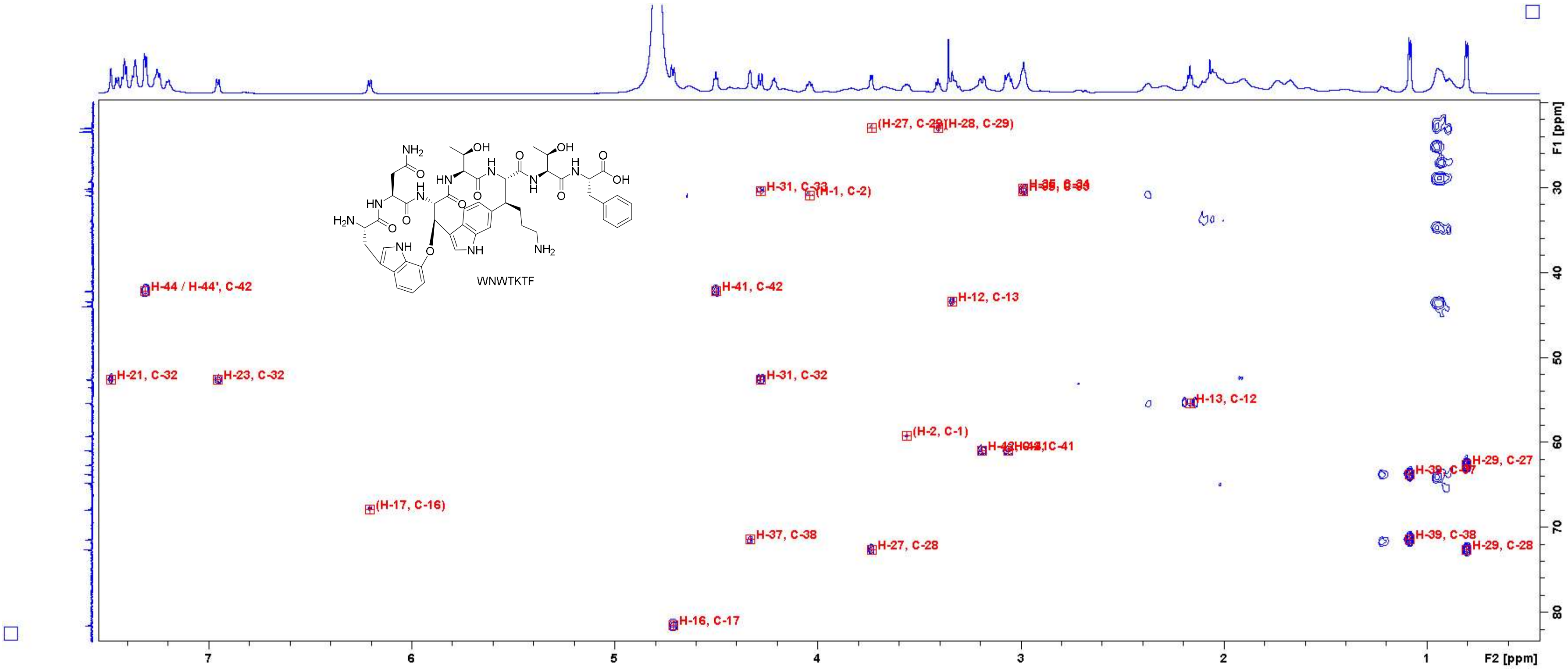

Figure S77: HMBC spectrum of **WNWTKTF** in $D_2O$ (700 MHz). Close-up in the region of 0.6 – 7.4 ppm (F2 axis) and 20 – 82 ppm (F1 axis) with peak assignments. For atom numbering, cf. Figure S3.

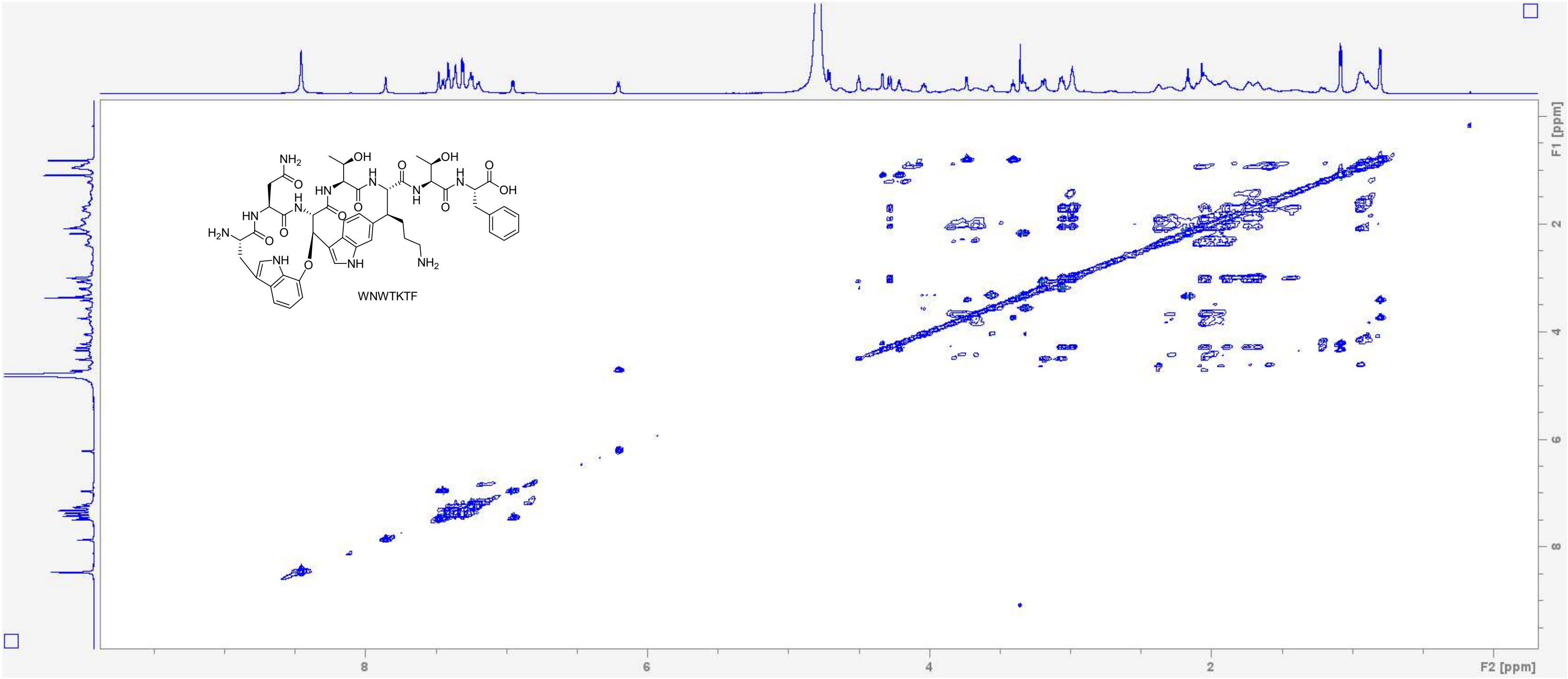

Figure S78: TOCSY spectrum of WNWTKTF in $D_2O$ (700 MHz, measured with $H_2O$ suppression).

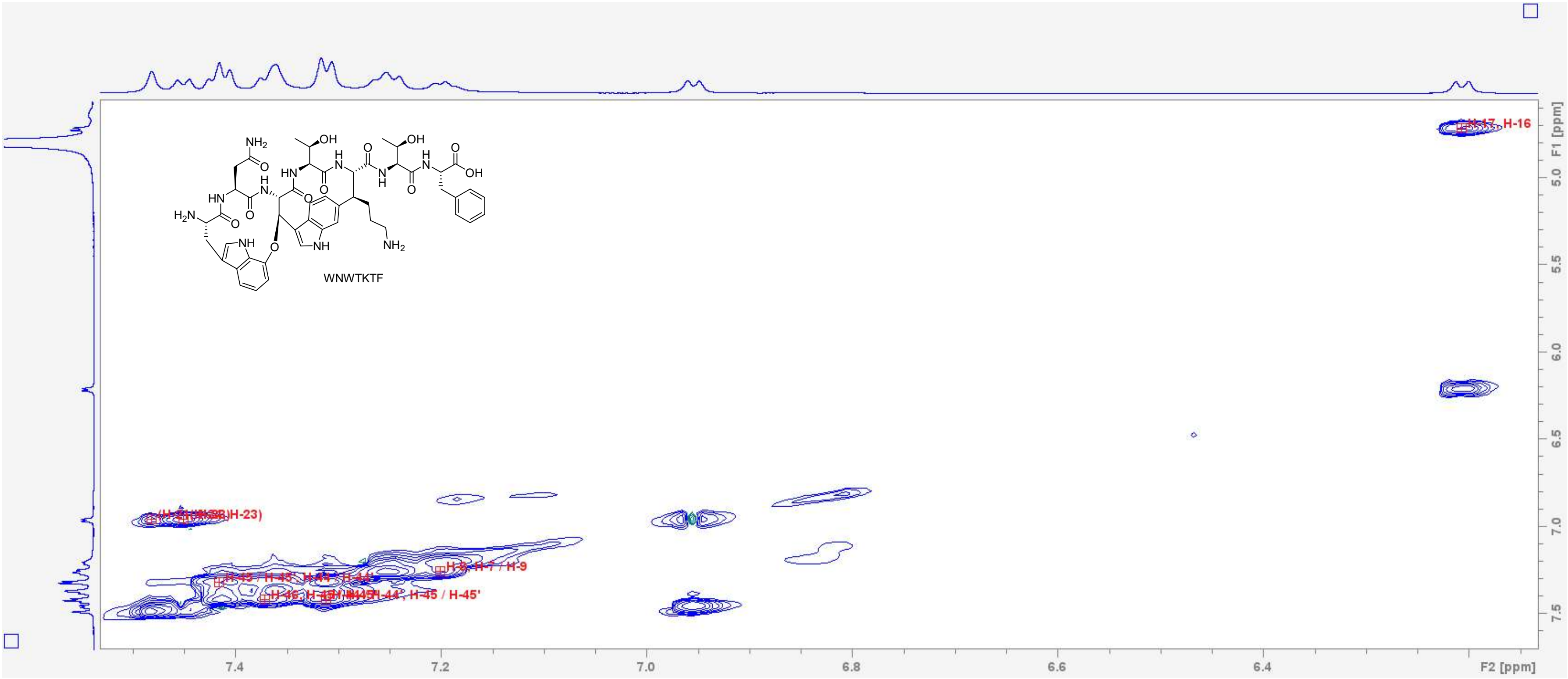

Figure S79: TOCSY spectrum of WNWTKTF in $D_2O$ (700 MHz, measured with $H_2O$ suppression). Close-up in the region of 6.15 – 7.50 ppm (F2 axis) and 4.6 – 7.6 ppm (F1 axis) with peak assignments. For atom numbering, cf. Figure S3.

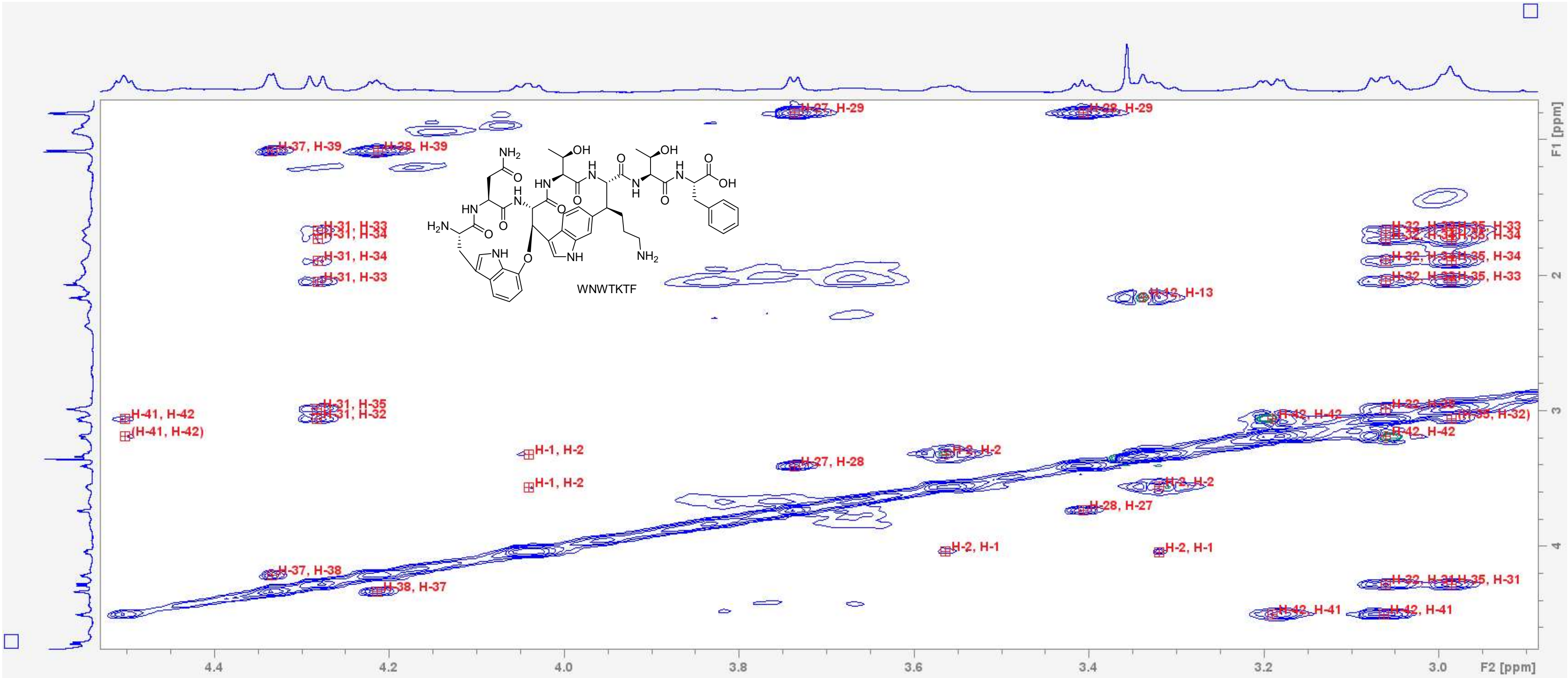

Figure S80: TOCSY spectrum of WNWTKTF in D2O (700 MHz, measured with H2O suppression). Close-up in the region of 2.9 – 4.5 ppm (F2 axis) and 0.8 – 4.6 ppm (F1 axis) with peak assignments. For atom numbering, cf. Figure S3.

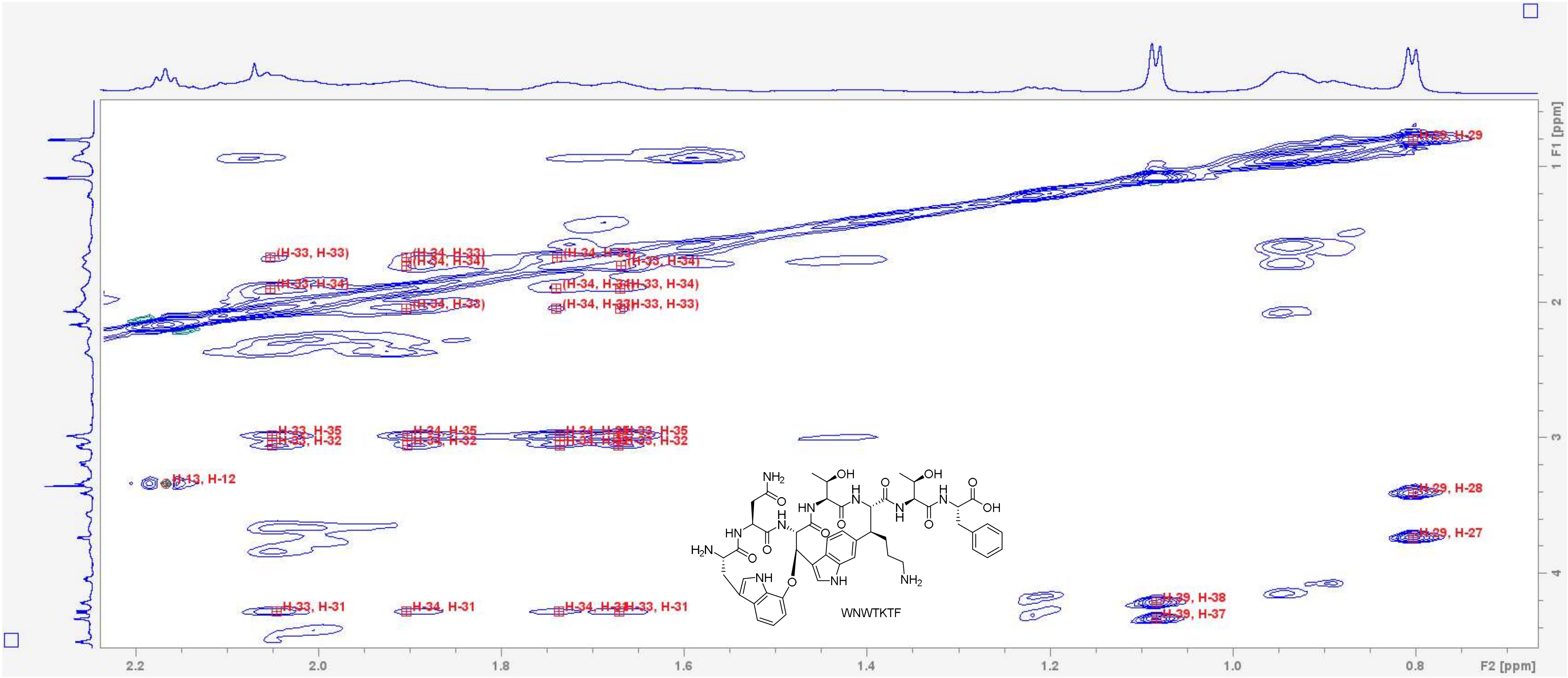

Figure S81: TOCSY spectrum of **WNWTKTF** in $D_2O$ (700 MHz, measured with $H_2O$ suppression). Close-up in the region of 0.7 – 2.2 ppm (F2 axis) and 0.6 – 4.4 ppm (F1 axis) with peak assignments. For atom numbering, cf. Figure S3.

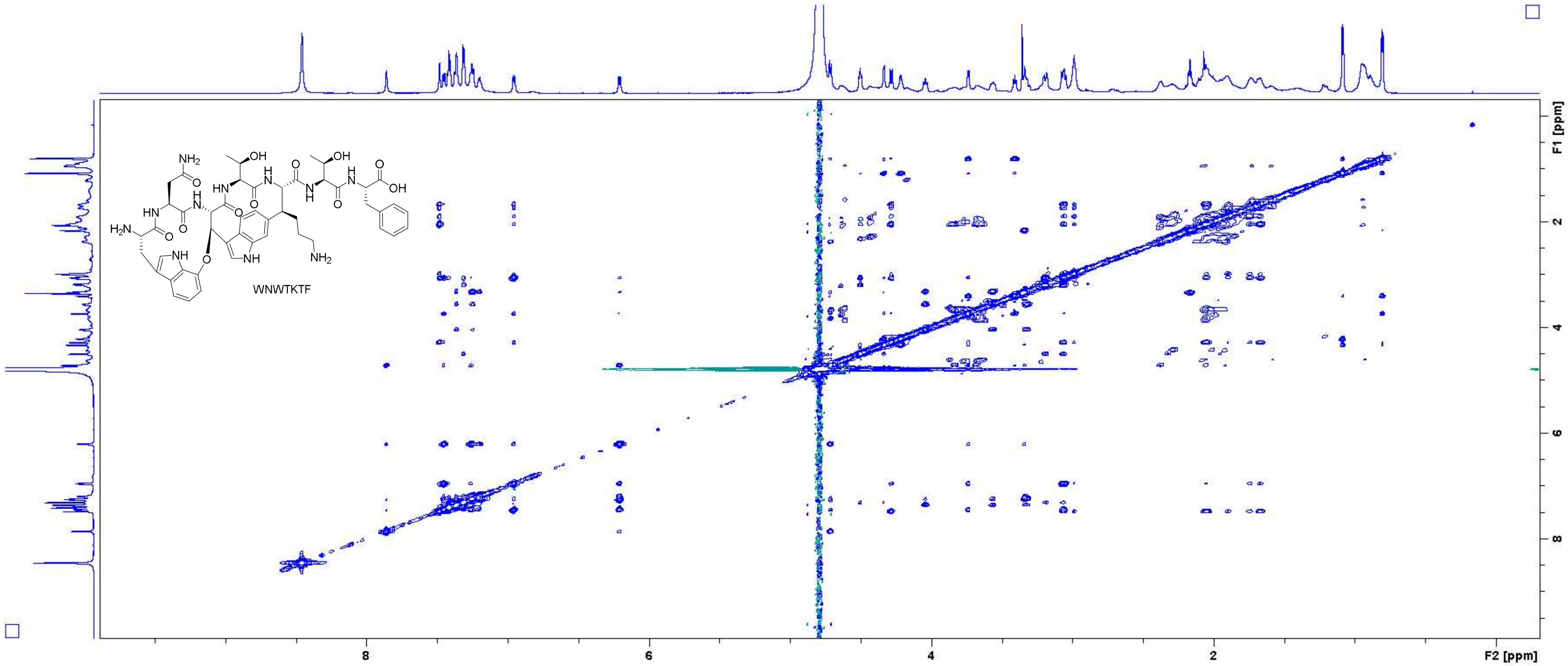

Figure S82: NOESY spectrum of **WNWTKTF** in $D_2O$ (700 MHz).

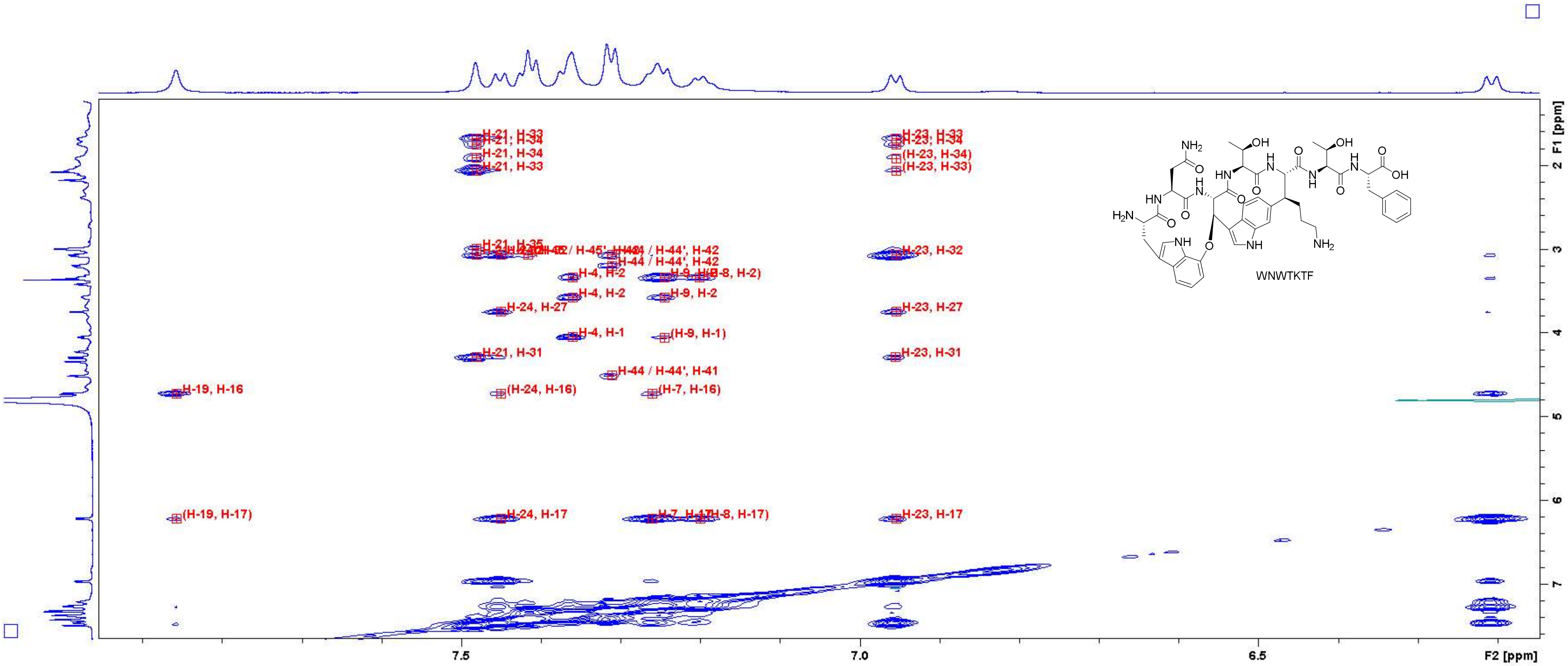

Figure S83: NOESY spectrum of **WNWTKTF** in $D_2O$ (700 MHz). Close-up in the region of 6.2 – 7.9 ppm (F2 axis) and 1.4 – 7.6 ppm (F1 axis) with peak assignments. For atom numbering, cf. Figure S3.

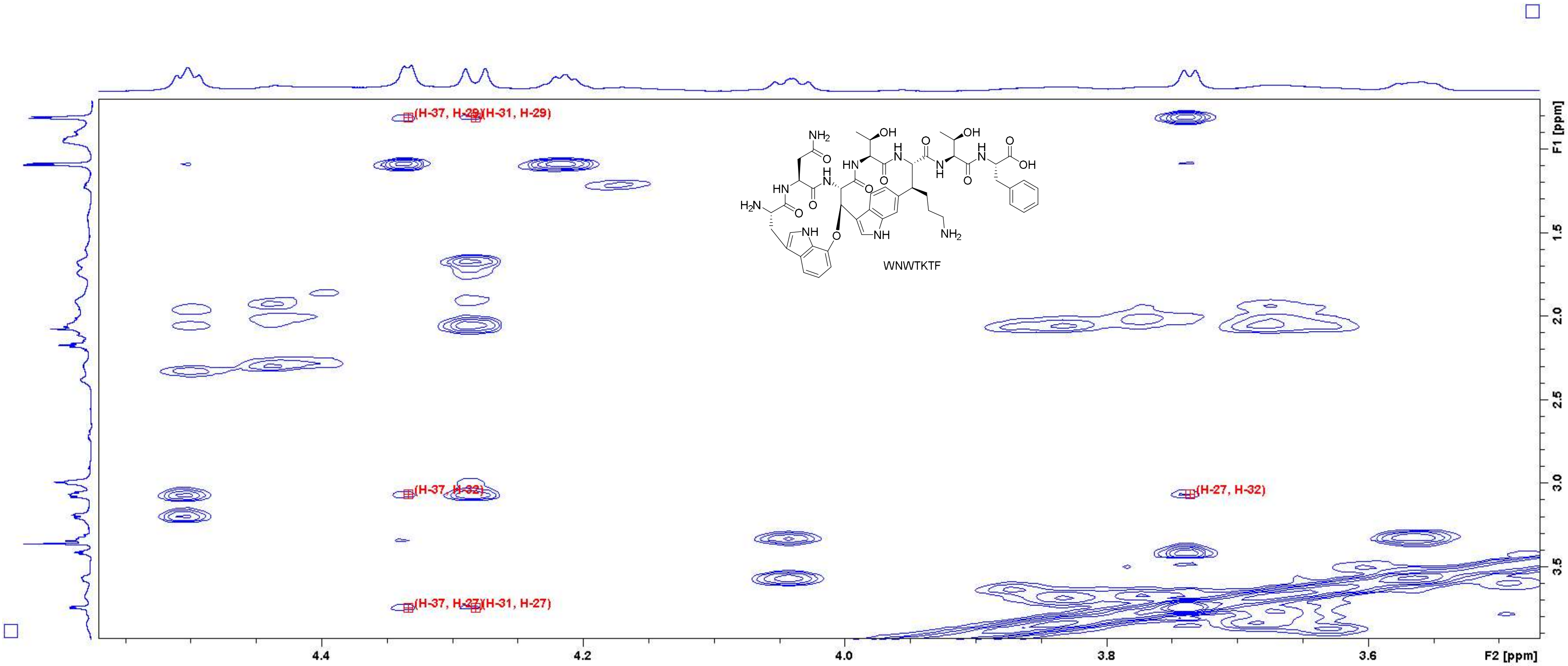

Figure S84: NOESY spectrum of WNWTKTF in $D_2O$ (700 MHz). Close-up in the region of 3.50 – 4.55 ppm (F2 axis) and 0.8 – 3.9 ppm (F1 axis) with peak assignments. For atom numbering, cf. Figure S3.

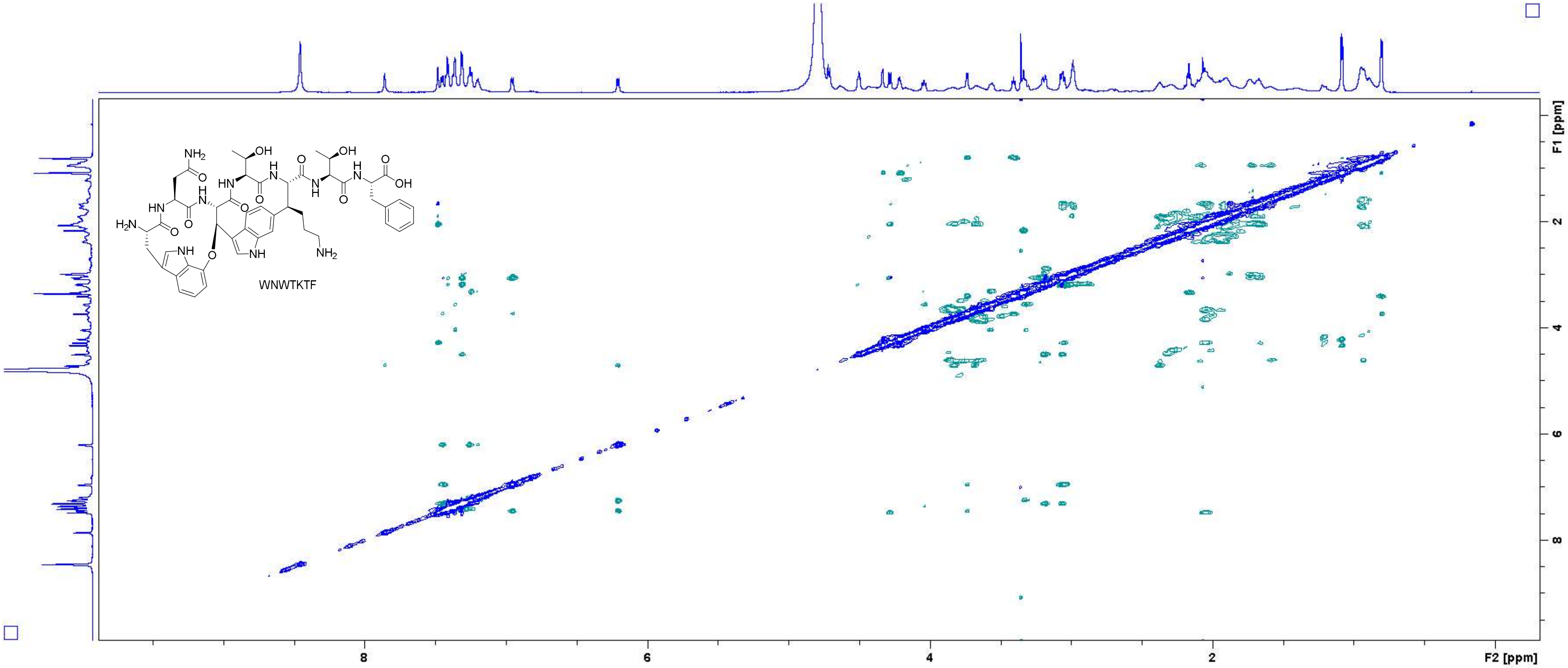

Figure S85: ROESY spectrum of WNWTKTF in $D_2O$ (700 MHz, measured with $H_2O$ suppression).

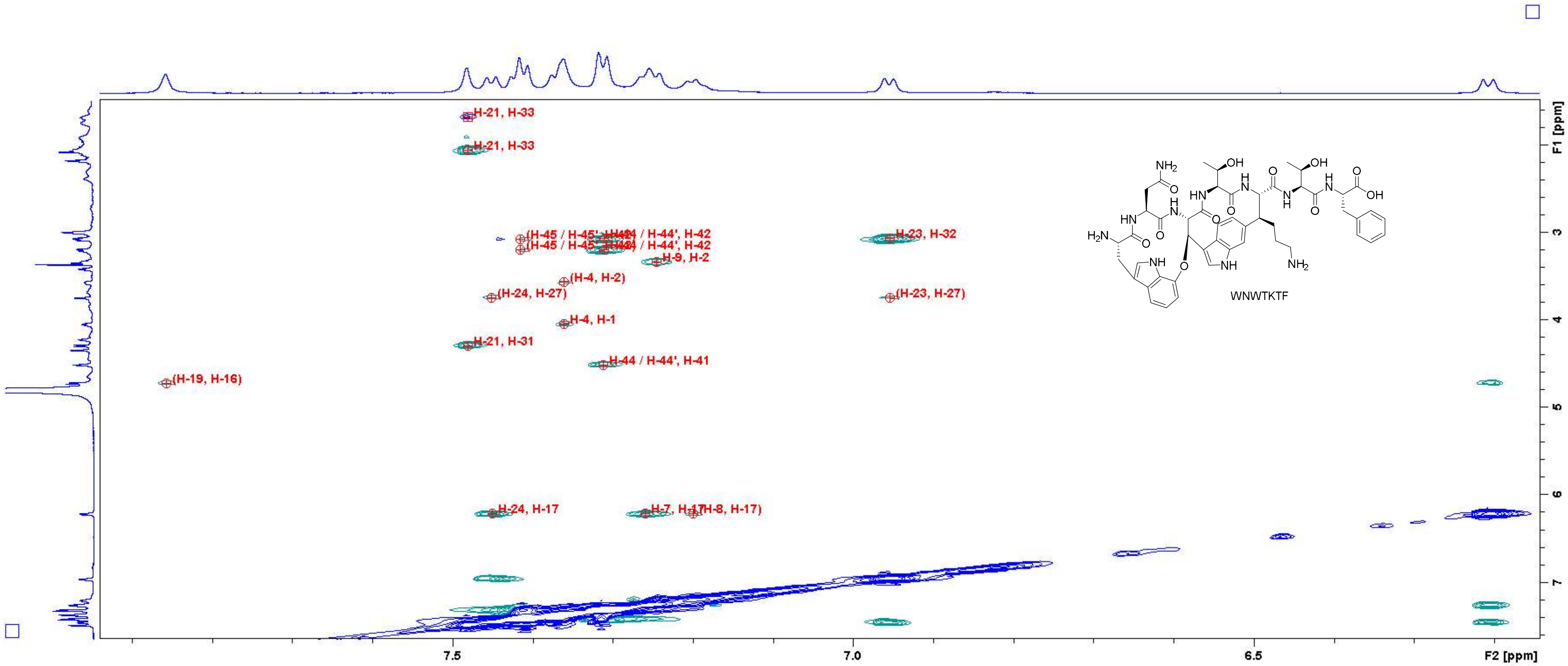

Figure S86: ROESY spectrum of **WNWTKTF** in $D_2O$ (700 MHz, measured with $H_2O$ suppression). Close-up in the region of 6.2 – 7.9 ppm (F2 axis) and 1.6 – 7.6 ppm (F1 axis) with peak assignments. For atom numbering, cf. Figure S3.

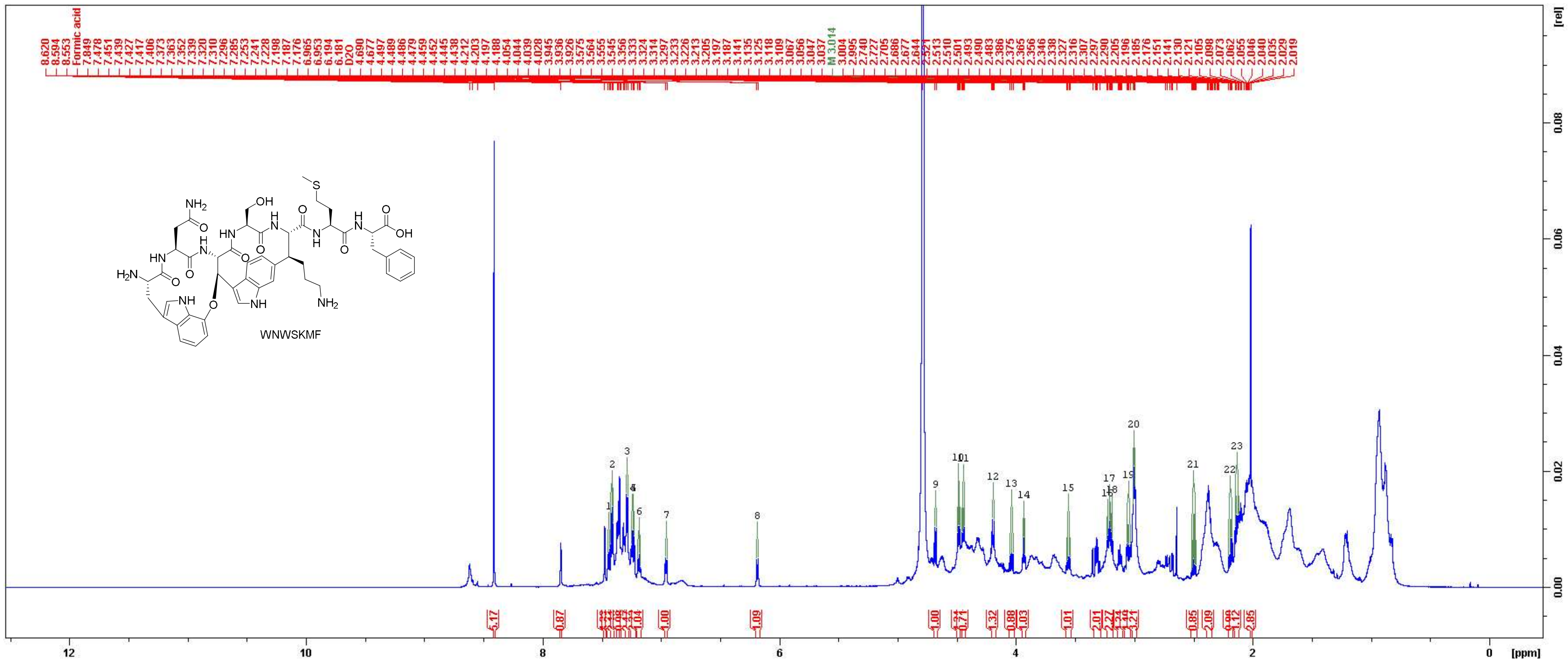

Figure S87: $^{1}$H NMR spectrum of **WNWSKMF** in $D_2O$ (700 MHz).

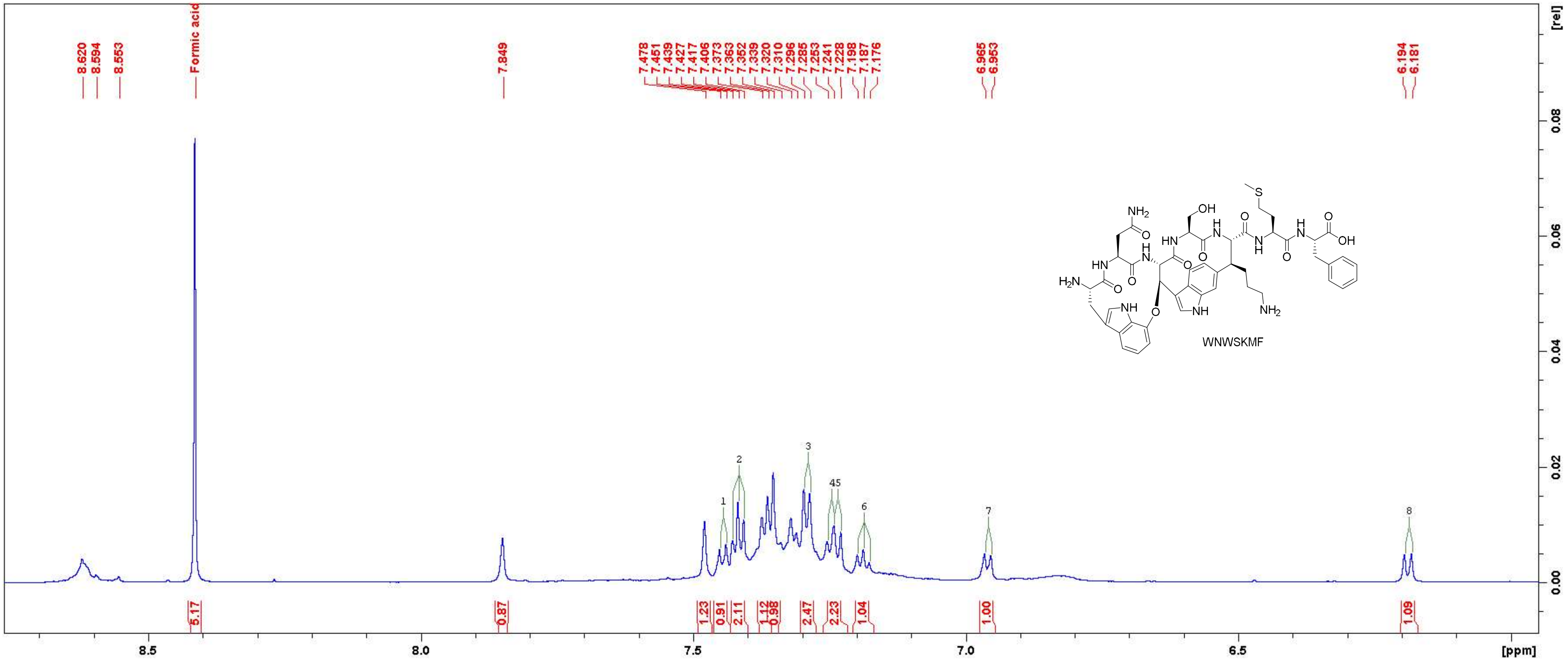

Figure S88: $^{1}$H NMR spectrum of WNWSKMF in $D_2O$ (700 MHz). Close-up in the region between 6.0 ppm and 8.7 ppm.

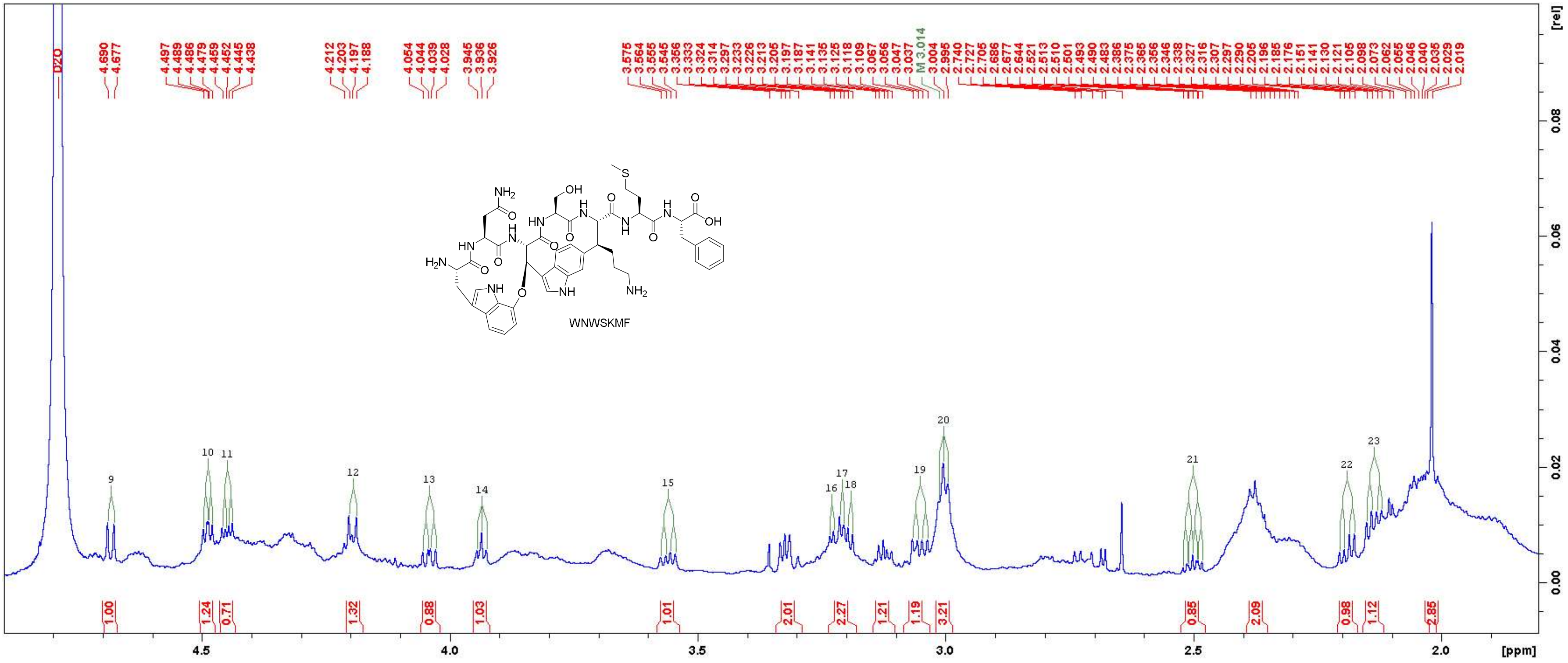

Figure S89: $^{1}$H NMR spectrum of WNWSKMF in $D_2O$ (700 MHz). Close-up in the region between 1.9 ppm and 4.8 ppm.

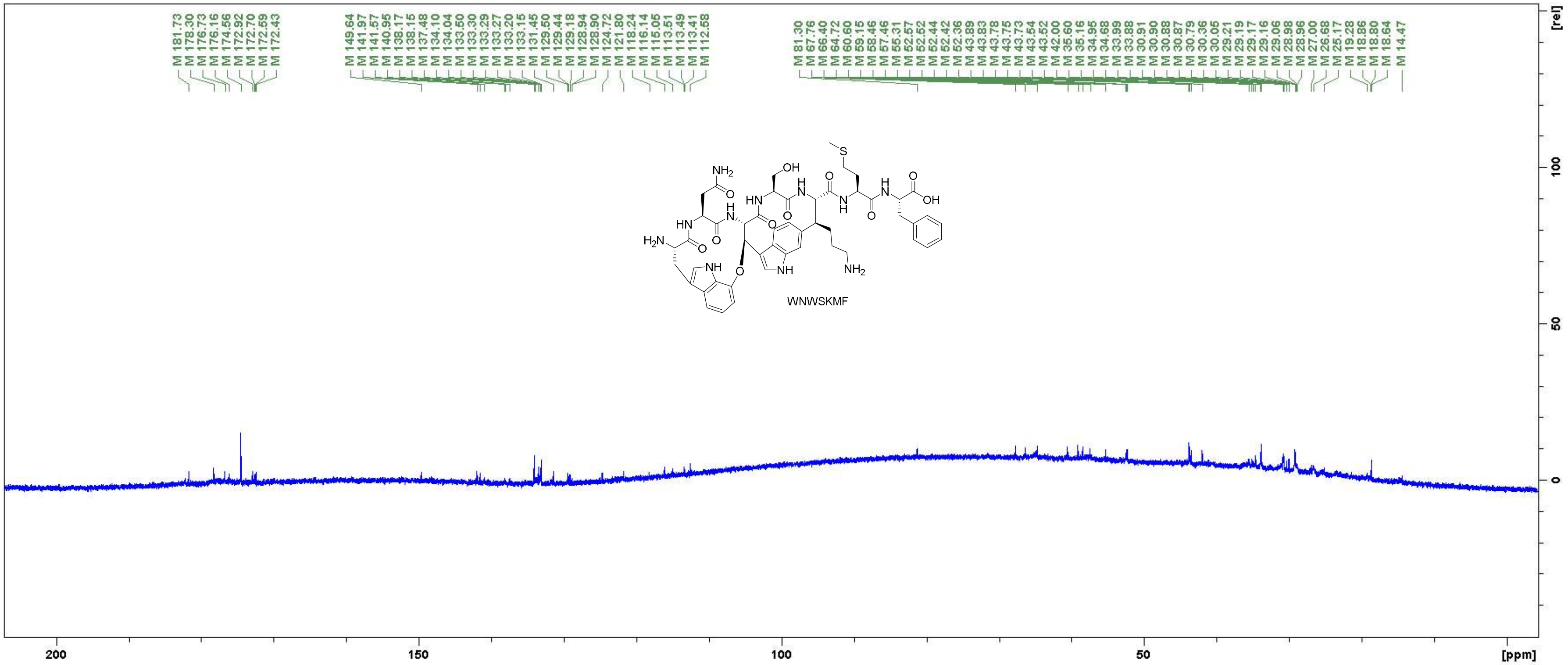

Figure S90: $^{13}C$ NMR spectrum of **WNWSKMF** in $D_2O$ (175 MHz).

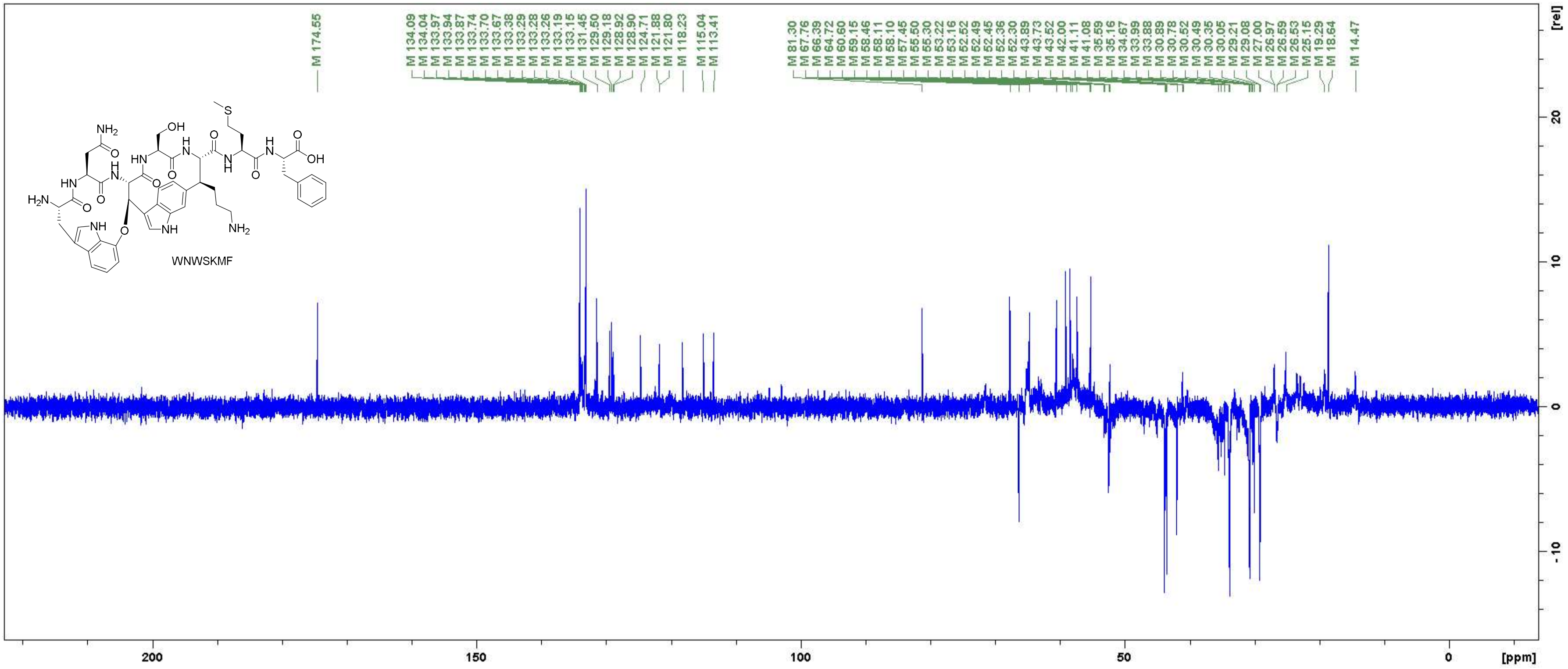

Figure S91: DEPT 135 spectrum of WNWSKMF in $D_2O$ (175 MHz).

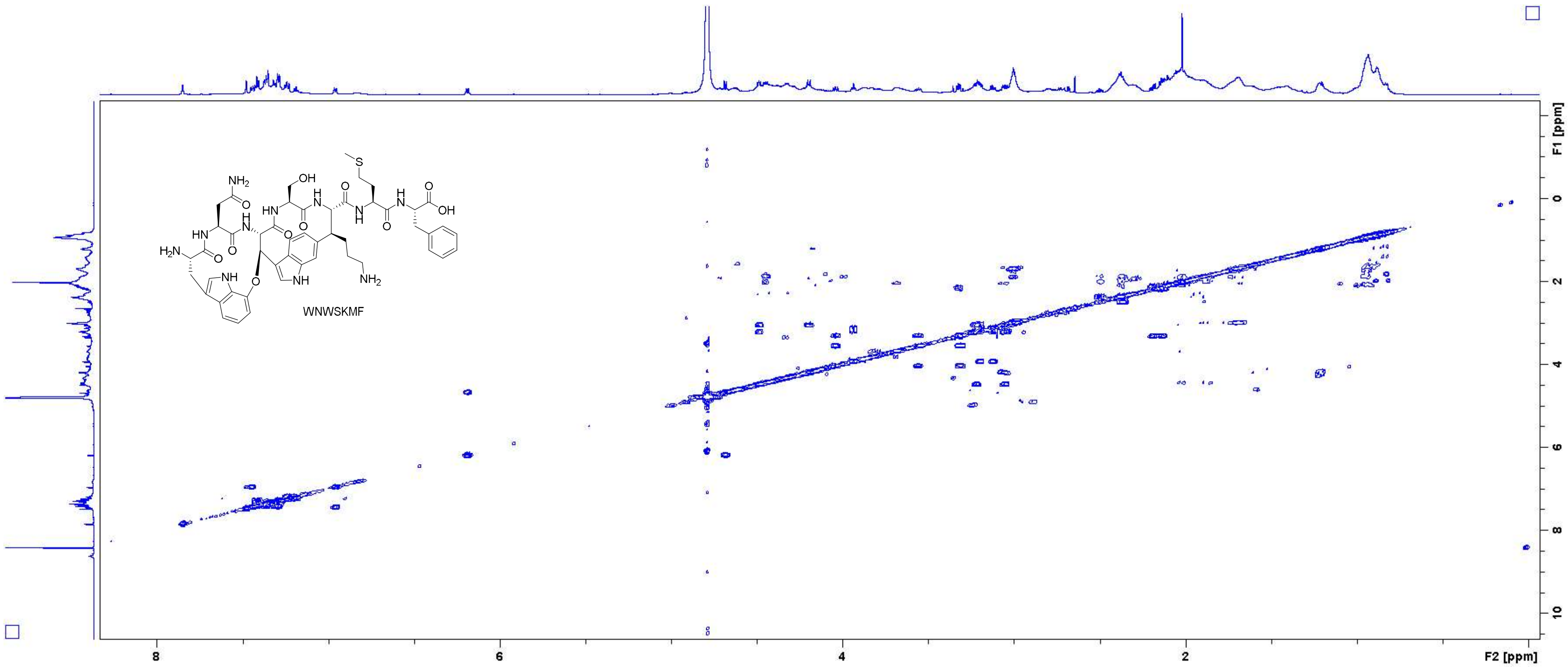

Figure S92: COSY spectrum of WNWSKMF in $D_2O$ (700 MHz).

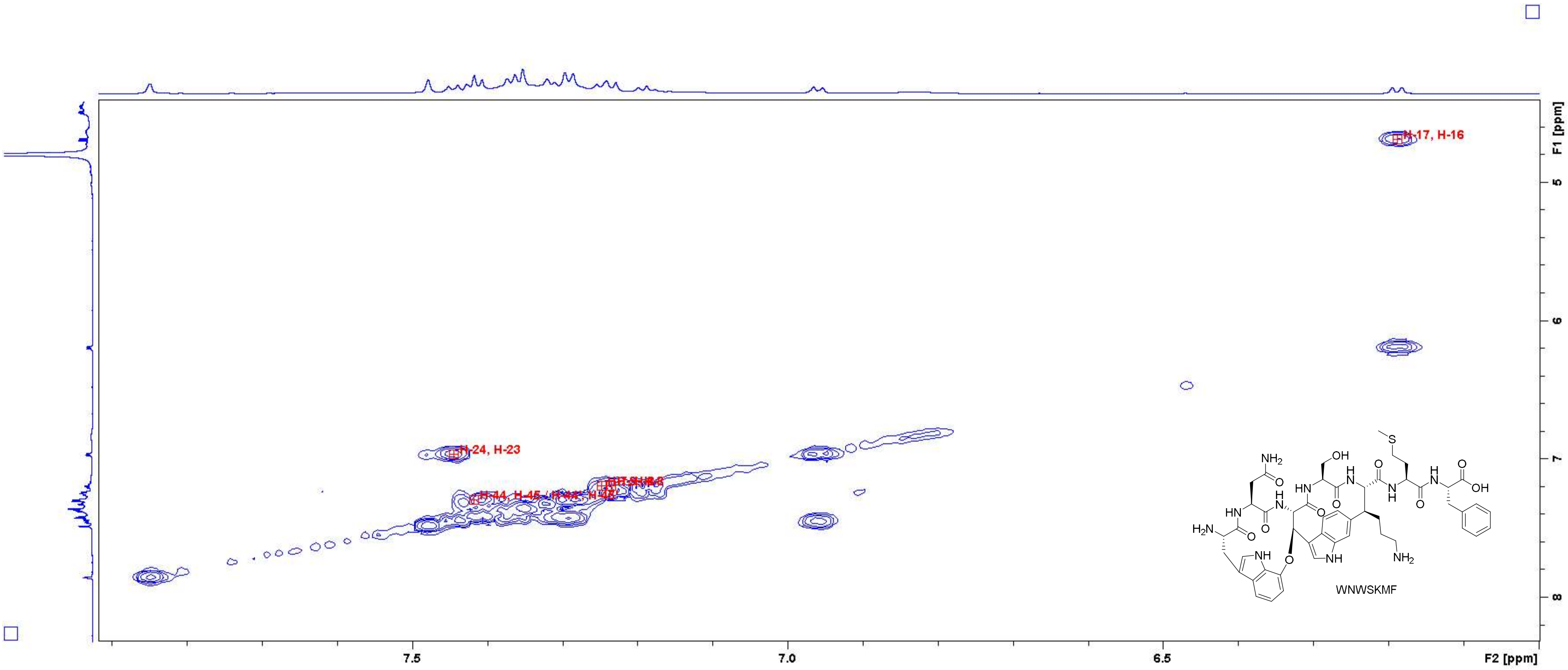

**Figure S93:** COSY spectrum of **WNWSKMF** in $D_2O$ (700 MHz). Close-up in the region of 6.0 – 7.9 ppm (F2 axis) and 4.6 – 8.2 ppm (F1 axis) with peak assignments. For atom numbering, cf. Figure S3.

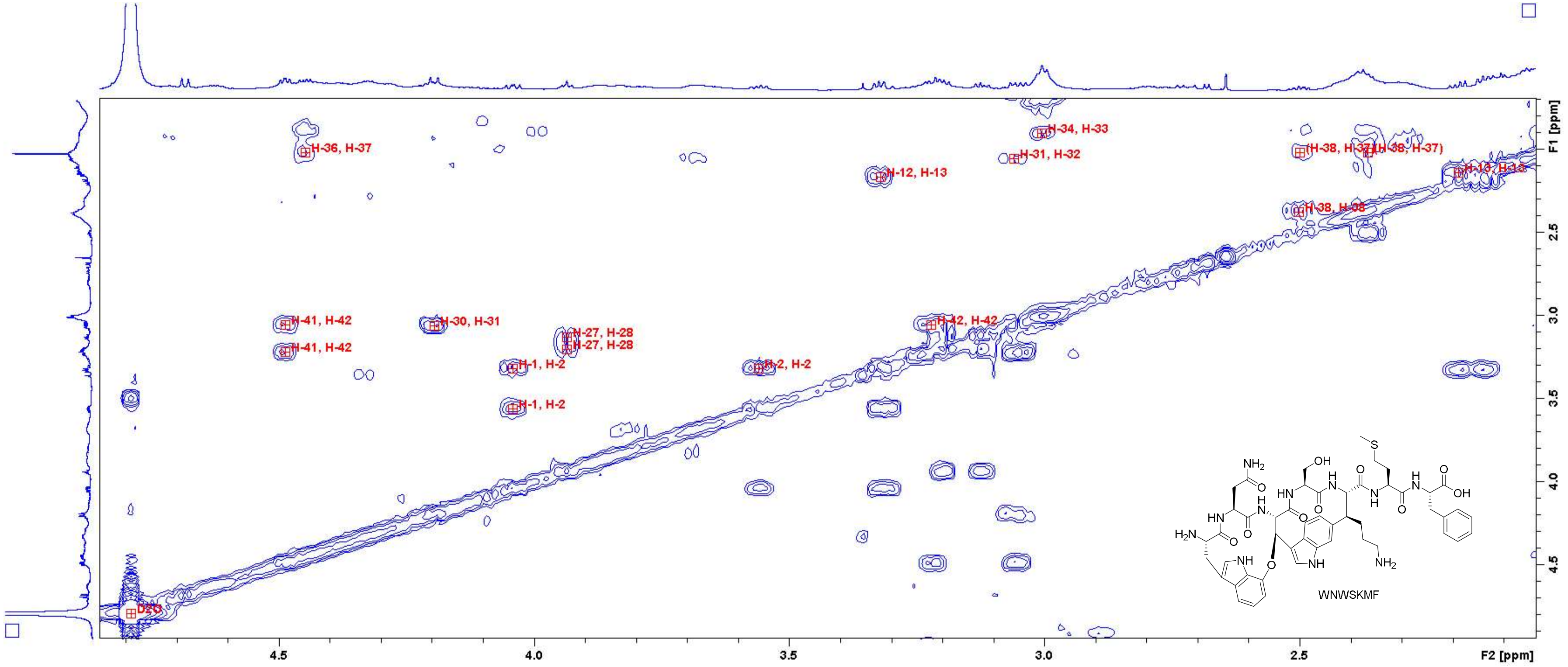

Figure S94: COSY spectrum of WNWSKMF in $D_2O$ (700 MHz). Close-up in the region of 2.1 – 4.8 ppm (F2 axis) and 1.7 – 4.9 ppm (F1 axis) with peak assignments. For atom numbering, cf. Figure S3.

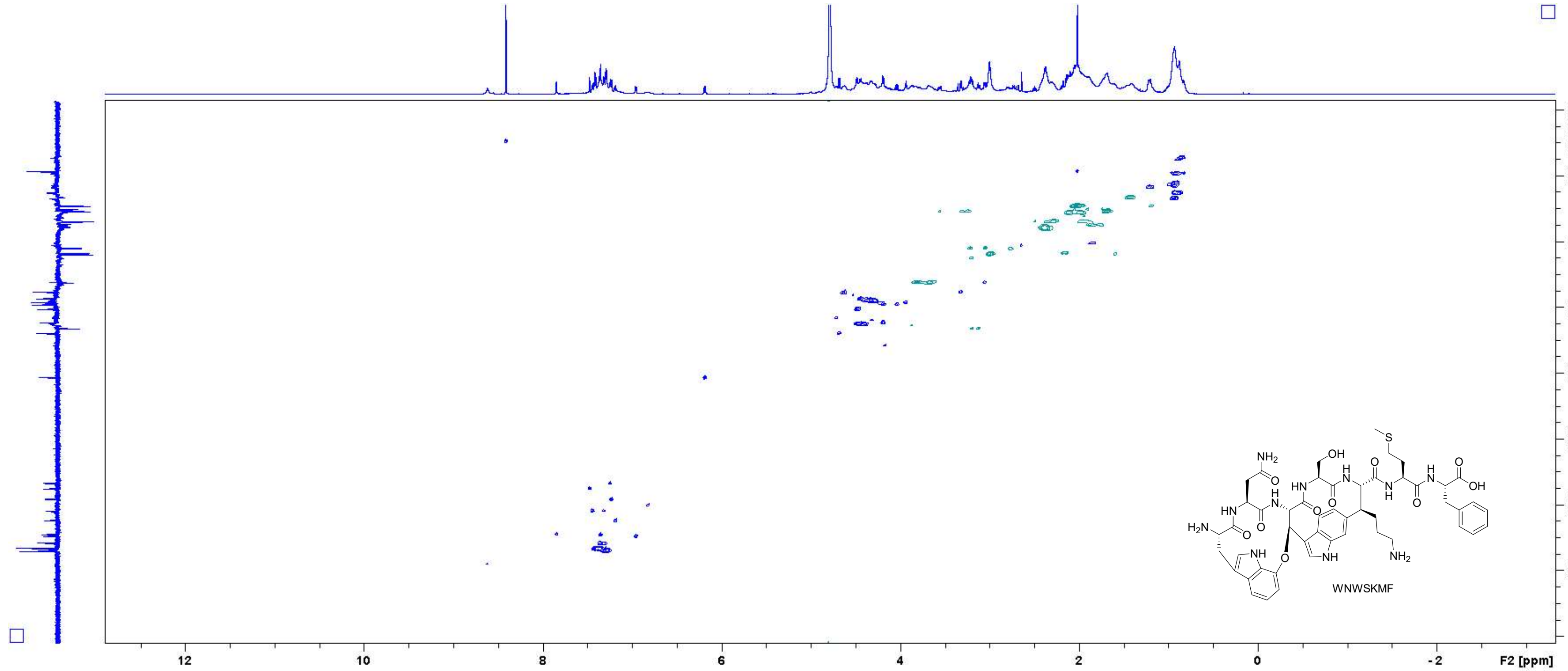

Figure S95: HSQC spectrum of WNWSKMF in $D_2O$ (700 MHz).

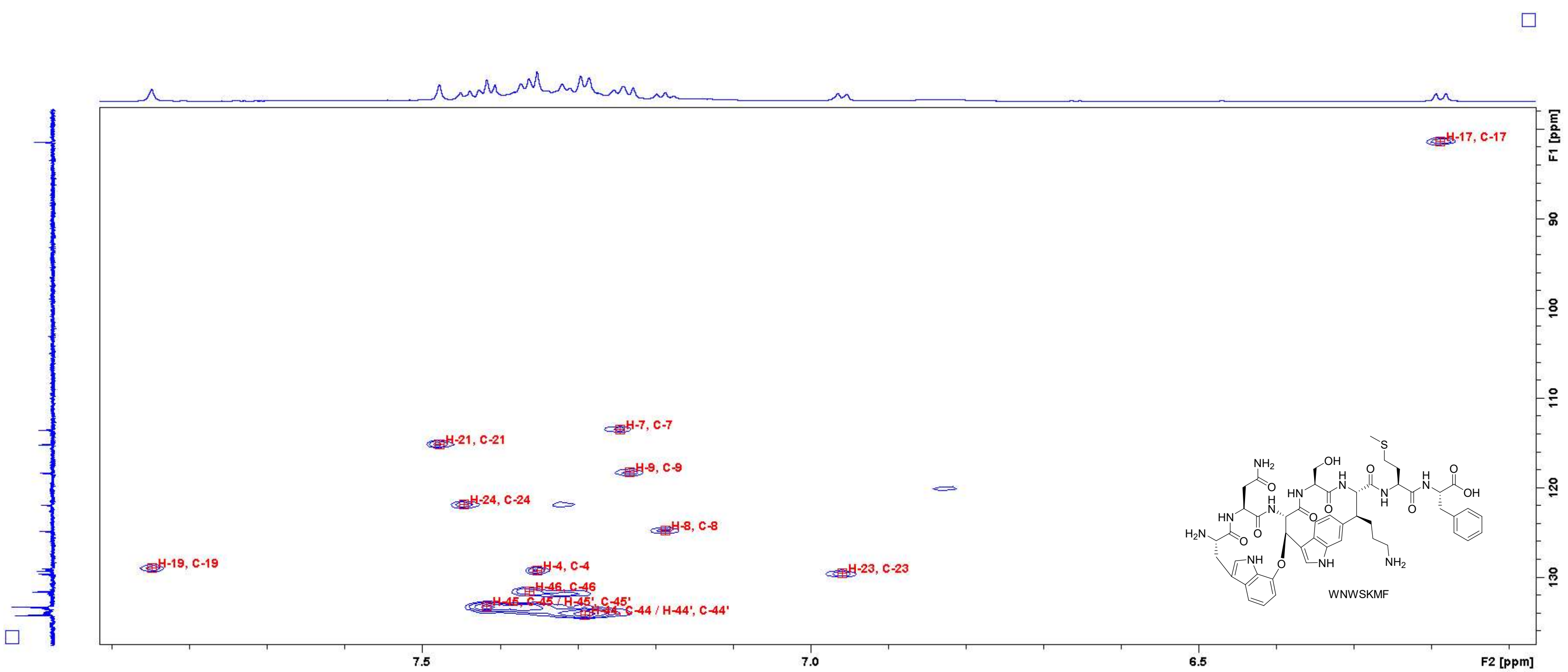

Figure S96: HSQC spectrum of WNWSKMF in $D_2O$ (700 MHz). Close-up in the region of 6.1 – 7.9 ppm (F2 axis) and 78 – 136 ppm (F1 axis) with peak assignments. For atom numbering, cf. Figure S3.

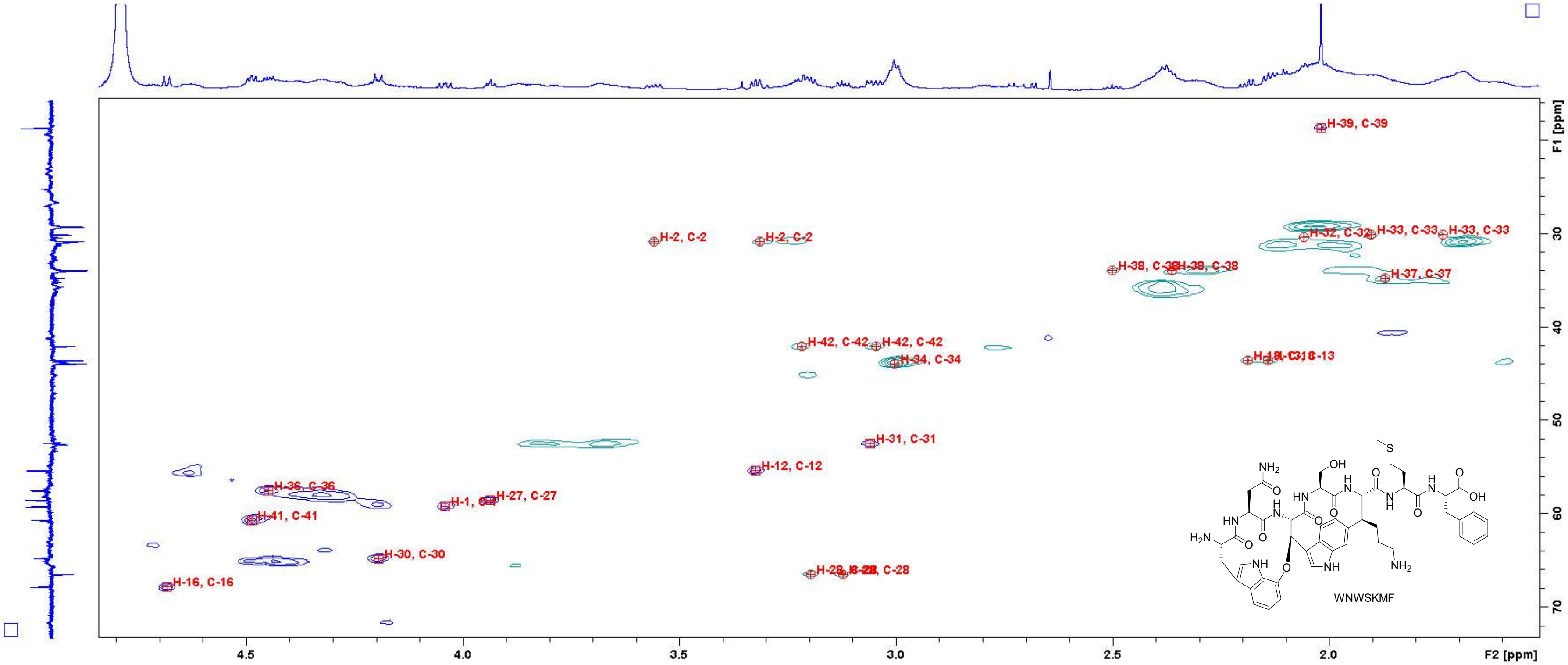

Figure S97: HSQC spectrum of WNWSKMF in D₂O (700 MHz). Close-up in the region of 1.6 – 4.8 ppm (F2 axis) and 16 – 72 ppm (F1 axis) with peak assignments. For atom numbering, cf. Figure S3.

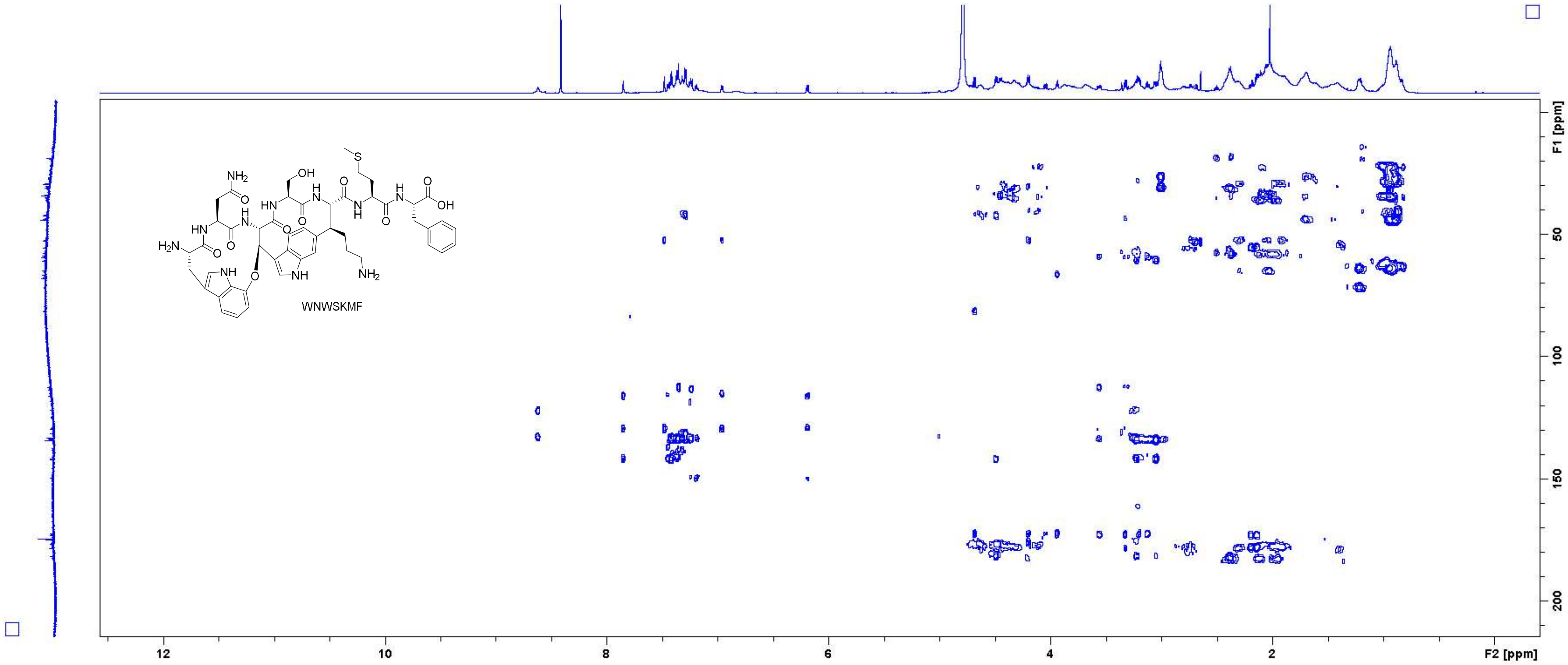

Figure S98: HMBC spectrum of **WNWSKMF** in $D_2O$ (700 MHz).

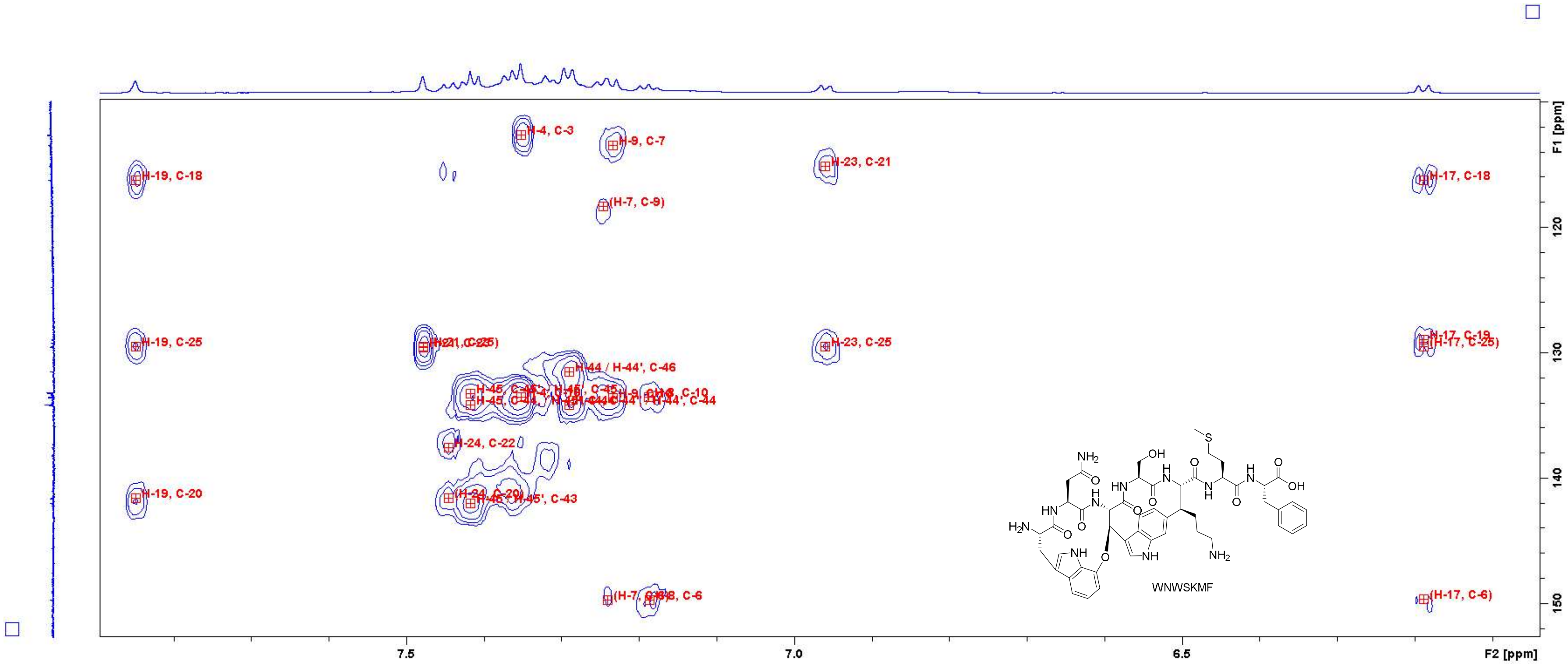

Figure S99: HMBC spectrum of WNWSKMF in D$_2$O (700 MHz). Close-up in the region of 6.1 – 7.8 ppm (F2 axis) and 110 – 152 ppm (F1 axis) with peak assignments. For atom numbering, cf. Figure S3.

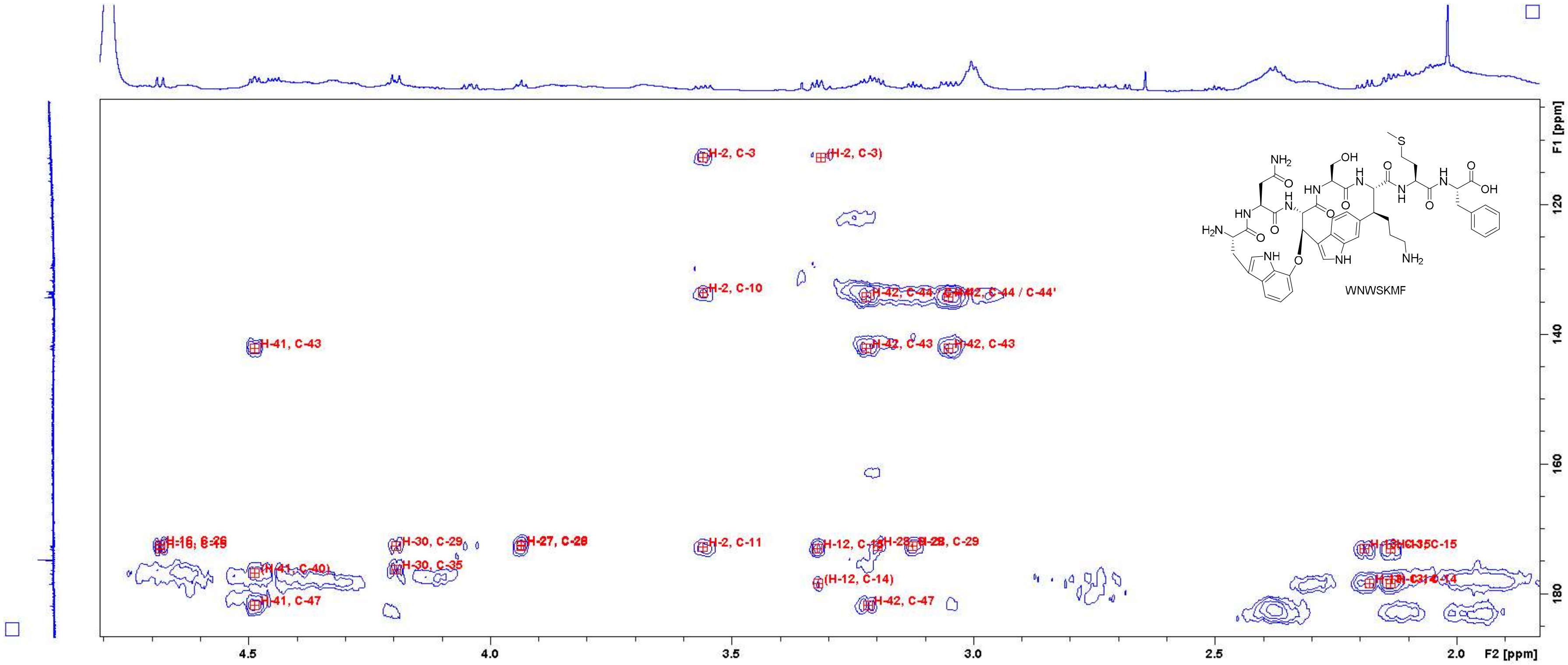

Figure S100: HMBC spectrum of WNWSKMF in D$_2$O (700 MHz). Close-up in the region of 1.9 – 4.8 ppm (F2 axis) and 105 – 185 ppm (F1 axis) with peak assignments. For atom numbering, cf. Figure S3.

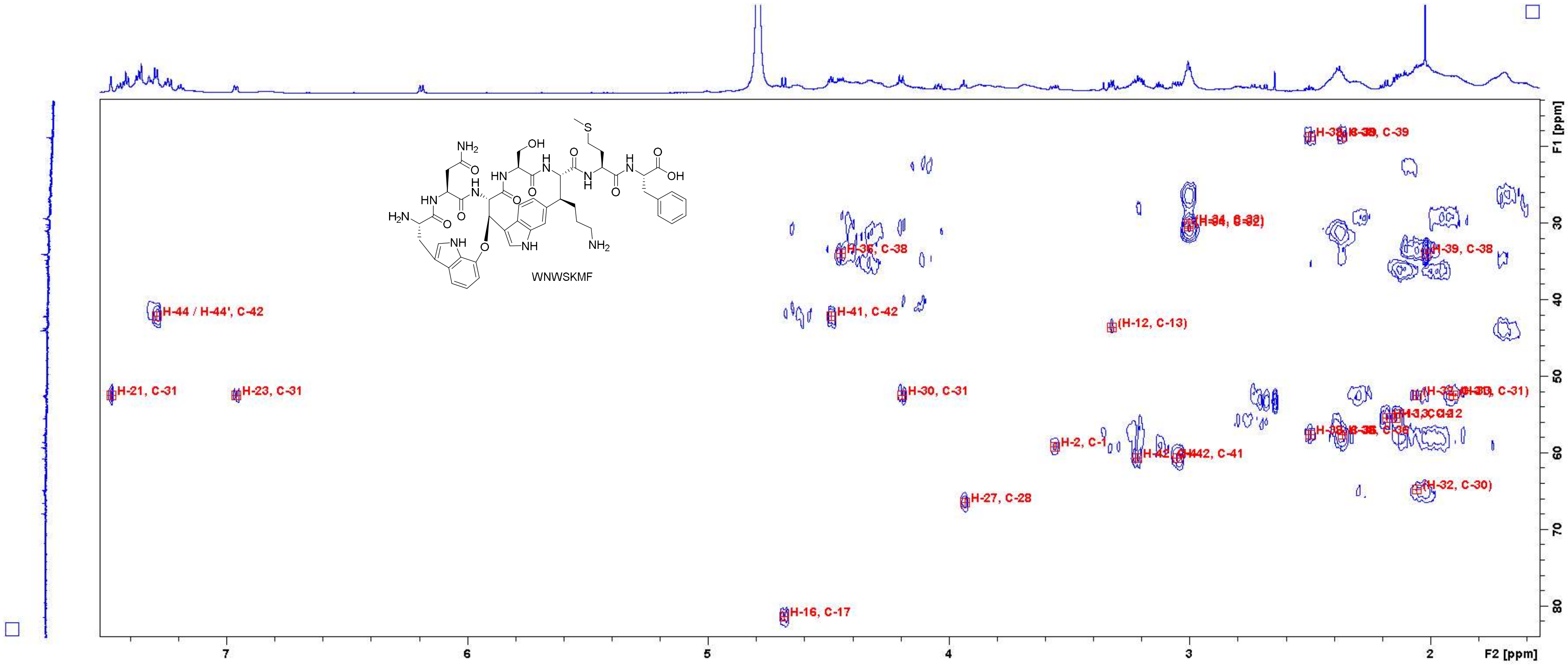

Figure S101: HMBC spectrum of WNWSKMF in $D_2O$ (700 MHz). Close-up in the region of 1.6 – 7.4 ppm (F2 axis) and 14 – 82 ppm (F1 axis) with peak assignments. For atom numbering, cf. Figure S3.

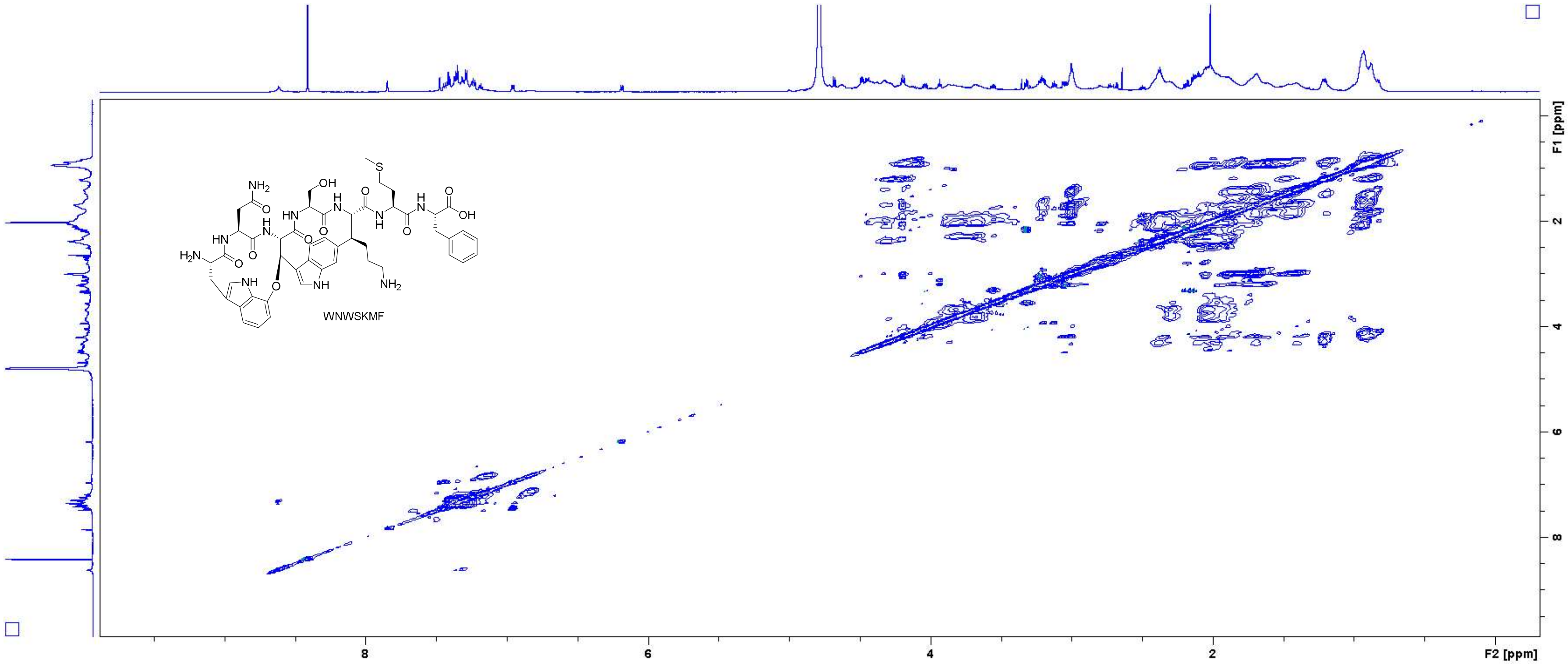

**Figure S102**: TOCSY spectrum of **WNWSKMF** in $D_2O$ (700 MHz, measured with $H_2O$ suppression).

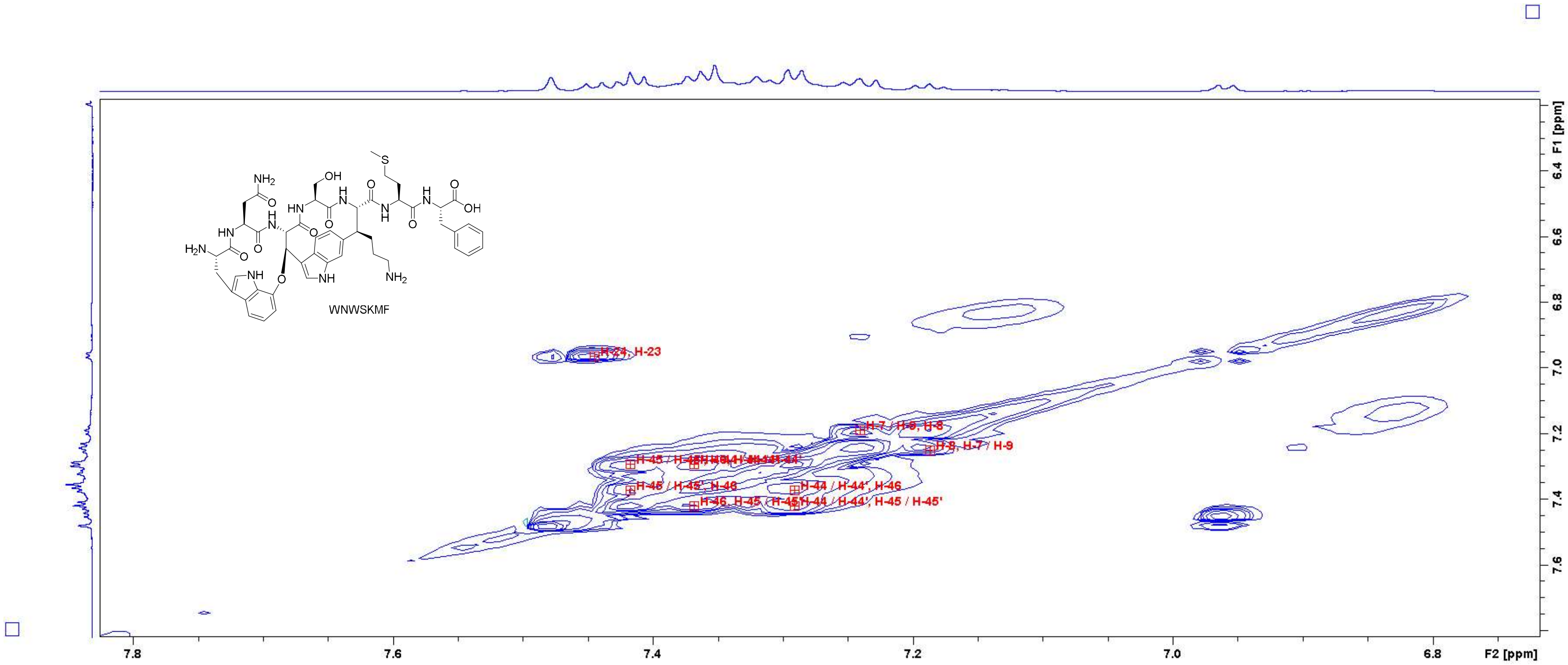

Figure S103: TOCSY spectrum of WNWSKMF in $D_2O$ (700 MHz, measured with $H_2O$ suppression). Close-up in the region of 6.75 – 7.80 ppm (F2 axis) and 6.2 – 7.8 ppm (F1 axis) with peak assignments. For atom numbering, cf. Figure S3.

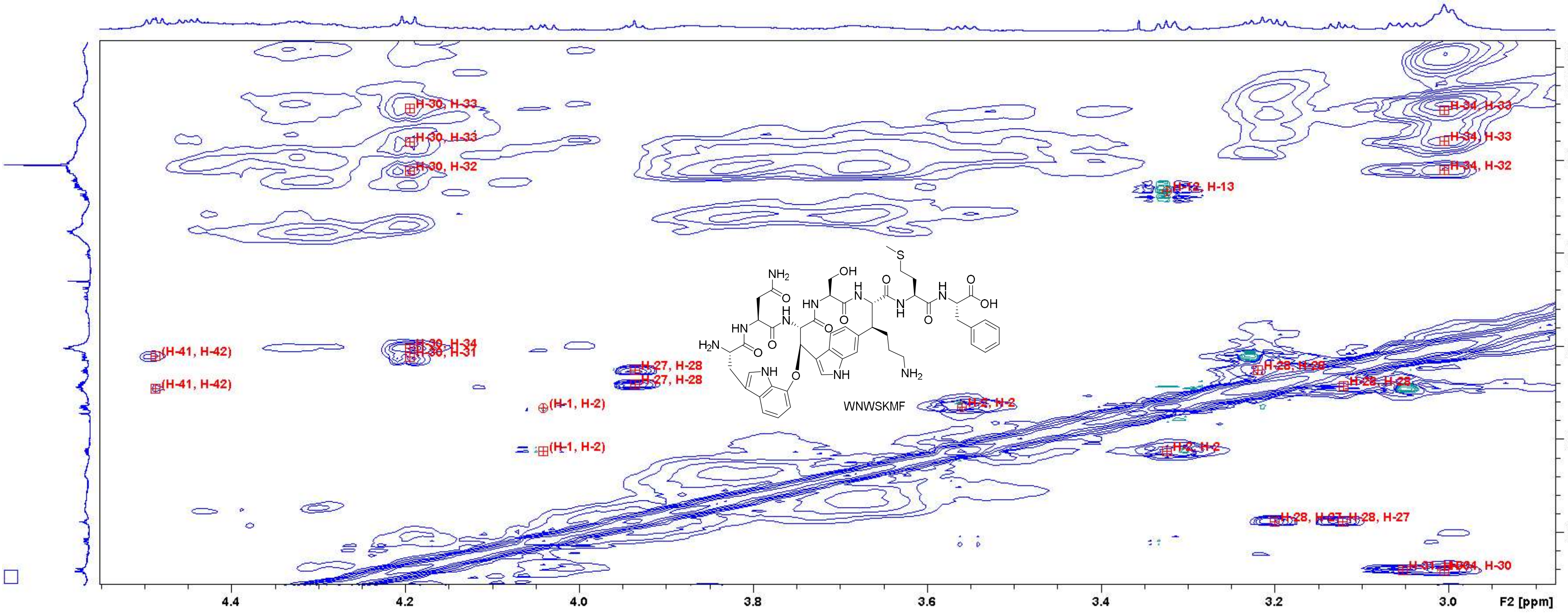

Figure S104: TOCSY spectrum of WNWSKMF in $D_2O$ (700 MHz, measured with $H_2O$ suppression). Close-up in the region of 2.90 – 4.55 ppm (F2 axis) and 1.4 – 4.2 ppm (F1 axis) with peak assignments. For atom numbering, cf. Figure S3.

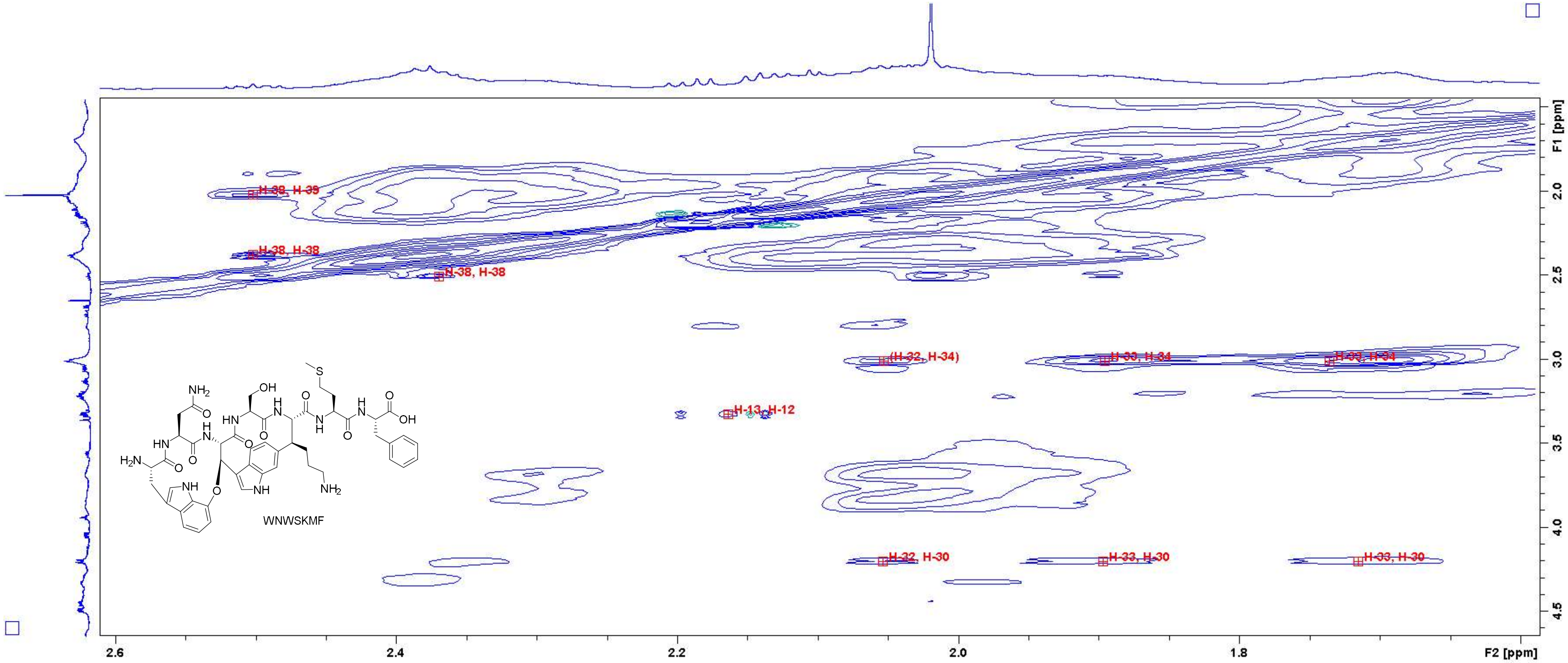

Figure S105: TOCSY spectrum of WNWSKMF in $D_2O$ (700 MHz, measured with $H_2O$ suppression). Close-up in the region of 1.6 – 2.6 ppm (F2 axis) and 1.5 – 4.6 ppm (F1 axis) with peak assignments. For atom numbering, cf. Figure S3.

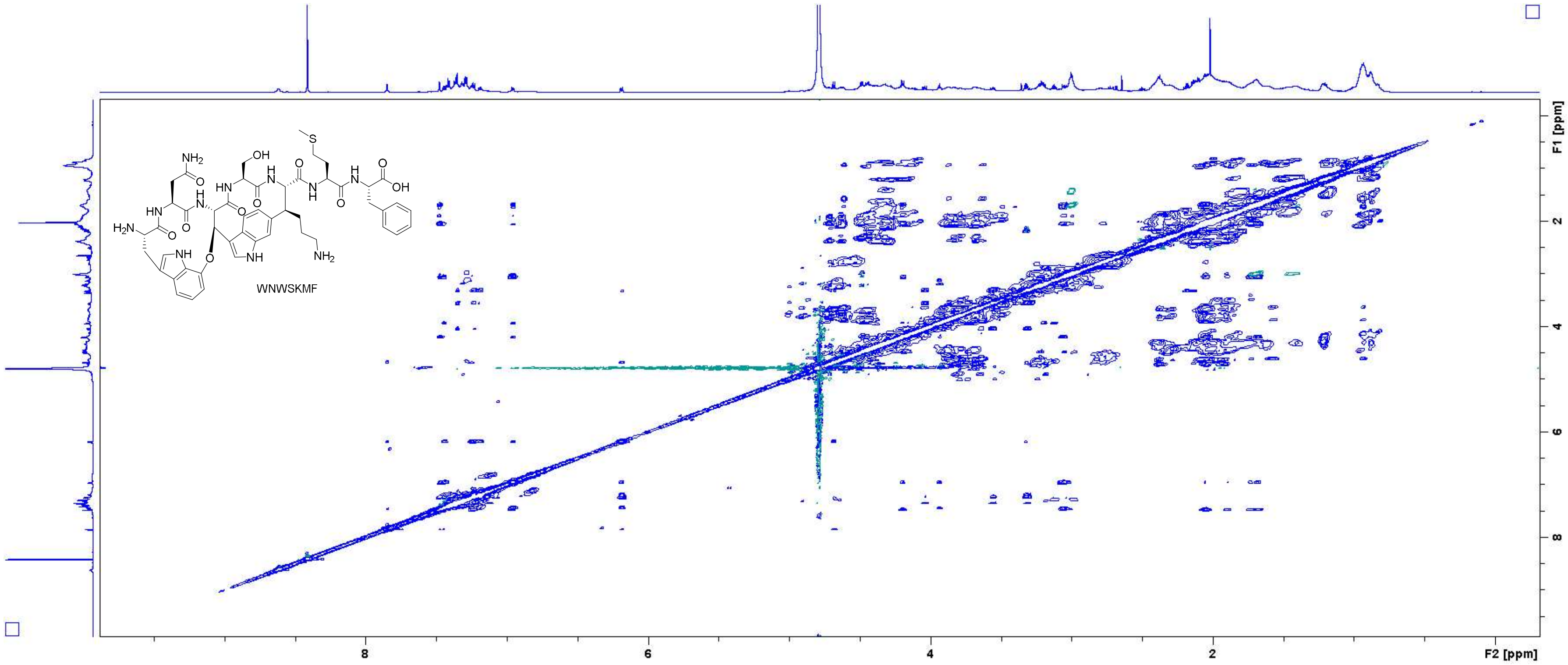

Figure S106: NOESY spectrum of WNWSKMF in $D_2O$ (700 MHz).

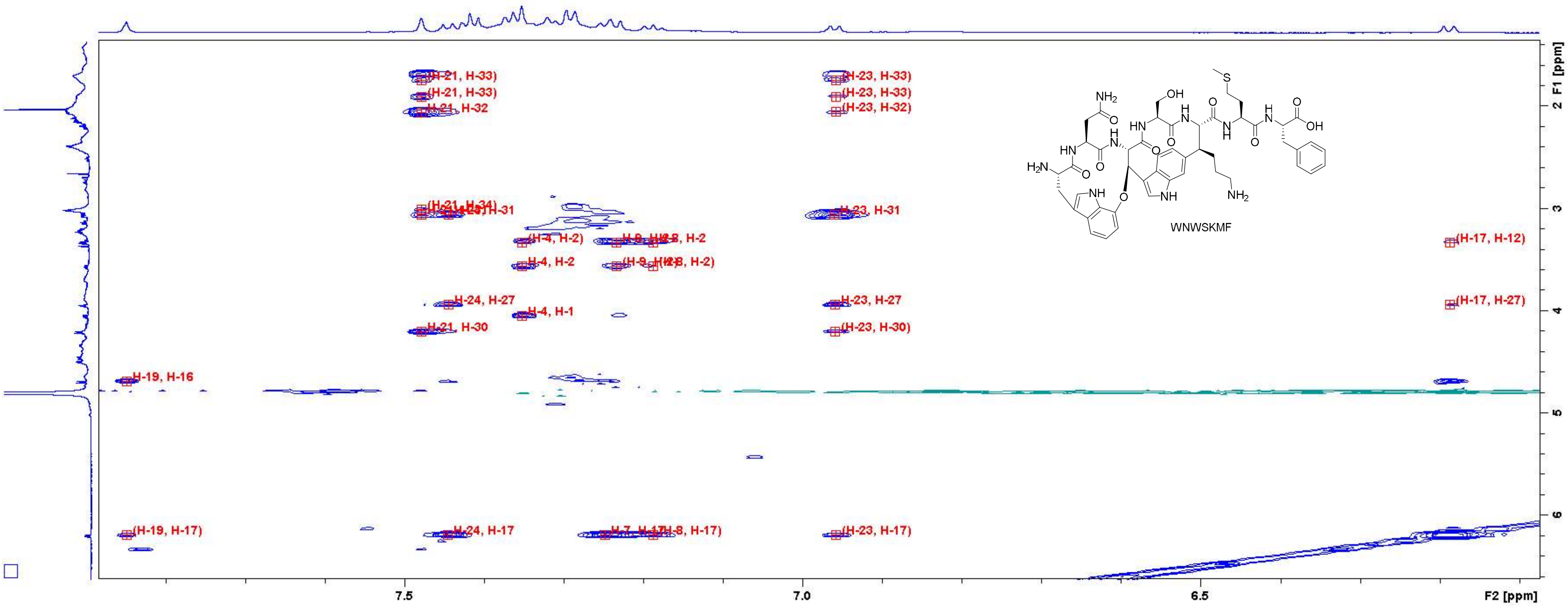

**Figure S107:** NOESY spectrum of **WNWSKMF** in $D_2O$ (700 MHz). Close-up in the region of 6.1 – 7.8 ppm (F2 axis) and 1.4 – 6.6 ppm (F1 axis) with peak assignments. For atom numbering, cf. Figure S3.

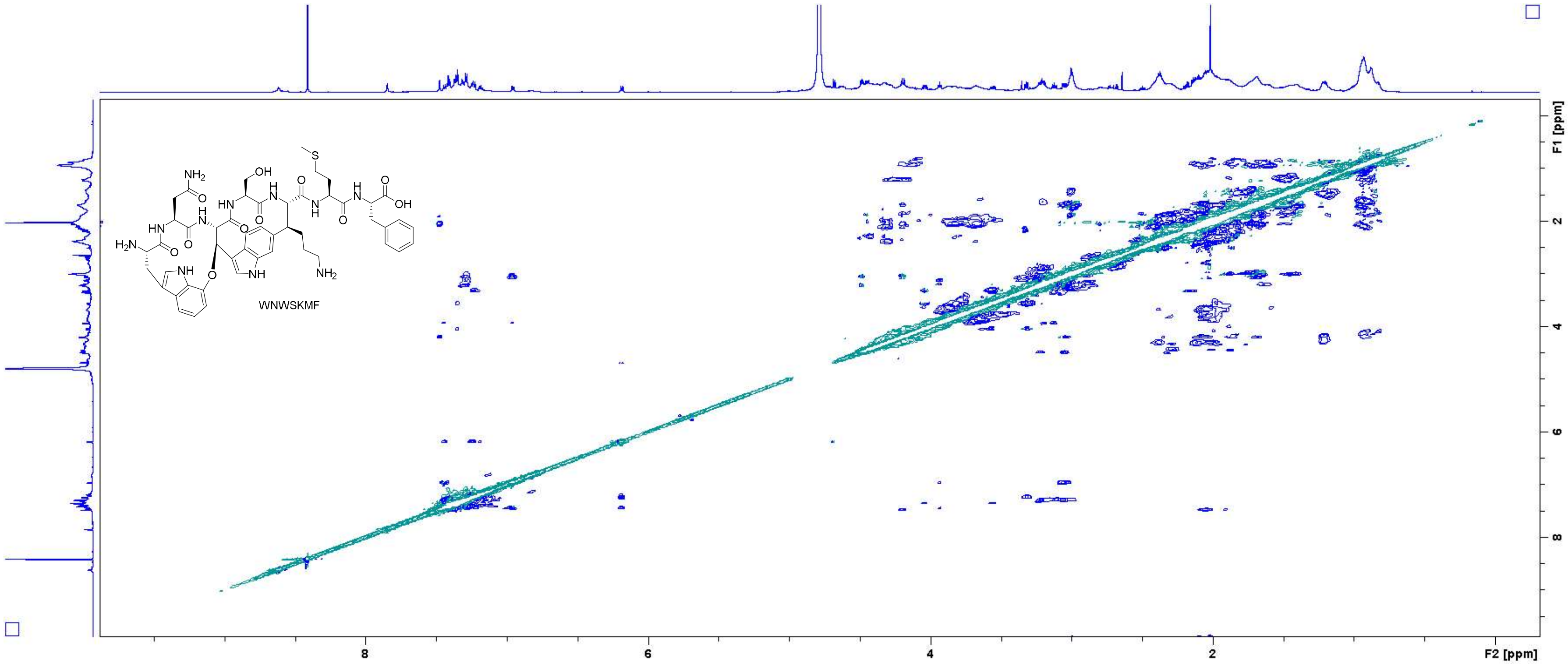

Figure S108: ROESY spectrum of WNWSKMF in $D_2O$ (700 MHz, measured with $H_2O$ suppression).

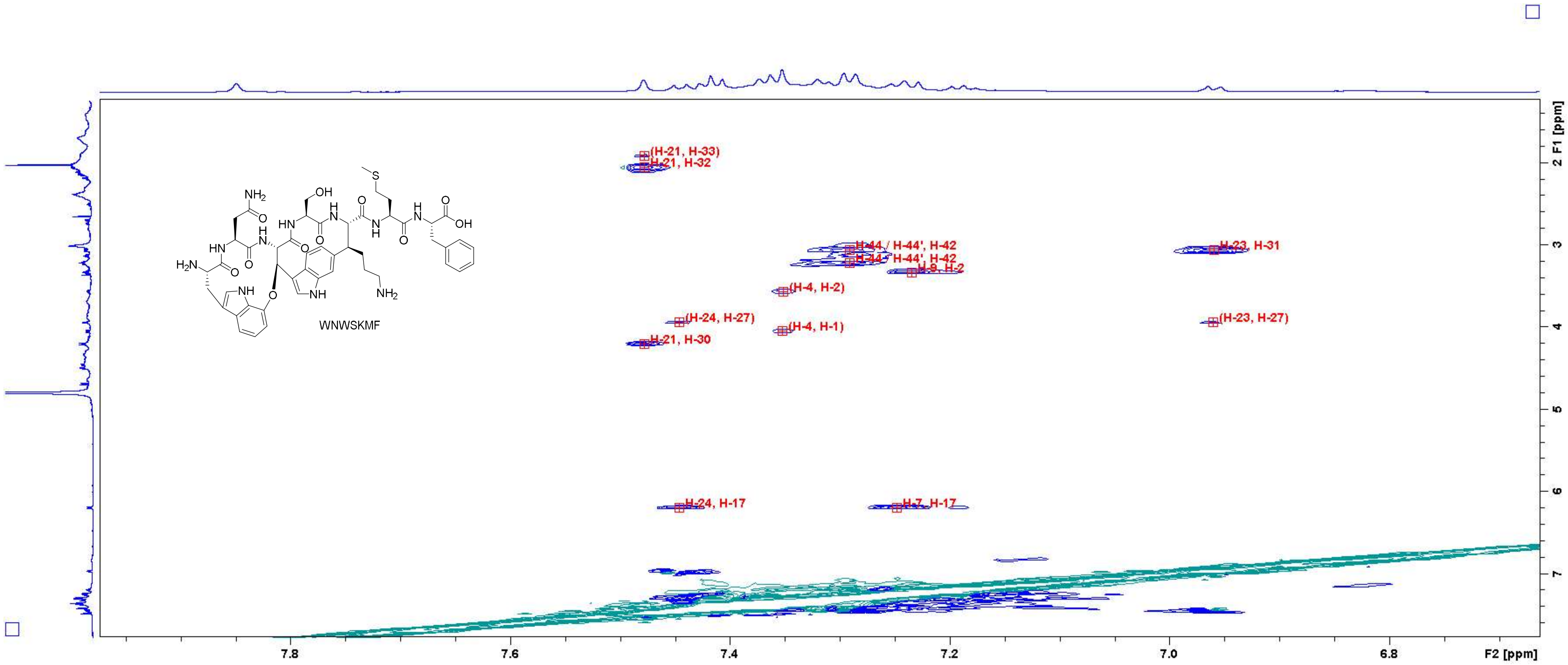

Figure S109: ROESY spectrum of WNWSKMF in $D_2O$ (700 MHz, measured with $H_2O$ suppression). Close-up in the region of 6.70 – 7.95 ppm (F2 axis) and 1.4 – 7.6 ppm (F1 axis) with peak assignments. For atom numbering, cf. Figure S3.
